# Allosteric Protein Chemical Shift Perturbations are Ubiquitous

**DOI:** 10.64898/2026.05.04.722792

**Authors:** Tiburon L. Benavides, Theresa A. Ramelot, Gaetano T. Montelione

## Abstract

While allosteric protein function has been appreciated for decades, the ubiquity of conformational shifts, particularly those distant from the interaction interface, has not been broadly characterized. For example, ligand binding frequently triggers allosteric effects far from the interaction interface, yet the prevalence of these conformational shifts underpinning protein function remain poorly documented. We systematically assessed the generality of allosteric effects as monitored by NMR Chemical Shift Perturbations (CSPs) distant from the interaction interface. In a set of 139 protein-protein complexes, a striking 74% of all significant CSPs are non-local to the binding site. Notably, more than 35% of significant CSPs outside the binding site occur in residues for which the shortest receptor-ligand interatomic distance is more than 10 Å. Every protein analyzed exhibits a significant fraction (> 8%) of CSPs distant from the binding site. This analysis across a large number of protein structures demonstrates and documents that structural plasticity is a ubiquitous and fundamental property of proteins.

**Significance Statement:** Studies of protein dynamics have had a profound impact on biology. Ruth Nussinov famously postulated that multiple protein conformations preexist in dynamic equilibrium, with interconversions that mediate function. While conformational flexibility has been characterized in many specific case studies, the extent to which structural plasticity can be considered a fundamental and ubiquitous property of proteins remains poorly documented. We address a central question: how common is protein structural plasticity? To do so, we compiled a database of protein-protein and protein-peptide complexes with NMR chemical shift data for both bound (holo) and unbound (apo) states. These data reveal the widespread prevalence of long-range structural perturbations induced by ligand binding, demonstrating that structural plasticity is a pervasive and fundamental property of proteins.

## Introduction

Studies of protein dynamics are having a significant impact on biology. The native structure of a protein is widely understood to be an ensemble of conformations that dynamically interconvert to provide the basis for its biochemical functions (1–3). The plasticity of protein structure has been observed in numerous experimental and theoretical studies of specific proteins. Dynamically-averaged backbone chemical shift predictions have also been shown to have better agreement with experimental values than predictions made from static X-ray structures (4). However, while multi-state properties and conformational flexibility have been characterized case-by-case for many individual proteins, the extent to which structural plasticity is a fundamental and ubiquitous property of all proteins is not well documented.

CSPs observed by NMR are a powerful and widely used probe of protein-ligand interactions. These perturbations report changes in local chemical environments of NMR active atomic nuclei upon ligand binding. CSPs are commonly measured using ^1^H, ^1^□N, ^13^C, and/or ^19^F chemical shifts recorded in NMR spectra of the apo (free) and holo (ligand-bound) states of the protein (5, 6). They are frequently and successfully employed in integrative structural modeling of complexes, particularly through information-driven docking tools like HADDOCK (7–9), ColabDock (10), or MELD-NMR (11, 12) that use CSP-derived ambiguous interaction restraints (AIRs) to define interface residues. Importantly, CSPs reflect not only direct interfacial contacts but also indirect conformational changes, including long-range allosteric effects propagated through the protein structure.

How general is protein structural plasticity? To assess how polypeptide ligand binding propagates through protein structures, we compiled a curated CSP Database (CSPdb) of protein-peptide interactions with backbone amide (^1^□N and ^1^H) chemical shifts determined for both bound (holo) and unbound (apo) states. In this set, 74% of all significant CSPs are non-local to the binding site. More than 35% of significant CSPs outside the binding site occur in residues for which the shortest receptor-ligand atomic distance exceeds 10 Å. Every protein in this set exhibits a significant fraction (> 8%) of CSPs distant from the binding site. These data demonstrate that structural plasticity is a pervasive and fundamental property of proteins. In addition, detailed residue-level CSP patterns and inferred allosteric sites for each complex are provided in the **Supplemental Information**, enabling receptor-specific analysis across the dataset.

## Results

Using the Biological Magnetic Resonance Bank (BMRB), we compiled a curated CSP database (CSPdb) of protein-protein and protein-peptide complexes with backbone amide (^1^□N and ^1^H) chemical shifts determined for both bound (holo) and unbound (apo) states. The initial data set of 260 protein pairs was filtered based on strict matching of solution conditions to 139 pairs. A subset of these [n=126] also include apo and holo ^13^Cα shifts. The complete CSPdb dataset is detailed in **SI Table S1** and available via GitHub (https://github.com/MontelioneLab/CSP_UBQ), with **Supplemental Information** providing residue-level CSP analysis and inferred allosteric perturbation sites for each protein complex.

The protein pairs included in CSPdb regulate diverse biological processes, including cell cycle progression, transcriptional control, translational regulation, autophagy, apoptosis, and cell adhesion. Disease-relevant targets address mechanisms in leukemia and other cancers, Alzheimer’s disease, and infectious diseases from viruses and bacteria, such as anthrax, chikungunya, SARS-CoV-2, HIV, and Ebola. Enzyme Commission (EC) classes for the receptors in the CSPdb include: 21 hydrolases (**SI Table S2, SI Figure S1**), 33 transferases (**SI Table S3, SI Figure S2**), 2 isomerases, 6 oxidoreductases, and 1 translocase. CSPdb receptors contain 21 all-α proteins (**SI Table S4, SI Figure S3**), 23 all-β proteins (**SI Table S5, SI Figure S4**), and 12 α+β proteins (**SI Table S6, SI Figure S5**). Additional highlights of the receptors included in CSPdb are summarized in **SI Text**.

We used the CSPdb to analyze the ubiquity of allosteric CSPs, calculated as weighted differences between chemical shifts observed in the bound and free forms of the receptor. Weighting factors followed standard conventions (6) to account for the different gyromagnetic ratios of ^1^H, ^15^N, and ^13^C (**Eqn. 1** of **Methods**). To correct for inconsistencies in spectral referencing in BMRB depositions, holo protein chemical shifts were globally adjusted relative to the apo state. The optimal offset was determined using a grid search that maximized the number of aligned peaks within < 0.05 ppm (as defined in **Methods**) between the apo and holo HSQC values (**Figure 1A**). This correction effectively minimizes artificial shift differences in disordered segments distant from the ligand binding site and other unperturbed regions due to inaccurate global referencing.

**Fig. 1.**
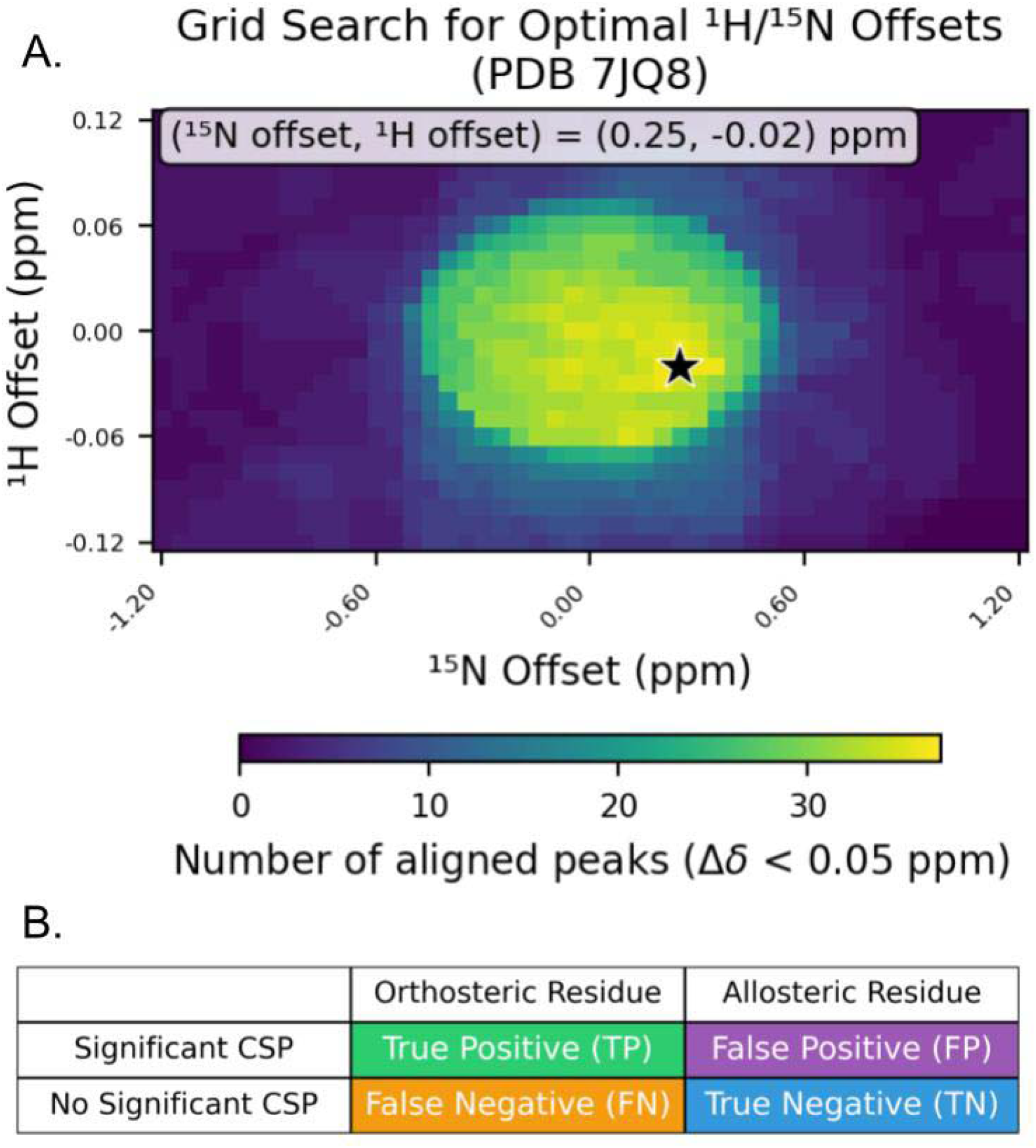
(A) Matching non-perturbed chemical shifts of apo and holo protein data. CSPs were calculated by comparing chemical shifts between bound and free forms of the receptor. In each case, apo and holo chemical shifts were referenced by applying offsets to the holo shifts. The optimal offset was determined by a grid search over possible ^1^H and ^1^□N (or ^13^C) offsets, selecting the values that maximized the number of aligned peaks between apo and holo HSQC peak lists, as described in **Methods**. Shown here is a representative grid search heat map for optimal N/H holo shift offsets for the ET-TP system (PDB ID 7JQ8). CSPs calculations follow previously published methods (6) (**Eqn. 1, Eqn. 2**). Offsets for all targets in the CSPdb are summarized in (**SI Figure S16**). (B). Confusion matrix used to assess CSPs. Orthosteric Residues are those within the binding site, Allosteric Residues are those outside of the binding site. See **Methods** for definition of binding site (orthosteric) residues.

We define a binary classification task using heuristic thresholds to define “significant” CSPs and structural definitions of binding site residues (see **Methods** for details) to construct a confusion matrix (**Figure 1B**), where significant CSPs are treated as predicted binding-site residues. Some specific case studies from this analysis are shown in **Figure 2**. Similar analyses are provided for each of the 139 complexes in the **SI**, effectively mapping receptor-specific allosteric CSP patterns across the CSPdb. In these molecular graphics images, residues with significant CSPs that are not in the ligand binding site (FPs) are colored purple. Aggregated counts for each quadrant of the confusion matrix across the entire CSP dataset are summarized in **Figure 3**. Overall, a striking 71% of significant CSPs (FP / (TP+FP)) occur outside the binding site. Among false positives (significant CSPs of residues not in the binding site), 36% are for residues with inter-chain Cα-Cα distances > 15 Å (**Figure 1A**), and 38% are for residues with minimum receptor-ligand interatomic distances (including sidechain - sidechain contacts) > 10 Å (**SI Figure S13**). The average FP rate across the CSPdb (FP / (FP+TP+FN+TN)) is 30%, and every protein in the dataset exhibits a substantial fraction of FP CSPs (minimum 8%) demonstrating that long-range ligand-induced perturbations of protein structure are ubiquitous.

**Fig. 2.**
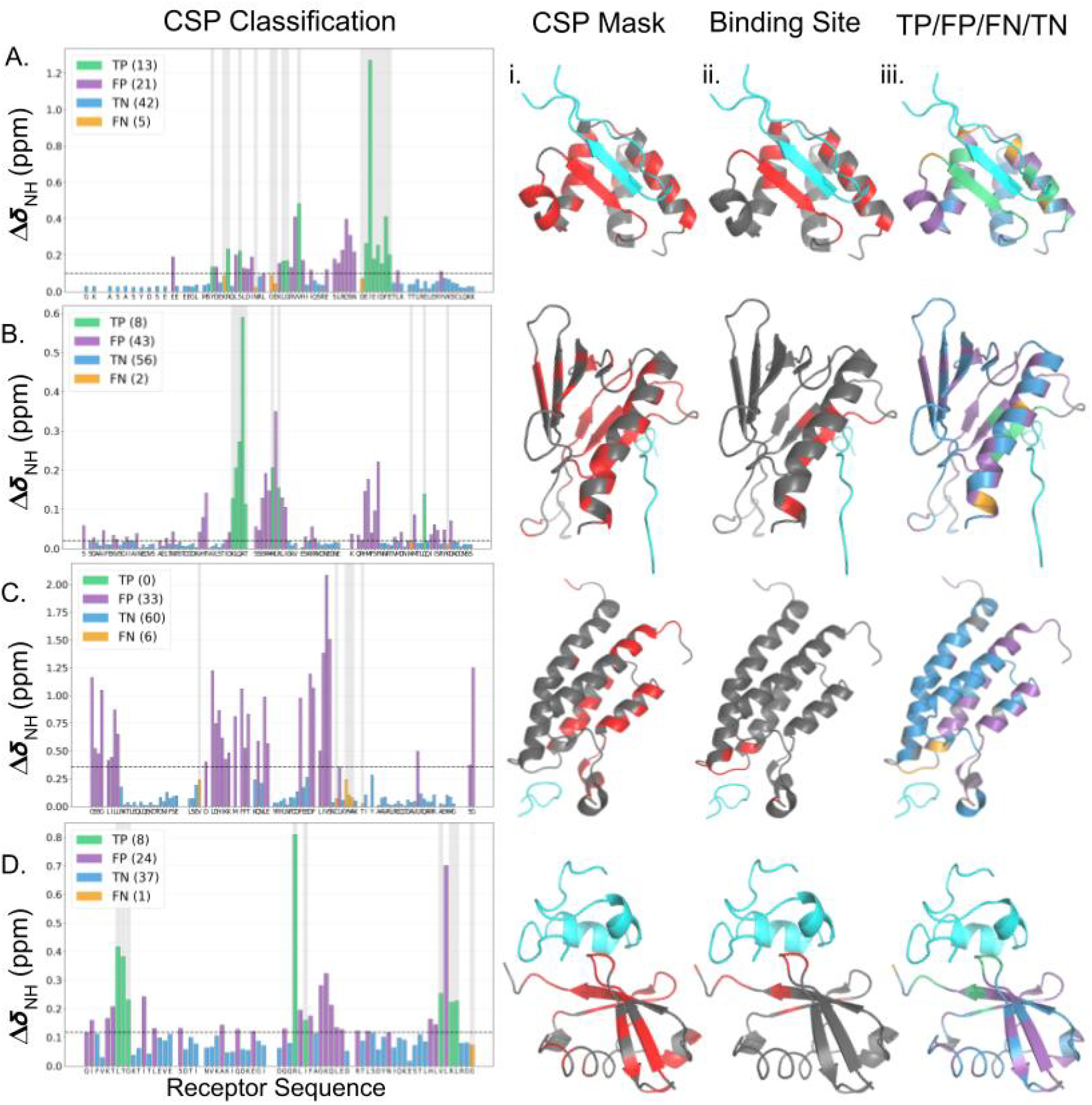
Representative case studies. (Left) Calculated CSPs for receptor residues along the sequence, bars colored by confusion matrix classification (see Fig. 1 for color key). Dashed horizontal lines indicate the significance threshold (Methods), and gray-shading highlights binding-site residues. Molecular graphics panels (left to right) show PyMOL renderings of receptor structures colored by (i) the mask of significant CSPs, with significant CSPs in red and residues without CSP measurements or insignificant CSPs shown in gray (ii) the binding-site residues (defined in Methods) in red and residues outside of the binding site in gray, and (iii) the confusion matrix classification projected onto the structure; ligands are colored cyan. Residues with allosteric CSPs are indicated in purple, and generally have patterns that radiate from the ligand binding site. (A) holo PDB 7JQ8; apo BMRB 30782; holo BMRB 30786. (B) holo PDB 2M14; apo BMRB 6225; holo BMRB 18842. (C) holo PDB 2RS9; apo BMRB 19125; holo BMRB 11463. (D) holo PDB 2KWV; apo BMRB 17769; holo BMRB 16885.

**Fig. 3.**
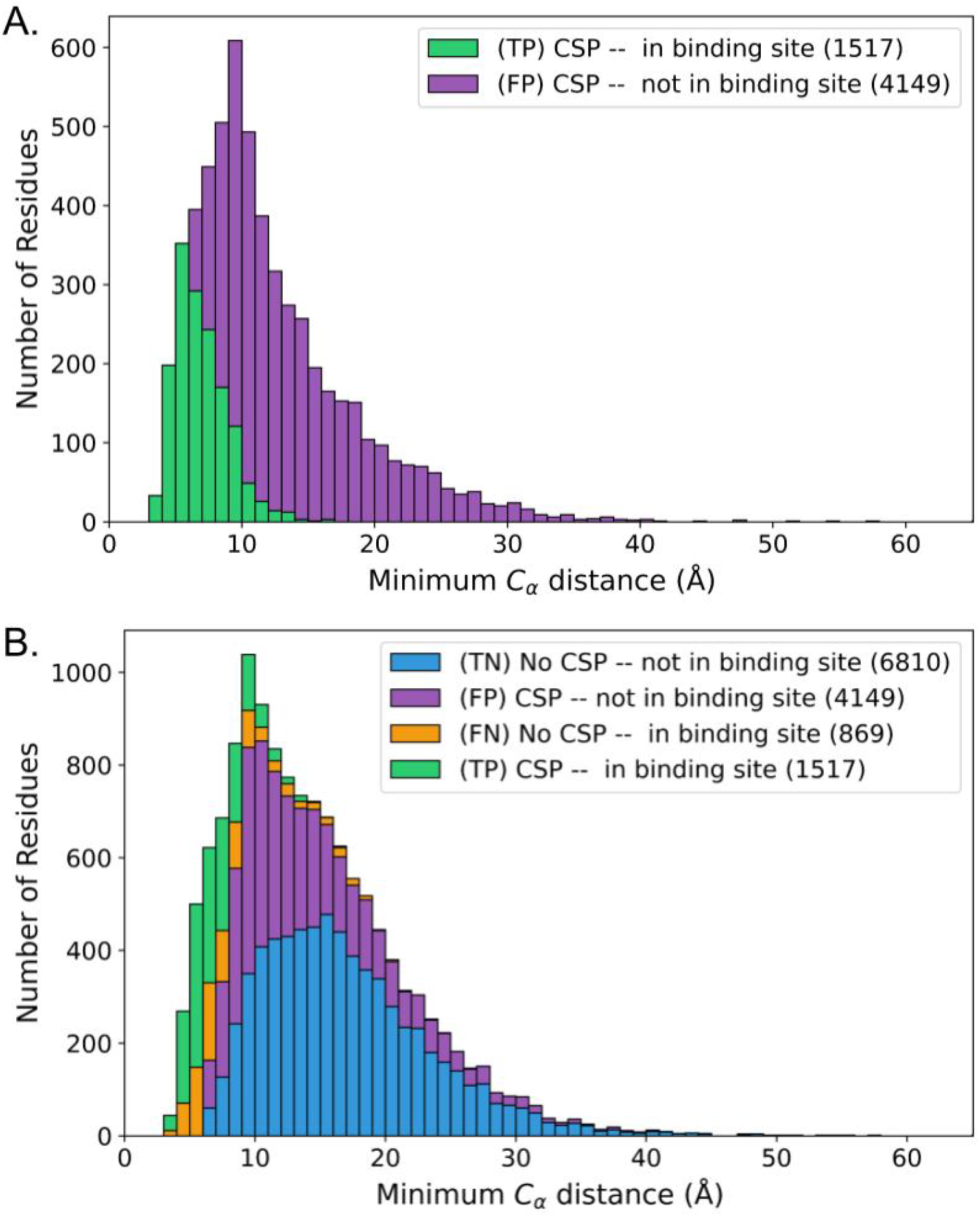
Stacked histograms summarizing the confusion matrix across the CSPdb. In both panels, the x-axis shows the minimum interchain Cα-Cα distance for residues in the receptor chain. **(A) Significant CSPs**. Green bars: residues within the binding site (TPs); purple bars: residues outside the binding site (FPs). Overall, 74% of significant CSPs occur outside the binding site. **(B) All residues**. In addition to the significant CSPs shown in (A), residues with no (significant) CSPs are included: blue bars: residues outside the binding site (TNs); orange bars: residues within the binding site (FNs). Calculating CSPs using Cα shifts yields similar results (**SI Figures S10, S11**). Similar plots using minimum interchain N-N distances and minimum interchain atomic distances are shown in **SI Figures S13, S14**. CSP are defined as described in **Methods**; binding-site residues are defined in **Methods**. Colors used throughout the manuscript correspond to the confusion matrix classification: TP (green), FP (purple), TN (blue), and FN (orange); aggregate counts per quadrant are in parentheses.

Although CSPs are often determined using only ^15^N-^1^H HSQC data, they can also be assessed using other chemical shift data, including ^13^C chemical shift data. For 126 of the 139 targets with available ^13^Cα apo and holo chemical shift data, calculating CSPs using **Eqn. 2** yields a similar proportion (72%; FP / (TP+FP)) of significant CSPs occur outside the ligand binding site (**SI Figure S10, SI Figure S11**). Backbone ^1^H^N, 15^N, and ^13^C^α^ resonances were each observed to be about equally sensitive to ligand-induced CSPs (**SI Figure S14**).

## Discussion

This study demonstrates, across a large and diverse set of NMR-characterized apo/holo protein-polypeptide complexes, the ubiquity of long-range perturbations due to ligand binding. It establishes the generality of binding-induced shifts in the conformational energy landscape and the fundamental *structural plasticity* of proteins. The non-local CSPs observed in our comprehensive analysis of 139 protein complexes align with extensive case-specific literature documenting allosteric networks and indirect effects in protein-ligand interactions.

As described in more detail in **SI Text**, numerous well-characterized systems in the CSPdb demonstrate mechanistically characterized long-range CSPs, often propagating through identifiable pathways or attributable to shifting dynamic ensembles. For example, calmodulin undergoes global reorganization upon peptide binding, resulting in widespread CSPs (13). SH3 domains show differential dynamics in peptide complexes (14), and the MAD2 spindle checkpoint protein features a substantial folding-upon-binding event to stabilize the formation of an inter-chain beta sheet. The findings highlighted here also corroborate results from our own studies of interactions with the extraterminal (ET) domain of BET proteins, which involve allosteric changes in loop stability, disorder to order transitions, binding-induced formation of secondary structure, and helix rearrangements upon complex formation (11, 15). Non-local structural changes accompanying complex formation have also been frequently reported in specific systems by amide hydrogen exchange measurements, and both increased and decreased rates of amide exchange at amide sites distant from the binding site have been reported (16–18).

Distant CSPs may arise from diverse phenomena, precluding exhaustive enumeration. Common mechanisms of signal propagation captured by receptors in the CSPdb include:

- Shifts in equilibria between multiple or alternative protein conformational states, leading to changes in ensemble-averaged chemical shifts (3).
- Binding-induced changes in conformational dynamics, such as folding-upon-binding or disorder-to-order transitions common in peptide recognition, propagating changes along the backbone and/or side chains (19).
- Rearrangement of hydrogen-bond networks, disrupting or forming extended hydrogen-bond chains that influence chemical environments far from the interface (20).
- Changes in rigidity, altering the average chemical environment at remote sites (15).
- Changes in helix dipole moments or electric fields affecting distant atomic resonance frequencies.

Despite the pervasiveness of CSPs distant from the ligand binding site and the observation that these often involve residues that are buried within the protein structure, in some cases other mechanisms may contribute to the observed chemical shift changes, including unrecognized secondary ligand-binding sites or ligand-induced changes in dimerization or oligomeric states. Examining the distributions of FP CSPs through 3D protein structures in the 139 case studies compiled in the **SI** reveals that in most cases these patterns radiate from the ligand binding site, consistent with binding-induced structural perturbations.

The structural amplitudes of ligand-induced non-local structural changes cannot be determined from CSP data alone. Even very small changes in structure (or the distribution of structures) can have significant effects on chemical shifts. Hence, while non-local CSPs provide evidence for conformational shifts and/or structural communication across the protein structure, the associated structural changes may be quite modest. Nonetheless, even small structural changes propagated across the protein structure can be functionally relevant. System-specific studies are required to assess if non-local CSPs have functional allosteric effects.

Characterization of the patterns of ligand-induced CSP-encoded structural changes can provide important insights into mechanisms of allosteric regulation (21). For example, CHEmical Shift Covariance Analysis (CHESCA), by exploiting correlated CSPs across a series of targeted perturbations (ligand analogs or mutations), can be used to map extended allosteric networks (22–24). By identifying clusters of residues exhibiting concerted chemical shift responses, CHESCA has revealed pathways of long-range communication in specific apo-holo protein pair comparisons (25, 26). In this context, sufficiently accurate prediction of chemical shifts from atomic structures needed to model allosteric structural changes consistent with the CSP data remains an important unsolved challenge.

Given the current state of development of AI models for modeling multiple conformations of proteins and for polypeptide and protein design, the results of this study are timely. These indirect, non-local effects on protein structure induced by polypeptide ligand binding remain difficult to predict and currently require experimental characterization. They reveal the intrinsically spongy nature of protein structures. The CSPdb provides a valuable resource for training machine learning models aimed at predicting non-local effects of ligand binding on protein structure and dynamics. Future work using CSPs together with structural ensemble modeling to characterize allosteric effects will also inform the development of allosteric modulators, a rapidly growing area of drug discovery.

## Materials and Methods

### Data collection

Targets were collected by manual curation from the PDB to identify targets annotated as two-chain heterodimeric solution NMR structures (PDB search settings configured as indicated in **SI Figure S15**). Targets were included based on the criteria that a receptor chain with identical sequence in the apo and holo states was deposited in the BMRB, and that a valid link existed between the holo PDB deposition and the holo BMRB deposition. As an additional criterion, the experimental conditions used in determining NMR assignments for the apo and holo protein (receptor) were required to be similar in pH (± 0.5 pH units) and temperature (± 5 °C). Differences in experimental conditions are known to cause systematic chemical shift changes not necessarily caused by ligand binding. Similar results for smaller numbers of target pairs were obtained using even tighter matching criteria. An additional 121 target pairs with experimental conditions outside of these matching thresholds were identified from the PDB and not included in the primary analysis of 139 pairs. We report summary statistics on this subset in **SI Table S10, SI Figure S9**.

### CSP calculation

CSPs were calculated by comparing chemical shifts between bound and free forms of the receptor using previously published methods (6) (**Eqn. 1, Eqn. 2**). Prior to CSP calculation, an offset was applied to holo chemical shifts to maximize the number of aligned peaks between the two spectra with 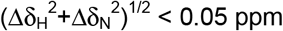. This offset is determined by a grid search over all possible ^1^H and ^1^□N offsets as outlined in **Figure 1**. ^1^H and ^1^□N holo shift offsets for all receptors in the CSPdb are summarized in **SI Figure S16**. An optimal offset for holo chemical shifts used in **Eqn. 2** was determined using a similar 3D grid search to maximize matched peaks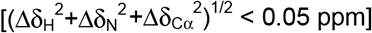. This alignment algorithm has the effect of matching chemical shifts of intrinsically disordered regions distant from the ligand binding site between the apo and holo data sets.

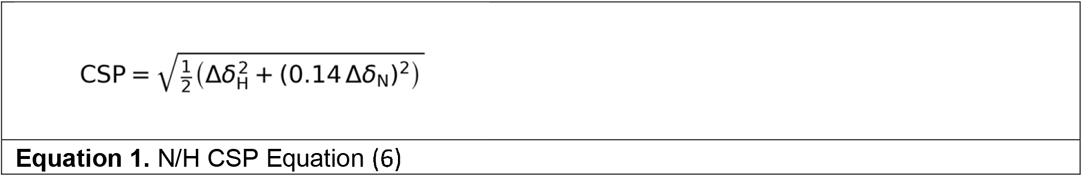

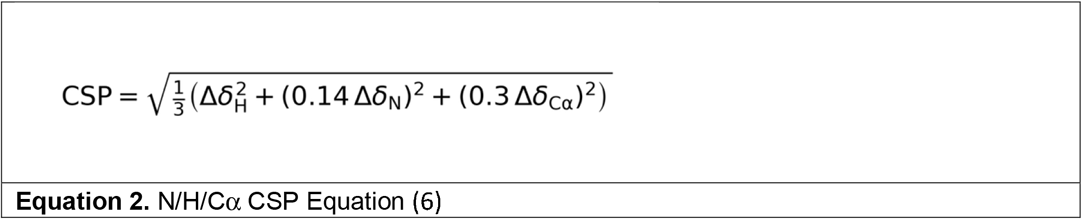

### Defining Significant CSPs

This method follows previously described protocols (27) Significance thresholds were determined as follows:

1. Calculate the list of chemical shift perturbations (28, 5, 6)
2. Iteratively remove outlier CSPs with a Z-score greater than 3 (σ > 3) until a stable subset with no outliers is obtained.
3. The final cutoff is defined as the mean CSP value of this outlier-free subset.

A histogram summarizing all CSP thresholds calculated across the dataset is available in **SI Figure S17**. The average significance cutoff across the CSPdb was 0.080 ppm. This is somewhat higher than the typical reproducibility of chemical shifts (∼0.020 ppm), likely reflecting small differences in conditions used for apo and holo NMR studies. Lowering these thresholds would identify more long-range CSPs, suggesting that our selection provides a conservative estimate of their extent.

### Defining the Binding Site

A residue is considered part of the binding site (i.e. orthosteric) if it satisfies any one of these criteria:

- has any interchain atom-atom distance < 2.0 Å.
- has interchain Cα-Cα distance < 6.0 Å.
- has Solvent Accessible Surface Area (SASA) occlusion, defined as a non-negligible change in SASA calculated using the Shrake–Rupley algorithm on the medoid holo model with and without the ligand atoms. Residues with a change in SASA for backbone N and H atoms are flagged as occluded.
- has hydrogen bonds with the ligand detected by using the Baker–Hubbard method (θ > 120°, H…A < 2.5 Å).
- has charge complementarity with the peptide (opposite charged groups within 4.5 Å).
- has *π* contacts (NH-to-aromatic distances within 6 Å).

### Binary Classification

We use the preceding definitions of Significant CSPs and the residues in the receptor’s binding site to define a confusion matrix from which F1 and MCC statistics were calculated (**Figure 1B**). A comparison of F1 and MCC across targets in the CSPdb is shown in **SI Figure S18**.

## Supporting information

Supplementary Information 2 - All cases

## Acknowledgments

We thank Dr. A. Bax, K. Fraga, Y.J. Huang, A. Perez, G.V.T. Swapna, and R. Tejero for helpful discussions, and S. Collen for computational systems support. We also acknowledge access to the RPI Center for Computational Innovations (CCI) computing infrastructure. This work was supported by NIH NIGMS grant (R35 GM141818 to G.T.M.) and by the RPI Bio-computing and Bio-informatics Constellation Chair Fund. T.L.B. was supported by the NIH NIGMS Biomolecular Science Engineering Training Program (T32 GM141865) and the NIH R25 RAFAELS Training Grant (R25AG088409).

## Code and Data Availability

Software used in this publication are available at GitHub (https://github.com/MontelioneLab/CSP_UBQ) and Zenodo (https://github.com/MontelioneLab/CSP_UBQ; https://doi.org/10.5281/zenodo.19334615) and the CSPdb is archived at Zenodo (https://doi.org/10.5281/zenodo.19821104)

## Notes

https://github.com/MontelioneLab/CSP_UBQ

https://doi.org/10.5281/zenodo.19334615

https://doi.org/10.5281/zenodo.19821104

### Competing Interest Statement

G.T.M. is a founder of Nexomics Biosciences, Inc. This does not represent a conflict of interest for this study.

## References

1. R. H. Austin, K. W. Beeson, L. Eisenstein, H. Frauenfelder, I. C. Gunsalus, Dynamics of ligand binding to myoglobin. Biochemistry 14, 5355–5373 (1975).

2. G. Wagner, K. Wüthrich, Dynamic model of globular protein conformations based on NMR studies in solution. Nature 275, 247–248 (1978).

3. C.-J. Tsai, B. Ma, R. Nussinov, Folding and binding cascades: Shifts in energy landscapes. Proc. Natl. Acad. Sci. 96, 9970–9972 (1999).

4. P. Robustelli, K. A. Stafford, A. G. Palmer, Interpreting protein structural dynamics from NMR chemical shifts. J. Am. Chem. Soc. 134, 6365–6374 (2012).

5. D. S. Wishart, Interpreting protein chemical shift data. Prog. Nucl. Magn. Reson. Spectrosc. 58, 62–87 (2011).

6. M. P. Williamson, Using chemical shift perturbation to characterise ligand binding. Prog. Nucl. Magn. Reson. Spectrosc. 73, 1–16 (2013).

7. C. Dominguez, R. Boelens, A. M. J. J. Bonvin, HADDOCK:L A proteinprotein docking approach based on biochemical or biophysical information. J. Am. Chem. Soc. 125, 1731–1737 (2003).

8. S. J. De Vries, et al., HADDOCK versus HADDOCK: New features and performance of HADDOCK2.0 on the CAPRI targets. Proteins Struct. Funct. Bioinforma. 69, 726–733 (2007).

9. C. Schmitz, et al., “Protein–Protein Docking with HADDOCK” in NMR of Biomolecules, 1st Ed., I. Bertini, K. S. McGreevy, G. Parigi, Eds. (Wiley, 2012), pp. 520–535.

10. S. Feng, et al., Integrated structure prediction of protein–protein docking with experimental restraints using ColabDock. Nat. Mach. Intell. 6, 924–935 (2024).

11. A. Mondal, et al., Structure determination of challenging protein–peptide complexes combining NMR chemical shift data and molecular dynamics simulations. J. Chem. Inf. Model. 63, 2058–2072 (2023).

12. J. Gaza, E. Brini, J. L. MacCallum, K. A. Dill, A. Perez, MELD in action: Harnessing data to accelerate molecular dynamics. J. Chem. Inf. Model. 65, 1685–1693 (2025).

13. T. R. Soderling, J. T. Stull, Structure and regulation of calcium/calmodulin-dependent protein kinases. Chem. Rev. 101, 2341–2352 (2001).

14. F. Malagrinò, F. Troilo, D. Bonetti, A. Toto, S. Gianni, Mapping the allosteric network within a SH3 domain. Sci. Rep. 9, 8279 (2019).

15. G. Alvarez, et al., Loop plasticity drives paralog-specific recognition in BET ET domains. J. Chem. Inf. Model. 66, 4685–4695 (2026).

16. J. Ray, S. W. Englander, Allosteric sensitivity in hemoglobin at the .alpha.-subunit N-terminus studied by hydrogen exchange. Biochemistry 25, 3000–3007 (1986).

17. W. Rist, C. Graf, B. Bukau, M. P. Mayer, Amide hydrogen exchange reveals conformational changes in Hsp70 chaperones important for allosteric regulation. J. Biol. Chem. 281, 16493–16501 (2006).

18. J. M. Aramini, et al., The RAS-binding domain of human BRAF protein serine/threonine kinase exhibits allosteric conformational changes upon binding HRAS. Structure 23, 1382–1393 (2015).

19. X. Luo, Z. Tang, J. Rizo, H. Yu, The Mad2 spindle checkpoint protein undergoes similar major conformational changes upon binding to either Mad1 or Cdc20. Mol. Cell 9, 59–71 (2002).

20. J. Guo, H.-X. Zhou, Protein allostery and conformational dynamics. Chem. Rev. 116, 6503–6515 (2016).

21. R. Nussinov, Introduction to protein ensembles and allostery. Chem. Rev. 116, 6263–6266 (2016).

22. R. Selvaratnam, S. Chowdhury, B. VanSchouwen, G. Melacini, Mapping allostery through the covariance analysis of NMR chemical shifts. Proc. Natl. Acad. Sci. 108, 6133–6138 (2011).

23. R. Selvaratnam, M. Akimoto, B. VanSchouwen, G. Melacini, cAMP-dependent allostery and dynamics in Epac: an NMR view. Biochem. Soc. Trans. 40, 219–223 (2012).

24. S. Boulton, R. Selvaratnam, R. Ahmed, G. Melacini, “Implementation of the NMR CHEmical Shift Covariance Analysis (CHESCA): A Chemical Biologist’s Approach to Allostery” in Protein NMR, Methods in Molecular Biology., R. Ghose, Ed. (Springer New York, 2018), pp. 391–405.

25. C. Olivieri, et al., An NMR portrait of functional and dysfunctional allosteric cooperativity in CAMP-dependent protein kinase A. FEBS Lett. 597, 1055–1072 (2023).

26. R. Selvaratnam, M. T. Mazhab-Jafari, R. Das, G. Melacini, The auto-inhibitory role of the EPAC hinge helix as mapped by NMR. PLoS ONE 7, e48707 (2012).

27. F. H. Schumann, et al., Combined chemical shift changes and amino acid specific chemical shift mapping of protein–protein interactions. J. Biomol. NMR 39, 275–289 (2007).

28. S. Grzesiek, et al., The solution structure of HIV-1 Nef reveals an unexpected fold and permits delineation of the binding surface for the SH3 domain of Hck tyrosine protein kinase. Nat. Struct. Mol. Biol. 3, 340–345 (1996).

