## Supplementary Information 1 for "Allosteric Protein Chemical Shift Perturbations are Ubiquitous"

| | | |
|---|---|---|
| Tiburon L. Benavides | orcid: 0000-0002-5795-4273 | |
| Theresa A. Ramelot | orcid: 0000-0002-0335-1573 | |
| Gaetano T. Montelione | orcid: 0000-0002-9440-3059 | |


**Table of Contents**

Supplementary Text: The CSPdb

SI Table S1. Data for all receptors in this study
SI Table S2. Data for Hydrolase Receptors
SI Fig S1. Confusion Matrix Histograms for Hydrolase Receptors
SI Table S3. Data for Transferase Receptors
SI Fig S2. Confusion Matrix Histograms for Transferase Receptors
SI Table S4. Data for  All Alpha Receptors
SI Fig S3. Confusion Matrix Histograms for All Alpha Receptors
SI Table S5. Data for All Beta Receptors
SI Fig S4. Confusion Matrix Histograms for All Beta Receptors
SI Table S6. Data for Alpha and Beta (a+b) Receptors
SI Fig S5. Confusion Matrix Histograms for Alpha and Beta (a+b) Receptors
SI Table S7. Data for BET-ET Domain Receptors
SI Fig S6. Confusion Matrix Histograms for BET-ET Domain Receptors
SI Table S8. Data for TFIIH Domain Receptors
SI Fig S7. Confusion Matrix Histograms for TFIIH Domain Receptors
SI Table S9. Data for Ubiquitin Domain Receptors
SI Fig S8. Confusion Matrix Histograms for Ubiquitin Domain Receptors
SI Table S10. Data for targets with dissimilar apo/holo experimental conditions
SI Fig S9. Confusion Matrix Histograms for targets with dissimilar apo/holo experimental conditions
SI Fig. S10. CA-inclusive CSP Confusion Matrix Histograms
SI Fig. S11. CA-inclusive vs exclusive CSP F1 scores
SI Fig. S12. CSP DB Confusion Matrix Histograms by closest interchain N-N distance
SI Fig. S13. CSP DB Confusion Matrix Histograms by closest interchain atom-atom distance
SI Fig. S14. F1 Scores for 1D CSPs
SI Fig. S15. PDB Advanced Search Settings
SI Fig. S16. Ideal N/H offsets

SI Fig. S17. CSP significance threshold Histogram
SI Fig. S18. F1 vs MCC scatterplot

References used in Supplementary Text

### Supplementary Text

#### The CSPdb

The CSPdb includes receptors that regulate diverse biological processes, including cell cycle progression (e.g., MAD2: 1KLQ), transcriptional control (e.g., Nrd1: 2MOW, 6GC3), translational regulation, autophagy, apoptosis (e.g., calcineurin: 2JOG; BCL-XL family: 2M04, 2LP8), and cell adhesion (e.g., tensin: 2LOZ). Disease-relevant targets address mechanisms in leukemia and other cancers (e.g., p53: 2MEJ), Alzheimer's disease (e.g., amyloid precursor protein-binding proteins: 2ROZ), and infectious diseases from viruses and bacteria, such as anthrax (2RUI), chikungunya (5I22), SARS-CoV-2 (7PKU), HIV (2L6E), and Ebola (2KQ0).

The CSPdb features several recurring protein domains that exhibit notable conformational changes upon ligand binding. For instance, it includes 23 interactions involving calmodulin (CaM) or CaM-like proteins with peptides (e.g., PDB IDs: 1SY9, 2JZI; full list in **SI Table S1**), which often involve substantial rearrangements upon binding described in detail elsewhere(1). Other receptors known for binding-induced conformational shifts or folding include the G protein Cdc42 (1CF4), the metaphase chromosome spindle checkpoint protein MAD2 (1KLQ), the KIX domain (2LXS), and CREB-binding proteins (e.g., 1R8U, 1L8C).

Additional recurring domains include the extra-terminal (ET) domains of bromodomain proteins BRD3 and BRD4 (e.g., 2NCZ, 2ND0; full list in **SI Table S7**, **SI Figure 6**), which bind intrinsically-disordered regions of viral integrase or chromatin regulator proteins. These interactions trigger subtle allosteric changes distal from the binding site, including the formation of a stabilizing secondary structure that reorients two allosteric helices (2, 3). Similarly, the PDZ2 domain (e.g., 1D5G, 1VJ6, 2M0V) shows allosteric effects mediated by the β2–β3 loop during peptide recognition (4).

Nine entries involve the PH domain of the TFIIH transcription initiation factor subunit Tfb1 (e.g., 2LOX, 2M14; full list in **SI Table S8**, **SI Figure 7**). In these complexes, the ligand is often modeled as a helical polypeptide bound orthogonally to an extended β-sheet, resulting in propagation of allosteric signals via hydrogen-bond rearrangements. Other targets which involve hydrogen bond rearrangement in β-sheets include the Cdc42 (1CF4), Cyclophilin A (2MS4), and SH3 domain (e.g., 2OJ2, 2ROL; full list in **SI Table S1**). An additional eight entries use ubiquitin or ubiquitin-like (e.g. SUMO and NEDD8) protein receptors. Ubiquitin has been used as a model system for NMR studies (5) and for studying allostery (6) (e.g., 2KWV, 2MUR; full list in **SI Table S9**, **SI Figure 8**).

In Other Supplementary Information, you will find a All_Case_Studies.pdf, which provides a concise summary for each of the 139 targets in the CSPdb. Each case study contains the following panels: A) a 15N/1H HSQC scatterplot overlaying the apo and holo assigned spectra with the optimal N/H offset applied; B) Calculated CSPs for receptor residues along the sequence, bars colored by confusion matrix classification; C) the mask of significant CSPs; D) the mask of binding-site residues; E) the confusion matrix classification projected onto the structure (see **Methods** for details). Residues without CSP measurements are shown in gray; ligands are colored cyan.

### SI Table S1. Data for all receptors in this study

| apo_pdb | holo_pdb | apo_bmrb | holo_bmrb | F1 | MCC |
|---|---|---|---|---|---|
| | 2MRE | 25071 | 25070 | 0.80 | 0.60 |
| | 2MCN | 16641 | 19447 | 0.68 | 0.58 |
| | 2N9P | 25913 | 25914 | 0.72 | 0.58 |
| | 2ROZ | 10235 | 10236 | 0.59 | 0.50 |
| | 5I22 | 4871 | 30010 | 0.55 | 0.49 |
| | 6IVU | 36220 | 36221 | 0.67 | 0.48 |
| | 2LBM | 15001 | 17569 | 0.60 | 0.46 |
| | 6FDP | 19757 | 34223 | 0.48 | 0.43 |
| 7JMY | 7JYN | 30782 | 30790 | 0.53 | 0.43 |
| | 2N8T | 25866 | 25865 | 0.64 | 0.42 |
| | 2K7A | 11018 | 16809 | 0.54 | 0.42 |
| | 2PLD | 5318 | 5310 | 0.46 | 0.40 |
| 7JMY | 7JYZ | 30782 | 30791 | 0.50 | 0.40 |
| | 2L6E | 16555 | 17307 | 0.52 | 0.40 |
| | 2RR4 | 11361 | 11115 | 0.56 | 0.40 |
| | 1N7T | 18785 | 5631 | 0.44 | 0.40 |
| | 5MF9 | 34068 | 34067 | 0.54 | 0.39 |
| | 6RH6 | 34394 | 34395 | 0.34 | 0.38 |
| 1W1F | 1WA7 | 6261 | 6456 | 0.51 | 0.38 |
| | 2L5E | 50148 | 17270 | 0.38 | 0.37 |
| 2LSY | 2N1G | 18455 | 25559 | 0.45 | 0.37 |
| | 2YKA | 16697 | 17693 | 0.36 | 0.37 |
| 7JMY | 7JQ8 | 30782 | 30786 | 0.51 | 0.37 |
| | 6G04 | 34247 | 34248 | 0.43 | 0.37 |
| 2JNS | 6BNH | 15125 | 30373 | 0.48 | 0.36 |
| | 2LI5 | 16835 | 17879 | 0.42 | 0.36 |
| | 9QLM | 15670 | 34986 | 0.59 | 0.34 |
| | 2ROL | 15710 | 11040 | 0.43 | 0.34 |
| | 6FDT | 19757 | 34224 | 0.33 | 0.34 |
| | 2KZU | 17018 | 17019 | 0.38 | 0.34 |
| | 2MC6 | 19428 | 19429 | 0.47 | 0.33 |
| | 2JXC | 4184 | 15554 | 0.58 | 0.33 |
| 2LSY | 2LSK | 18455 | 18434 | 0.38 | 0.32 |
| | 2KSP | 15279 | 16671 | 0.40 | 0.32 |
| | 2MUR | 17769 | 25230 | 0.40 | 0.32 |
| | 2M0U | 18824 | 18825 | 0.35 | 0.31 |
| | 6OQJ | 30599 | 30605 | 0.47 | 0.31 |
| | 2BN5 | 6691 | 6690 | 0.28 | 0.31 |
| | 2LVO | 18581 | 18582 | 0.50 | 0.30 |
| | 2KGI | 16209 | 16210 | 0.62 | 0.30 |
| | 6E5N | 25544 | 30500 | 0.40 | 0.30 |
| | 2RQG | 4915 | 10019 | 0.43 | 0.30 |
| | 7KLR | 30808 | 30809 | 0.55 | 0.29 |
| | 2KPL | 6911 | 16559 | 0.40 | 0.28 |
| | 2OJ2 | 4122 | 6581 | 0.37 | 0.28 |
| | 6GC3 | 17173 | 34260 | 0.29 | 0.28 |
| | 5VF0 | 17769 | 30276 | 0.31 | 0.27 |
| | 2LOZ | 10318 | 17364 | 0.38 | 0.27 |
| | 2M0G | 19034 | 18808 | 0.43 | 0.27 |
| | 2K17 | 15670 | 15671 | 0.53 | 0.27 |
| | 2KC8 | 16066 | 16065 | 0.52 | 0.27 |
| | 2L29 | 17128 | 17127 | 0.26 | 0.26 |
| | 2LP0 | 25142 | 17407 | 0.49 | 0.26 |
| | 2MUR | 25229 | 25230 | 0.57 | 0.26 |
| | 2K6Q | 5958 | 15877 | 0.36 | 0.26 |
| | 7X5C | 51441 | 36473 | 0.31 | 0.26 |
| | 2N3K | 15125 | 25649 | 0.39 | 0.26 |
| | 2LXM | 18681 | 18682 | 0.30 | 0.25 |
| 2JNS | 2NCZ | 15125 | 26041 | 0.40 | 0.25 |
| | 2LGK | 17813 | 17812 | 0.52 | 0.25 |
| | 1H8B | 17627 | 4453 | 0.33 | 0.25 |
| | 7PKU | 50446 | 34661 | 0.42 | 0.25 |
| | 6R5G | 28070 | 34384 | 0.39 | 0.25 |
| | 7S5J | 30926 | 30950 | 0.39 | 0.25 |
| | 2IPA | 6075 | 15028 | 0.18 | 0.23 |
| | 2L4T | 17254 | 17255 | 0.31 | 0.23 |
| | 8SG2 | 27579 | 31080 | 0.44 | 0.23 |
| | 2MZW | 25368 | 25504 | 0.29 | 0.23 |
| | 1DDM | 4683 | 4263 | 0.22 | 0.22 |
| | 2JZI | 51289 | 15624 | 0.43 | 0.22 |
| | 2CZY | 27227 | 6749 | 0.38 | 0.22 |
| 7QCX | 1D5G | 34688 | 4516 | 0.27 | 0.22 |
| | 2M5B | 50942 | 19045 | 0.30 | 0.22 |
| | 1KLQ | 52275 | 5299 | 0.32 | 0.22 |
| 1SSF | 2LVM | 5878 | 18579 | 0.31 | 0.20 |
| | 6U19 | 16620 | 27875 | 0.43 | 0.20 |
| | 2B0F | 5659 | 6823 | 0.27 | 0.18 |
| | 2YS5 | 11095 | 11094 | 0.35 | 0.18 |
| 6VED | 2L3R | 30703 | 17200 | 0.30 | 0.18 |
| | 5J8H | 51289 | 30063 | 0.43 | 0.18 |
| 1SSF | 2MWP | 5878 | 25348 | 0.25 | 0.17 |
| | 6H8C | 18827 | 34307 | 0.29 | 0.17 |
| | 2LXS | 6095 | 18694 | 0.32 | 0.17 |
| | 2FFK | 6809 | 7024 | 0.16 | 0.15 |
| | 2KWI | 15524 | 15525 | 0.40 | 0.15 |
| | 1VJ6 | 5131 | 6060 | 0.29 | 0.15 |
| | 2LP8 | 18250 | 18238 | 0.18 | 0.15 |
| | 2M0K | 51289 | 17771 | 0.41 | 0.14 |
| | 2KTB | 51472 | 16694 | 0.20 | 0.13 |
| | 1PD7 | 6899 | 5808 | 0.37 | 0.13 |
| 1V49 | 2N9X | 5958 | 25919 | 0.28 | 0.13 |
| | 6QXZ | 27250 | 27251 | 0.42 | 0.13 |
| | 2NMB | 4683 | 4263 | 0.15 | 0.13 |
| | 2JW1 | 15503 | 15504 | 0.30 | 0.13 |
| 1SSF | 2MWO | 5878 | 25347 | 0.26 | 0.12 |
| | 1ONV | 27288 | 5685 | 0.34 | 0.12 |
| | 2MES | 51289 | 19238 | 0.45 | 0.11 |
| 2KQ3 | 2KHS | 16585 | 15357 | 0.31 | 0.11 |
| | 2LE8 | 16396 | 17702 | 0.21 | 0.10 |
| | 2KOH | 27205 | 16507 | 0.30 | 0.10 |
| | 2FIN | 6809 | 7024 | 0.10 | 0.09 |
| 2JNS | 2ND0 | 15125 | 26042 | 0.31 | 0.09 |
| | 2HUG | 6592 | 7241 | 0.35 | 0.09 |
| | 2K7A | 5461 | 16809 | 0.21 | 0.09 |
| | 2L0Y | 15763 | 17067 | 0.27 | 0.09 |
| | 2N55 | 52209 | 25694 | 0.44 | 0.07 |
| | 2MJV | 15057 | 19738 | 0.30 | 0.07 |
| | 2RUI | 16811 | 11570 | 0.16 | 0.06 |
| | 1SY9 | 51289 | 5480 | 0.36 | 0.06 |
| | 2M56 | 4154 | 19038 | 0.12 | 0.04 |
| | 2G35 | 15792 | 7061 | 0.17 | 0.04 |
| | 9R3Y | 26643 | 34992 | 0.25 | 0.04 |
| | 2LSP | 15057 | 18439 | 0.26 | 0.03 |
| | 2M0J | 51289 | 17771 | 0.35 | 0.03 |
| 2L5U | 2L75 | 17285 | 17344 | 0.44 | 0.02 |
| | 7SFT | 36097 | 30958 | 0.27 | 0.02 |
| | 2MNZ | 19913 | 19914 | 0.44 | 0.02 |
| | 5U5S | 50149 | 30206 | 0.19 | 0.01 |
| | 2MSR | 34179 | 25130 | 0.12 | 0.00 |
| | 2KVM | 15674 | 16778 | 0.33 | -0.00 |
| | 2JMX | 6564 | 15072 | 0.09 | -0.01 |
| | 2N80 | 25883 | 25829 | 0.26 | -0.02 |
| | 2LGG | 17813 | 17808 | 0.37 | -0.03 |
| | 2I94 | 5332 | 7293 | 0.17 | -0.03 |
| | 2K8F | 34231 | 15944 | 0.22 | -0.05 |
| 2JNS | 2ND1 | 15125 | 26043 | 0.23 | -0.06 |
| | 5OEO | 51289 | 34161 | 0.16 | -0.07 |
| 2JNS | 6BGG | 15125 | 30367 | 0.14 | -0.08 |

(continued)

| apo_pdb | holo_pdb | apo_bmrb | holo_bmrb | F1 | MCC | apo_pdb | holo_pdb | apo_bmrb | holo_bmrb | F1 | MCC |
|---|---|---|---|---|---|---|---|---|---|---|---|
| | 2M0G | 18802 | 18808 | 0.29 | 0.25 | | 6U4N | 17827 | 30659 | 0.10 | -0.08 |
| | 2MOW | 17173 | 19954 | 0.21 | 0.25 | | 2LGF | 51289 | 17807 | 0.28 | -0.09 |
| | 2K7L | 27288 | 15919 | 0.39 | 0.25 | | 9C5E | 18642 | 31177 | 0.35 | -0.11 |
| | 2N3A | 34537 | 25639 | 0.37 | 0.24 | | 2RSE | 16931 | 11471 | 0.00 | -0.16 |
| 1U2N | 1R8U | 6268 | 5987 | 0.56 | 0.24 | | 2K6D | 52080 | 15866 | 0.15 | -0.18 |
| | 2KYL | 15972 | 16967 | 0.41 | 0.23 | | | | | | |

### SI Table S2. Data for Hydrolase Receptors

| apo_pdb | holo_pdb | apo_bmrb | holo_bmrb | F1 | MCC | apo_pdb | holo_pdb | apo_bmrb | holo_bmrb | F1 | MCC |
|---|---|---|---|---|---|---|---|---|---|---|---|
| | 2LBM | 15001 | 17569 | 0.60 | 0.46 | | 2JZI | 51289 | 15624 | 0.43 | 0.22 |
| | 6FDP | 19757 | 34223 | 0.48 | 0.43 | 7QCX | 1D5G | 34688 | 4516 | 0.27 | 0.22 |
| 2GS0 | 2LOX | 6225 | 18229 | 0.47 | 0.42 | | 6OQJ | 4276 | 30605 | 0.32 | 0.22 |
| | 2PLD | 5318 | 5310 | 0.46 | 0.40 | | 2KWI | 15524 | 15525 | 0.40 | 0.15 |
| | 6OQJ | 30599 | 30605 | 0.47 | 0.31 | | 1VJ6 | 5131 | 6060 | 0.29 | 0.15 |
| | 2RQG | 4915 | 10019 | 0.43 | 0.30 | | 1ONV | 27288 | 5685 | 0.34 | 0.12 |
| | 6GC3 | 17173 | 34260 | 0.29 | 0.28 | 2KQ3 | 2KHS | 16585 | 15357 | 0.31 | 0.11 |
| | 2LOZ | 10318 | 17364 | 0.38 | 0.27 | | 2LE8 | 16396 | 17702 | 0.21 | 0.10 |
| | 1CF4 | 18251 | 4700 | 0.49 | 0.27 | | 2HUG | 6592 | 7241 | 0.35 | 0.09 |
| | 2KC8 | 16066 | 16065 | 0.52 | 0.27 | | 2LTO | 10276 | 18490 | 0.14 | 0.09 |
| | 6R5G | 28070 | 34384 | 0.39 | 0.25 | 2L5U | 2L75 | 17285 | 17344 | 0.44 | 0.02 |
| | 2K7L | 27288 | 15919 | 0.39 | 0.25 | | 7NQC | 51289 | 34608 | 0.34 | 0.01 |
| | 2KYL | 15972 | 16967 | 0.41 | 0.23 | | 2RMK | 5511 | 11010 | 0.18 | -0.02 |
| | 2KWI | 15230 | 15525 | 0.36 | 0.23 | 2JNS | 6BGG | 15125 | 30367 | 0.14 | -0.08 |

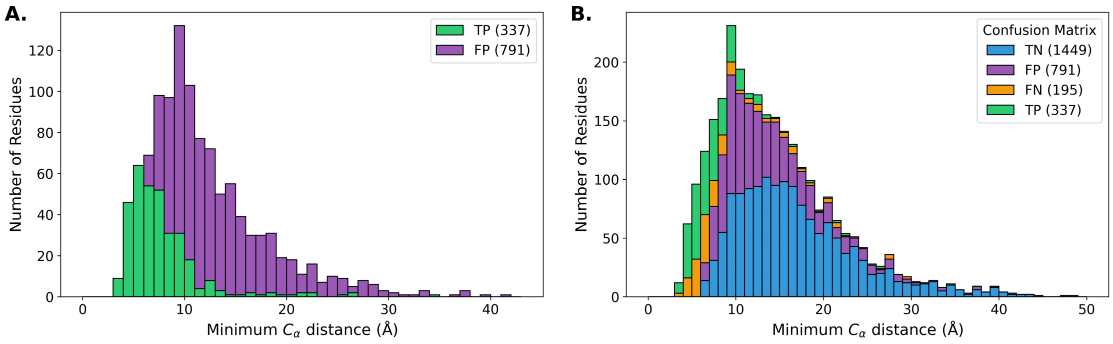


**SI Fig. S1. Confusion Matrix Histograms for Hydrolase Receptors**

### SI Table S3. Data for Transferase Receptors

| apo_pdb | holo_pdb | apo_bmrb | holo_bmrb | F1 | MCC | apo_pdb | holo_pdb | apo_bmrb | holo_bmrb | F1 | MCC |
|---|---|---|---|---|---|---|---|---|---|---|---|
| | 2MRE | 25071 | 25070 | 0.80 | 0.60 | | 2LVO | 18610 | 18582 | 0.34 | 0.19 |
| | 2N9P | 25913 | 25914 | 0.72 | 0.58 | | 2LAS | 50128 | 25553 | 0.28 | 0.19 |
| 7JMY | 7JYN | 30782 | 30790 | 0.53 | 0.43 | 2L0I | 5M9D | 17044 | 34058 | 0.33 | 0.18 |
| | 2N8T | 25866 | 25865 | 0.64 | 0.42 | | 2YS5 | 11095 | 11094 | 0.35 | 0.18 |
| | 2K7A | 11018 | 16809 | 0.54 | 0.42 | 6VED | 2L3R | 30703 | 17200 | 0.30 | 0.18 |
| | 2C52 | 15398 | 6874 | 0.56 | 0.41 | | 5J8H | 51289 | 30063 | 0.43 | 0.18 |
| | 2PLD | 5318 | 5310 | 0.46 | 0.40 | | 2N83 | 25828 | 25833 | 0.36 | 0.17 |
| 5AHT | 2KPZ | 25349 | 16574 | 0.55 | 0.40 | | 5IAY | 30703 | 30019 | 0.22 | 0.17 |
| 1W1F | 1WA7 | 6261 | 6456 | 0.51 | 0.38 | | 2LXS | 6095 | 18694 | 0.32 | 0.17 |
| 2LSY | 2N1G | 18455 | 25559 | 0.45 | 0.37 | | 2MV7 | 18094 | 19516 | 0.38 | 0.15 |
| | 2KWU | 17769 | 16880 | 0.40 | 0.37 | | 7ZEY | 16983 | 34727 | 0.44 | 0.14 |
| 2KM4 | 2L0I | 16411 | 17044 | 0.36 | 0.36 | | 2MRE | 18610 | 25070 | 0.28 | 0.13 |
| | 2KWV | 17769 | 16885 | 0.39 | 0.34 | | 5JYV | 30092 | 30093 | 0.31 | 0.13 |
| 2LSG | 2LSJ | 18431 | 18433 | 0.35 | 0.34 | | 9C5E | 18610 | 31177 | 0.32 | 0.13 |
| | 2MC6 | 19428 | 19429 | 0.47 | 0.33 | | 6QXZ | 27250 | 27251 | 0.42 | 0.13 |
| 2LSY | 2LSK | 18455 | 18434 | 0.38 | 0.32 | | 2H7D | 15792 | 7150 | 0.24 | 0.11 |
| | 2LVO | 18581 | 18582 | 0.50 | 0.30 | | 2M86 | 15945 | 19230 | 0.25 | 0.10 |
| 5AHT | 2KQ0 | 25349 | 16575 | 0.50 | 0.30 | | 2N1A | 6304 | 25553 | 0.10 | 0.10 |
| | 2OJ2 | 4122 | 6581 | 0.37 | 0.28 | | 2RSE | 50325 | 11471 | 0.10 | 0.10 |
| | 5VF0 | 17769 | 30276 | 0.31 | 0.27 | | 2KA6 | 4789 | 16015 | 0.51 | 0.09 |
| | 5JYV | 30091 | 30093 | 0.45 | 0.27 | | 2LTO | 10276 | 18490 | 0.14 | 0.09 |
| | 1CF4 | 18251 | 4700 | 0.49 | 0.27 | | 2K7A | 5461 | 16809 | 0.21 | 0.09 |
| 2JNS | 2NCZ | 15125 | 26041 | 0.40 | 0.25 | | 2KJE | 4789 | 16318 | 0.45 | 0.07 |
| | 2LGK | 17813 | 17812 | 0.52 | 0.25 | | 2MZD | 34231 | 25484 | 0.35 | 0.06 |
| | 2MOW | 17173 | 19954 | 0.21 | 0.25 | | 2RSY | 7141 | 11508 | 0.30 | 0.05 |
| | 2MBB | 15410 | 19394 | 0.43 | 0.24 | | 2G35 | 15792 | 7061 | 0.17 | 0.04 |
| 2L0I | 5LVF | 17044 | 34041 | 0.31 | 0.24 | | 2MSR | 34179 | 25130 | 0.12 | 0.00 |
| 1U2N | 1R8U | 6268 | 5987 | 0.56 | 0.24 | | 2RMK | 5511 | 11010 | 0.18 | -0.02 |
| | 1I5H | 26698 | 4963 | 0.40 | 0.23 | | 2LGG | 17813 | 17808 | 0.37 | -0.03 |
| | 2KYL | 15972 | 16967 | 0.41 | 0.23 | | 2I94 | 5332 | 7293 | 0.17 | -0.03 |
| | 8SG2 | 27579 | 31080 | 0.44 | 0.23 | | 2K8F | 34231 | 15944 | 0.22 | -0.05 |
| 1U2N | 1L8C | 6268 | 5327 | 0.58 | 0.22 | 2JNS | 2ND1 | 15125 | 26043 | 0.23 | -0.06 |
| | 7ZEY | 34724 | 34727 | 0.36 | 0.22 | | 2LV6 | 51289 | 18556 | 0.35 | -0.07 |
| 1U2N | 2KA4 | 6268 | 16014 | 0.55 | 0.21 | | 2N83 | 25883 | 25833 | 0.29 | -0.10 |
| | 2MH0 | 34231 | 19610 | 0.33 | 0.21 | | 9C5E | 18642 | 31177 | 0.35 | -0.11 |
| | 2MPS | 6612 | 18876 | 0.34 | 0.20 | | 2RSE | 16931 | 11471 | 0.00 | -0.16 |
| | 6U19 | 16620 | 27875 | 0.43 | 0.20 | | 2JMF | 6262 | 15016 | 0.13 | -0.17 |

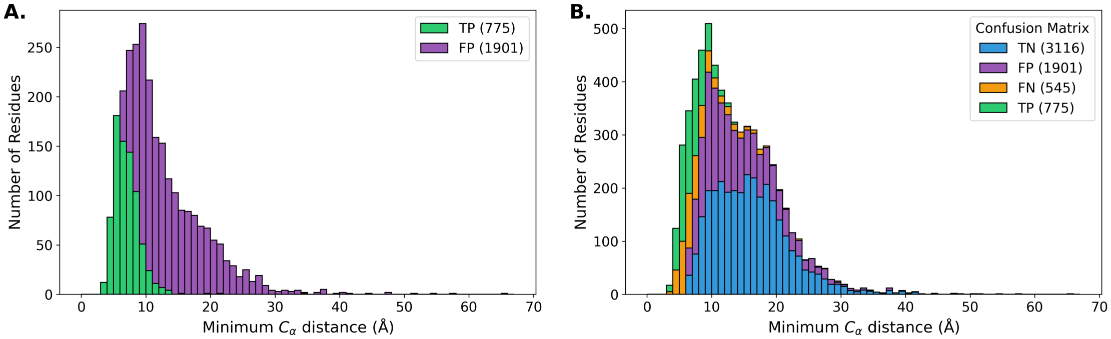


**SI Fig. S2. Confusion Matrix Histograms for Transferase Receptors**

### SI Table S4. Data for All Alpha Receptors

| apo_pdb | holo_pdb | apo_bmrb | holo_bmrb | F1 | MCC |
|---|---|---|---|---|---|
| | 6FDP | 19757 | 34223 | 0.48 | 0.43 |
| | 2M5A | 6804 | 19043 | 0.62 | 0.42 |
| | 2C52 | 15398 | 6874 | 0.56 | 0.41 |
| | 2L6E | 16555 | 17307 | 0.52 | 0.40 |
| | 6FDT | 19757 | 34224 | 0.33 | 0.34 |
| | 2M5A | 5656 | 19043 | 0.53 | 0.34 |
| | 2JXC | 4184 | 15554 | 0.58 | 0.33 |
| | 2KSP | 15279 | 16671 | 0.40 | 0.32 |
| | 2RQG | 4915 | 10019 | 0.43 | 0.30 |
| | 2LY4 | 11532 | 18709 | 0.40 | 0.30 |
| | 2MG5 | 51289 | 19586 | 0.44 | 0.27 |
| | 2LP0 | 25142 | 17407 | 0.49 | 0.26 |
| | 1H8B | 17627 | 4453 | 0.33 | 0.25 |
| | 2KNH | 7396 | 16467 | 0.30 | 0.24 |
| | 2JZI | 51289 | 15624 | 0.43 | 0.22 |
| | 2LL6 | 51289 | 18027 | 0.39 | 0.21 |
| | 2MPS | 6612 | 18876 | 0.34 | 0.20 |
| | 2M56 | 17415 | 19038 | 0.12 | 0.18 |
| | 5J8H | 51289 | 30063 | 0.43 | 0.18 |
| | 2N83 | 25828 | 25833 | 0.36 | 0.17 |
| | 2LL7 | 51289 | 18028 | 0.33 | 0.17 |
| | 2LXS | 6095 | 18694 | 0.32 | 0.17 |
| | 1LXF | 30966 | 5386 | 0.37 | 0.14 |
| | 2M0K | 51289 | 17771 | 0.41 | 0.14 |
| | 2KTB | 51472 | 16694 | 0.20 | 0.13 |
| | 1PD7 | 6899 | 5808 | 0.37 | 0.13 |
| | 2KNE | 51289 | 16465 | 0.36 | 0.13 |
| | 1ONV | 27288 | 5685 | 0.34 | 0.12 |
| | 2MES | 51289 | 19238 | 0.45 | 0.11 |
| | 2MGU | 51289 | 19604 | 0.42 | 0.09 |
| | 2L53 | 51289 | 17264 | 0.27 | 0.06 |
| | 1SY9 | 51289 | 5480 | 0.36 | 0.06 |
| | 6BUT | 51289 | 27095 | 0.25 | 0.04 |
| | 2M56 | 4154 | 19038 | 0.12 | 0.04 |
| | 2M0J | 51289 | 17771 | 0.35 | 0.03 |
| | 2MSR | 34179 | 25130 | 0.12 | 0.00 |
| | 2N80 | 25883 | 25829 | 0.26 | -0.02 |
| | 2I94 | 5332 | 7293 | 0.17 | -0.03 |
| | 5TP6 | 51289 | 30196 | 0.21 | -0.03 |
| | 1NPQ | 4232 | 5738 | 0.24 | -0.03 |
| | 2N83 | 25883 | 25833 | 0.29 | -0.10 |
| | 2N80 | 51835 | 25829 | 0.04 | -0.13 |

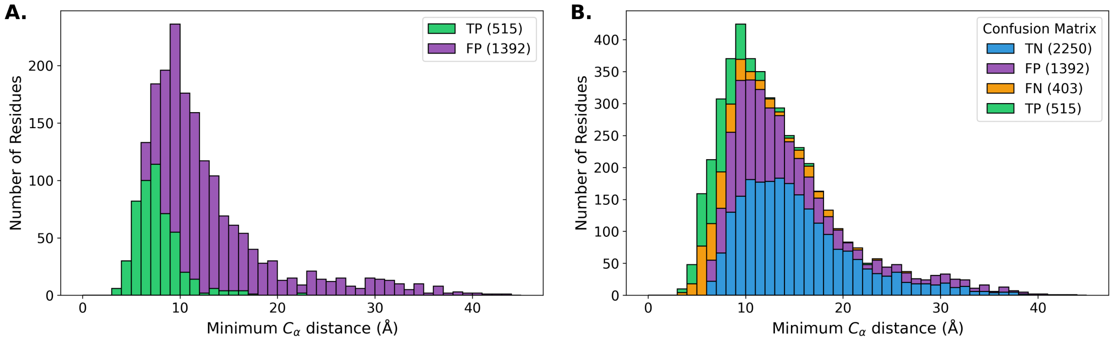


SI Fig. S3. Confusion Matrix Histograms for All Alpha Receptors

### SI Table S5. Data for All Beta Receptors

| apo_pdb | holo_pdb | apo_bmrb | holo_bmrb | F1 | MCC |
|---|---|---|---|---|---|
| | 1YWI | 6658 | 6659 | 0.78 | 0.72 |
| | 2MCN | 16641 | 19447 | 0.68 | 0.58 |
| | 2FIN | 5073 | 7024 | 0.65 | 0.52 |
| | 2ROZ | 10235 | 10236 | 0.59 | 0.50 |
| | 5I22 | 4871 | 30010 | 0.55 | 0.49 |
| | 2AFF | 5959 | 6748 | 0.62 | 0.48 |
| 2RQT | 2RQU | 11081 | 11082 | 0.60 | 0.42 |
| 2GS0 | 2LOX | 6225 | 18229 | 0.47 | 0.42 |
| | 2K7A | 11018 | 16809 | 0.54 | 0.42 |
| | 1N7T | 18785 | 5631 | 0.44 | 0.40 |
| 1W1F | 1WA7 | 6261 | 6456 | 0.51 | 0.38 |
| 2GS0 | 2MKR | 6225 | 19791 | 0.29 | 0.33 |
| 2GS0 | 2N0Y | 6225 | 25540 | 0.37 | 0.33 |
| | 2M0U | 18824 | 18825 | 0.35 | 0.31 |
| | 2RUK | 4901 | 11578 | 0.44 | 0.30 |
| | 2MS4 | 2208 | 25104 | 0.29 | 0.29 |
| | 2KPL | 6911 | 16559 | 0.40 | 0.28 |
| | 2LOZ | 10318 | 17364 | 0.38 | 0.27 |
| | 2L29 | 17128 | 17127 | 0.26 | 0.26 |
| 2GS0 | 2M14 | 6225 | 18842 | 0.24 | 0.23 |
| | 1I5H | 26698 | 4963 | 0.40 | 0.23 |
| | 2L4T | 17254 | 17255 | 0.31 | 0.23 |
| 2GS0 | 5URN | 6225 | 30243 | 0.29 | 0.23 |
| | 2M3M | 17373 | 17942 | 0.42 | 0.22 |
| | 1DDM | 4683 | 4263 | 0.22 | 0.22 |
| 7QCX | 1D5G | 34688 | 4516 | 0.27 | 0.22 |
| 2GS0 | 2N23 | 6225 | 25584 | 0.17 | 0.22 |
| 1SSF | 2LVM | 5878 | 18579 | 0.31 | 0.20 |
| | 2RVB | 4901 | 11594 | 0.44 | 0.20 |
| | 2KVQ | 15490 | 16788 | 0.25 | 0.16 |
| | 2FFK | 6809 | 7024 | 0.16 | 0.15 |
| | 1VJ6 | 5131 | 6060 | 0.29 | 0.15 |
| | 2NMB | 4683 | 4263 | 0.15 | 0.13 |
| 2KQ3 | 2KHS | 16585 | 15357 | 0.31 | 0.11 |
| | 2MCN | 18610 | 19447 | 0.40 | 0.10 |
| | 2KOH | 27205 | 16507 | 0.30 | 0.10 |
| | 2FIN | 6809 | 7024 | 0.10 | 0.09 |
| | 5XV8 | 4901 | 36101 | 0.37 | 0.09 |
| | 2HUG | 6592 | 7241 | 0.35 | 0.09 |
| | 2K7A | 5461 | 16809 | 0.21 | 0.09 |
| | 2LZ6 | 18610 | 18737 | 0.29 | 0.08 |
| | 2G35 | 15792 | 7061 | 0.17 | 0.04 |
| | 2LZ6 | 15407 | 18737 | 0.28 | 0.02 |
| | 2KVM | 15674 | 16778 | 0.33 | -0.00 |
| | 2N80 | 25883 | 25829 | 0.26 | -0.02 |
| | 2MNU | 4283 | 19906 | 0.19 | -0.04 |
| | 2L29 | 34000 | 17127 | 0.16 | -0.08 |
| | 2N80 | 51835 | 25829 | 0.04 | -0.13 |
| | 2JMF | 6262 | 15016 | 0.13 | -0.17 |

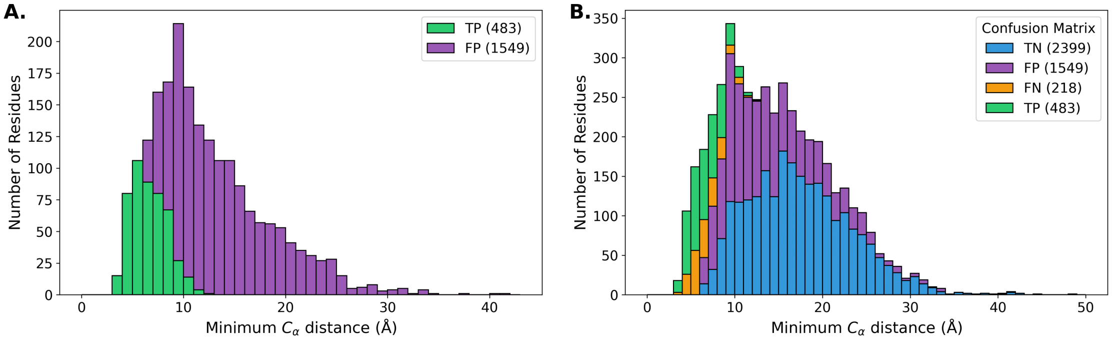


**SI Fig. S4. Confusion Matrix Histograms for All Beta Receptors**

### SI Table S6. Data for Alpha and Beta (a+b) Receptors

| apo_pdb | holo_pdb | apo_bmrb | holo_bmrb | F1 | MCC | apo_pdb | holo_pdb | apo_bmrb | holo_bmrb | F1 | MCC |
|---|---|---|---|---|---|---|---|---|---|---|---|
| | 2MRE | 25071 | 25070 | 0.80 | 0.60 | | 2MBB | 15410 | 19394 | 0.43 | 0.24 |
| | 2MCN | 16641 | 19447 | 0.68 | 0.58 | | 1KLQ | 52275 | 5299 | 0.32 | 0.22 |
| | 2FIN | 5073 | 7024 | 0.65 | 0.52 | | 2LVO | 18610 | 18582 | 0.34 | 0.19 |
| | 2K7A | 11018 | 16809 | 0.54 | 0.42 | | 2M56 | 17415 | 19038 | 0.12 | 0.18 |
| | 2PLD | 5318 | 5310 | 0.46 | 0.40 | | 6H8C | 18827 | 34307 | 0.29 | 0.17 |
| 2LD9 | 2MBH | 15410 | 19399 | 0.40 | 0.40 | | 2MRE | 18610 | 25070 | 0.28 | 0.13 |
| | 2KWU | 17769 | 16880 | 0.40 | 0.37 | | 2MCN | 18610 | 19447 | 0.40 | 0.10 |
| | 2LI5 | 16835 | 17879 | 0.42 | 0.36 | | 2N1A | 6304 | 25553 | 0.10 | 0.10 |
| | 2G3Q | 4769 | 7002 | 0.38 | 0.35 | | 2FIN | 6809 | 7024 | 0.10 | 0.09 |
| | 2KWV | 17769 | 16885 | 0.39 | 0.34 | | 2K7A | 5461 | 16809 | 0.21 | 0.09 |
| | 1M4P | 50765 | 5532 | 0.29 | 0.32 | | 6XMN | 30316 | 50331 | 0.40 | 0.09 |
| | 2MUR | 17769 | 25230 | 0.40 | 0.32 | | 2LZ6 | 18610 | 18737 | 0.29 | 0.08 |
| | 2LVO | 18581 | 18582 | 0.50 | 0.30 | | 2N55 | 52209 | 25694 | 0.44 | 0.07 |
| | 5VF0 | 17769 | 30276 | 0.31 | 0.27 | | 2RSY | 7141 | 11508 | 0.30 | 0.05 |
| | 2M0G | 19034 | 18808 | 0.43 | 0.27 | | 2M56 | 4154 | 19038 | 0.12 | 0.04 |
| | 2MUR | 25229 | 25230 | 0.57 | 0.26 | | 2LZ6 | 15407 | 18737 | 0.28 | 0.02 |
| | 2K6Q | 5958 | 15877 | 0.36 | 0.26 | | 2K6D | 52080 | 15866 | 0.15 | -0.18 |
| | 2M0G | 18802 | 18808 | 0.29 | 0.25 | | | | | | |

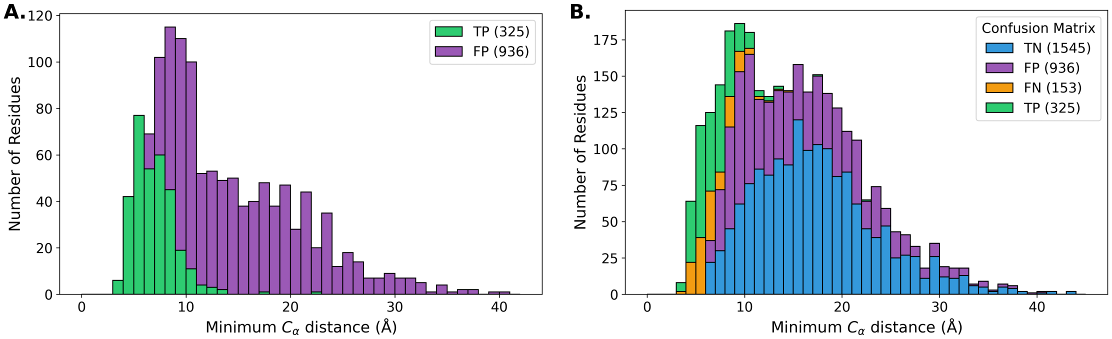


**SI Fig. S5. Confusion Matrix Histograms for Alpha and Beta (a+b) Receptors**

### SI Table S7. Data for BET-ET Domain Receptors

| apo_pdb | holo_pdb | apo_bmrb | holo_bmrb | F1 | MCC | apo_pdb | holo_pdb | apo_bmrb | holo_bmrb | F1 | MCC |
|---|---|---|---|---|---|---|---|---|---|---|---|
| 7JMY | 7JYN | 30782 | 30790 | 0.53 | 0.43 | 2JNS | 2NCZ | 15125 | 26041 | 0.40 | 0.25 |
| 7JMY | 7JYZ | 30782 | 30791 | 0.50 | 0.40 | 2JNS | 2ND0 | 15125 | 26042 | 0.31 | 0.09 |
| 7JMY | 7JQ8 | 30782 | 30786 | 0.51 | 0.37 | 2JNS | 2ND1 | 15125 | 26043 | 0.23 | -0.06 |
| 2JNS | 6BNH | 15125 | 30373 | 0.48 | 0.36 | 2JNS | 6BGG | 15125 | 30367 | 0.14 | -0.08 |

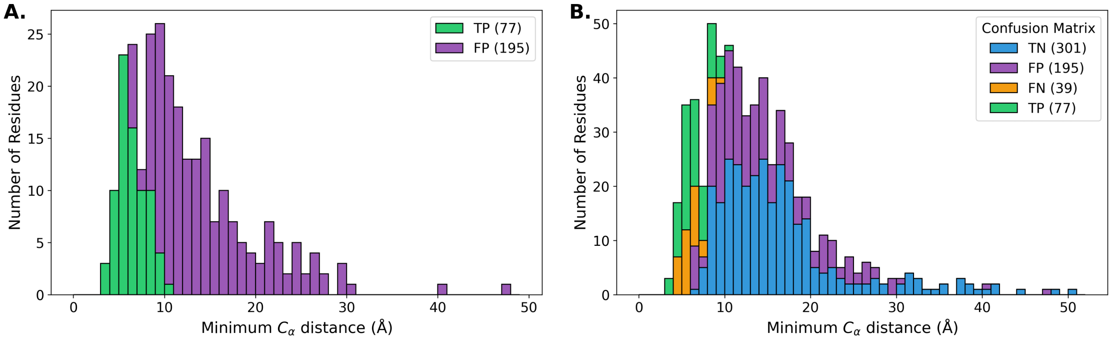


SI Fig. S6. Confusion Matrix Histograms for BET-ET Domain Receptors

### SI Table S8. Data for TFIIH Domain Receptors

| apo_pdb | holo_pdb | apo_bmrb | holo_bmrb | F1 | MCC | apo_pdb | holo_pdb | apo_bmrb | holo_bmrb | F1 | MCC |
|---|---|---|---|---|---|---|---|---|---|---|---|
| 2GS0 | 2LOX | 6225 | 18229 | 0.47 | 0.42 | 2GS0 | 5URN | 6225 | 30243 | 0.29 | 0.23 |
| 2GS0 | 2MKR | 6225 | 19791 | 0.29 | 0.33 | 2GS0 | 2N23 | 6225 | 25584 | 0.17 | 0.22 |
| 2GS0 | 2N0Y | 6225 | 25540 | 0.37 | 0.33 | | 2RVB | 4901 | 11594 | 0.44 | 0.20 |
| | 2RUK | 4901 | 11578 | 0.44 | 0.30 | | 5XV8 | 4901 | 36101 | 0.37 | 0.09 |
| 2GS0 | 2M14 | 6225 | 18842 | 0.24 | 0.23 | | | | | | |

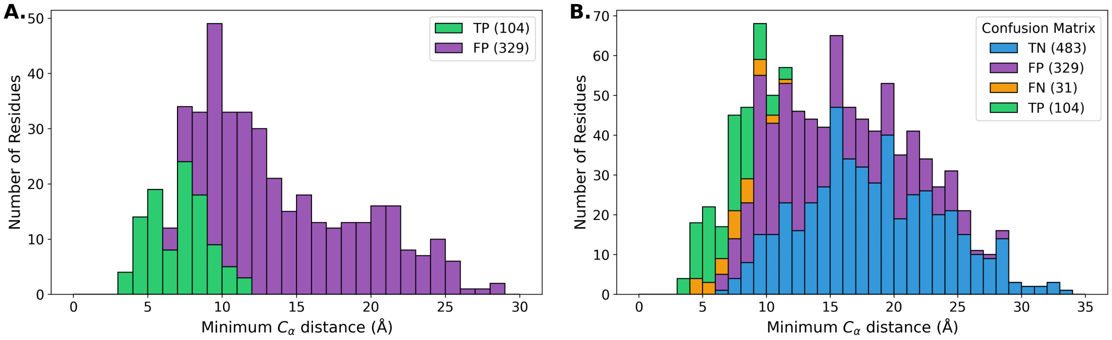


**SI Fig. S7. Confusion Matrix Histograms for TFIIH Domain Receptors**

### SI Table S9. Data for Ubiquitin Domain Receptors

| apo_pdb | holo_pdb | apo_bmrb | holo_bmrb | F1 | MCC | apo_pdb | holo_pdb | apo_bmrb | holo_bmrb | F1 | MCC |
|---|---|---|---|---|---|---|---|---|---|---|---|
| | 2MUR | 17769 | 25230 | 0.40 | 0.32 | | 2MUR | 25229 | 25230 | 0.57 | 0.26 |
| | 5VF0 | 17769 | 30276 | 0.31 | 0.27 | | | | | | |

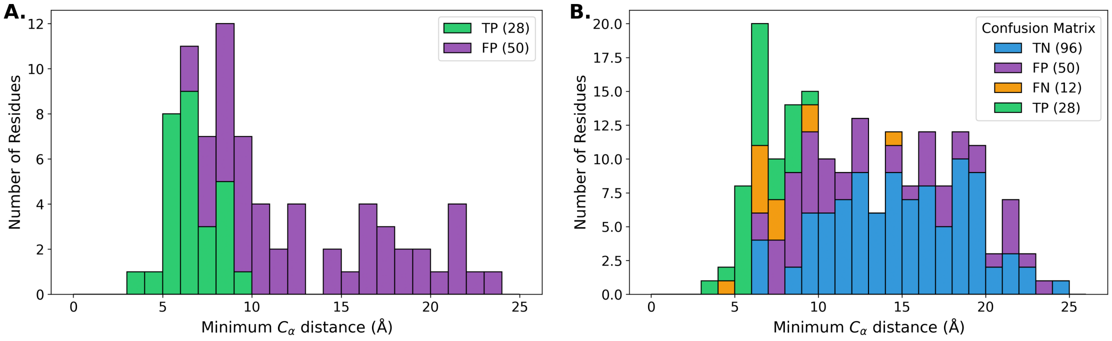


**SI Fig. S8. Confusion Matrix Histograms for Ubiquitin Domain Receptors**

### SI Table S10. Data for targets with dissimilar apo/holo experimental conditions

| apo_pdb | holo_pdb | apo_bmrb | holo_bmrb | F1 | MCC |
|---|---|---|---|---|---|
| | 1YWI | 6658 | 6659 | 0.78 | 0.72 |
| | 2FIN | 5073 | 7024 | 0.65 | 0.52 |
| | 2AFF | 5959 | 6748 | 0.62 | 0.48 |
| 1XWH | 2KE1 | 6374 | 16510 | 0.68 | 0.46 |
| 2RQT | 2RQU | 11081 | 11082 | 0.60 | 0.42 |
| 2GS0 | 2LOX | 6225 | 18229 | 0.47 | 0.42 |
| | 7OVC | 6546 | 34638 | 0.49 | 0.42 |
| | 2M5A | 6804 | 19043 | 0.62 | 0.42 |
| | 2C52 | 15398 | 6874 | 0.56 | 0.41 |
| 5AHT | 2KPZ | 25349 | 16574 | 0.55 | 0.40 |
| | 2KID | 4879 | 16270 | 0.39 | 0.40 |
| 2LD9 | 2MBH | 15410 | 19399 | 0.40 | 0.40 |
| | 2KWU | 17769 | 16880 | 0.40 | 0.37 |
| 2KM4 | 2L0I | 16411 | 17044 | 0.36 | 0.36 |
| | 2G3Q | 4769 | 7002 | 0.38 | 0.35 |
| | 2KWV | 17769 | 16885 | 0.39 | 0.34 |
| | 2M5A | 5656 | 19043 | 0.53 | 0.34 |
| 2LSG | 2LSJ | 18431 | 18433 | 0.35 | 0.34 |
| 2AWT | 2N9E | 6801 | 25901 | 0.30 | 0.33 |
| 2GS0 | 2MKR | 6225 | 19791 | 0.29 | 0.33 |
| 2GS0 | 2N0Y | 6225 | 25540 | 0.37 | 0.33 |
| | 1M4P | 50765 | 5532 | 0.29 | 0.32 |
| | 6K5T | 25299 | 36260 | 0.46 | 0.32 |
| 2LPC | 2M04 | 18250 | 18793 | 0.45 | 0.31 |
| | 2N4Q | 18094 | 25677 | 0.40 | 0.30 |
| 5AHT | 2KQ0 | 25349 | 16575 | 0.50 | 0.30 |
| | 2KJ4 | 4276 | 16311 | 0.38 | 0.30 |
| | 2RUK | 4901 | 11578 | 0.44 | 0.30 |
| | 2LY4 | 11532 | 18709 | 0.40 | 0.30 |
| | 2MS4 | 2208 | 25104 | 0.29 | 0.29 |
| | 2MPM | 4155 | 19989 | 0.41 | 0.28 |
| | 6K5R | 6801 | 36259 | 0.43 | 0.28 |
| | 2MG5 | 51289 | 19586 | 0.44 | 0.27 |
| | 5JYV | 30091 | 30093 | 0.45 | 0.27 |
| 5YDY | 2LAW | 36115 | 17538 | 0.50 | 0.27 |
| | 1CF4 | 18251 | 4700 | 0.49 | 0.27 |
| | 2MBB | 15410 | 19394 | 0.43 | 0.24 |
| | 2KNH | 7396 | 16467 | 0.30 | 0.24 |
| 2L0I | 5LVF | 17044 | 34041 | 0.31 | 0.24 |
| 2GS0 | 2M14 | 6225 | 18842 | 0.24 | 0.23 |
| | 1I5H | 26698 | 4963 | 0.40 | 0.23 |
| | 2KWI | 15230 | 15525 | 0.36 | 0.23 |
| 2GS0 | 5URN | 6225 | 30243 | 0.29 | 0.23 |
| | 2M3M | 17373 | 17942 | 0.42 | 0.22 |
| 1U2N | 1L8C | 6268 | 5327 | 0.58 | 0.22 |
| | 7ZEY | 34724 | 34727 | 0.36 | 0.22 |
| 2GS0 | 2N23 | 6225 | 25584 | 0.17 | 0.22 |
| | 6OQJ | 4276 | 30605 | 0.32 | 0.22 |
| | 2LL6 | 51289 | 18027 | 0.39 | 0.21 |
| 1U2N | 2KA4 | 6268 | 16014 | 0.55 | 0.21 |
| | 6FGP | 16803 | 34232 | 0.38 | 0.21 |
| | 7L8V | 26682 | 27692 | 0.29 | 0.21 |
| | 2MH0 | 34231 | 19610 | 0.33 | 0.21 |
| | 2MPS | 6612 | 18876 | 0.34 | 0.20 |
| | 2RVB | 4901 | 11594 | 0.44 | 0.20 |
| | 6CO4 | 30824 | 25979 | 0.31 | 0.20 |
| | 2LVO | 18610 | 18582 | 0.34 | 0.19 |
| | 2LAS | 50128 | 25553 | 0.28 | 0.19 |
| | 2M56 | 17415 | 19038 | 0.12 | 0.18 |
| 2L0I | 5M9D | 17044 | 34058 | 0.33 | 0.18 |
| | 2MMA | 19849 | 19850 | 0.30 | 0.18 |
| | 5GOW | 4901 | 36013 | 0.39 | 0.18 |
| | 2N83 | 25828 | 25833 | 0.36 | 0.17 |
| | 2LL7 | 51289 | 18028 | 0.33 | 0.17 |
| | 5IAY | 30703 | 30019 | 0.22 | 0.17 |
| | 5J7J | 51289 | 30062 | 0.48 | 0.17 |
| | 2KVQ | 15490 | 16788 | 0.25 | 0.16 |
| | 2L1R | 30966 | 17103 | 0.35 | 0.16 |
| 2M03 | 6IJQ | 18792 | 36133 | 0.18 | 0.16 |
| | 2MV7 | 18094 | 19516 | 0.38 | 0.15 |
| 1N3H | 2L1C | 5566 | 17080 | 0.34 | 0.15 |
| | 2MZW | 17503 | 25504 | 0.29 | 0.14 |
| | 1LXF | 30966 | 5386 | 0.37 | 0.14 |
| 2JQ6 | 2KFH | 15279 | 16181 | 0.27 | 0.14 |
| | 7ZEY | 16983 | 34727 | 0.44 | 0.14 |
| | 2MRE | 18610 | 25070 | 0.28 | 0.13 |
| | 5JYV | 30092 | 30093 | 0.31 | 0.13 |
| | 9C5E | 18610 | 31177 | 0.32 | 0.13 |
| | 2KNE | 51289 | 16465 | 0.36 | 0.13 |
| 2KJD | 2M0V | 16637 | 18826 | 0.20 | 0.11 |
| | 2JQR | 4566 | 15301 | 0.24 | 0.11 |
| | 2H7D | 15792 | 7150 | 0.24 | 0.11 |
| 1V49 | 2LUE | 5958 | 18518 | 0.22 | 0.11 |
| | 2M86 | 15945 | 19230 | 0.25 | 0.10 |
| | 2MCN | 18610 | 19447 | 0.40 | 0.10 |
| | 2N1A | 6304 | 25553 | 0.10 | 0.10 |
| | 2RSE | 50325 | 11471 | 0.10 | 0.10 |
| | 5XV8 | 4901 | 36101 | 0.37 | 0.09 |
| | 2KA6 | 4789 | 16015 | 0.51 | 0.09 |
| | 2LTO | 10276 | 18490 | 0.14 | 0.09 |
| | 6XMN | 30316 | 50331 | 0.40 | 0.09 |
| | 2MGU | 51289 | 19604 | 0.42 | 0.09 |
| | 2LZ6 | 18610 | 18737 | 0.29 | 0.08 |
| | 2KJE | 4789 | 16318 | 0.45 | 0.07 |
| | 2L53 | 51289 | 17264 | 0.27 | 0.06 |
| | 2MZD | 34231 | 25484 | 0.35 | 0.06 |
| | 2KFT | 6374 | 16878 | 0.52 | 0.06 |
| | 2RSY | 7141 | 11508 | 0.30 | 0.05 |
| | 6BUT | 51289 | 27095 | 0.25 | 0.04 |
| | 2MZP | 30966 | 25495 | 0.17 | 0.04 |
| | 2MKP | 25495 | 19789 | 0.27 | 0.04 |
| | 2LZ6 | 15407 | 18737 | 0.28 | 0.02 |
| | 7NQC | 51289 | 34608 | 0.34 | 0.01 |
| | 7F7X | 17769 | 36427 | 0.20 | 0.00 |
| | 6B1G | 27072 | 30345 | 0.26 | -0.01 |
| 2E5E | 2L7U | 7364 | 17378 | 0.14 | -0.01 |
| 5YDX | 2LAY | 36114 | 17540 | 0.29 | -0.01 |
| | 2N8J | 51289 | 25852 | 0.19 | -0.01 |
| | 2M55 | 51289 | 19036 | 0.24 | -0.01 |
| | 2RMK | 5511 | 11010 | 0.18 | -0.02 |
| | 2NBV | 30824 | 25995 | 0.15 | -0.02 |
| | 2MWN | 15792 | 25343 | 0.17 | -0.03 |
| | 5TP6 | 51289 | 30196 | 0.21 | -0.03 |
| | 1NPQ | 4232 | 5738 | 0.24 | -0.03 |
| | 2MNU | 4283 | 19906 | 0.19 | -0.04 |
| | 2LV6 | 51289 | 18556 | 0.35 | -0.07 |
| | 2L29 | 34000 | 17127 | 0.16 | -0.08 |
| | 2N83 | 25883 | 25833 | 0.29 | -0.10 |
| | 2N80 | 51835 | 25829 | 0.04 | -0.13 |
| | 2JMF | 6262 | 15016 | 0.13 | -0.17 |
| | 2RS9 | 19125 | 11463 | 0.00 | -0.17 |

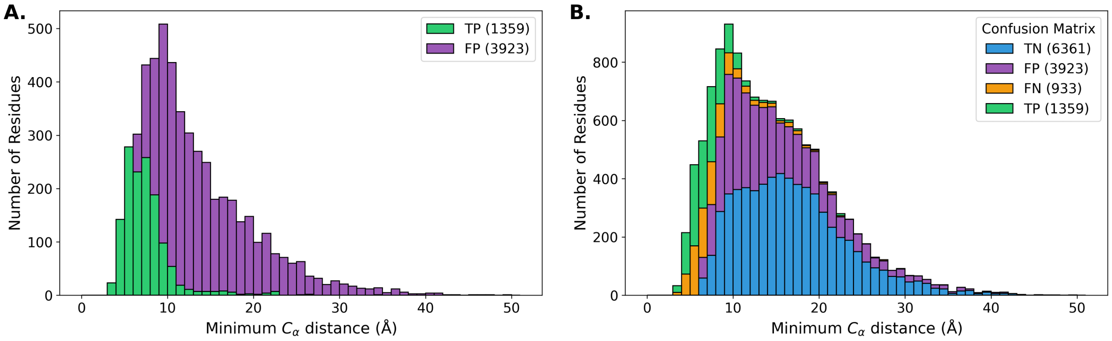


**SI Fig. S9. Confusion Matrix Histograms for targets with dissimilar apo/holo experimental conditions**

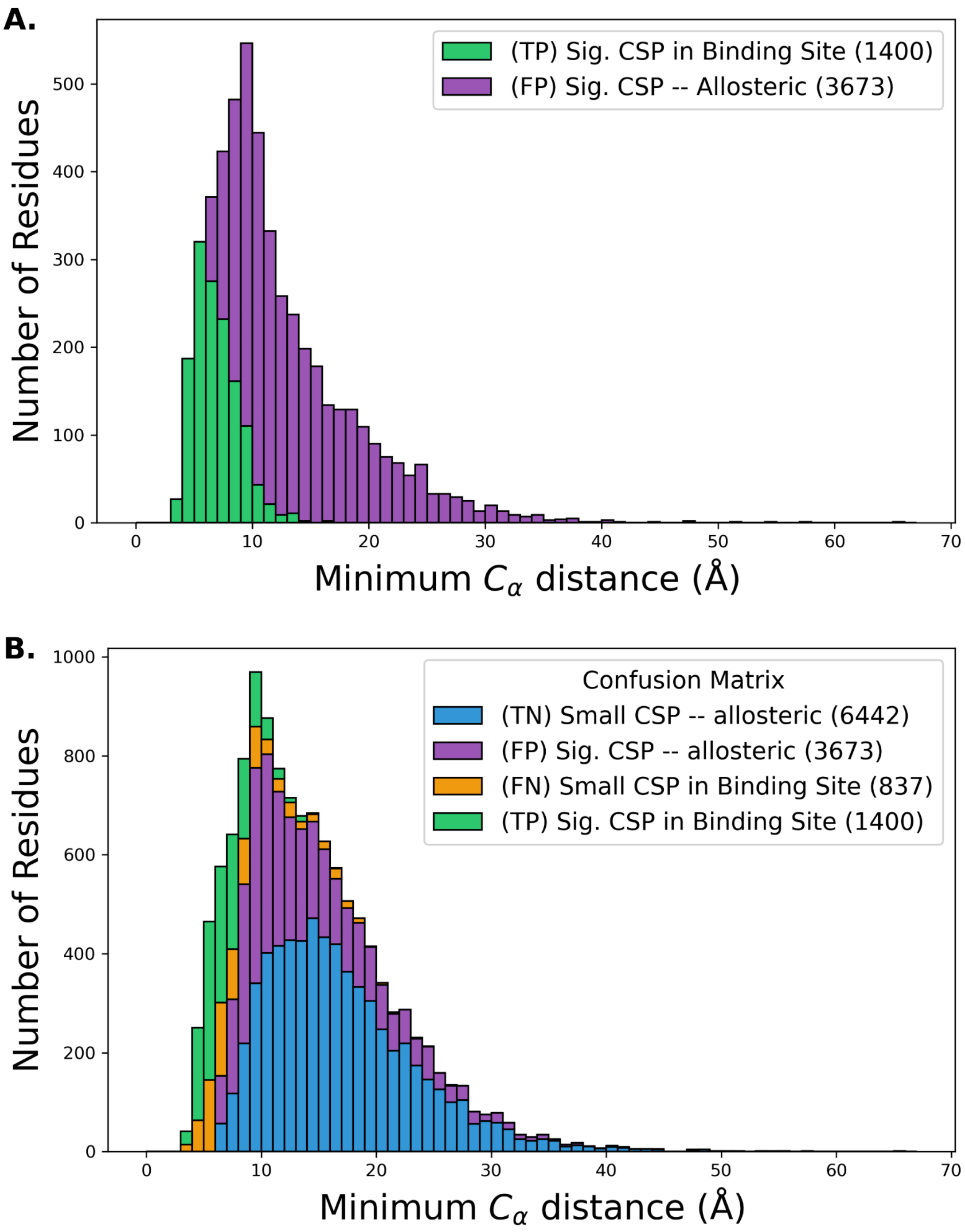
A.
(TP) Sig. CSP in Binding Site (1400)
(FP) Sig. CSP -- Allosteric (3673)
Number of Residues
Minimum $C_\alpha$ distance (Å)
B.
Confusion Matrix
(TN) Small CSP -- allosteric (6442)
(FP) Sig. CSP -- allosteric (3673)
(FN) Small CSP in Binding Site (837)
(TP) Sig. CSP in Binding Site (1400)
Number of Residues
Minimum $C_\alpha$ distance (Å)

**SI Fig. S10**. **CA-inclusive CSP Confusion Matrix Histograms**. Calculating CSPs with CAs included in the formula does drastically not change the percentage of allosteric CSPs ~72%. CSPs are calculated using formula described in **Eqn. 2**.

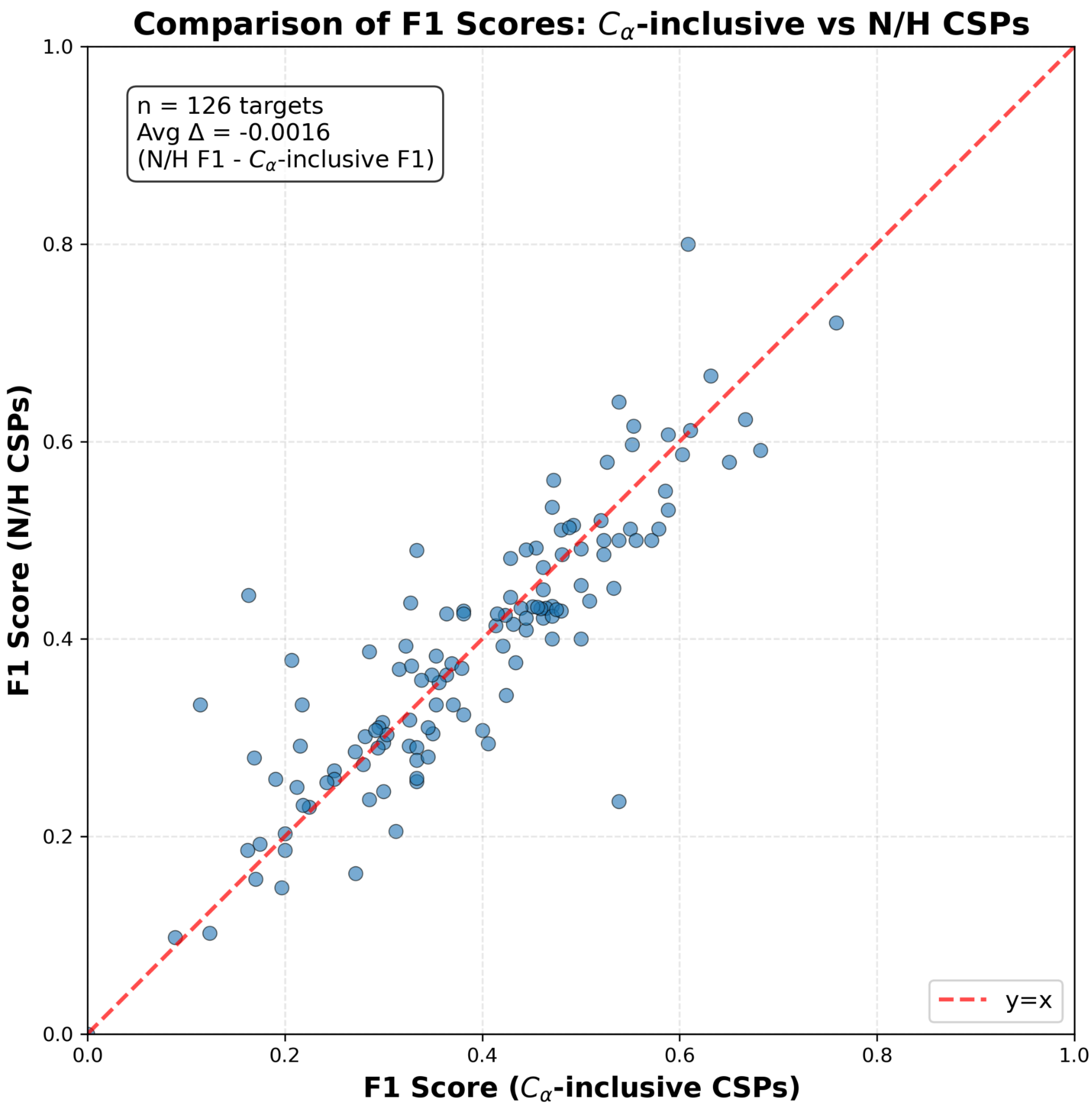


**SI Fig. S11. Change in F1 scores when considering CA-inclusive CSPs**. This plot shows that over the targets in the CSP DB, the average change in F1 score is negligible when comparing CSPs calculated from N/H shifts vs CSPs calculated with N/H/Ca shifts. CA-inclusive CSPs are calculated following **Eqn. 2**.

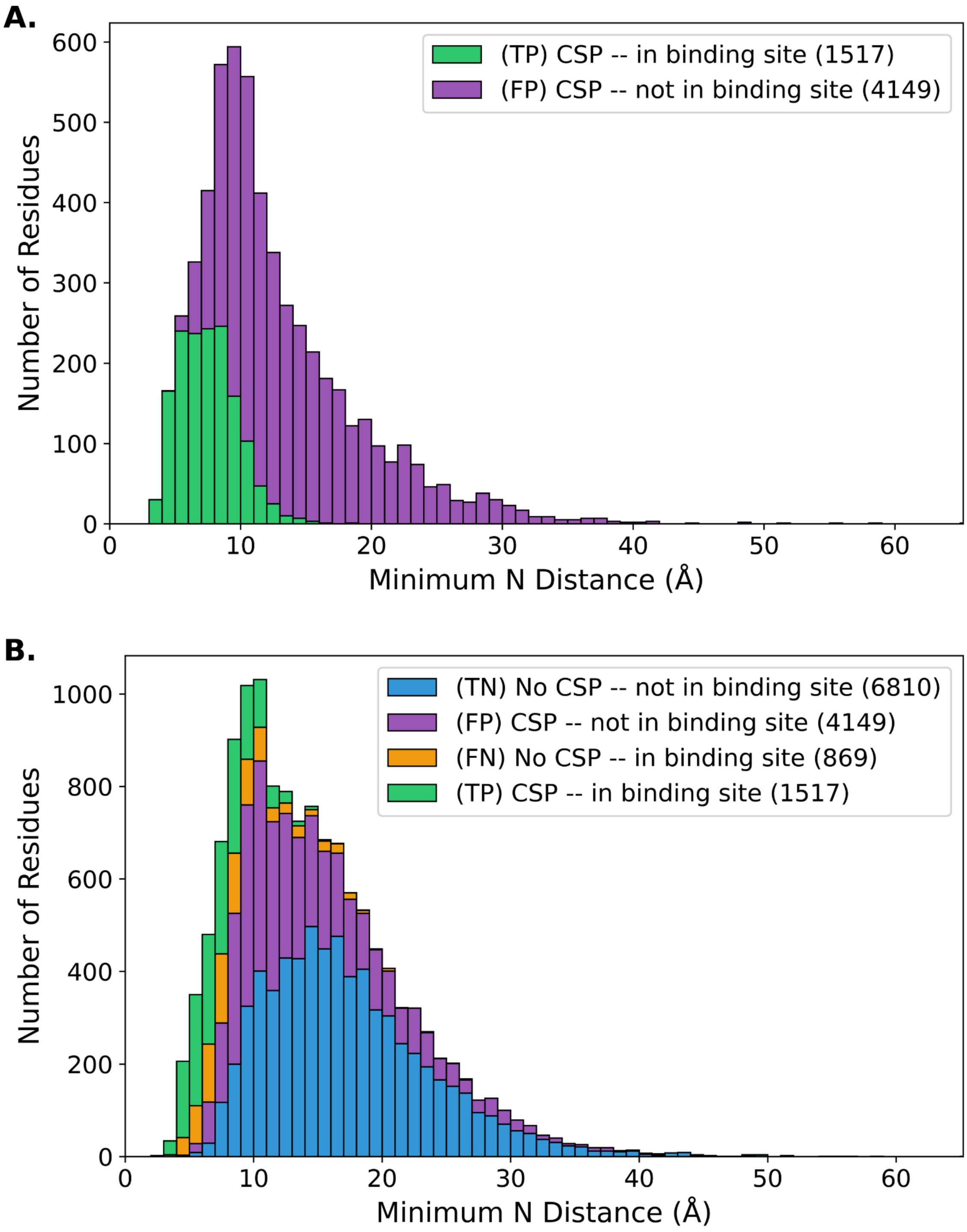


**SI Fig. S12. CSP DB Confusion Matrix Histograms by closest interchain N-N distance.**

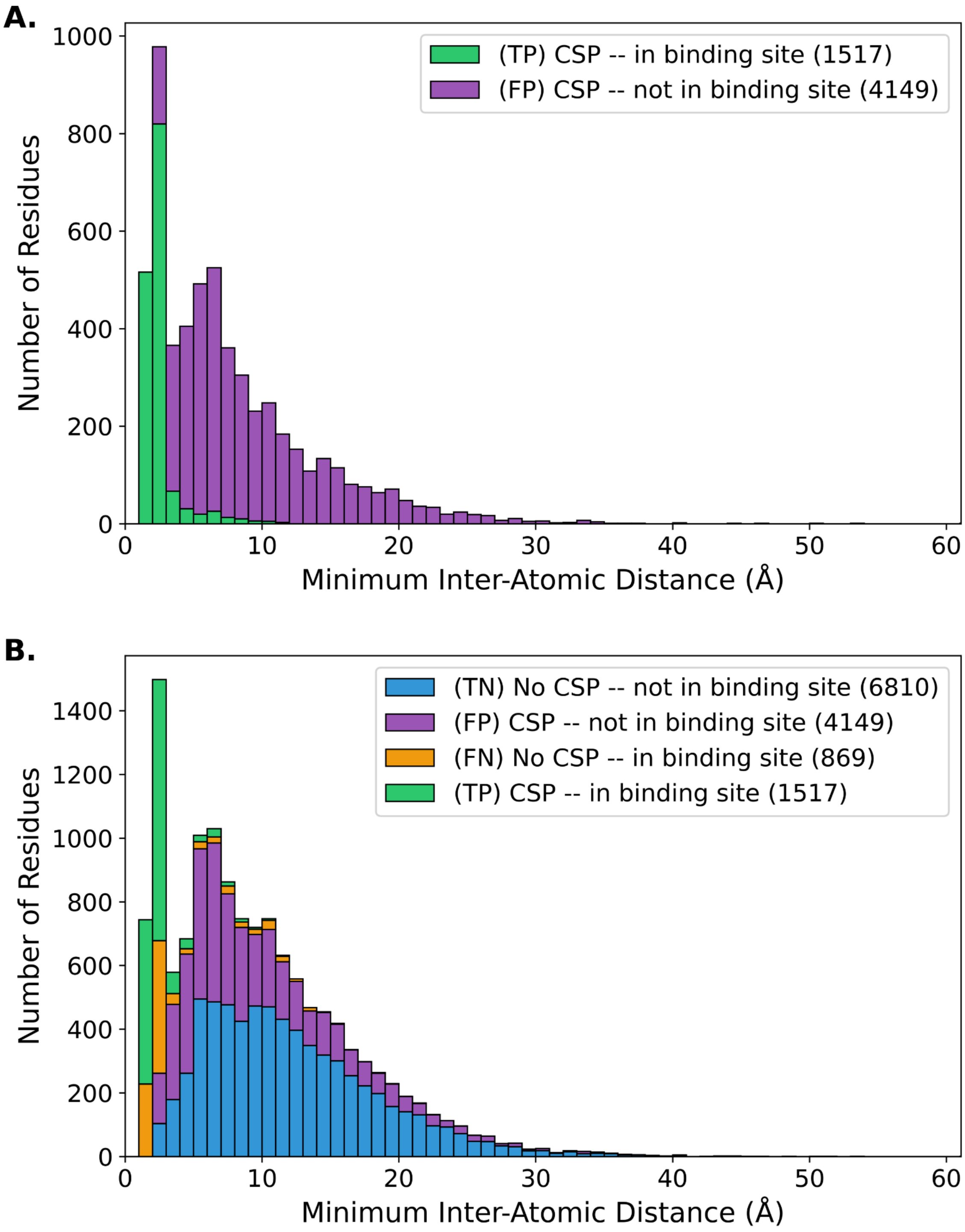


**SI Fig. S13. CSP DB Confusion Matrix Histograms by closest interchain atom-atom distance.**

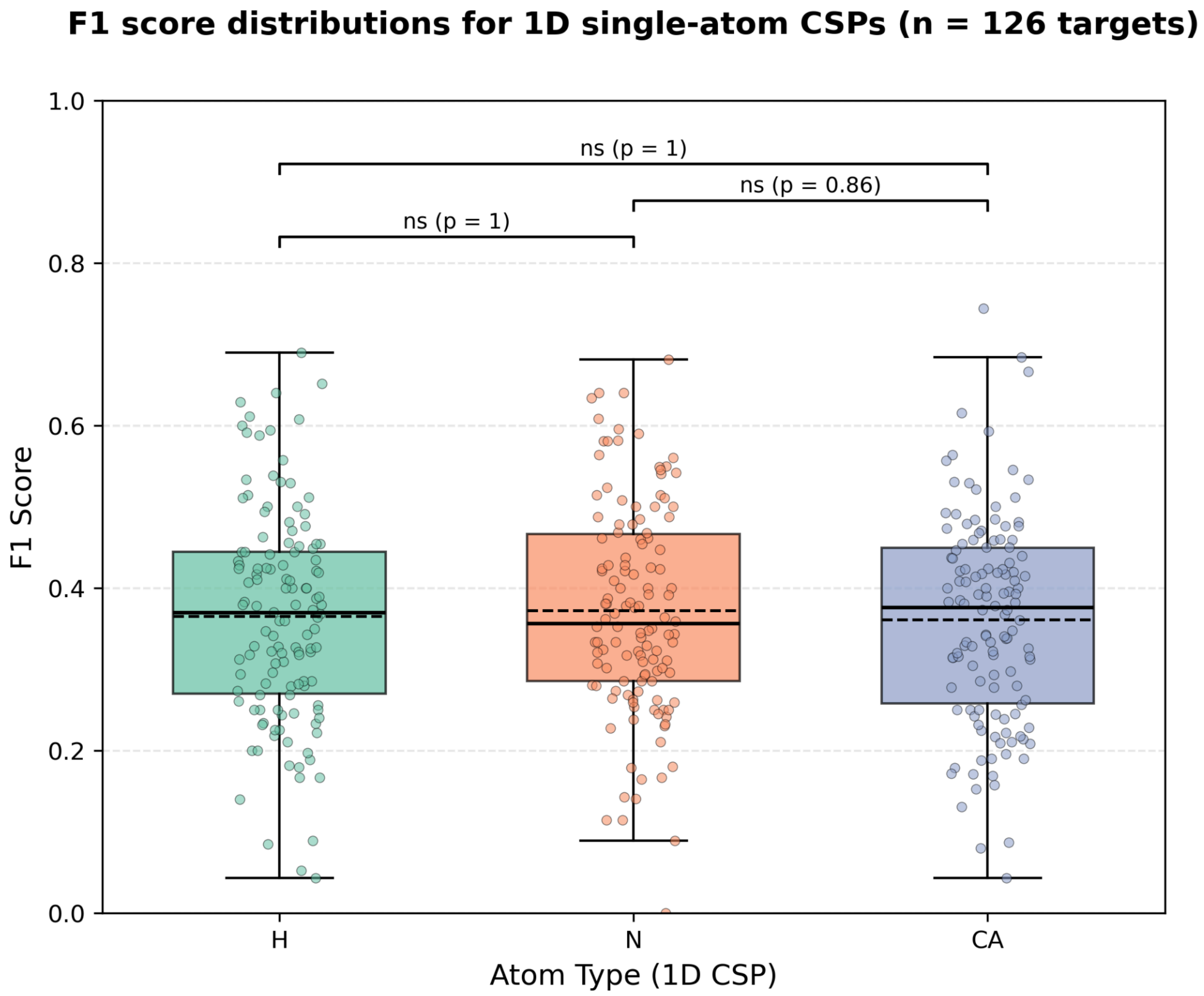


**SI Fig. S14. F1 Scores for 1D CSPs.** This plot shows there are no statistical differences between distributions of calculated F1 scores from 1D CSPs H/N/CA atom types. CSPs are calculated as root-squared differences after applying an ideal holo offset for a 3D H/N/CA offset grid (see **Methods** for details). Statistical analysis is performed using paired Wilcoxon tests with Holm correction.

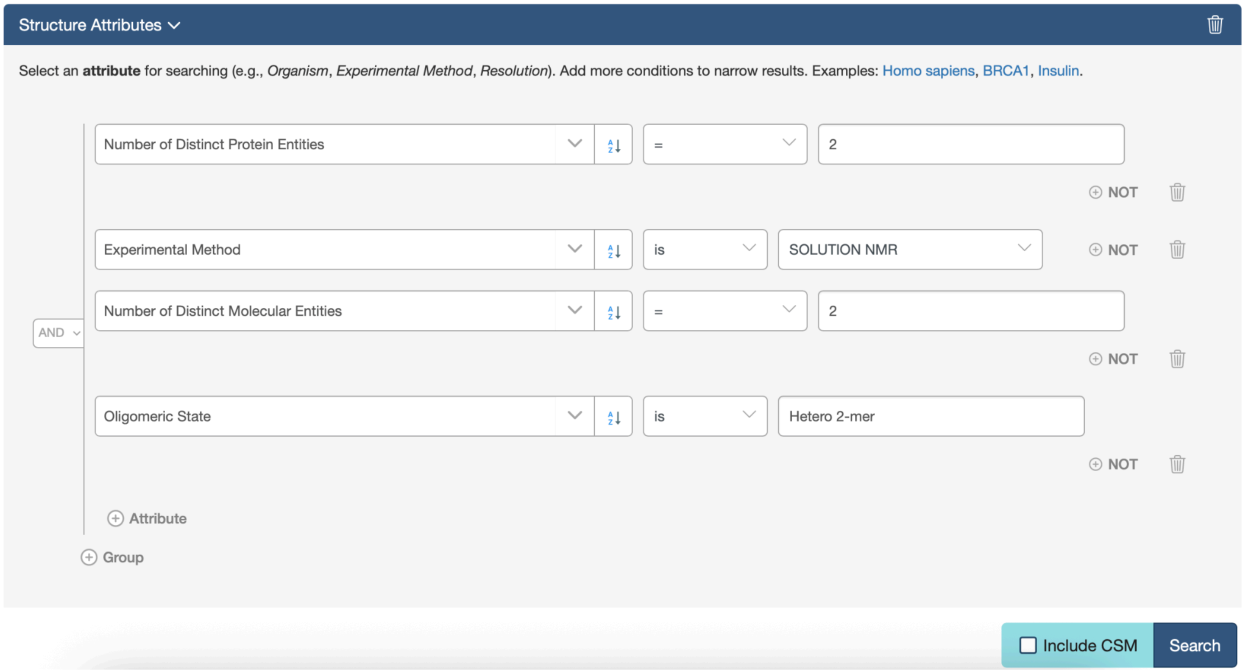


**SI Fig. S15. PDB Advanced Search Settings.** For the subset of targets manually scraped from the PDB, we restrict our analysis to these settings: entries resolved by Solution NMR, with two and only two distinct protein chains. At the time of curation, this advanced search resulted in 720 hits on the RCSB PDB. From this list of 720, entries were included in the CSP DB if there were N and H chemical shifts deposited in the BMRB for an identical sequence which was evaluated in both apo and holo experiments.

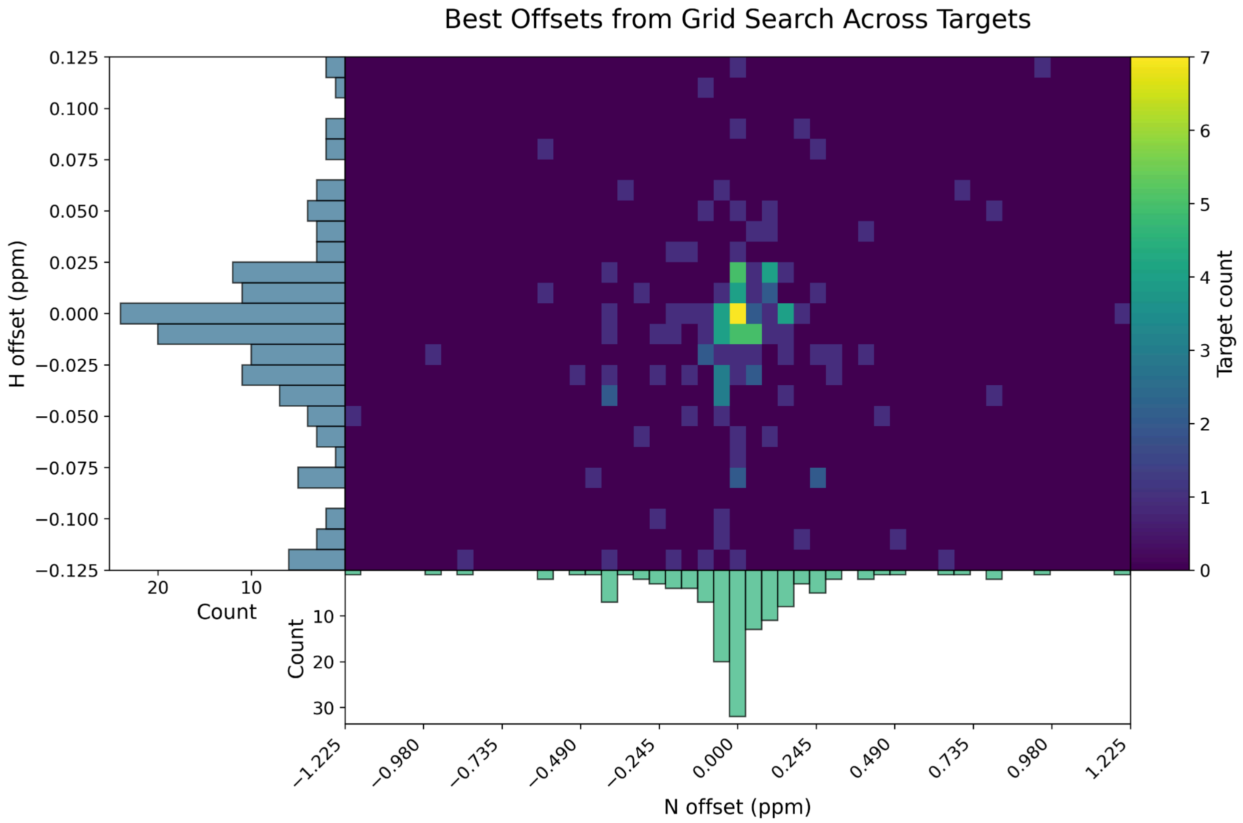


**SI Fig. S16**. **Ideal offsets across the whole dataset for N/H**. This heatmap shows the offsets used to align the apo/holo HSQC plots. The majority of targets require small adjustments within 0.025 ppm on the Hydrogen axis and 0.245 ppm on the Nitrogen axis.

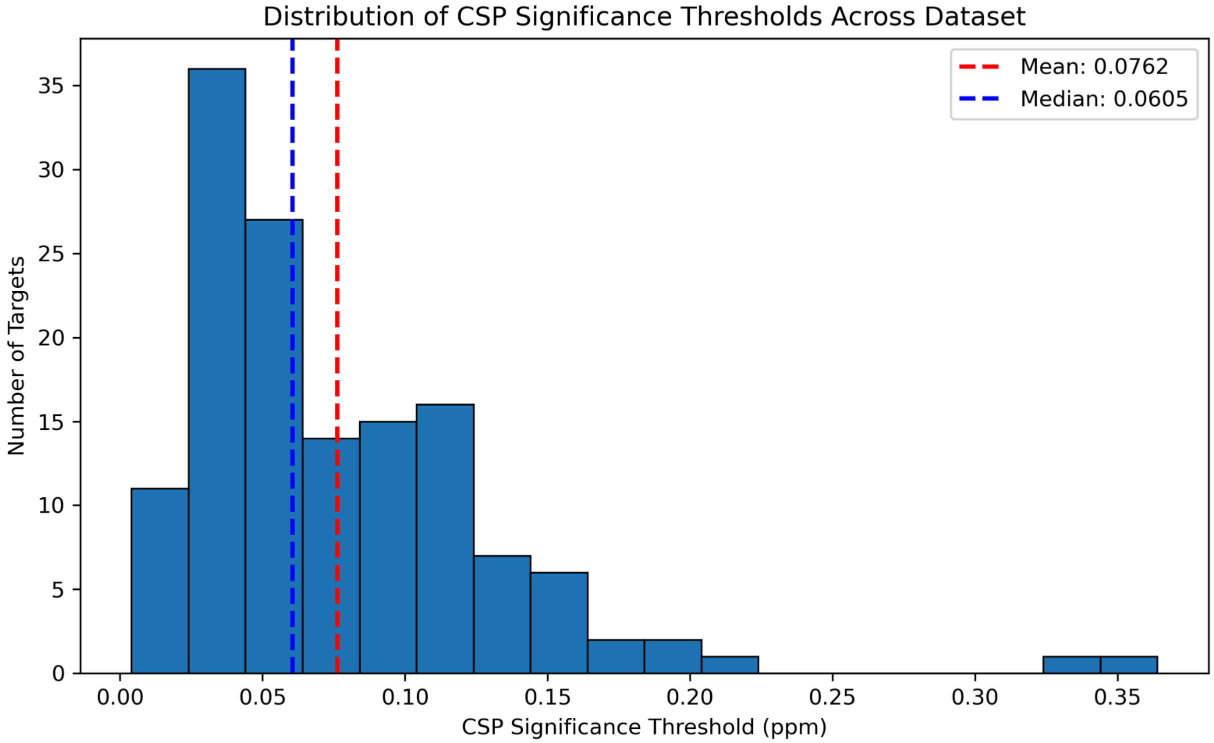


**SI Fig. S17. CSP significance threshold Histogram.** This histogram summarizes the thresholds for determining whether a CSP for a given receptor is significant. For a small number of targets, the calculated significance threshold is significantly higher than one would expect, we attribute this to differences in experimental conditions between the apo and holo NMR experiments which aberrantly perturbs all observed shifts. We prune from the CSPdb any targets with a calculated significance threshold > 0.6 ppm. This includes the following targets: 2LMC, 1OO9, 2L12, 2L00, 2N01, 2MP0, 8VOI, 8U2Y, 6OSW.

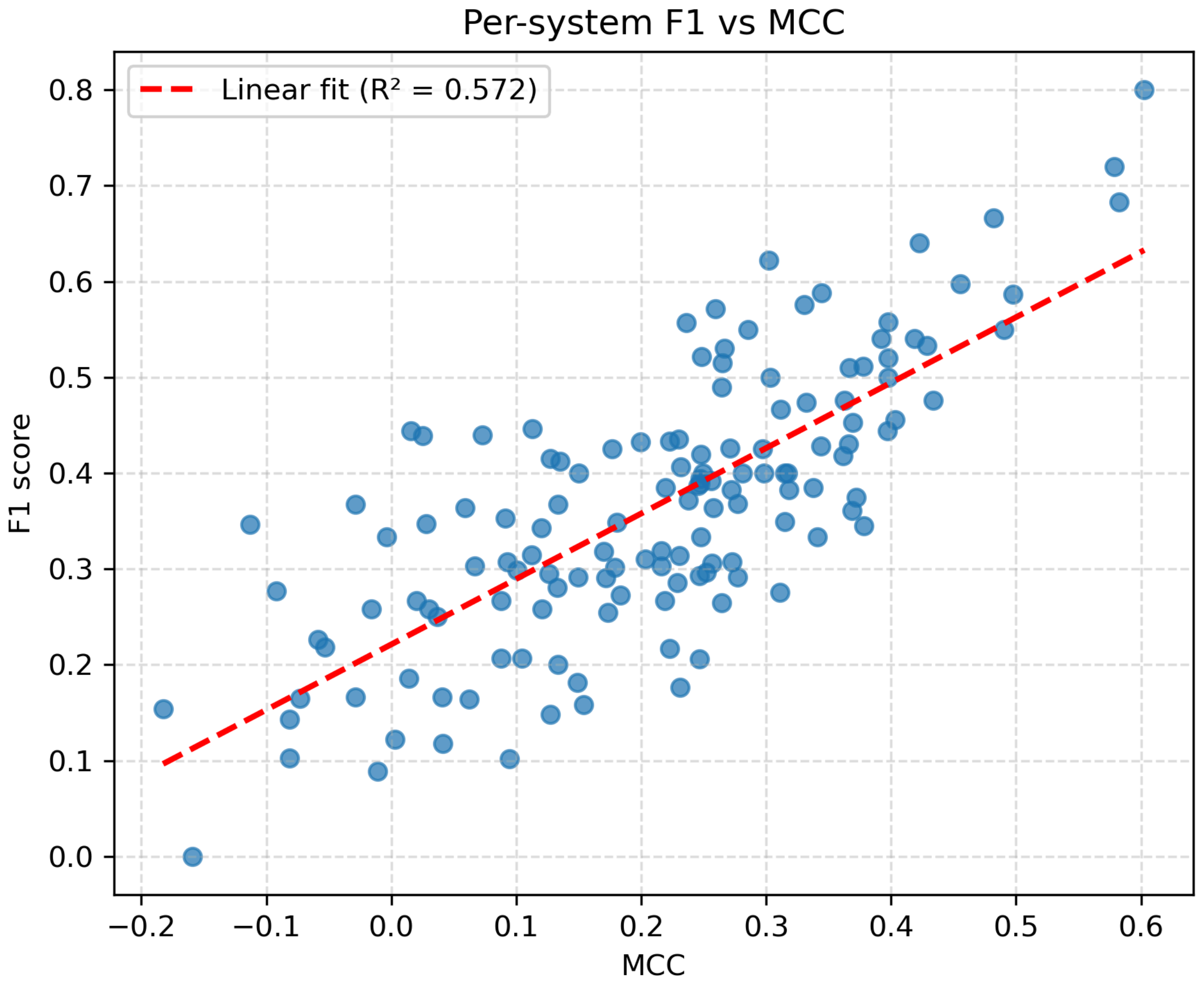


**SI Fig. S18. F1 vs MCC score comparison across CSP DB targets.** This plot shows the correlation between calculated F1 and MCC metrics. MCC is a more reliable metric for larger systems with many True Negatives (small CSPs outside of the binding site) (7).

**References**


1. N. Juranic, *et al.*, Calmodulin Wraps around Its Binding Domain in the Plasma Membrane Ca2+ Pump Anchored by a Novel 18-1 Motif. *J. Biol. Chem.* **285**, 4015–4024 (2010).

2. A. Mondal, *et al.*, Structure Determination of Challenging Protein–Peptide Complexes Combining NMR Chemical Shift Data and Molecular Dynamics Simulations. *J. Chem. Inf. Model.* **63**, 2058–2072 (2023).

3. G. Alvarez, *et al.*, Loop Plasticity Drives Paralog-Specific Recognition in BET ET Domains. *J. Chem. Inf. Model.* **66**, 4685–4695 (2026).

4. S. Gianni, *et al.*, Demonstration of Long-Range Interactions in a PDZ Domain by NMR, Kinetics, and Protein Engineering. *Structure* **14**, 1801–1809 (2006).

5. R. Agrata, D. Komander, Ubiquitin—A structural perspective. *Mol. Cell* **85**, 323–346 (2025).

6. A. Hacisuleyman, B. Erman, Entropy Transfer between Residue Pairs and Allostery in Proteins: Quantifying Allosteric Communication in Ubiquitin. *PLOS Comput. Biol.* **13**, e1005319 (2017).

7. D. Chicco, M. J. Warrens, G. Jurman, The Matthews Correlation Coefficient (MCC) is More Informative Than Cohen's Kappa and Brier Score in Binary Classification Assessment. *IEEE Access* **9**, 78368–78381 (2021).
