## Supplementary Information 2 - All cases for "Allosteric Protein Chemical Shift Perturbations are Ubiquitous"

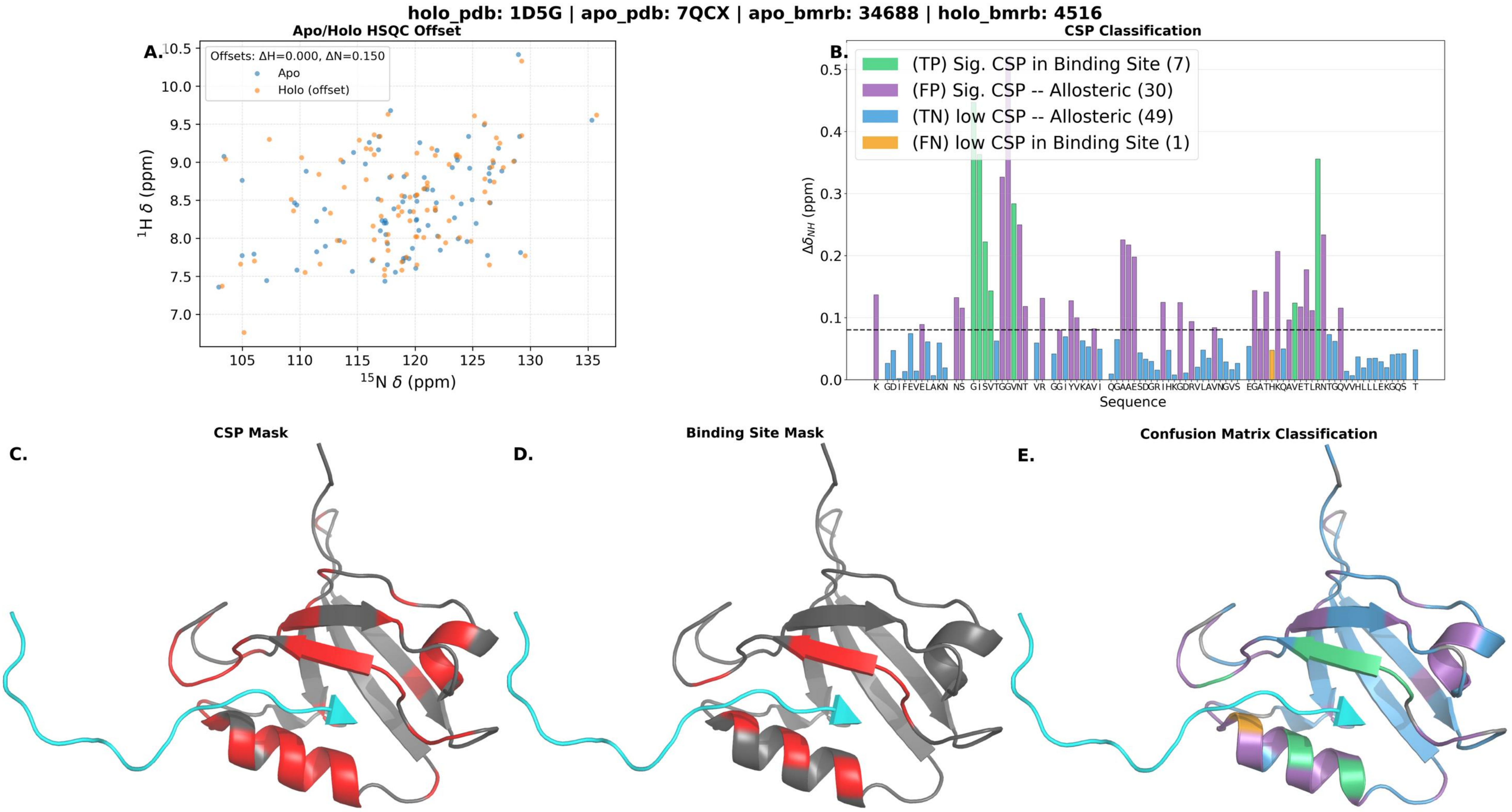

holo_pdb: 1D5G | apo_pdb: 7QCX | apo_bmrb: 34688 | holo_bmrb: 4516
A.
Apo/Holo HSQC Offset
Offsets: ΔH=0.000, ΔN=0.150
Apo
Holo (offset)
$^{1}$H δ (ppm)
$^{15}$N δ (ppm)
B.
CSP Classification
(TP) Sig. CSP in Binding Site (7)
(FP) Sig. CSP -- Allosteric (30)
(TN) low CSP -- Allosteric (49)
(FN) low CSP in Binding Site (1)
$\Delta\delta_{NH}$ (ppm)
Sequence
C.
CSP Mask
D.
Binding Site Mask
E.
Confusion Matrix Classification

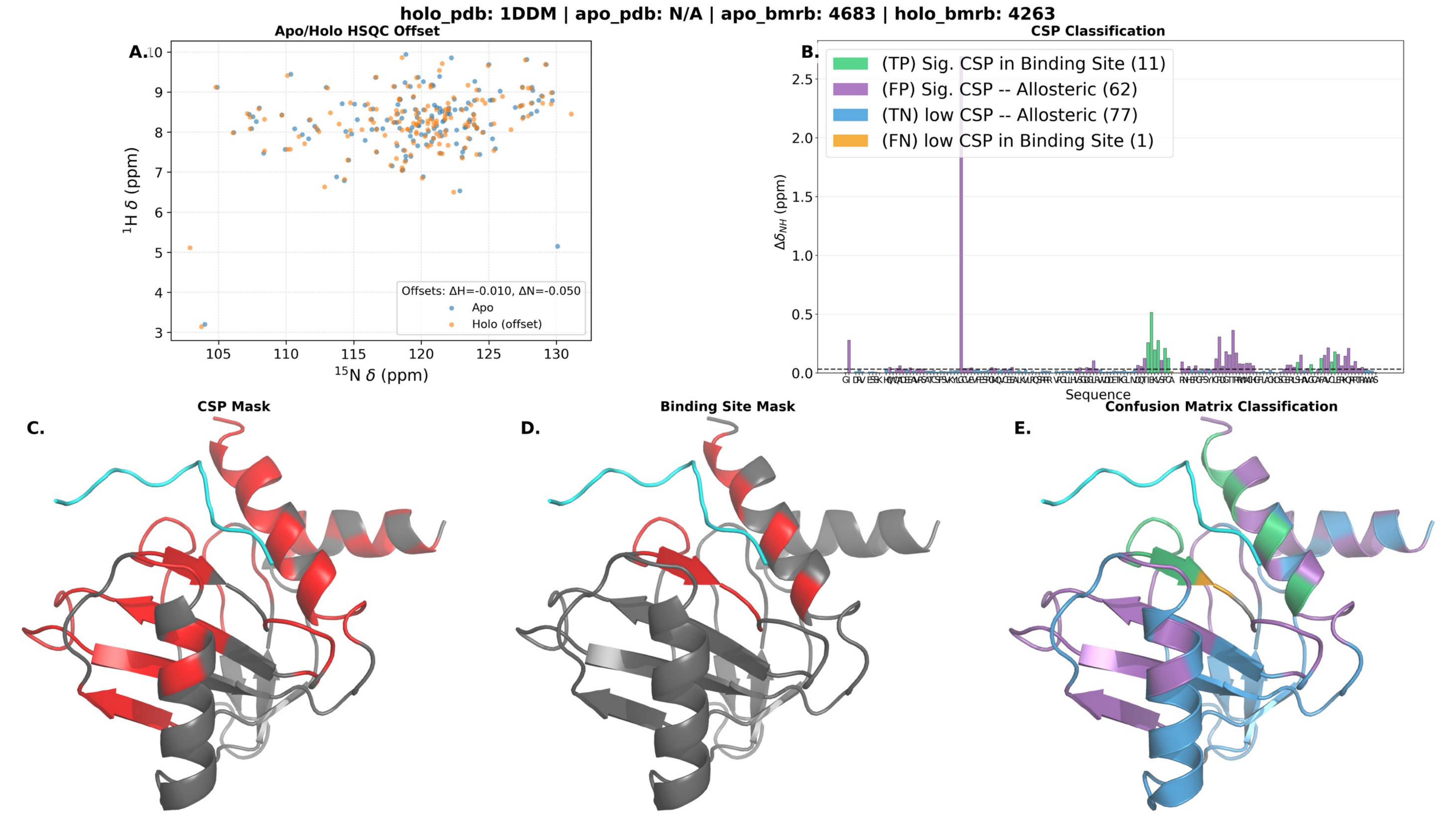
holo_pdb: 1DDM | apo_pdb: N/A | apo_bmrb: 4683 | holo_bmrb: 4263
A.
Apo/Holo HSQC Offset
$^{1}$H $\delta$ (ppm)
$^{15}$N $\delta$ (ppm)
Offsets: ΔH=-0.010, ΔN=-0.050
Apo
Holo (offset)
B.
CSP Classification
(TP) Sig. CSP in Binding Site (11)
(FP) Sig. CSP -- Allosteric (62)
(TN) low CSP -- Allosteric (77)
(FN) low CSP in Binding Site (1)
$\Delta\delta_{NH}$ (ppm)
Sequence
C.
CSP Mask
D.
Binding Site Mask
E.
Confusion Matrix Classification

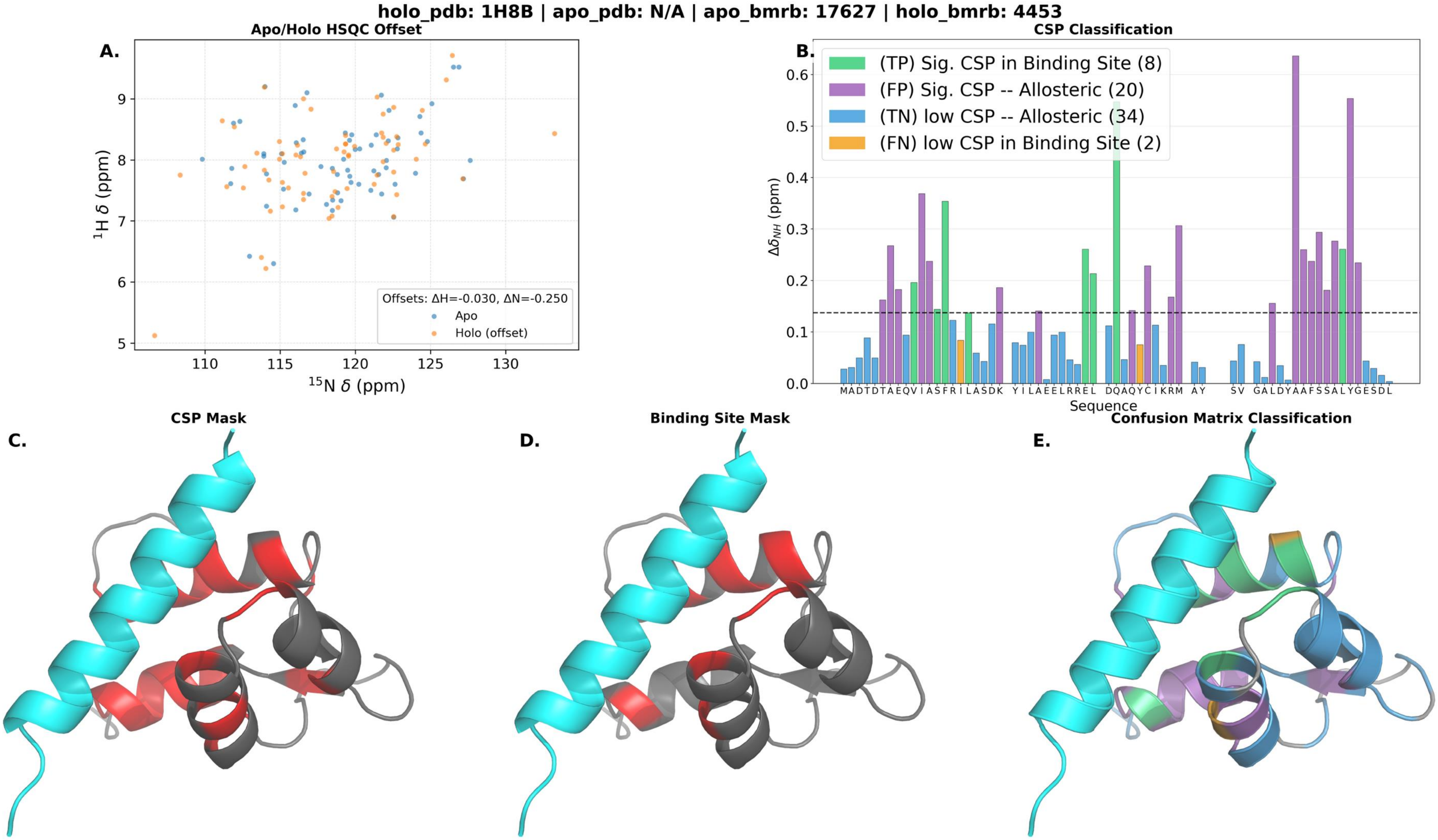

holo_pdb: 1H8B | apo_pdb: N/A | apo_bmrb: 17627 | holo_bmrb: 4453
A.
Apo/Holo HSQC Offset
$^{1}$H δ (ppm)
$^{15}$N δ (ppm)
Offsets: ΔH=-0.030, ΔN=-0.250
Apo
Holo (offset)
B.
CSP Classification
(TP) Sig. CSP in Binding Site (8)
(FP) Sig. CSP -- Allosteric (20)
(TN) low CSP -- Allosteric (34)
(FN) low CSP in Binding Site (2)
$\Delta\delta_{NH}$ (ppm)
Sequence
C.
CSP Mask
D.
Binding Site Mask
E.
Confusion Matrix Classification

**holo_pdb: 1KLQ | apo_pdb: N/A | apo_bmrb: 52275 | holo_bmrb: 5299**

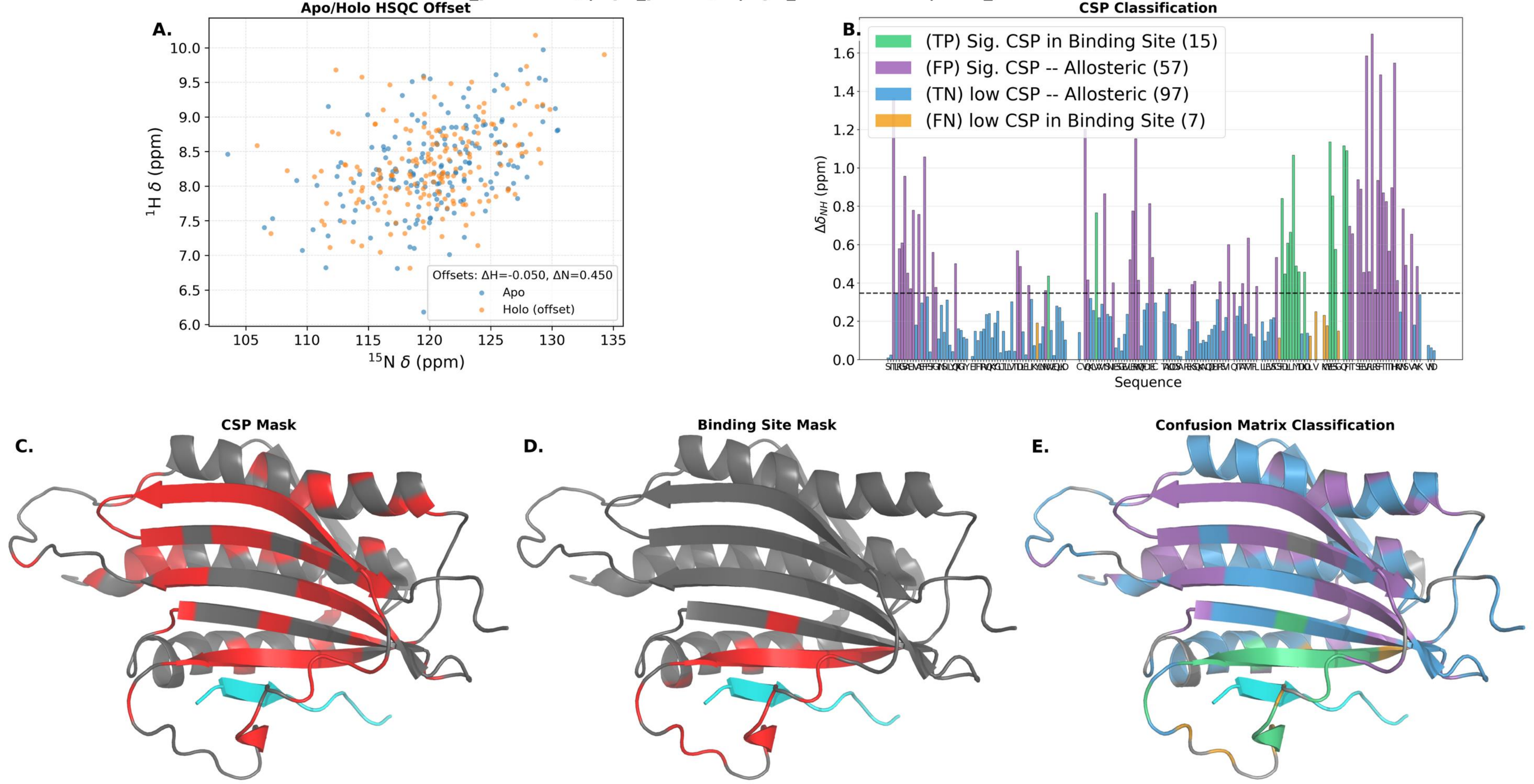

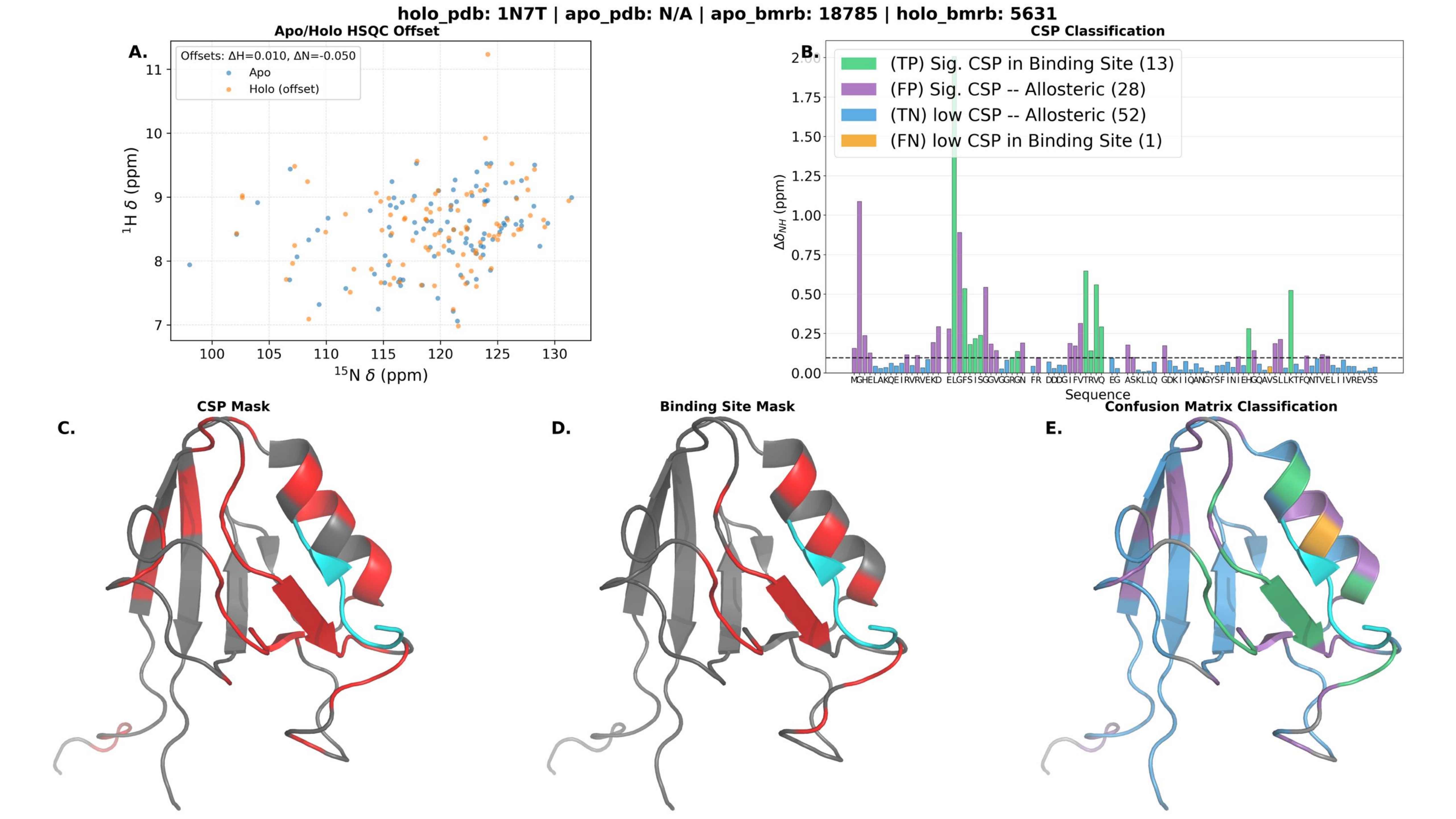
holo_pdb: 1N7T | apo_pdb: N/A | apo_bmrb: 18785 | holo_bmrb: 5631
Apo/Holo HSQC Offset
A.
Offsets: ΔH=0.010, ΔN=-0.050
Apo
Holo (offset)
$^{1}$H δ (ppm)
$^{15}$N δ (ppm)
CSP Classification
B.
(TP) Sig. CSP in Binding Site (13)
(FP) Sig. CSP -- Allosteric (28)
(TN) low CSP -- Allosteric (52)
(FN) low CSP in Binding Site (1)
$\Delta\delta_{NH}$ (ppm)
Sequence
CSP Mask
C.
Binding Site Mask
D.
Confusion Matrix Classification
E.

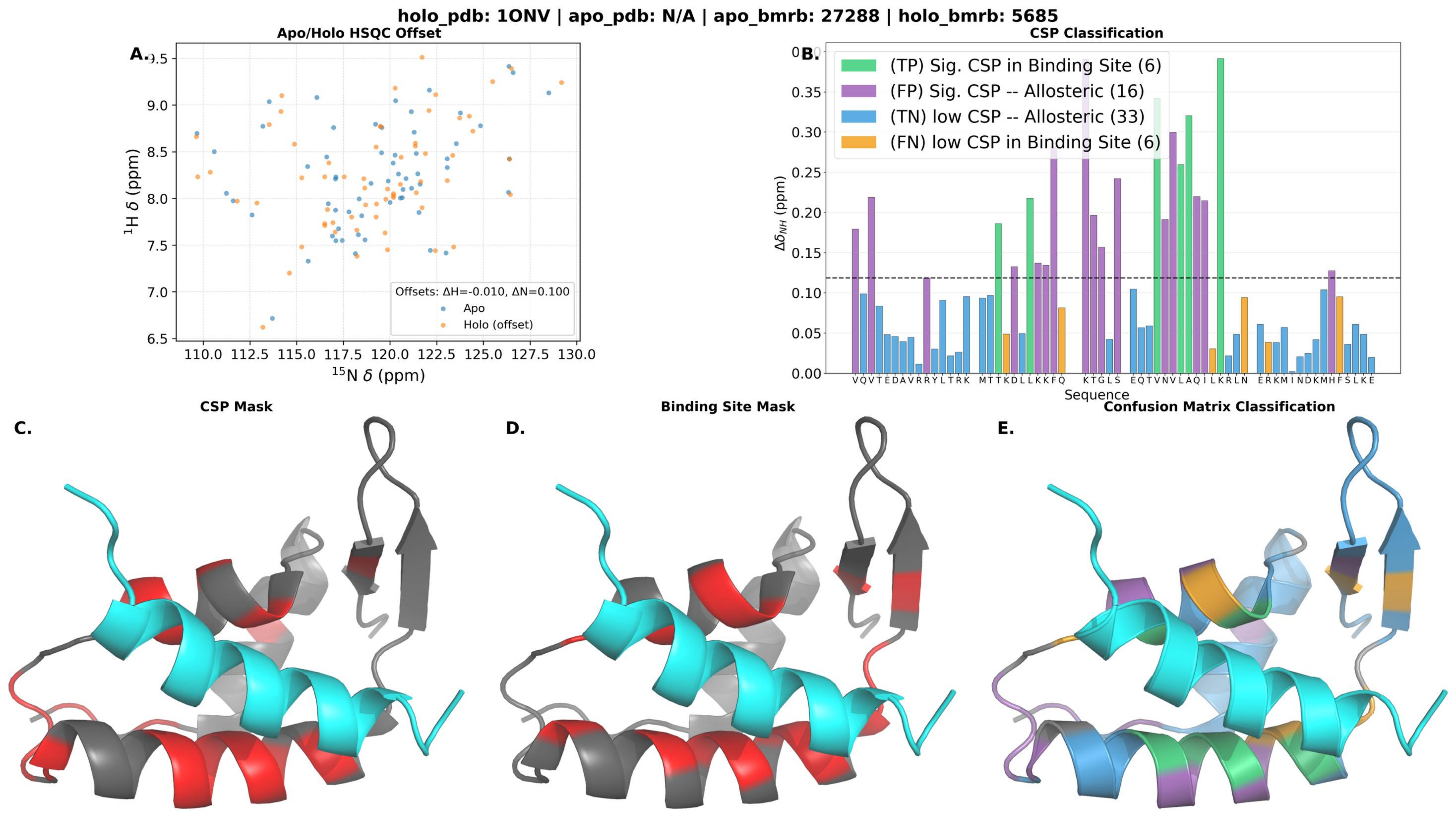
holo_pdb: 1ONV | apo_pdb: N/A | apo_bmrb: 27288 | holo_bmrb: 5685
A.
Apo/Holo HSQC Offset
Offsets: ΔH=-0.010, ΔN=0.100
Apo
Holo (offset)
$^{1}$H δ (ppm)
$^{15}$N δ (ppm)
B.
CSP Classification
(TP) Sig. CSP in Binding Site (6)
(FP) Sig. CSP -- Allosteric (16)
(TN) low CSP -- Allosteric (33)
(FN) low CSP in Binding Site (6)
$\Delta\delta_{NH}$ (ppm)
Sequence
C.
CSP Mask
D.
Binding Site Mask
E.
Confusion Matrix Classification

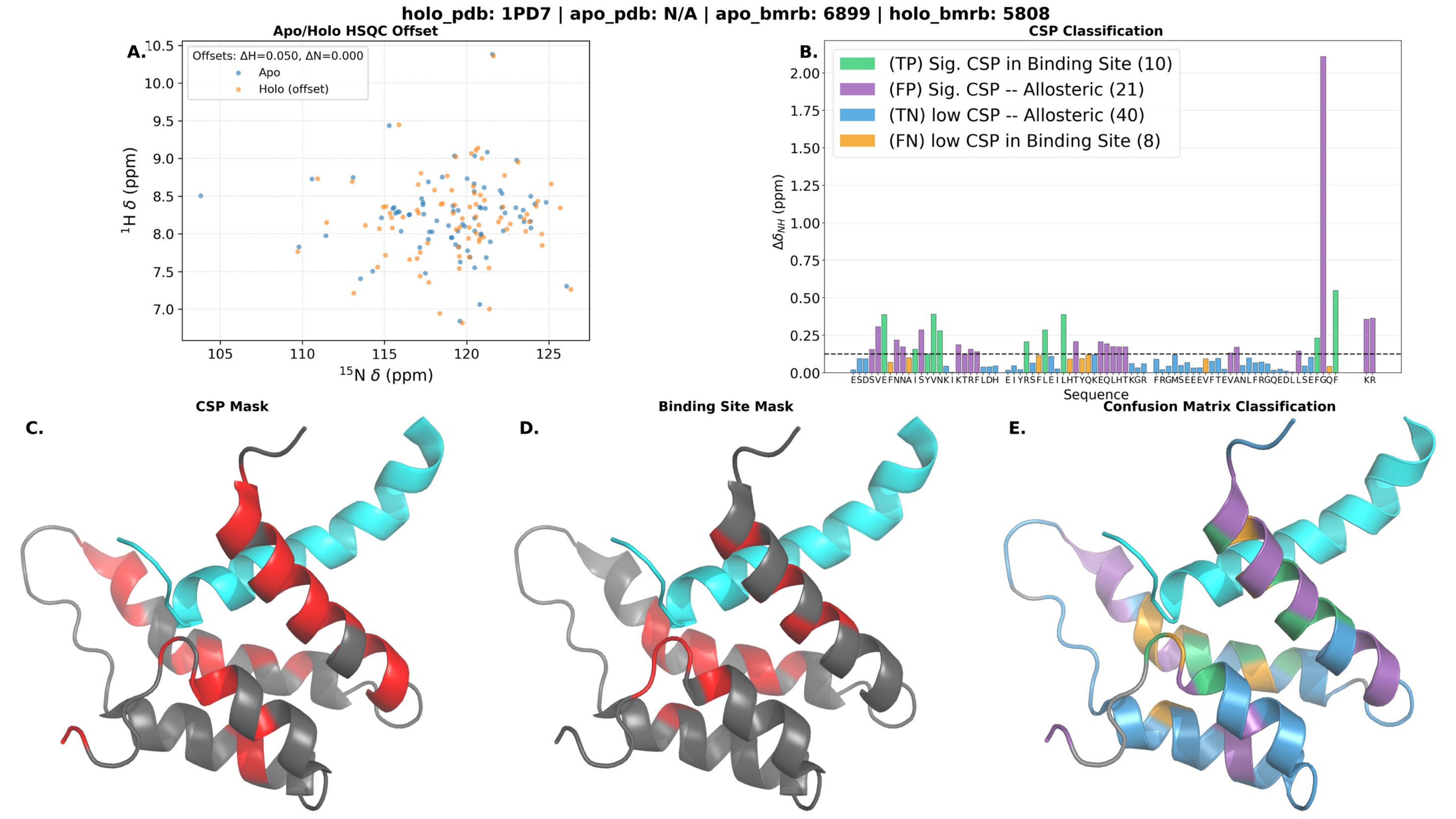

holo_pdb: 1PD7 | apo_pdb: N/A | apo_bmrb: 6899 | holo_bmrb: 5808
A.
Apo/Holo HSQC Offset
Offsets: ΔH=0.050, ΔN=0.000
Apo
Holo (offset)
1H δ (ppm)
15N δ (ppm)
B.
CSP Classification
(TP) Sig. CSP in Binding Site (10)
(FP) Sig. CSP -- Allosteric (21)
(TN) low CSP -- Allosteric (40)
(FN) low CSP in Binding Site (8)
ΔδNH (ppm)
Sequence
C.
CSP Mask
D.
Binding Site Mask
E.
Confusion Matrix Classification

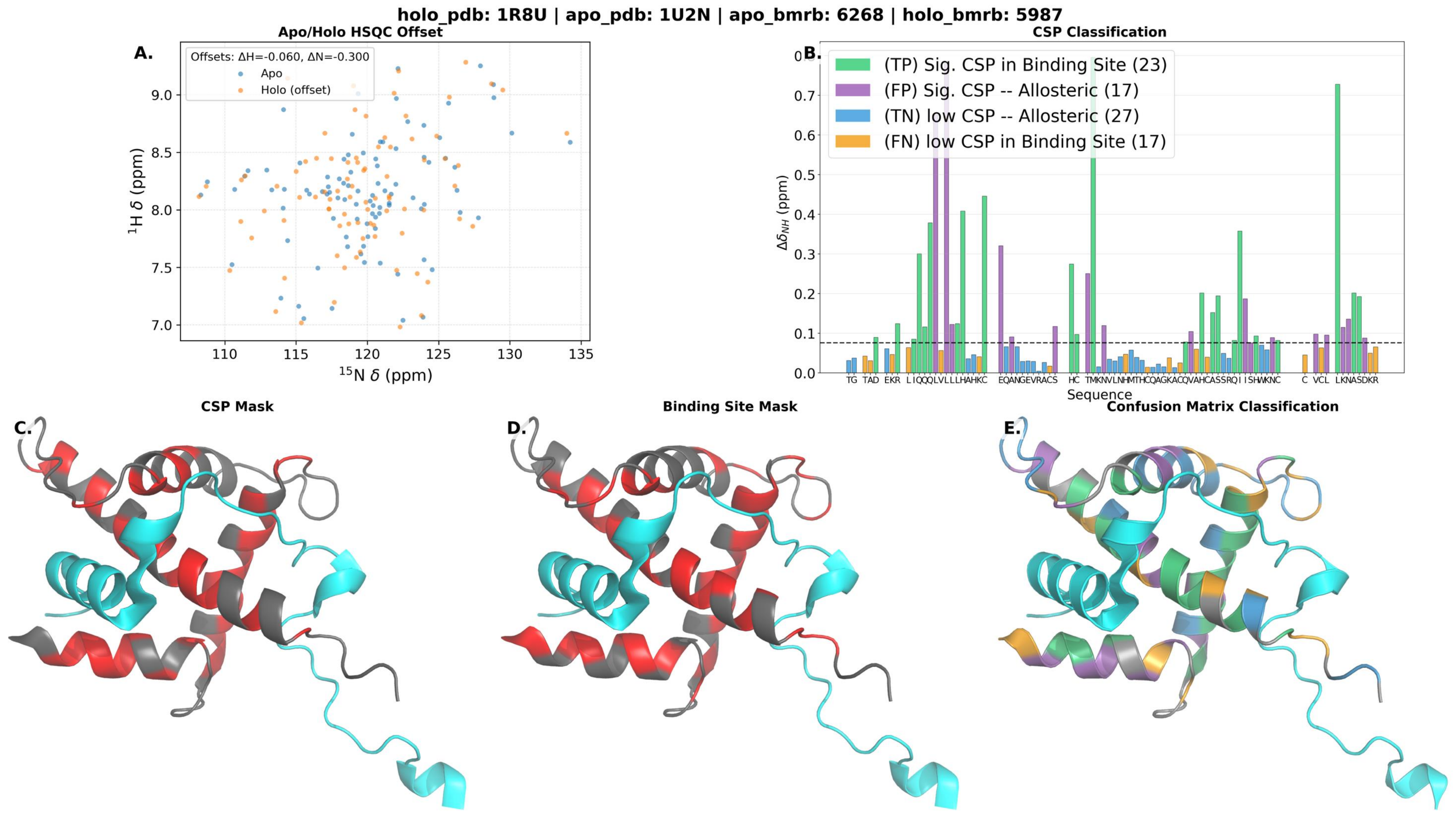
holo_pdb: 1R8U | apo_pdb: 1U2N | apo_bmrb: 6268 | holo_bmrb: 5987
A.
Apo/Holo HSQC Offset
Offsets: ΔH=-0.060, ΔN=-0.300
Apo
Holo (offset)
$^{1}$H δ (ppm)
$^{15}$N δ (ppm)
B.
CSP Classification
(TP) Sig. CSP in Binding Site (23)
(FP) Sig. CSP -- Allosteric (17)
(TN) low CSP -- Allosteric (27)
(FN) low CSP in Binding Site (17)
$\Delta\delta_{NH}$ (ppm)
Sequence
C.
CSP Mask
D.
Binding Site Mask
E.
Confusion Matrix Classification

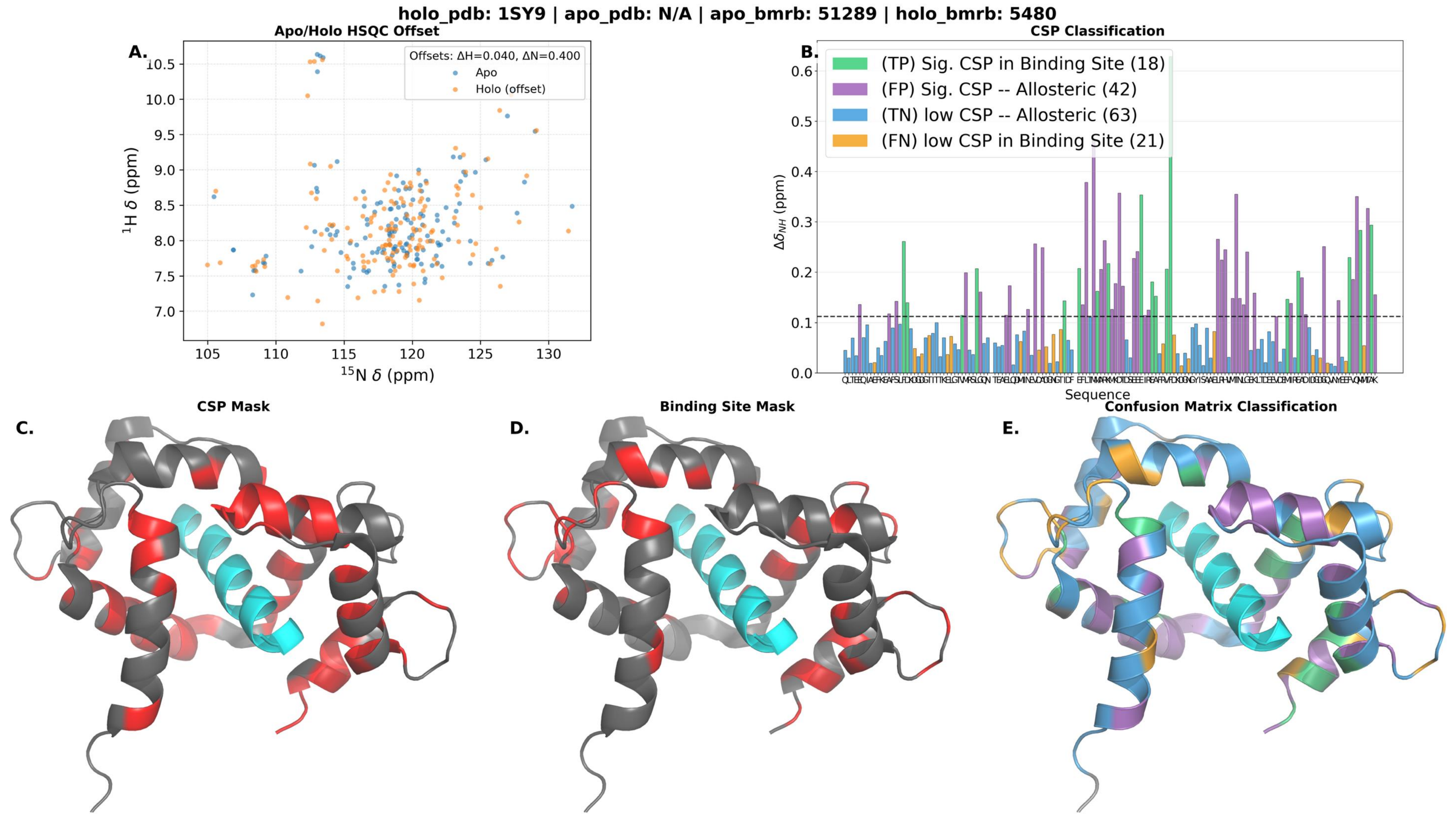

holo_pdb: 1SY9 | apo_pdb: N/A | apo_bmrb: 51289 | holo_bmrb: 5480
A.
Apo/Holo HSQC Offset
Offsets: ΔH=0.040, ΔN=0.400
Apo
Holo (offset)
1H δ (ppm)
15N δ (ppm)
B.
CSP Classification
(TP) Sig. CSP in Binding Site (18)
(FP) Sig. CSP -- Allosteric (42)
(TN) low CSP -- Allosteric (63)
(FN) low CSP in Binding Site (21)
Δδ_NH (ppm)
Sequence
C.
CSP Mask
D.
Binding Site Mask
E.
Confusion Matrix Classification

**holo_pdb: 1VJ6 | apo_pdb: N/A | apo_bmrb: 5131 | holo_bmrb: 6060**

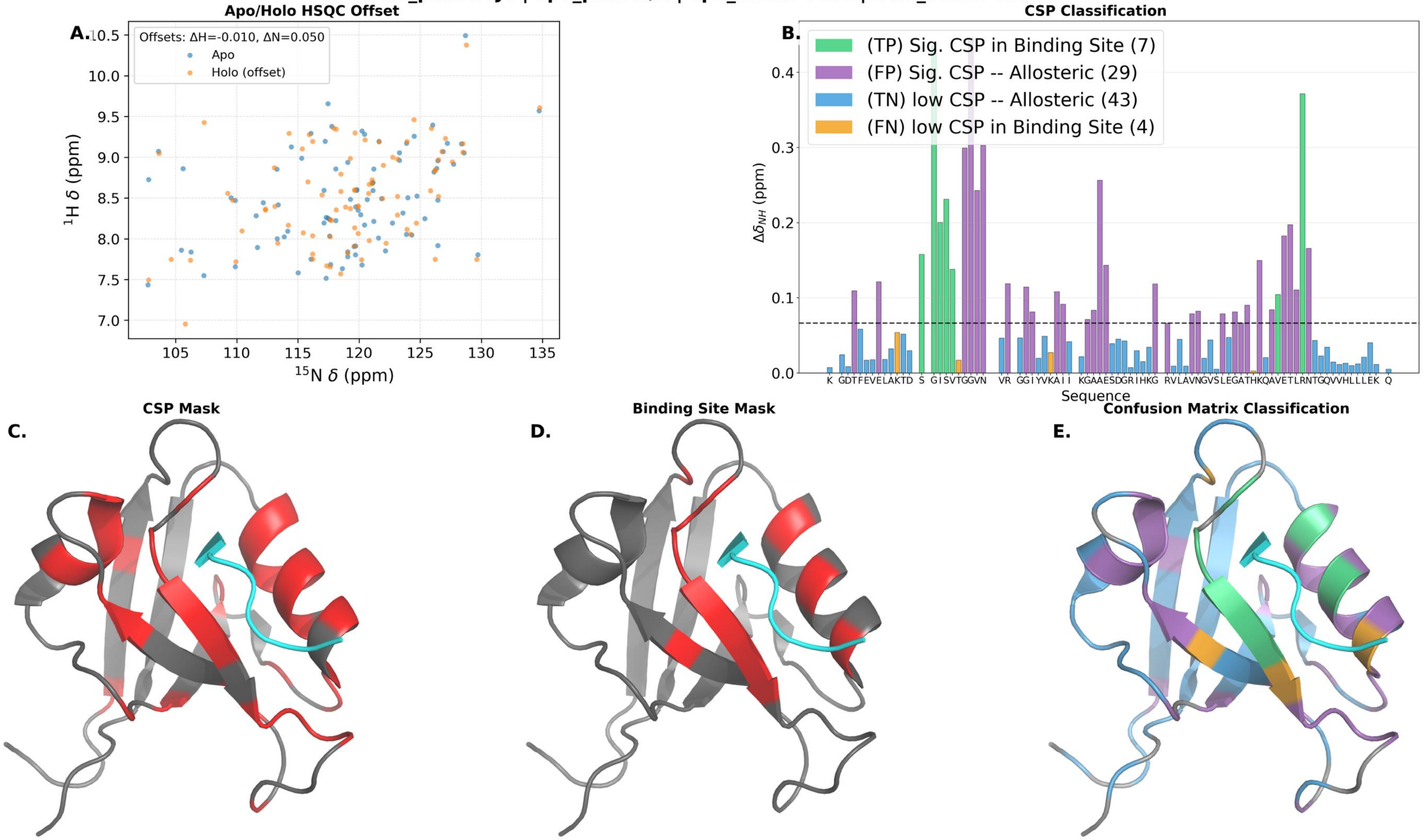

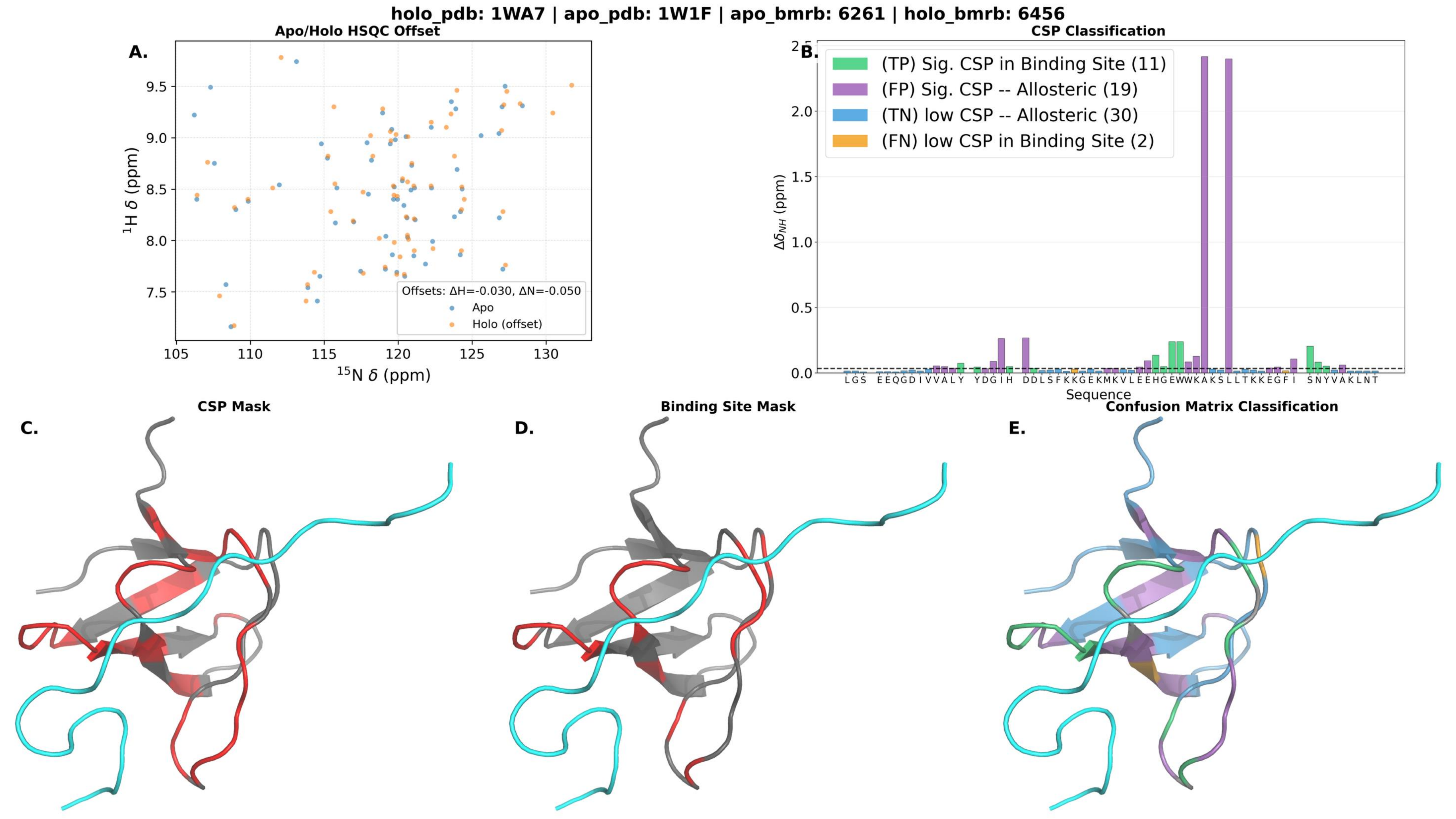
holo_pdb: 1WA7 | apo_pdb: 1W1F | apo_bmrb: 6261 | holo_bmrb: 6456
A.
Apo/Holo HSQC Offset
Offsets: ΔH=-0.030, ΔN=-0.050
Apo
Holo (offset)
1H δ (ppm)
15N δ (ppm)
B.
CSP Classification
(TP) Sig. CSP in Binding Site (11)
(FP) Sig. CSP -- Allosteric (19)
(TN) low CSP -- Allosteric (30)
(FN) low CSP in Binding Site (2)
ΔδNH (ppm)
Sequence
C.
CSP Mask
D.
Binding Site Mask
E.
Confusion Matrix Classification

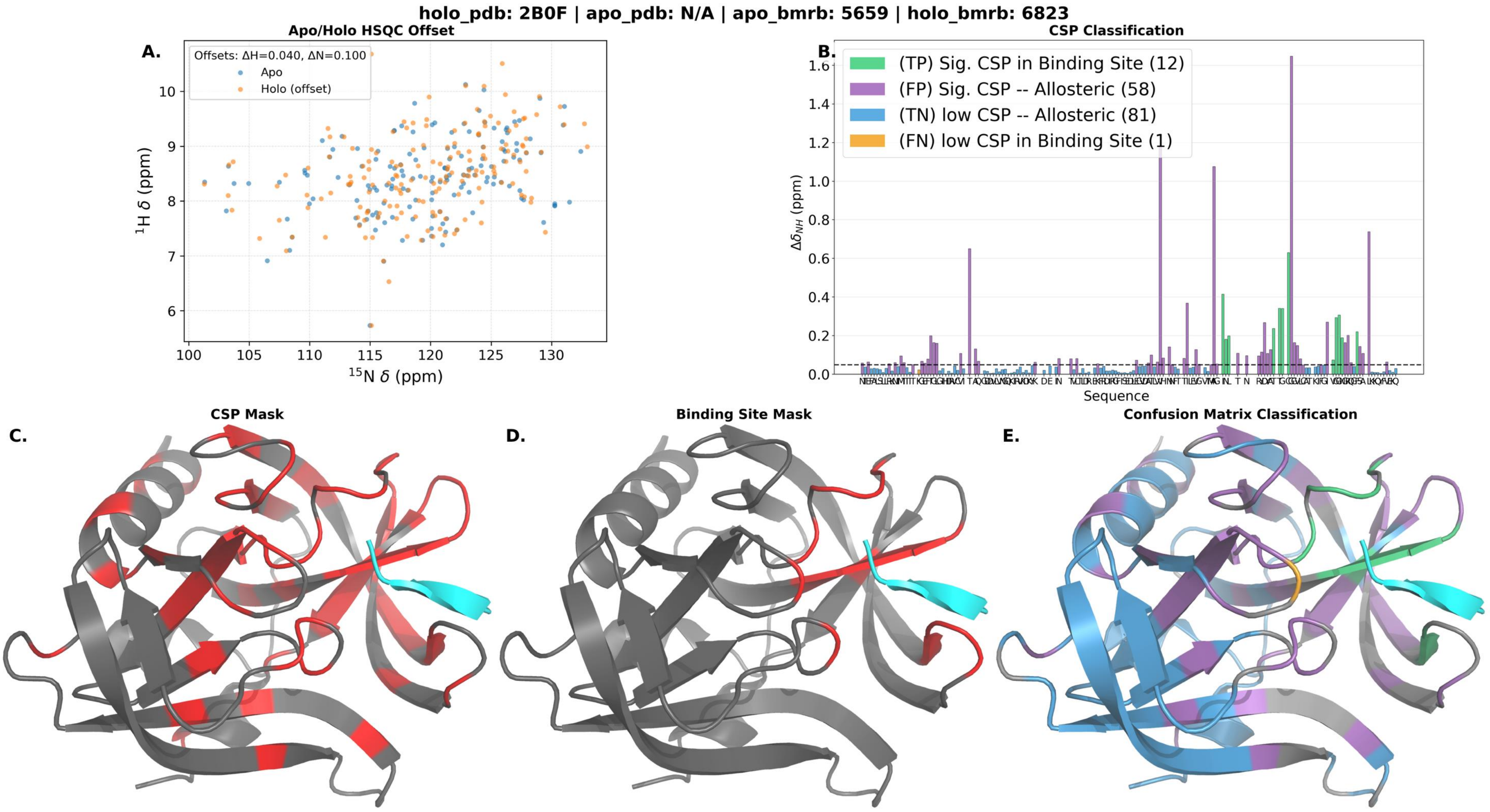
holo_pdb: 2B0F | apo_pdb: N/A | apo_bmrb: 5659 | holo_bmrb: 6823
A.
Apo/Holo HSQC Offset
Offsets: ΔH=0.040, ΔN=0.100
Apo
Holo (offset)
1H δ (ppm)
15N δ (ppm)
B.
CSP Classification
(TP) Sig. CSP in Binding Site (12)
(FP) Sig. CSP -- Allosteric (58)
(TN) low CSP -- Allosteric (81)
(FN) low CSP in Binding Site (1)
ΔδNH (ppm)
Sequence
C.
CSP Mask
D.
Binding Site Mask
E.
Confusion Matrix Classification

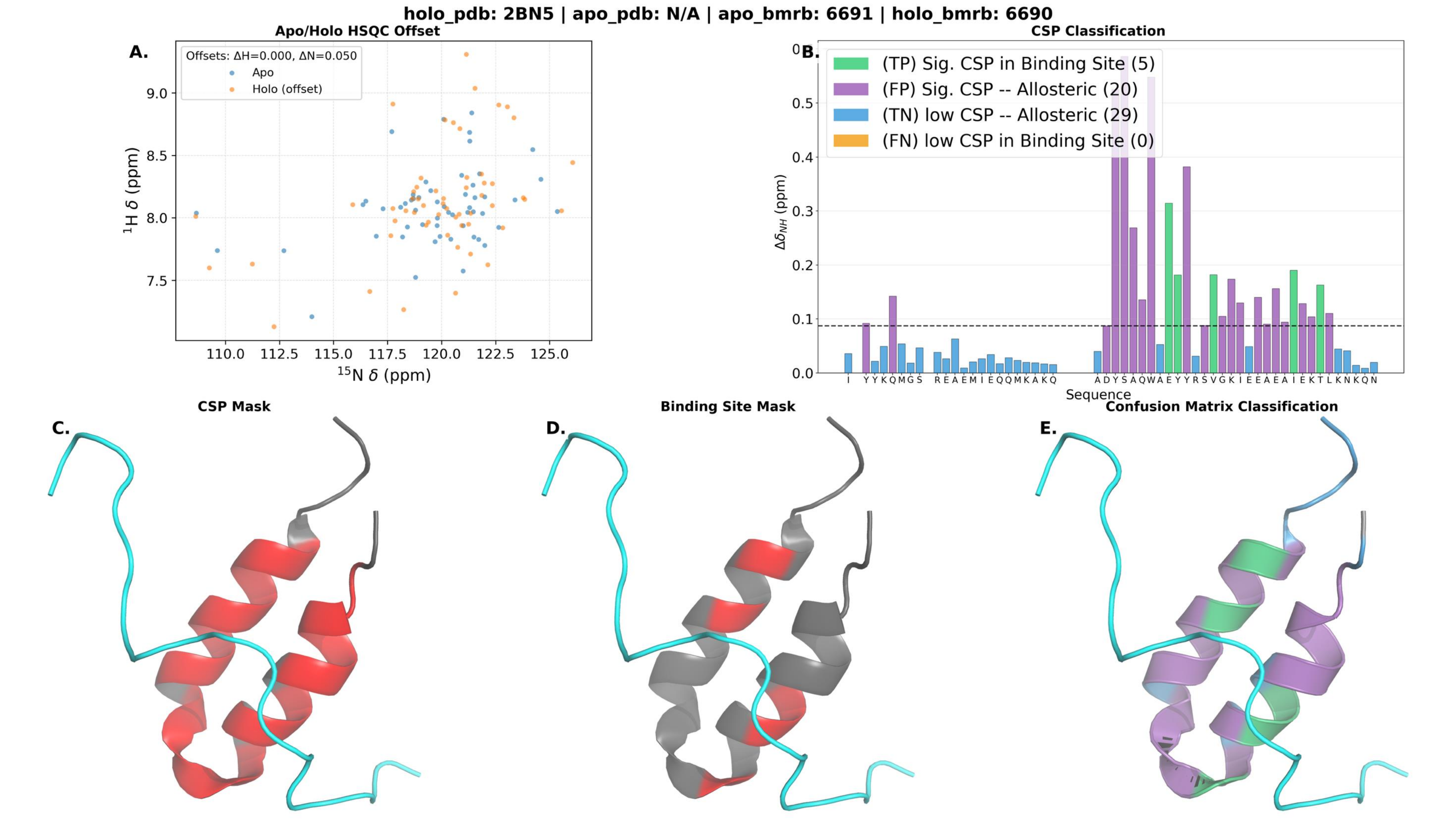
holo_pdb: 2BN5 | apo_pdb: N/A | apo_bmrb: 6691 | holo_bmrb: 6690
A.
Apo/Holo HSQC Offset
Offsets: ΔH=0.000, ΔN=0.050
Apo
Holo (offset)
$^{1}$H δ (ppm)
$^{15}$N δ (ppm)
B.
CSP Classification
(TP) Sig. CSP in Binding Site (5)
(FP) Sig. CSP -- Allosteric (20)
(TN) low CSP -- Allosteric (29)
(FN) low CSP in Binding Site (0)
$\Delta\delta_{NH}$ (ppm)
Sequence
C.
CSP Mask
D.
Binding Site Mask
E.
Confusion Matrix Classification

holo_pdb: 2CZY | apo_pdb: N/A | apo_bmrb: 27227 | holo_bmrb: 6749

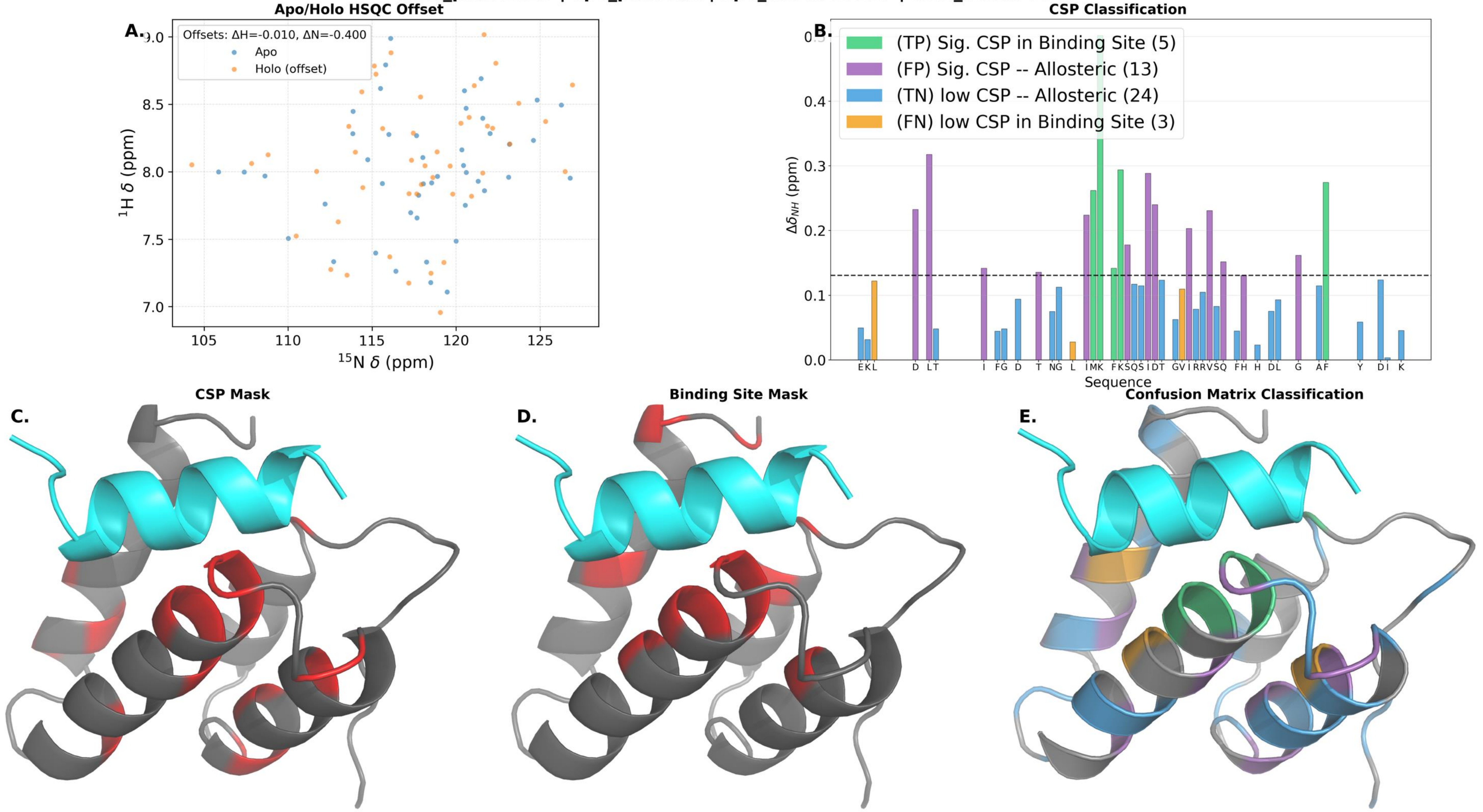

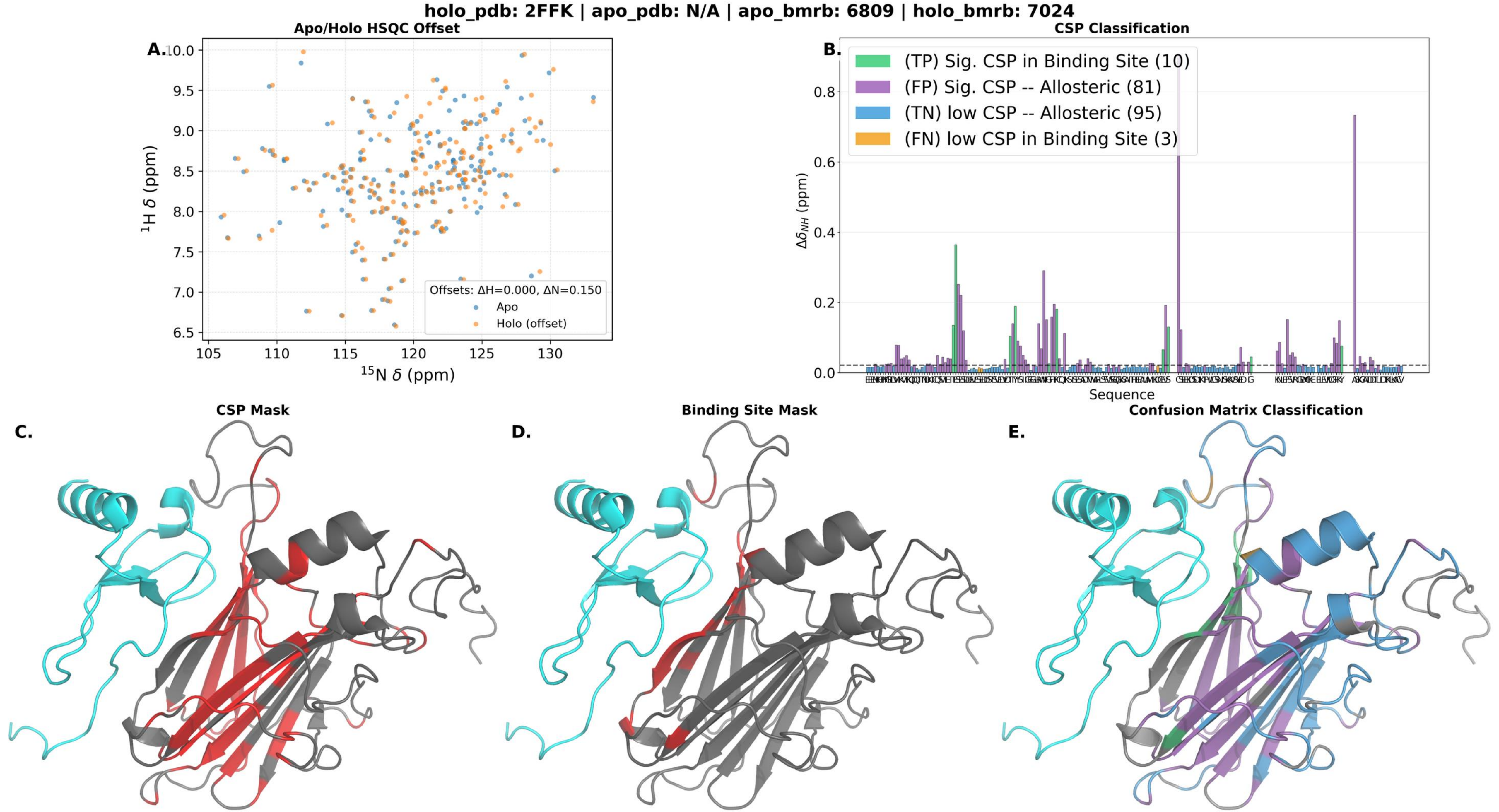

holo_pdb: 2FFK | apo_pdb: N/A | apo_bmrb: 6809 | holo_bmrb: 7024
A.
Apo/Holo HSQC Offset
Offsets: ΔH=0.000, ΔN=0.150
Apo
Holo (offset)
$^{1}$H δ (ppm)
$^{15}$N δ (ppm)
B.
CSP Classification
(TP) Sig. CSP in Binding Site (10)
(FP) Sig. CSP -- Allosteric (81)
(TN) low CSP -- Allosteric (95)
(FN) low CSP in Binding Site (3)
$\Delta\delta_{NH}$ (ppm)
Sequence
C.
CSP Mask
D.
Binding Site Mask
E.
Confusion Matrix Classification

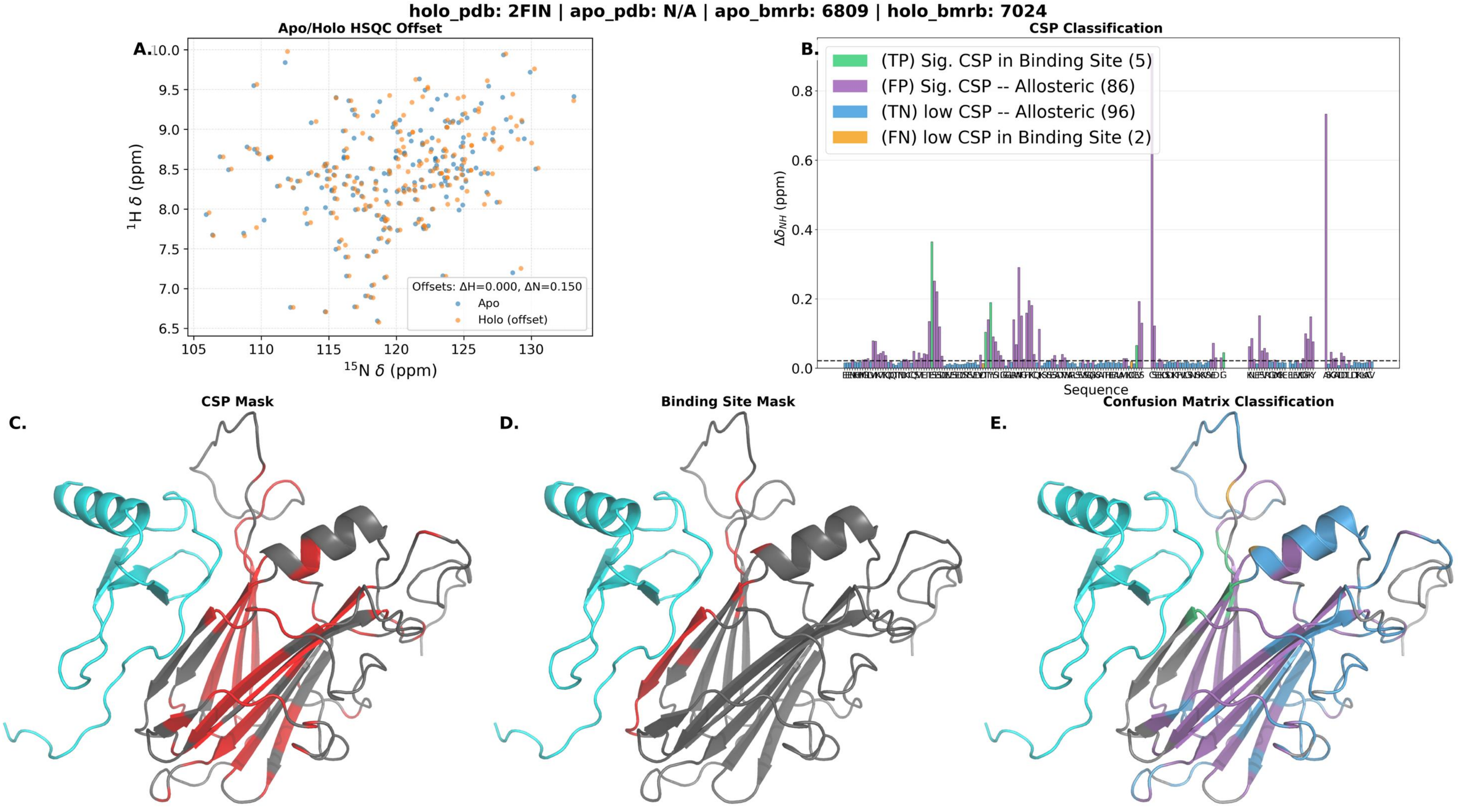
holo_pdb: 2FIN | apo_pdb: N/A | apo_bmrb: 6809 | holo_bmrb: 7024
Apo/Holo HSQC Offset
A.
Offsets: ΔH=0.000, ΔN=0.150
Apo
Holo (offset)
$^{1}$H δ (ppm)
$^{15}$N δ (ppm)
CSP Classification
B.
(TP) Sig. CSP in Binding Site (5)
(FP) Sig. CSP -- Allosteric (86)
(TN) low CSP -- Allosteric (96)
(FN) low CSP in Binding Site (2)
$\Delta\delta_{NH}$ (ppm)
Sequence
CSP Mask
C.
Binding Site Mask
D.
Confusion Matrix Classification
E.

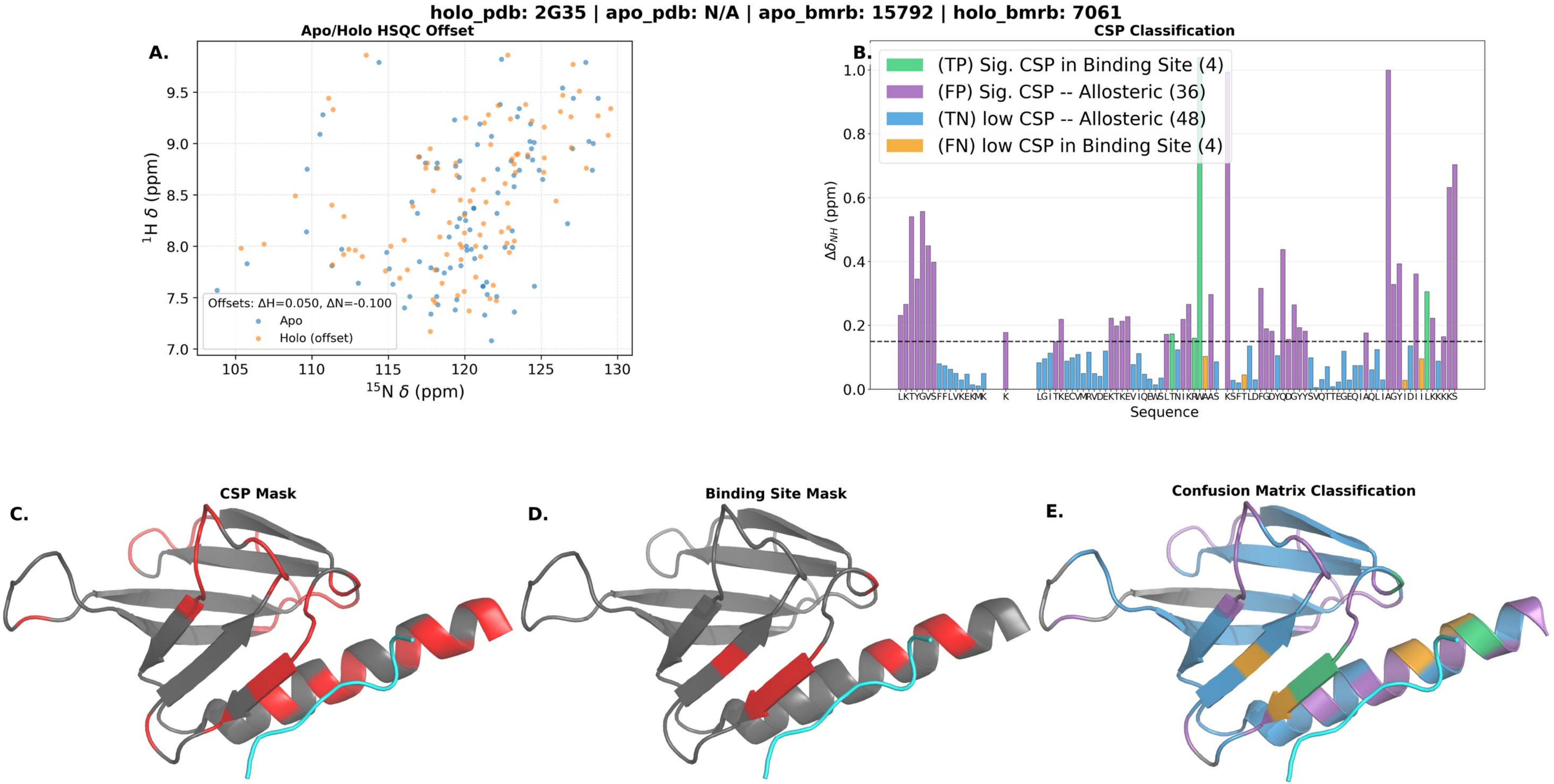
holo_pdb: 2G35 | apo_pdb: N/A | apo_bmrb: 15792 | holo_bmrb: 7061
A.
Apo/Holo HSQC Offset
Offsets: ΔH=0.050, ΔN=-0.100
Apo
Holo (offset)
1H δ (ppm)
15N δ (ppm)
B.
CSP Classification
(TP) Sig. CSP in Binding Site (4)
(FP) Sig. CSP -- Allosteric (36)
(TN) low CSP -- Allosteric (48)
(FN) low CSP in Binding Site (4)
ΔδNH (ppm)
Sequence
C.
CSP Mask
D.
Binding Site Mask
E.
Confusion Matrix Classification

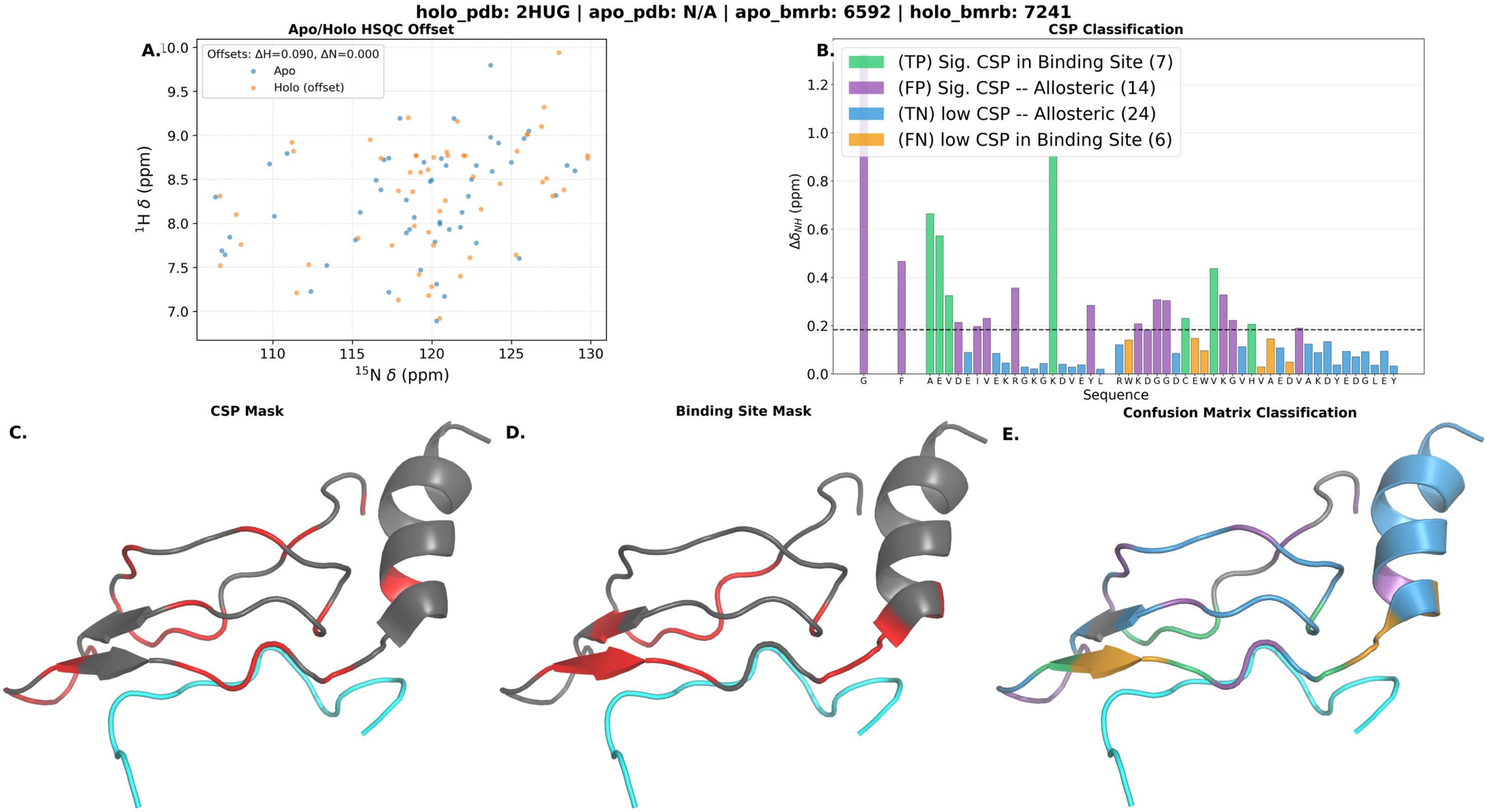

holo_pdb: 2HUG | apo_pdb: N/A | apo_bmrb: 6592 | holo_bmrb: 7241
A.
Apo/Holo HSQC Offset
Offsets: ΔH=0.090, ΔN=0.000
Apo
Holo (offset)
$^{1}$H δ (ppm)
$^{15}$N δ (ppm)
B.
CSP Classification
(TP) Sig. CSP in Binding Site (7)
(FP) Sig. CSP -- Allosteric (14)
(TN) low CSP -- Allosteric (24)
(FN) low CSP in Binding Site (6)
$\Delta\delta_{NH}$ (ppm)
Sequence
G F A E V D E I V E K R G K G K D V E Y L R W K D G G D C E W V K G V H V A E D V A K D Y E D G L E Y
C.
CSP Mask
D.
Binding Site Mask
E.
Confusion Matrix Classification

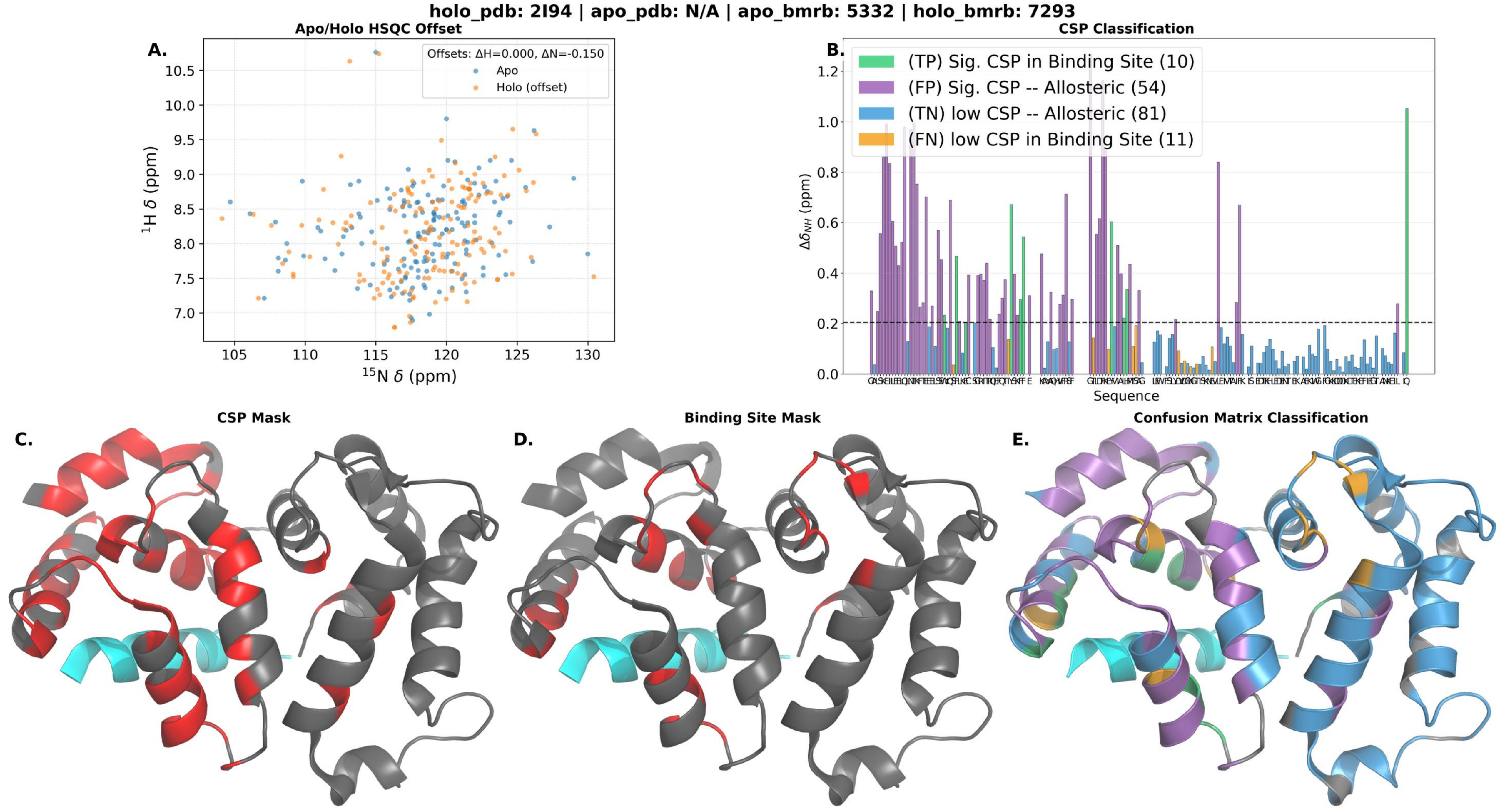

holo_pdb: 2I94 | apo_pdb: N/A | apo_bmrb: 5332 | holo_bmrb: 7293
A.
Apo/Holo HSQC Offset
Offsets: ΔH=0.000, ΔN=-0.150
Apo
Holo (offset)
$^{1}$H δ (ppm)
$^{15}$N δ (ppm)
B.
CSP Classification
(TP) Sig. CSP in Binding Site (10)
(FP) Sig. CSP -- Allosteric (54)
(TN) low CSP -- Allosteric (81)
(FN) low CSP in Binding Site (11)
$\Delta\delta_{NH}$ (ppm)
Sequence
C.
CSP Mask
D.
Binding Site Mask
E.
Confusion Matrix Classification

holo_pdb: 2IPA | apo_pdb: N/A | apo_bmrb: 6075 | holo_bmrb: 15028

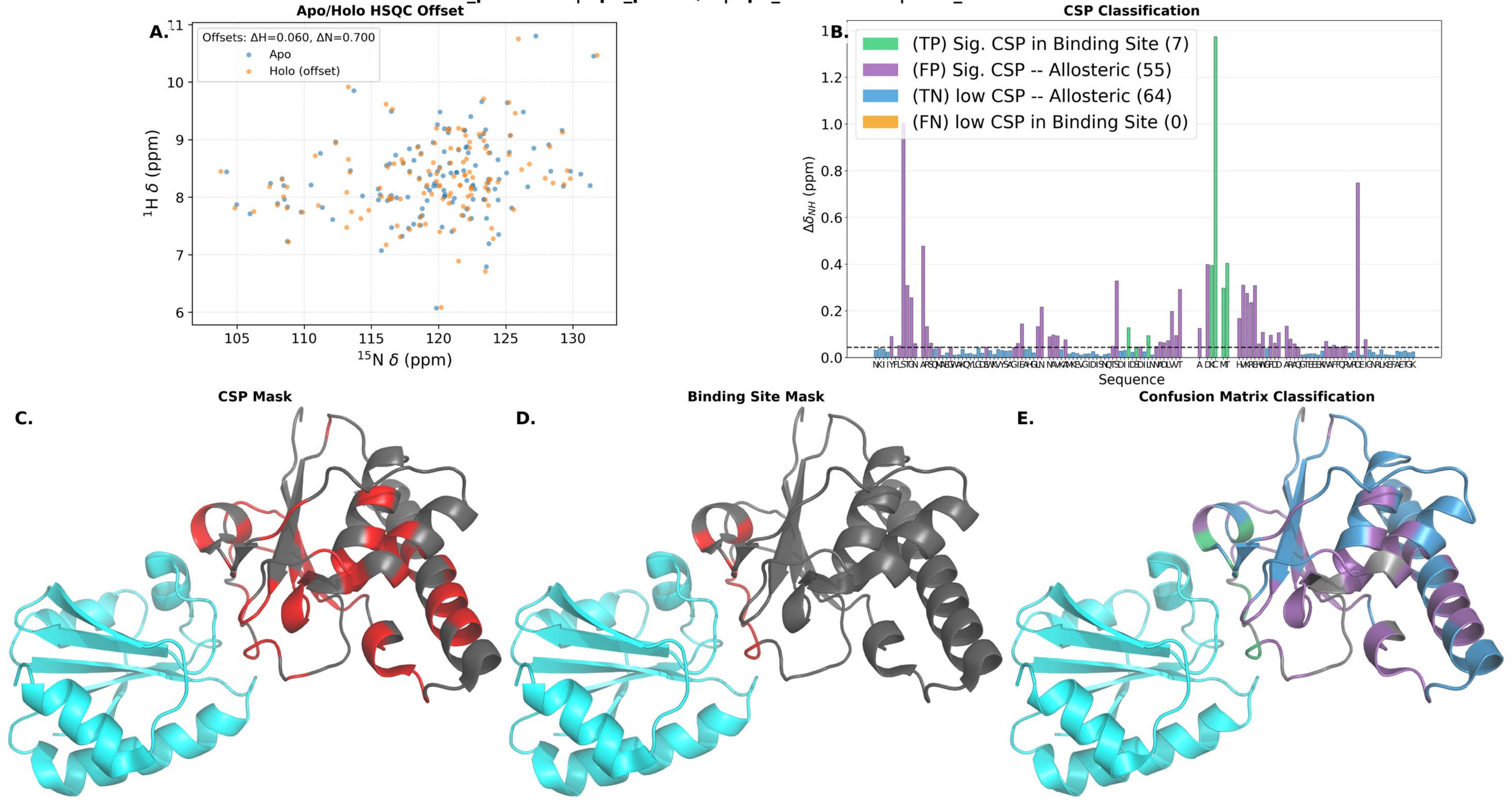

holo_pdb: 2JMX | apo_pdb: N/A | apo_bmrb: 6564 | holo_bmrb: 15072

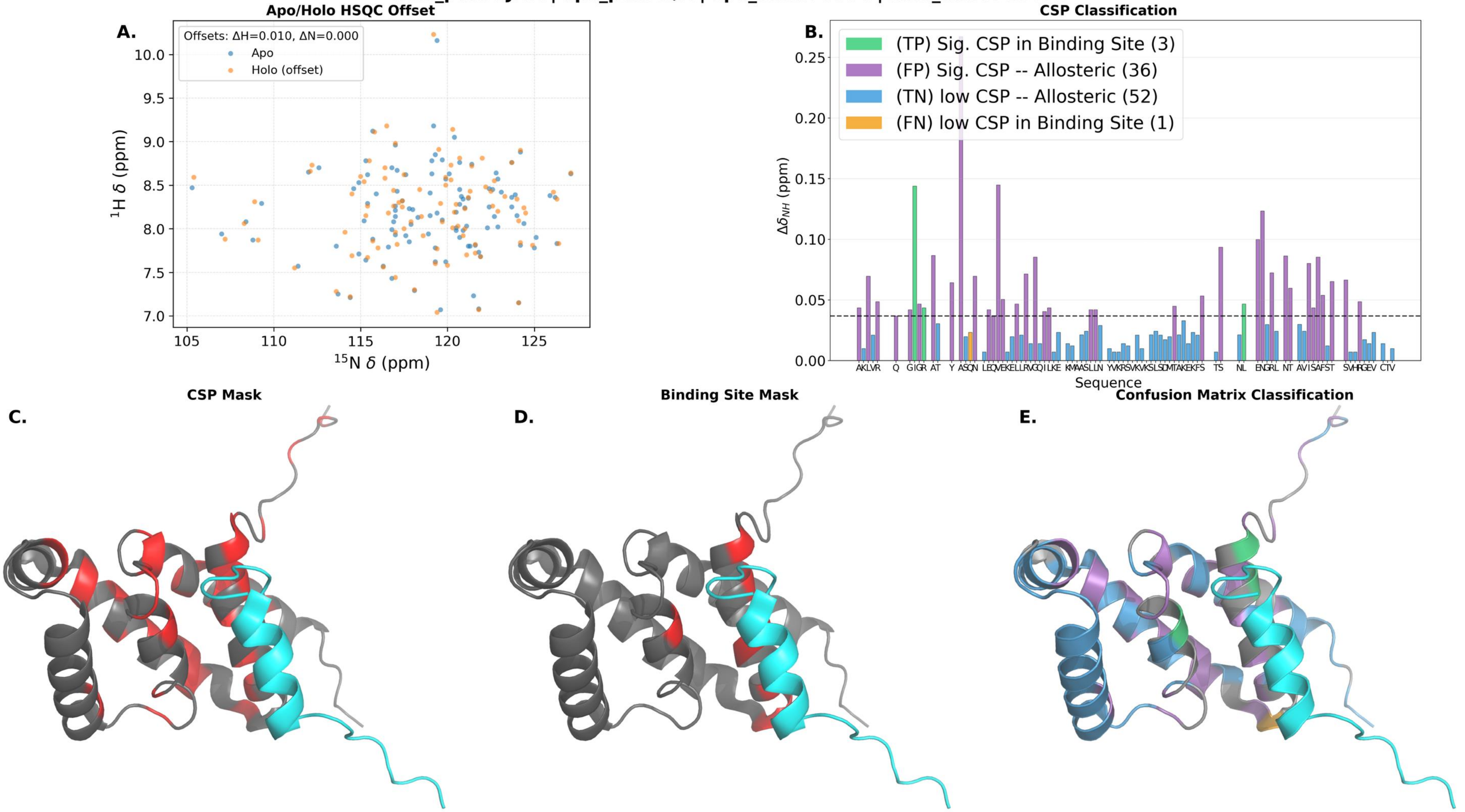

**holo_pdb: 2JW1 | apo_pdb: N/A | apo_bmrb: 15503 | holo_bmrb: 15504**

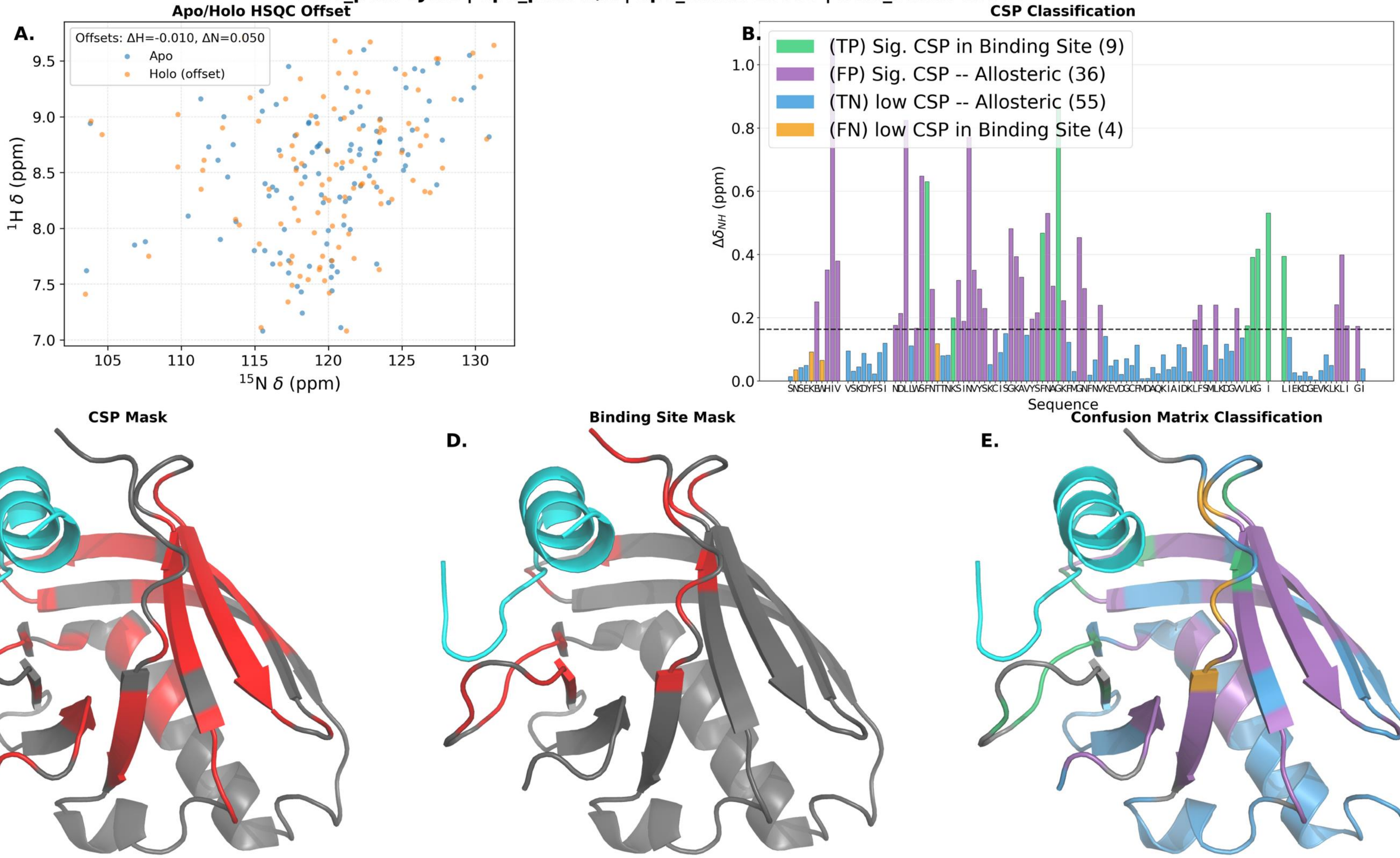

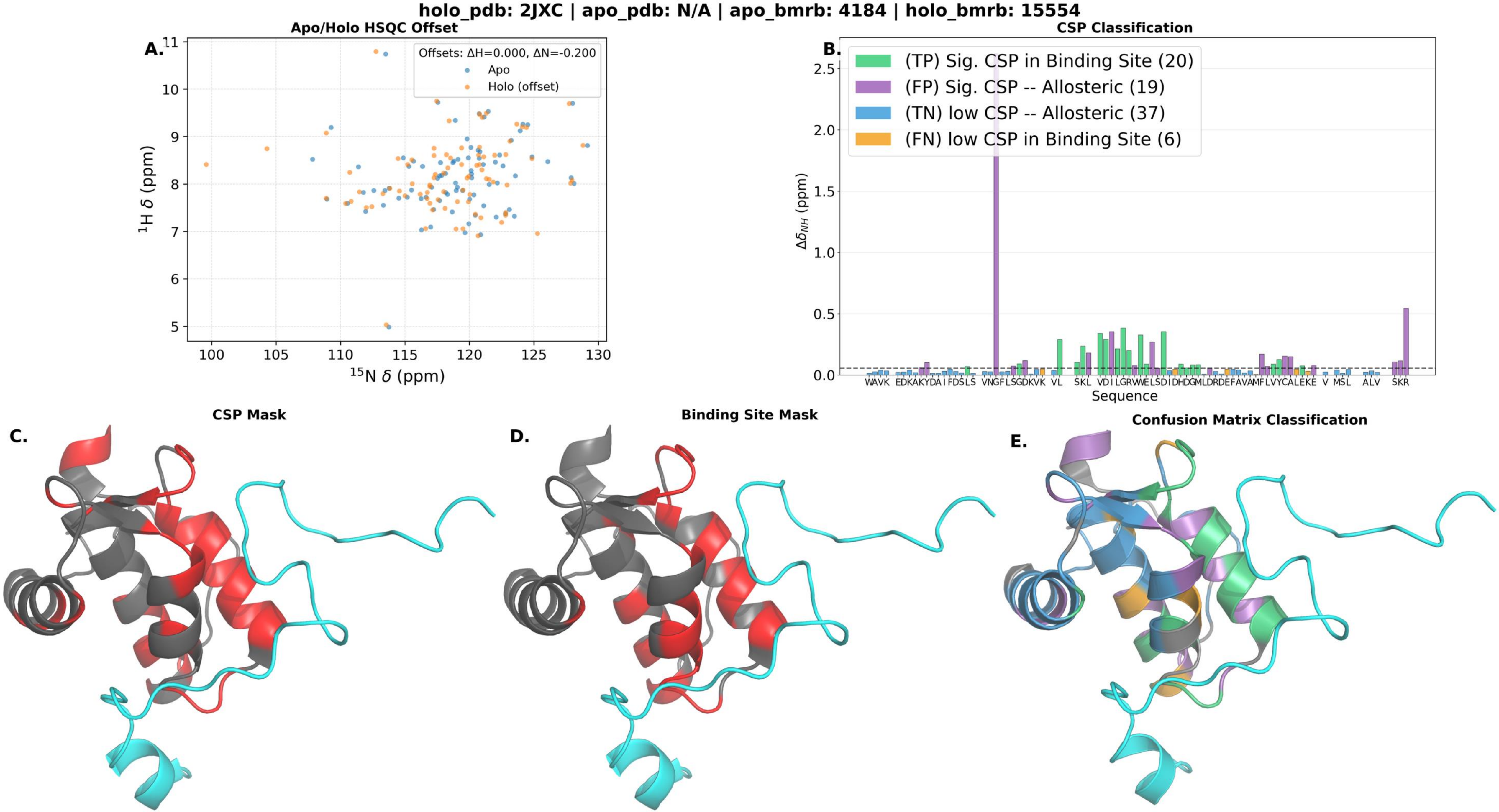

holo_pdb: 2JXC | apo_pdb: N/A | apo_bmrb: 4184 | holo_bmrb: 15554
A.
Apo/Holo HSQC Offset
Offsets: ΔH=0.000, ΔN=-0.200
Apo
Holo (offset)
$^{1}$H δ (ppm)
$^{15}$N δ (ppm)
11
10
9
8
7
6
5
100
105
110
115
120
125
130
B.
CSP Classification
(TP) Sig. CSP in Binding Site (20)
(FP) Sig. CSP -- Allosteric (19)
(TN) low CSP -- Allosteric (37)
(FN) low CSP in Binding Site (6)
$\Delta\delta_{NH}$ (ppm)
2.5
2.0
1.5
1.0
0.5
0.0
WAVK EDKAKYDAIFDSLS VNGFLSGDKVK VL SKL VDILGRWELSDIDHDGMLDRDEFAVAMFLVYCALEKE V MSL ALV SKR
Sequence
C.
CSP Mask
D.
Binding Site Mask
E.
Confusion Matrix Classification

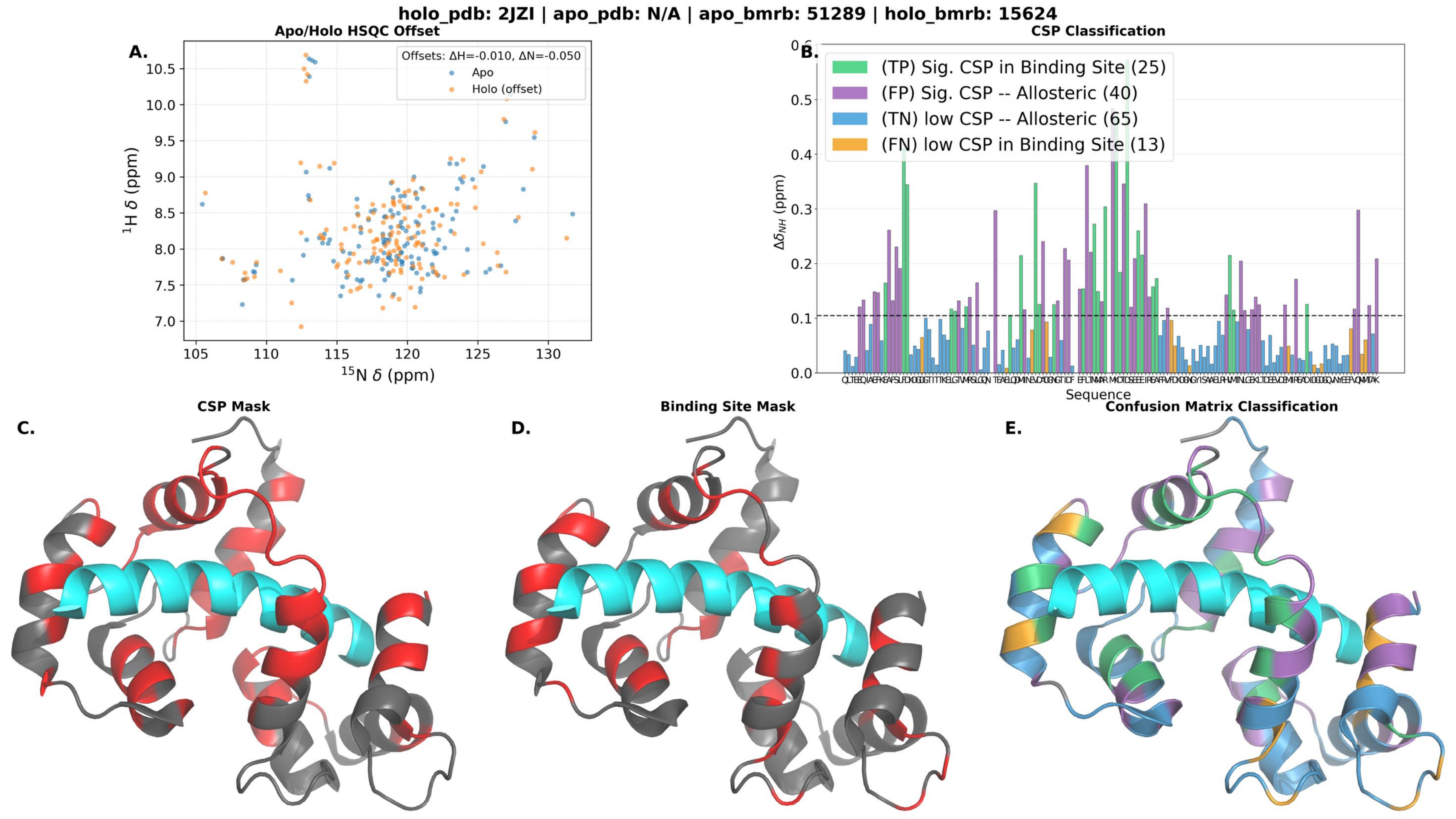

holo_pdb: 2JZI | apo_pdb: N/A | apo_bmrb: 51289 | holo_bmrb: 15624
A.
Apo/Holo HSQC Offset
Offsets: ΔH=-0.010, ΔN=-0.050
Apo
Holo (offset)
$^{1}$H δ (ppm)
$^{15}$N δ (ppm)
B.
CSP Classification
(TP) Sig. CSP in Binding Site (25)
(FP) Sig. CSP -- Allosteric (40)
(TN) low CSP -- Allosteric (65)
(FN) low CSP in Binding Site (13)
$\Delta\delta_{NH}$ (ppm)
Sequence
C.
CSP Mask
D.
Binding Site Mask
E.
Confusion Matrix Classification

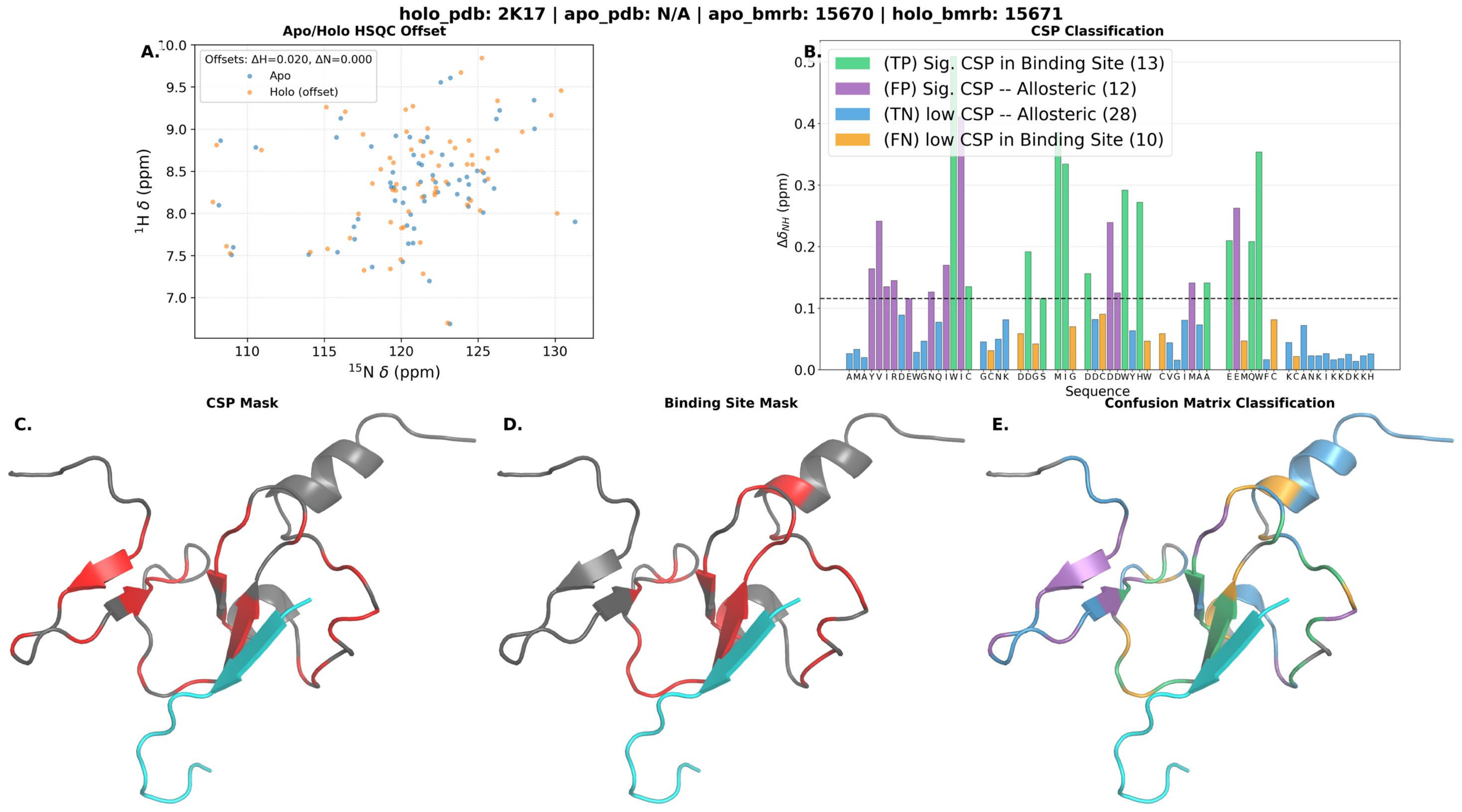

holo_pdb: 2K17 | apo_pdb: N/A | apo_bmrb: 15670 | holo_bmrb: 15671
A.
Apo/Holo HSQC Offset
Offsets: ΔH=0.020, ΔN=0.000
Apo
Holo (offset)
$^{1}$H δ (ppm)
$^{15}$N δ (ppm)
B.
CSP Classification
(TP) Sig. CSP in Binding Site (13)
(FP) Sig. CSP -- Allosteric (12)
(TN) low CSP -- Allosteric (28)
(FN) low CSP in Binding Site (10)
$\Delta\delta_{NH}$ (ppm)
Sequence
C.
CSP Mask
D.
Binding Site Mask
E.
Confusion Matrix Classification

**holo_pdb: 2K6D | apo_pdb: N/A | apo_bmrb: 52080 | holo_bmrb: 15866**

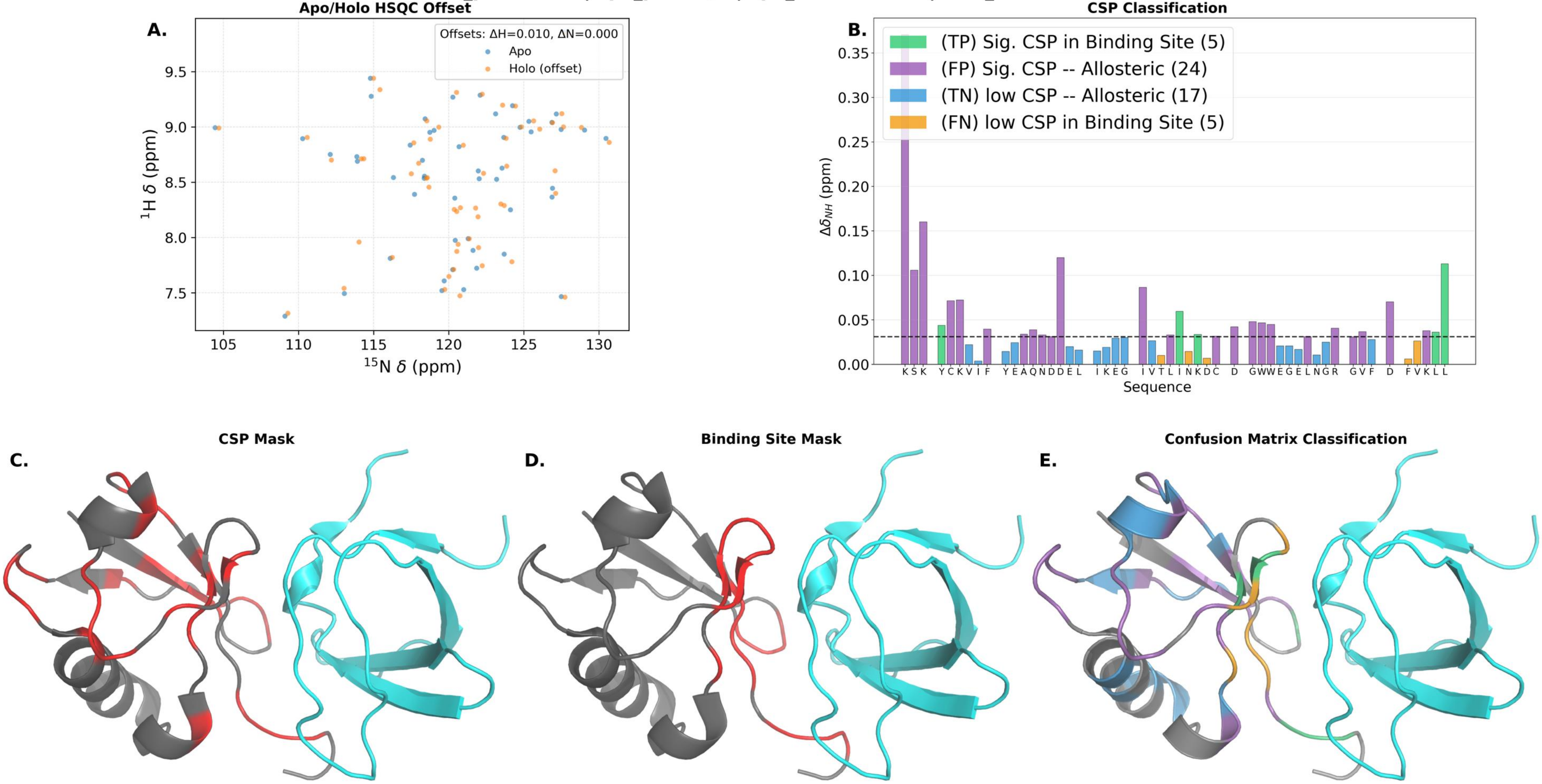

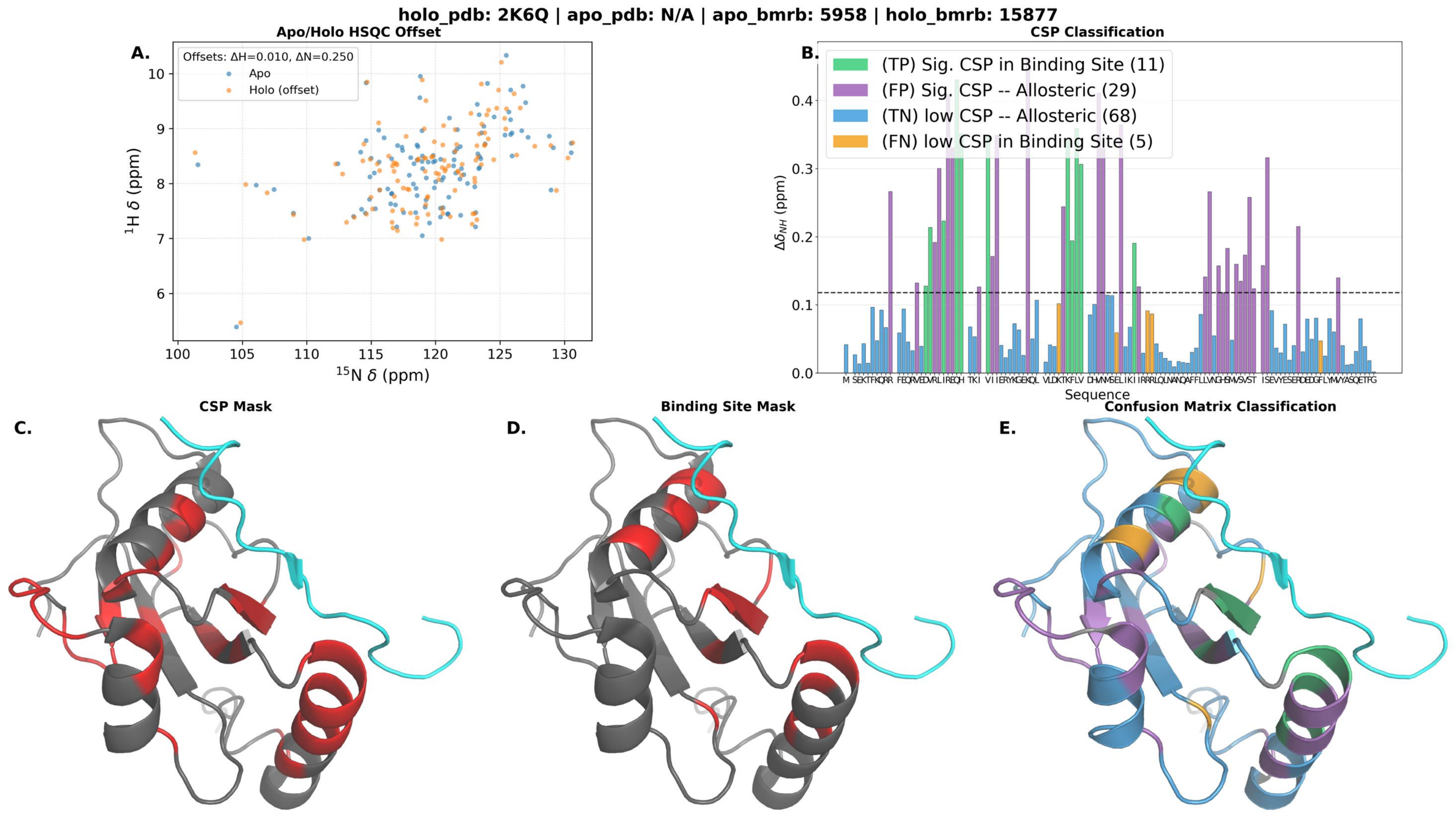

holo_pdb: 2K6Q | apo_pdb: N/A | apo_bmrb: 5958 | holo_bmrb: 15877
A.
Apo/Holo HSQC Offset
Offsets: ΔH=0.010, ΔN=0.250
Apo
Holo (offset)
1H δ (ppm)
15N δ (ppm)
B.
CSP Classification
(TP) Sig. CSP in Binding Site (11)
(FP) Sig. CSP -- Allosteric (29)
(TN) low CSP -- Allosteric (68)
(FN) low CSP in Binding Site (5)
ΔδNH (ppm)
Sequence
C.
CSP Mask
D.
Binding Site Mask
E.
Confusion Matrix Classification

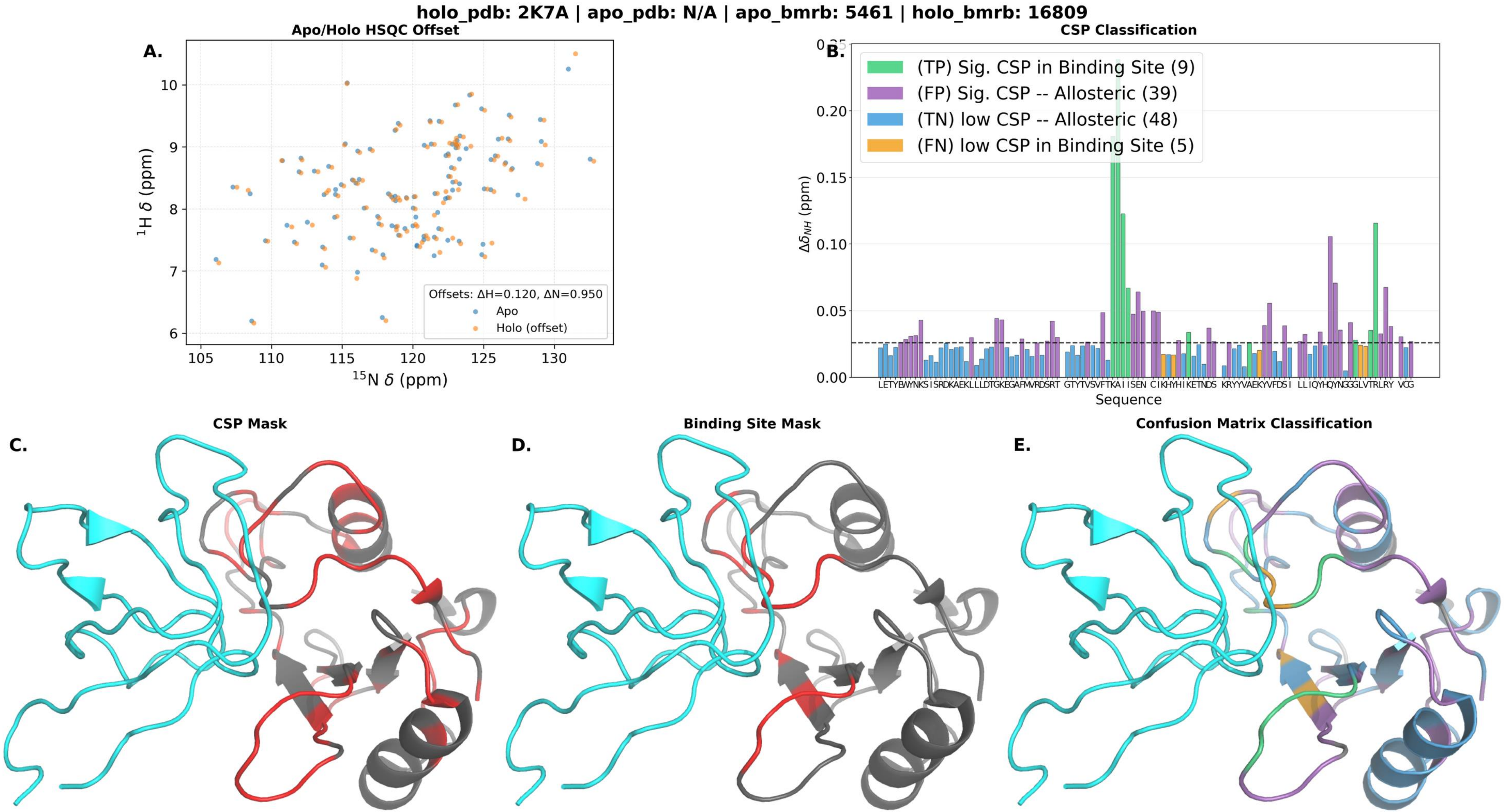
holo_pdb: 2K7A | apo_pdb: N/A | apo_bmrb: 5461 | holo_bmrb: 16809
A.
Apo/Holo HSQC Offset
$^{1}$H $\delta$ (ppm)
$^{15}$N $\delta$ (ppm)
Offsets: ΔH=0.120, ΔN=0.950
Apo
Holo (offset)
B.
CSP Classification
(TP) Sig. CSP in Binding Site (9)
(FP) Sig. CSP -- Allosteric (39)
(TN) low CSP -- Allosteric (48)
(FN) low CSP in Binding Site (5)
$\Delta\delta_{NH}$ (ppm)
Sequence
C.
CSP Mask
D.
Binding Site Mask
E.
Confusion Matrix Classification

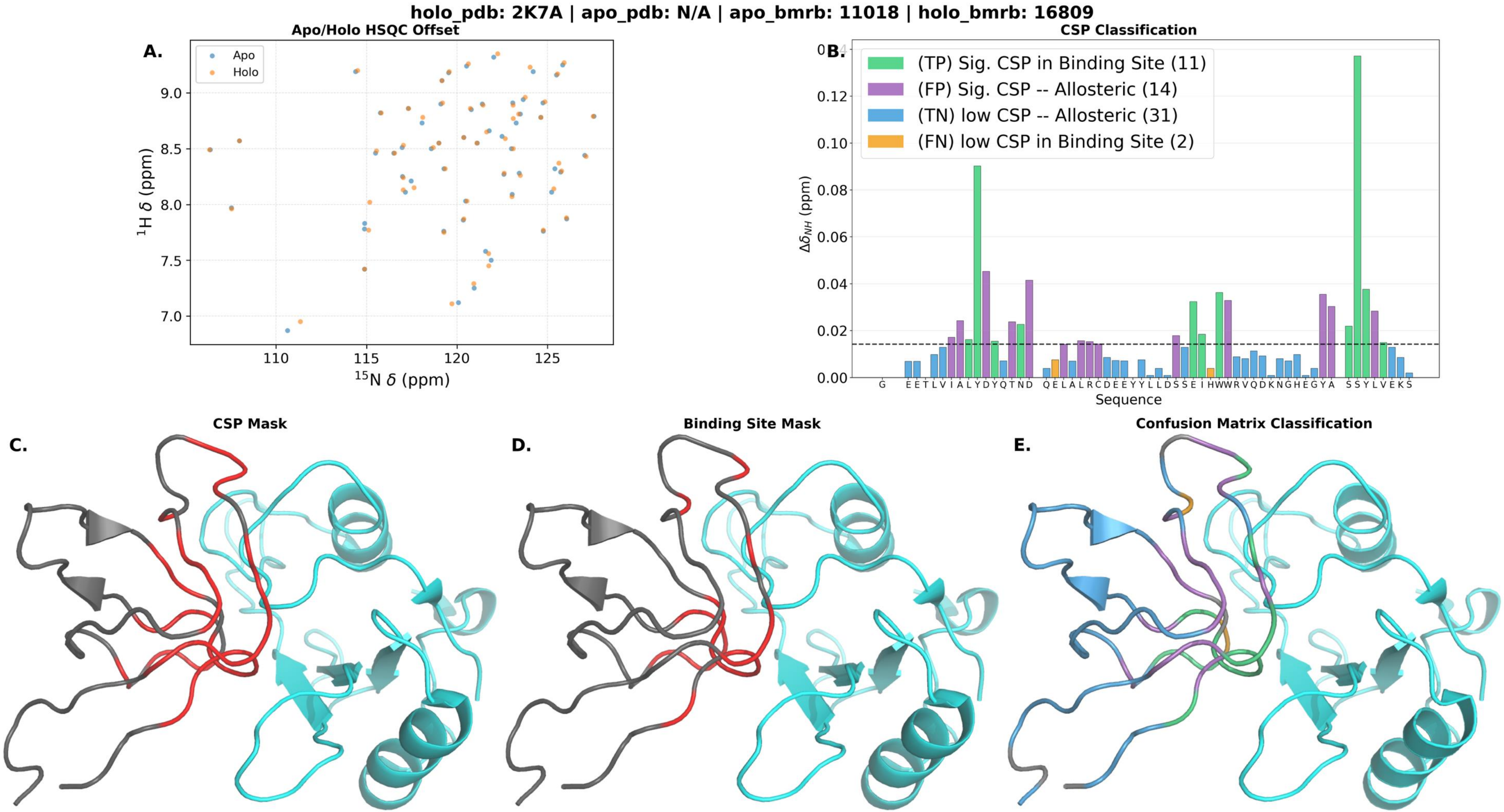
holo_pdb: 2K7A | apo_pdb: N/A | apo_bmrb: 11018 | holo_bmrb: 16809
A.
Apo/Holo HSQC Offset
Apo
Holo
$^{1}$H δ (ppm)
$^{15}$N δ (ppm)
B.
CSP Classification
(TP) Sig. CSP in Binding Site (11)
(FP) Sig. CSP -- Allosteric (14)
(TN) low CSP -- Allosteric (31)
(FN) low CSP in Binding Site (2)
$\Delta\delta_{NH}$ (ppm)
Sequence
C.
CSP Mask
D.
Binding Site Mask
E.
Confusion Matrix Classification

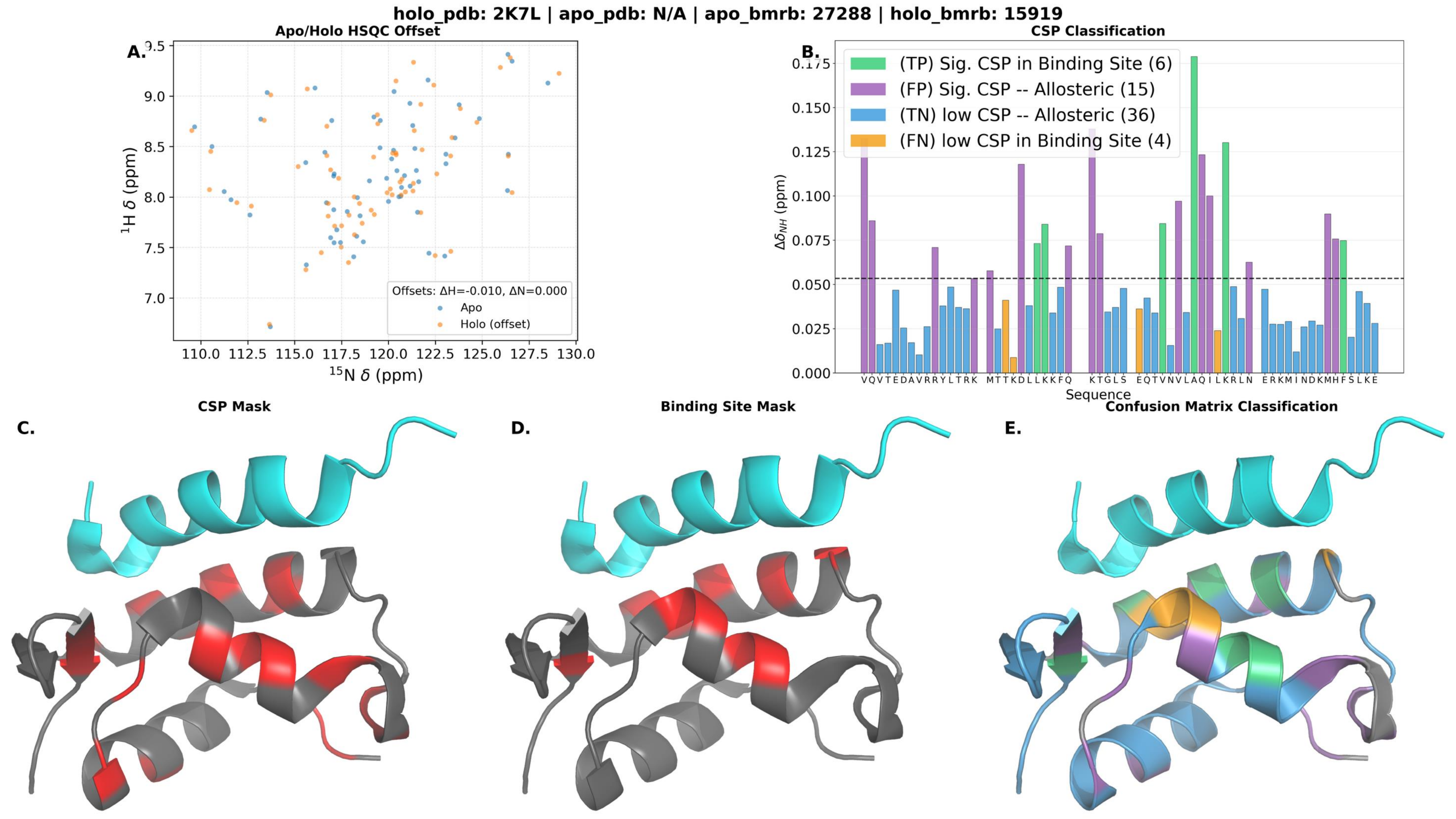

holo_pdb: 2K7L | apo_pdb: N/A | apo_bmrb: 27288 | holo_bmrb: 15919
Apo/Holo HSQC Offset
A.
Offsets: ΔH=-0.010, ΔN=0.000
Apo
Holo (offset)
$^{1}$H δ (ppm)
$^{15}$N δ (ppm)
CSP Classification
B.
(TP) Sig. CSP in Binding Site (6)
(FP) Sig. CSP -- Allosteric (15)
(TN) low CSP -- Allosteric (36)
(FN) low CSP in Binding Site (4)
$\Delta\delta_{NH}$ (ppm)
Sequence
VQVTEDAVRRYLTRK MTTKDLLKKFQ KTGLS EQTVNVLAQILKRLN ERKMINDKMHFSLKE
CSP Mask
C.
Binding Site Mask
D.
Confusion Matrix Classification
E.

**holo_pdb: 2K8F | apo_pdb: N/A | apo_bmrb: 34231 | holo_bmrb: 15944**

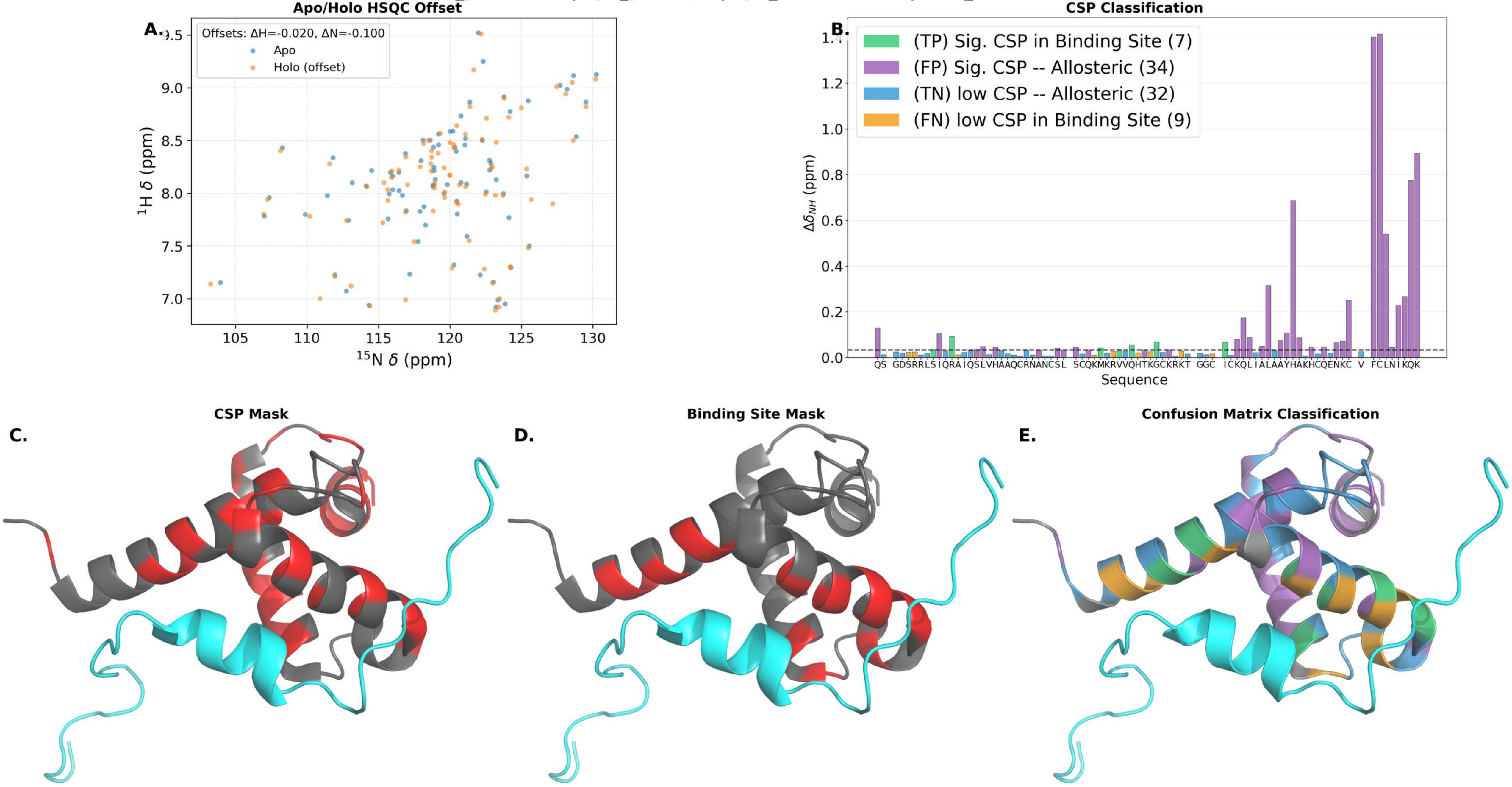

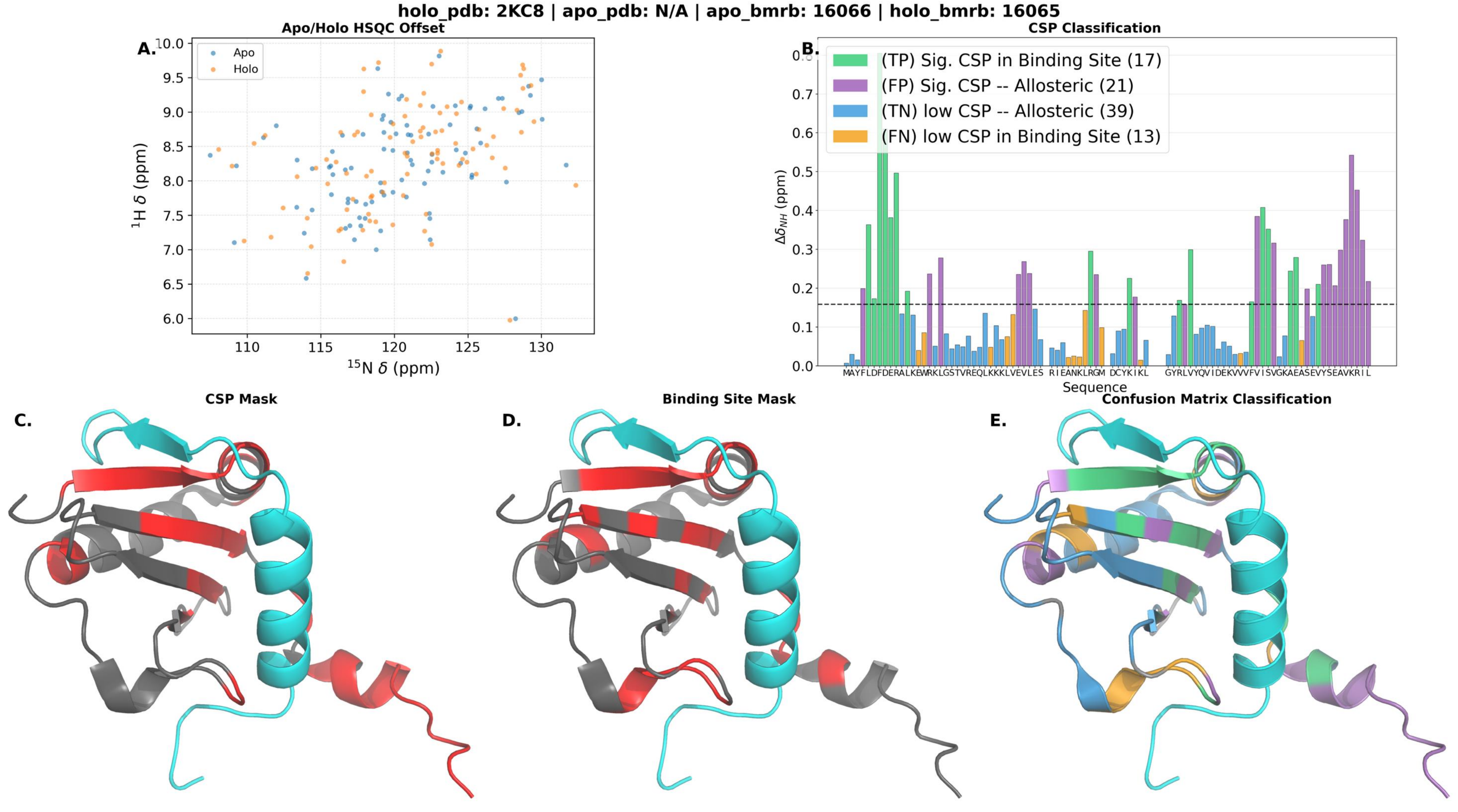

holo_pdb: 2KC8 | apo_pdb: N/A | apo_bmrb: 16066 | holo_bmrb: 16065
A.
Apo/Holo HSQC Offset
Apo
Holo
$^1$H δ (ppm)
$^{15}$N δ (ppm)
B.
CSP Classification
(TP) Sig. CSP in Binding Site (17)
(FP) Sig. CSP -- Allosteric (21)
(TN) low CSP -- Allosteric (39)
(FN) low CSP in Binding Site (13)
$\Delta\delta_{NH}$ (ppm)
Sequence
C.
CSP Mask
D.
Binding Site Mask
E.
Confusion Matrix Classification

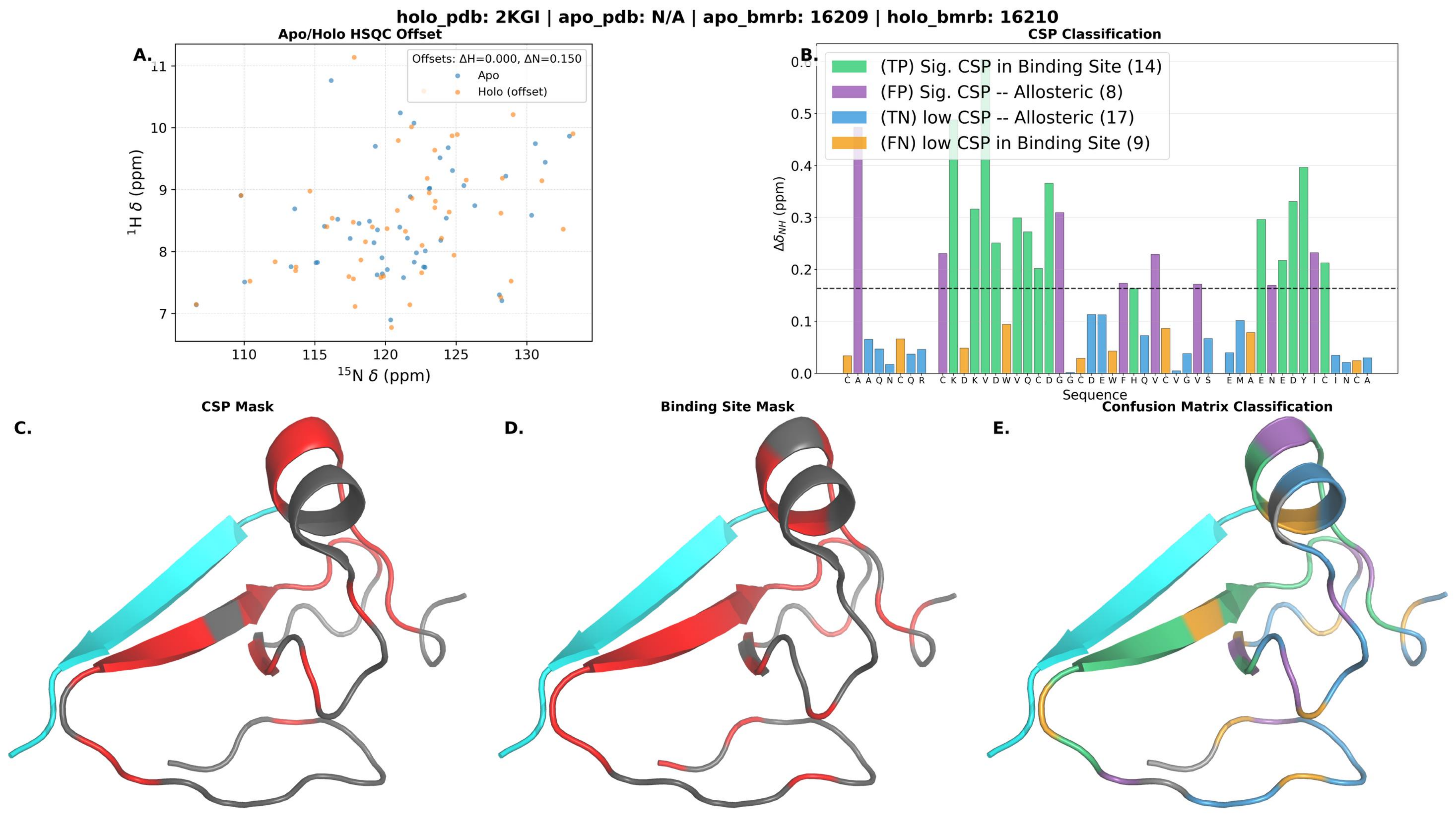
holo_pdb: 2KGI | apo_pdb: N/A | apo_bmrb: 16209 | holo_bmrb: 16210
A.
Apo/Holo HSQC Offset
Offsets: ΔH=0.000, ΔN=0.150
Apo
Holo (offset)
$^{1}$H δ (ppm)
$^{15}$N δ (ppm)
B.
CSP Classification
(TP) Sig. CSP in Binding Site (14)
(FP) Sig. CSP -- Allosteric (8)
(TN) low CSP -- Allosteric (17)
(FN) low CSP in Binding Site (9)
$\Delta\delta_{NH}$ (ppm)
Sequence
C A A Q N C Q R C K D K V D W V Q C D G G G C D E W F H Q V C V G V S E M A E N E D Y I C I N C A
C.
CSP Mask
D.
Binding Site Mask
E.
Confusion Matrix Classification

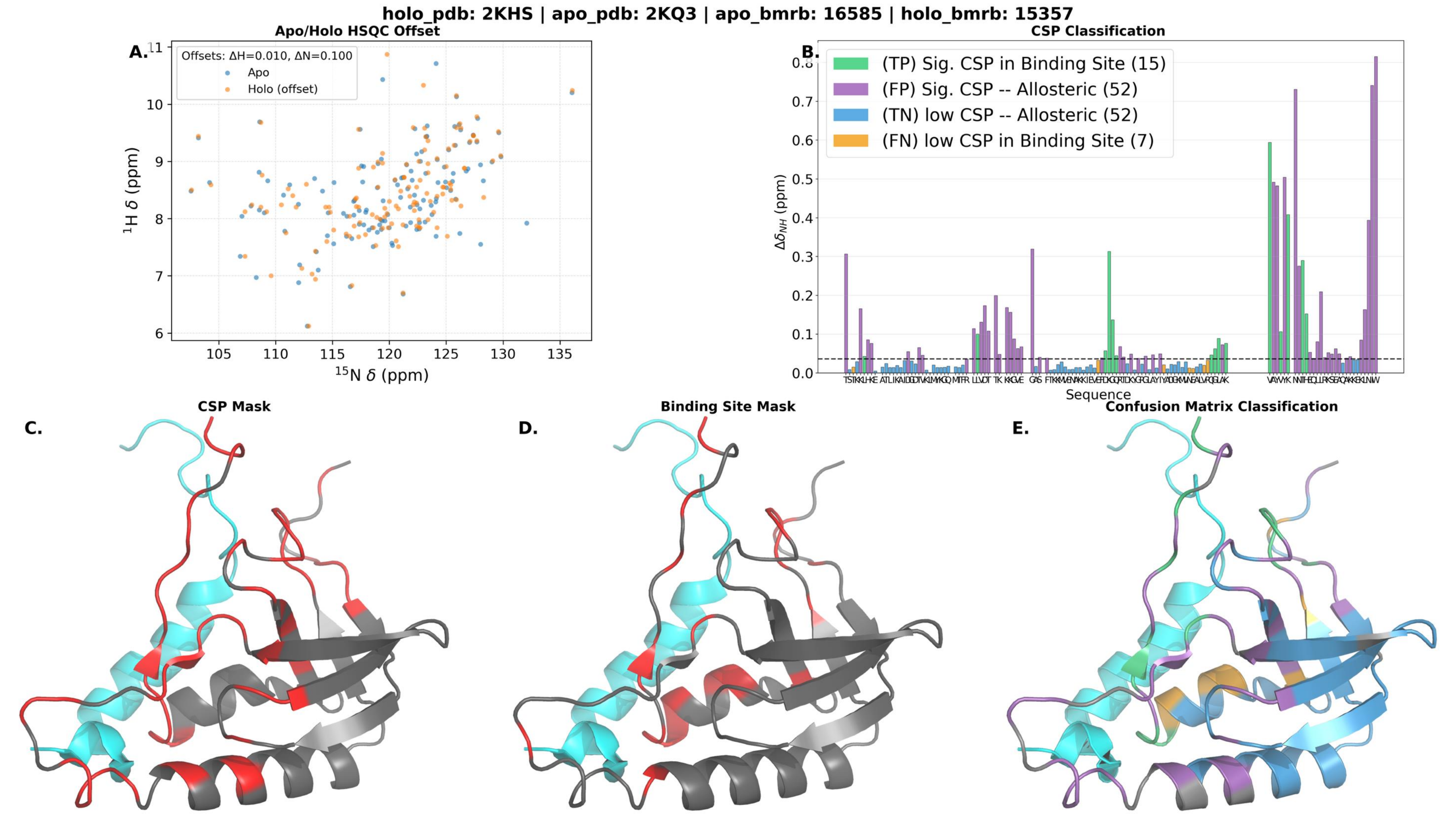
holo_pdb: 2KHS | apo_pdb: 2KQ3 | apo_bmrb: 16585 | holo_bmrb: 15357
A.
Apo/Holo HSQC Offset
Offsets: ΔH=0.010, ΔN=0.100
Apo
Holo (offset)
1H δ (ppm)
15N δ (ppm)
B.
CSP Classification
(TP) Sig. CSP in Binding Site (15)
(FP) Sig. CSP -- Allosteric (52)
(TN) low CSP -- Allosteric (52)
(FN) low CSP in Binding Site (7)
ΔδNH (ppm)
Sequence
C.
CSP Mask
D.
Binding Site Mask
E.
Confusion Matrix Classification

holo_pdb: 2KOH | apo_pdb: N/A | apo_bmrb: 27205 | holo_bmrb: 16507

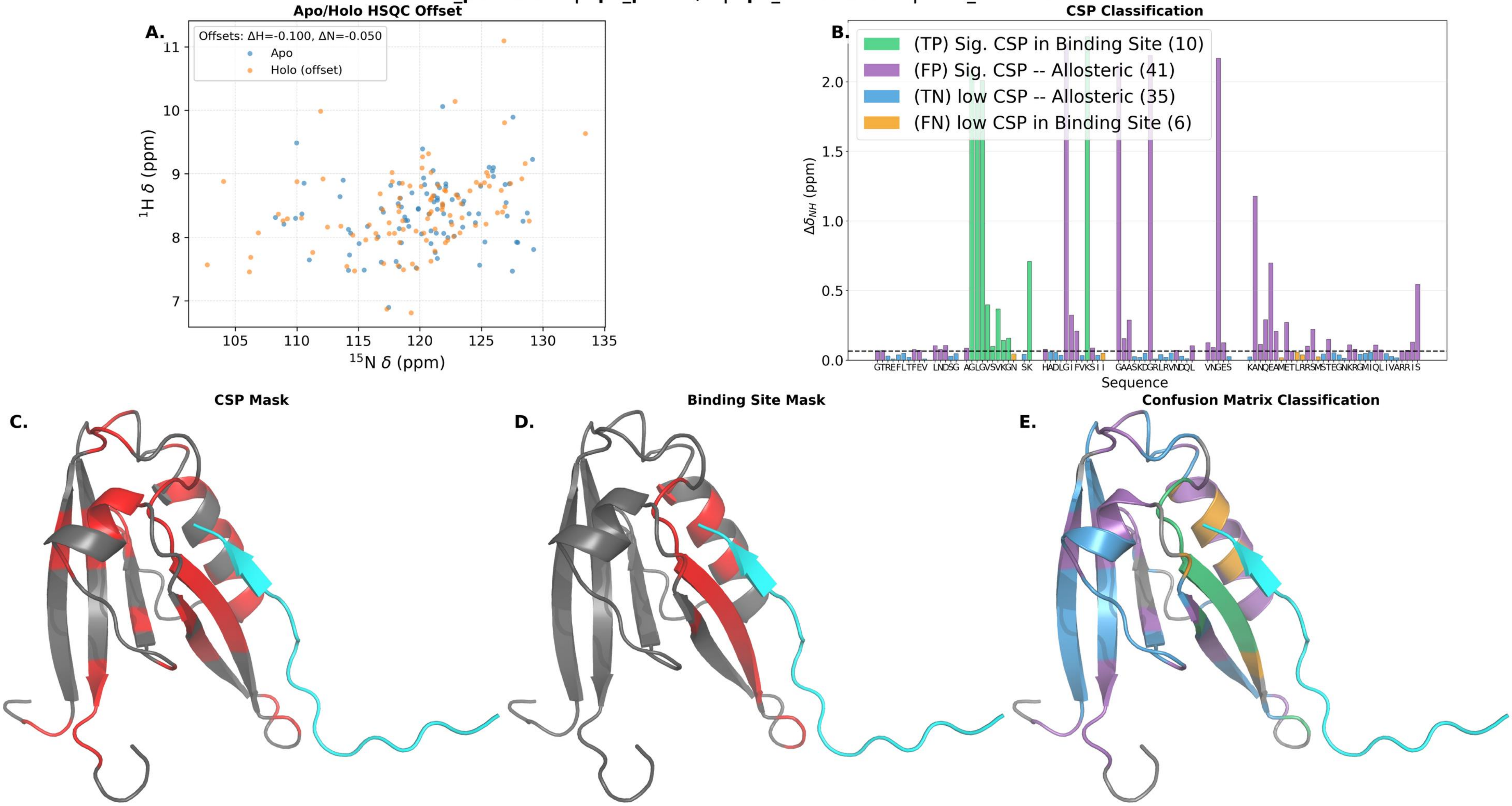

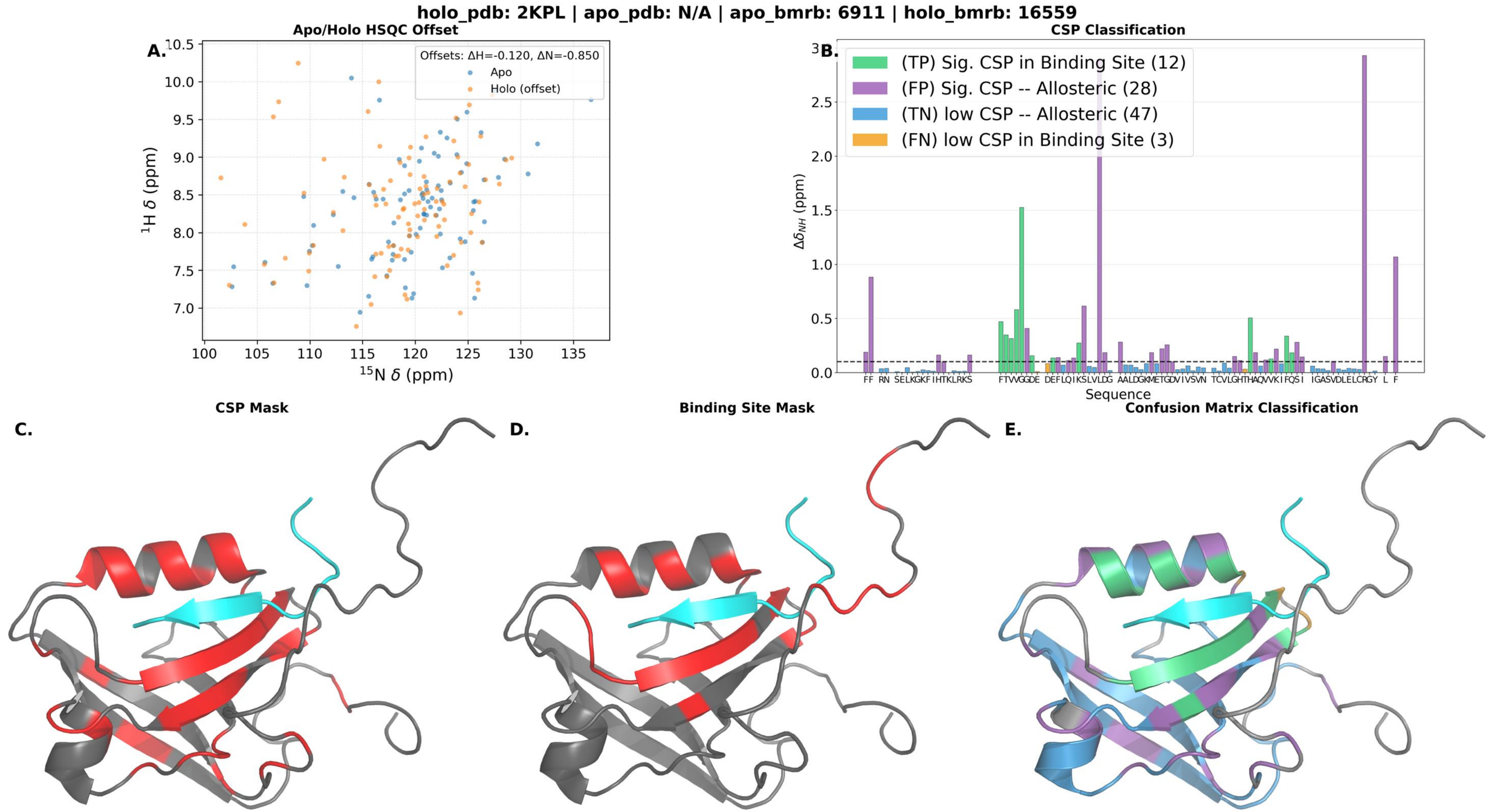

holo_pdb: 2KPL | apo_pdb: N/A | apo_bmrb: 6911 | holo_bmrb: 16559
A.
Apo/Holo HSQC Offset
Offsets: ΔH=-0.120, ΔN=-0.850
Apo
Holo (offset)
¹H δ (ppm)
¹⁵N δ (ppm)
B.
CSP Classification
(TP) Sig. CSP in Binding Site (12)
(FP) Sig. CSP -- Allosteric (28)
(TN) low CSP -- Allosteric (47)
(FN) low CSP in Binding Site (3)
ΔδNH (ppm)
Sequence
C.
CSP Mask
D.
Binding Site Mask
E.
Confusion Matrix Classification

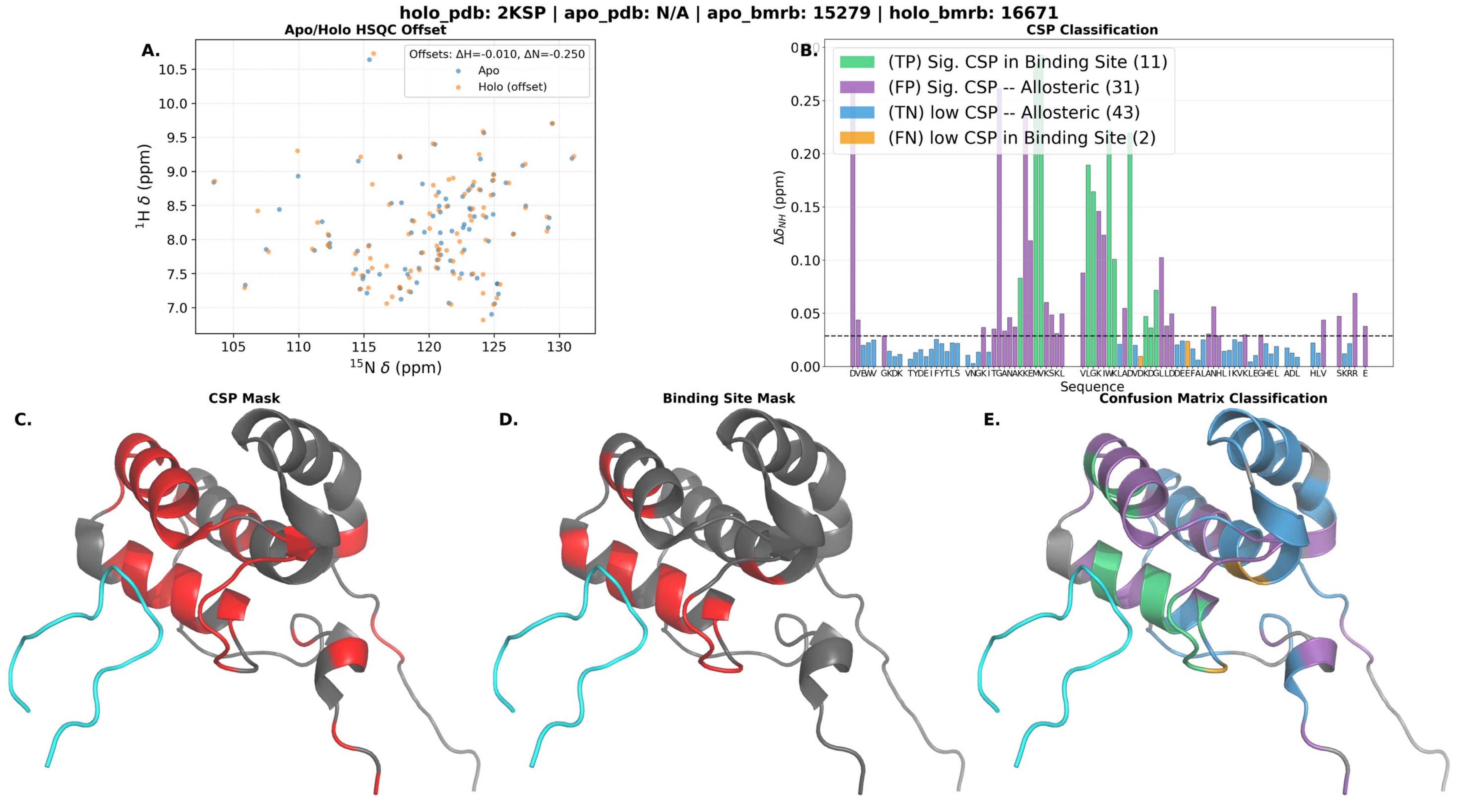
holo_pdb: 2KSP | apo_pdb: N/A | apo_bmrb: 15279 | holo_bmrb: 16671
A.
Apo/Holo HSQC Offset
Offsets: ΔH=-0.010, ΔN=-0.250
Apo
Holo (offset)
¹H δ (ppm)
¹⁵N δ (ppm)
B.
CSP Classification
(TP) Sig. CSP in Binding Site (11)
(FP) Sig. CSP -- Allosteric (31)
(TN) low CSP -- Allosteric (43)
(FN) low CSP in Binding Site (2)
Δδ_NH (ppm)
Sequence
C.
CSP Mask
D.
Binding Site Mask
E.
Confusion Matrix Classification

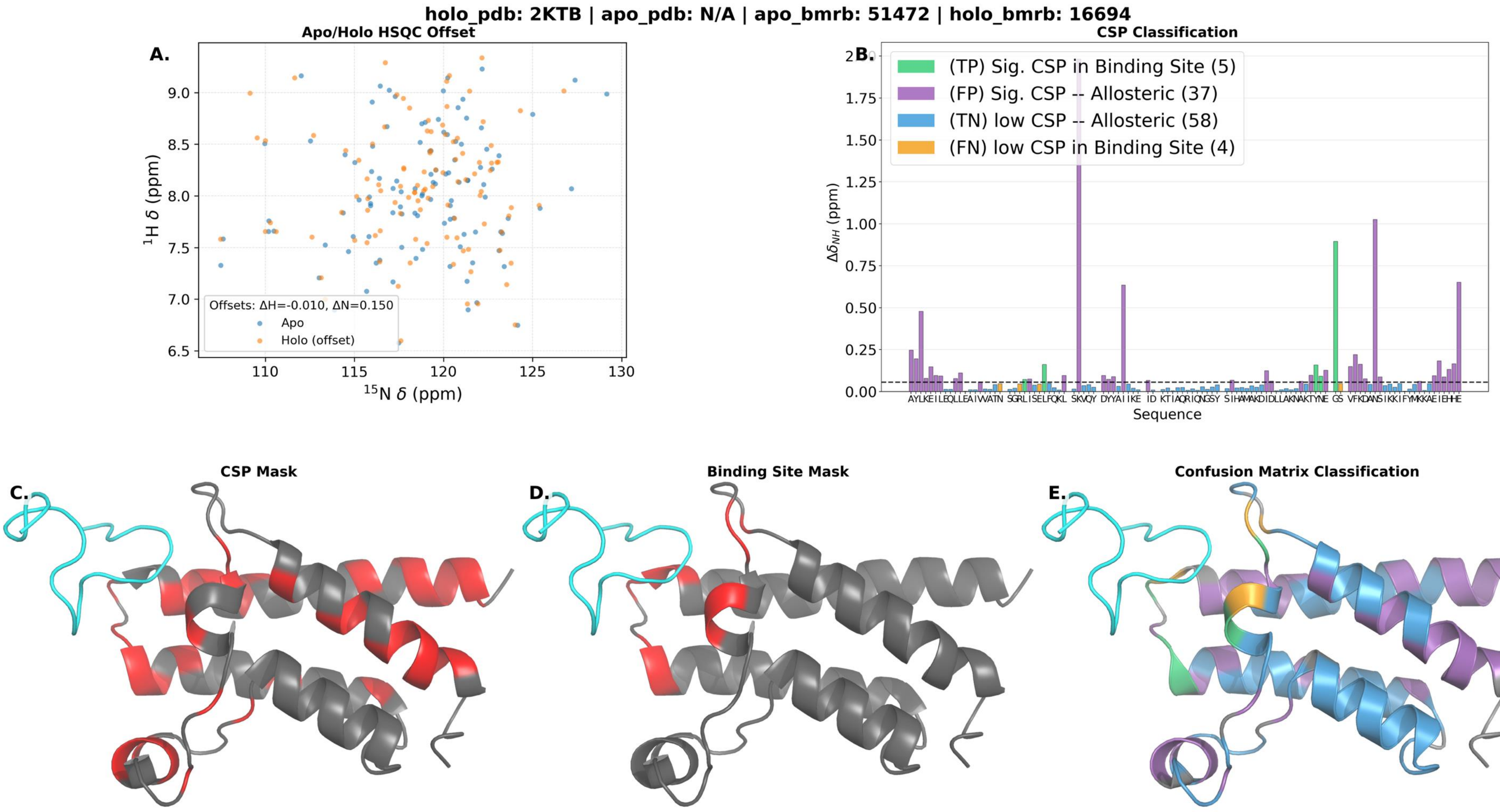

holo_pdb: 2KTB | apo_pdb: N/A | apo_bmrb: 51472 | holo_bmrb: 16694
A.
Apo/Holo HSQC Offset
Offsets: ΔH=-0.010, ΔN=0.150
Apo
Holo (offset)
$^{1}$H δ (ppm)
$^{15}$N δ (ppm)
B.
CSP Classification
(TP) Sig. CSP in Binding Site (5)
(FP) Sig. CSP -- Allosteric (37)
(TN) low CSP -- Allosteric (58)
(FN) low CSP in Binding Site (4)
$\Delta\delta_{NH}$ (ppm)
Sequence
C.
CSP Mask
D.
Binding Site Mask
E.
Confusion Matrix Classification

holo_pdb: 2KVM | apo_pdb: N/A | apo_bmrb: 15674 | holo_bmrb: 16778

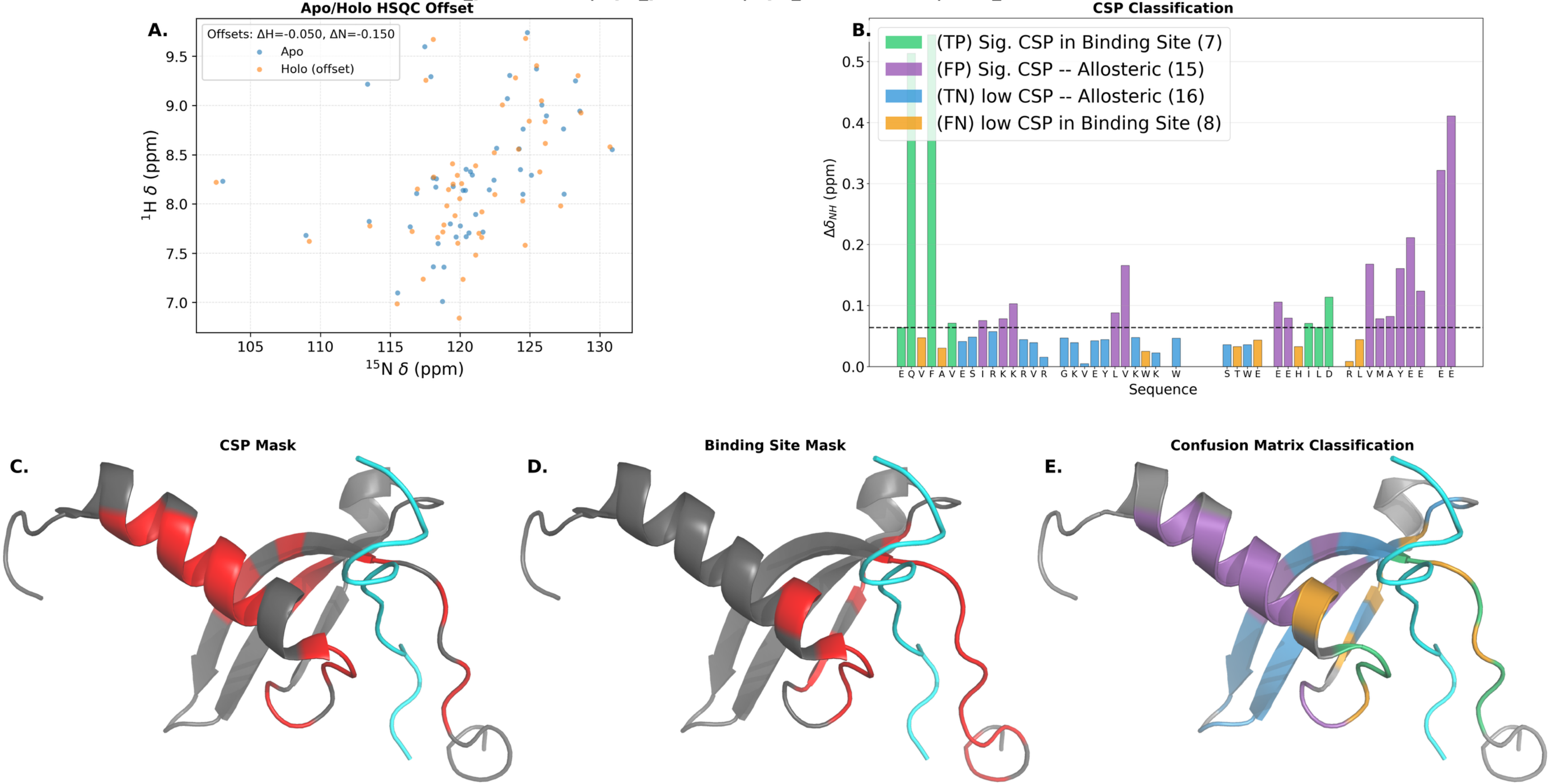

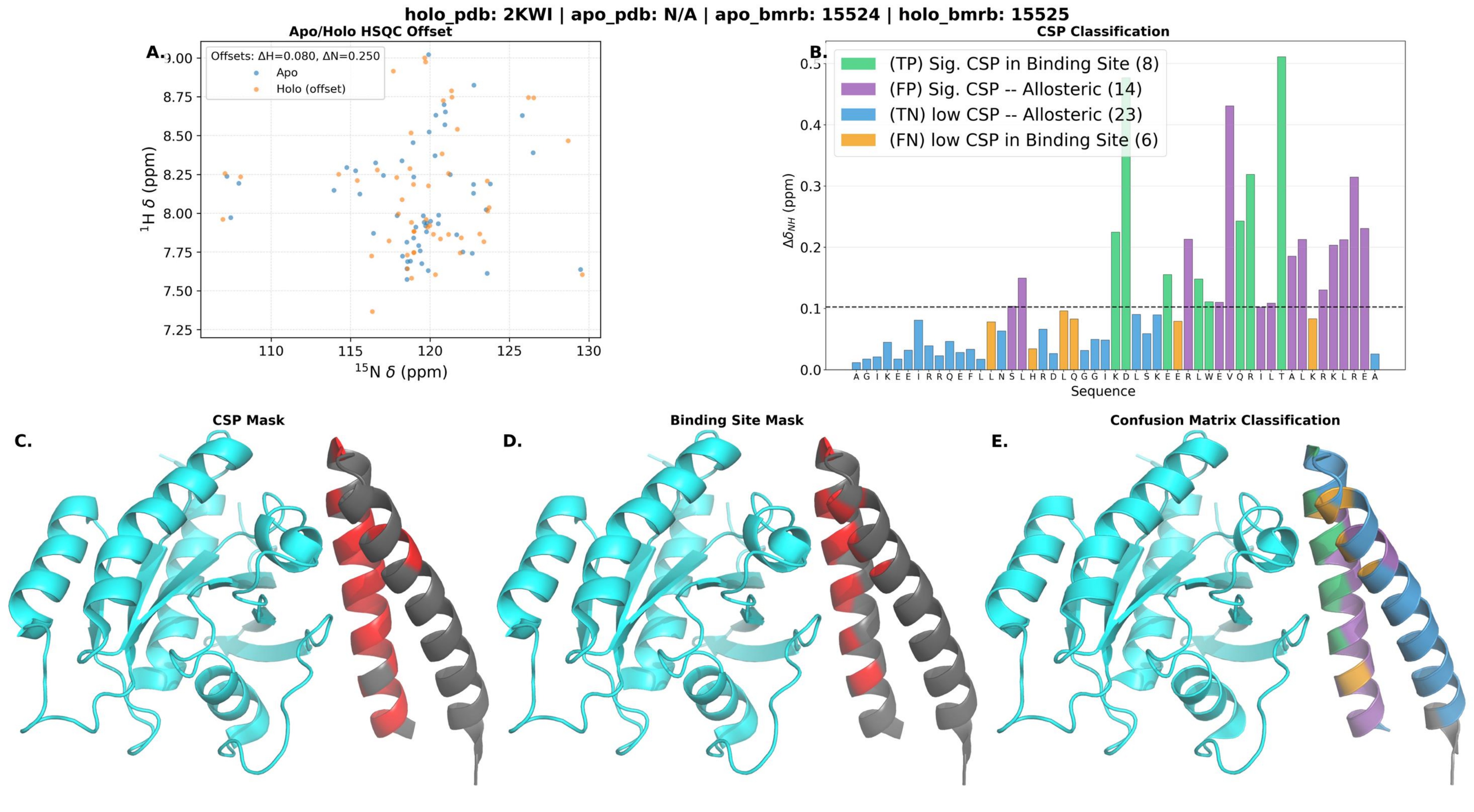
holo_pdb: 2KWI | apo_pdb: N/A | apo_bmrb: 15524 | holo_bmrb: 15525
A.
Apo/Holo HSQC Offset
Offsets: ΔH=0.080, ΔN=0.250
Apo
Holo (offset)
$^{1}$H δ (ppm)
$^{15}$N δ (ppm)
9.00
8.75
8.50
8.25
8.00
7.75
7.50
7.25
110
115
120
125
130
B.
CSP Classification
(TP) Sig. CSP in Binding Site (8)
(FP) Sig. CSP -- Allosteric (14)
(TN) low CSP -- Allosteric (23)
(FN) low CSP in Binding Site (6)
$\Delta\delta_{NH}$ (ppm)
0.5
0.4
0.3
0.2
0.1
0.0
A G I K E E I R R Q E F L L N S L H R D L Q G G I K D L S K E E R L W E V Q R I L T A L K R K L R E A
Sequence
C.
CSP Mask
D.
Binding Site Mask
E.
Confusion Matrix Classification

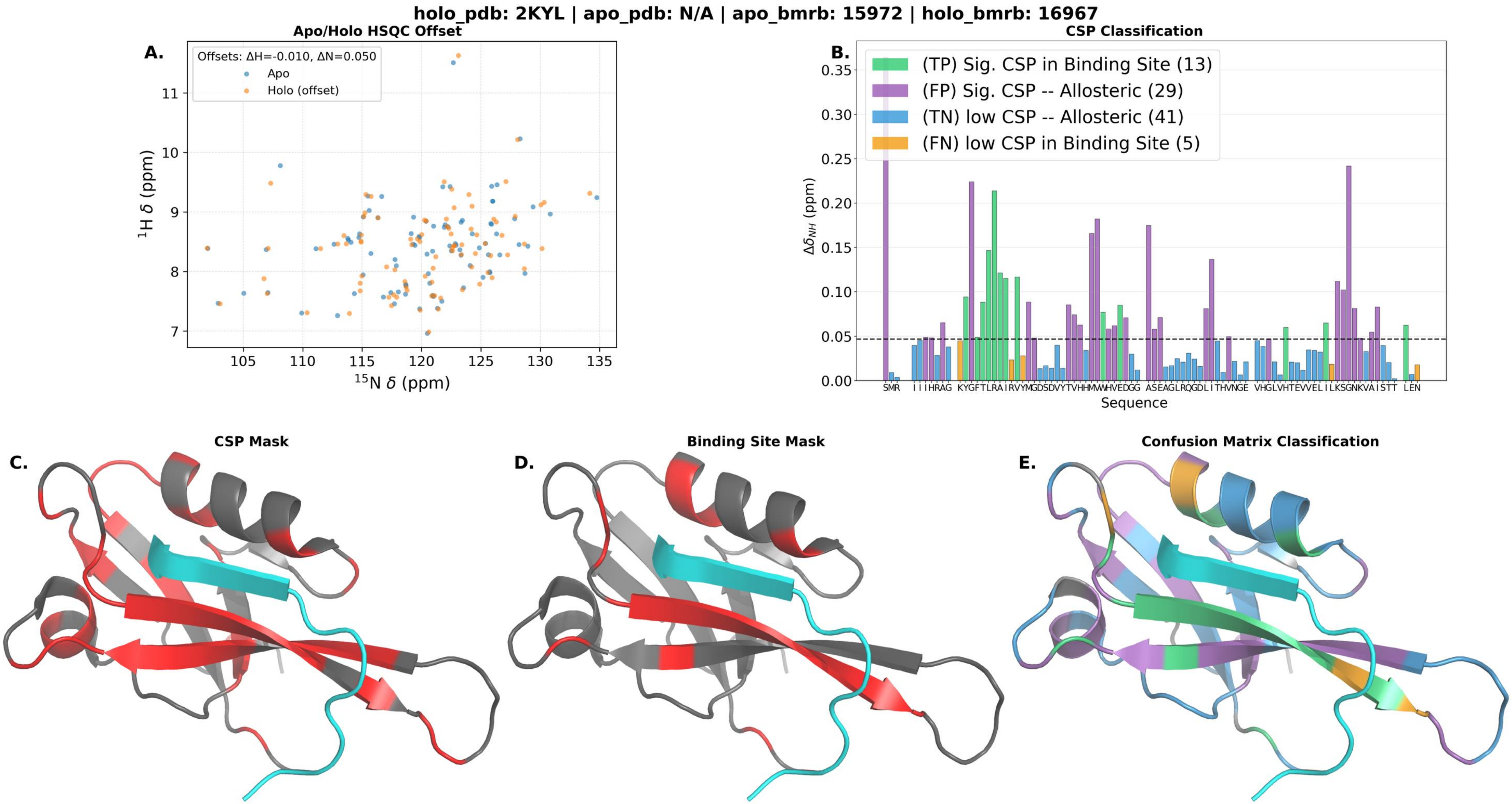
holo_pdb: 2KYL | apo_pdb: N/A | apo_bmrb: 15972 | holo_bmrb: 16967
A.
Apo/Holo HSQC Offset
Offsets: ΔH=-0.010, ΔN=0.050
Apo
Holo (offset)
1H δ (ppm)
15N δ (ppm)
B.
CSP Classification
(TP) Sig. CSP in Binding Site (13)
(FP) Sig. CSP -- Allosteric (29)
(TN) low CSP -- Allosteric (41)
(FN) low CSP in Binding Site (5)
ΔδNH (ppm)
Sequence
C.
CSP Mask
D.
Binding Site Mask
E.
Confusion Matrix Classification

holo_pdb: 2KZU | apo_pdb: N/A | apo_bmrb: 17018 | holo_bmrb: 17019

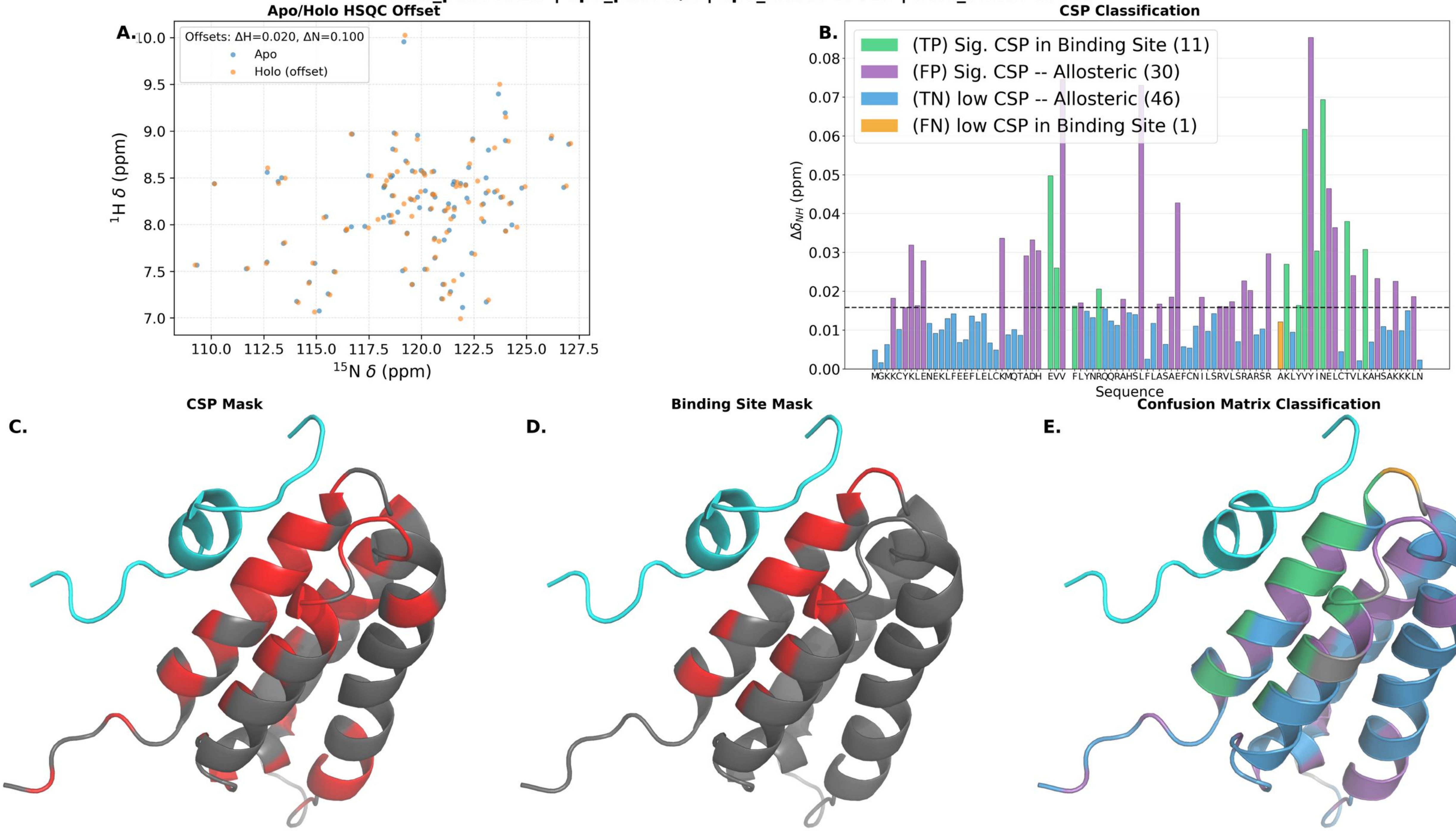

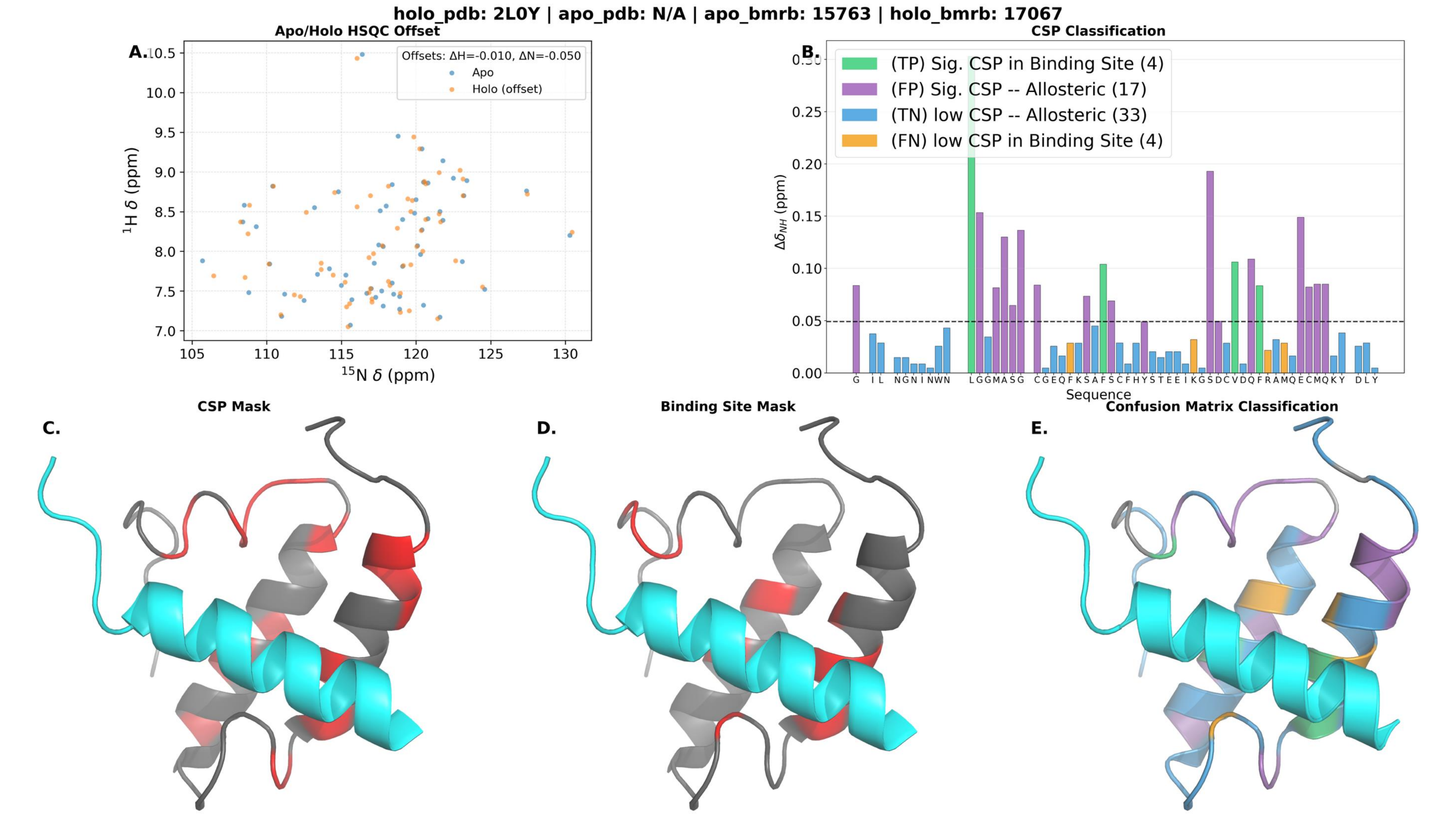
holo_pdb: 2L0Y | apo_pdb: N/A | apo_bmrb: 15763 | holo_bmrb: 17067
A.
Apo/Holo HSQC Offset
Offsets: ΔH=-0.010, ΔN=-0.050
Apo
Holo (offset)
$^{1}$H δ (ppm)
$^{15}$N δ (ppm)
B.
CSP Classification
(TP) Sig. CSP in Binding Site (4)
(FP) Sig. CSP -- Allosteric (17)
(TN) low CSP -- Allosteric (33)
(FN) low CSP in Binding Site (4)
$\Delta\delta_{NH}$ (ppm)
Sequence
C.
CSP Mask
D.
Binding Site Mask
E.
Confusion Matrix Classification

**holo_pdb: 2L29 | apo_pdb: N/A | apo_bmrb: 17128 | holo_bmrb: 17127**

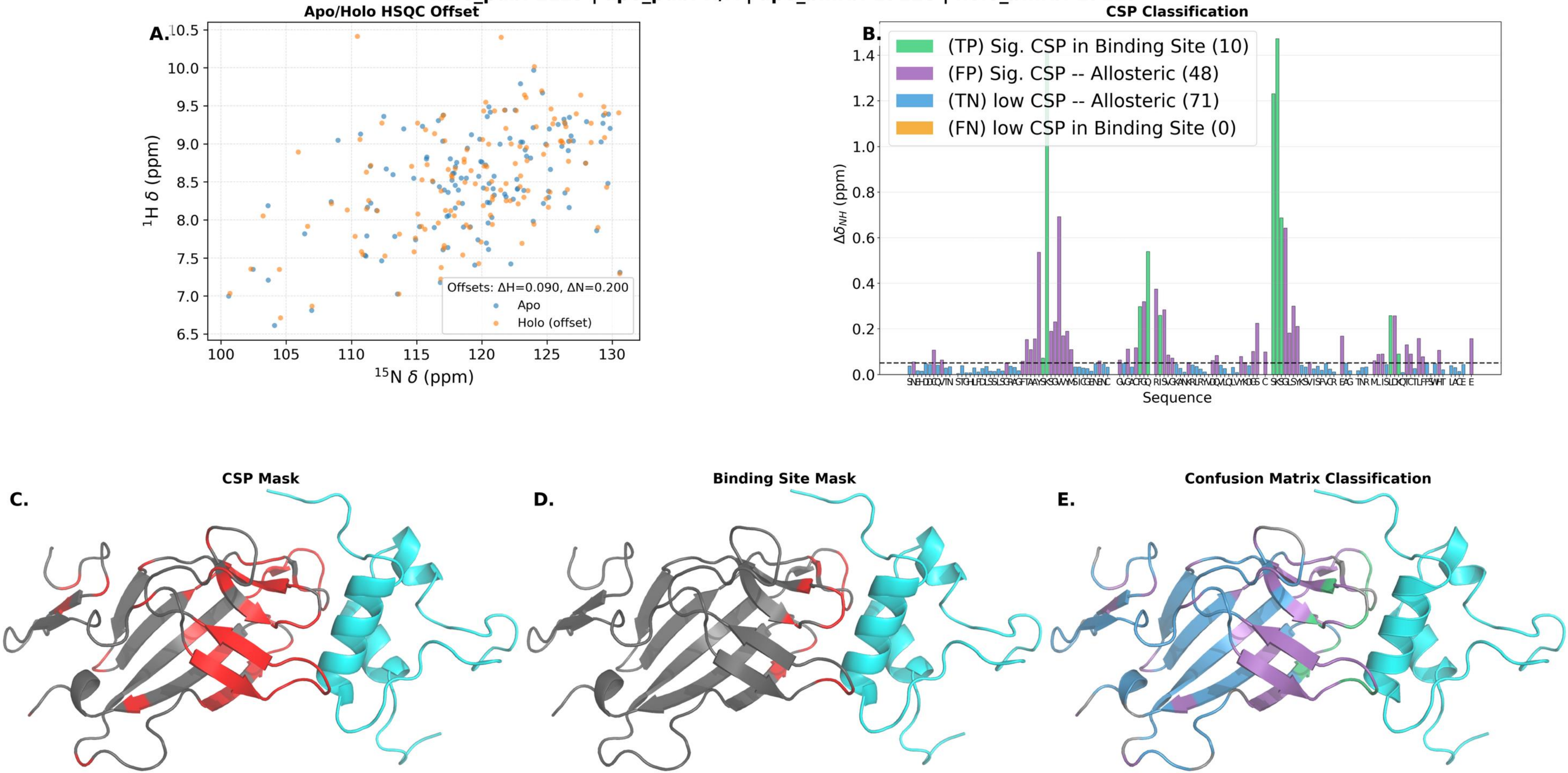

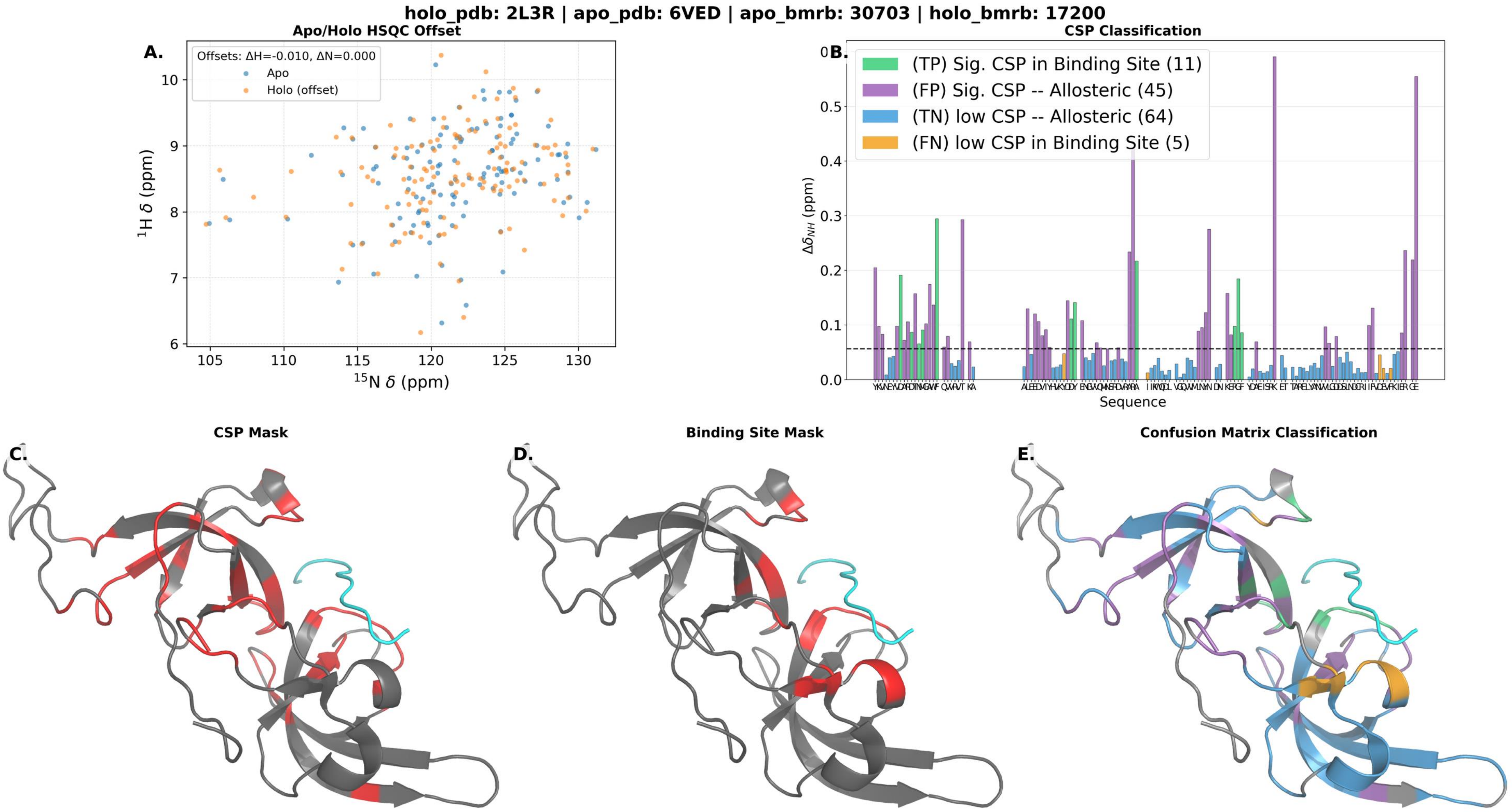

holo_pdb: 2L3R | apo_pdb: 6VED | apo_bmrb: 30703 | holo_bmrb: 17200
A.
Apo/Holo HSQC Offset
Offsets: ΔH=-0.010, ΔN=0.000
Apo
Holo (offset)
$^{1}$H δ (ppm)
$^{15}$N δ (ppm)
B.
CSP Classification
(TP) Sig. CSP in Binding Site (11)
(FP) Sig. CSP -- Allosteric (45)
(TN) low CSP -- Allosteric (64)
(FN) low CSP in Binding Site (5)
$\Delta\delta_{NH}$ (ppm)
Sequence
C.
CSP Mask
D.
Binding Site Mask
E.
Confusion Matrix Classification

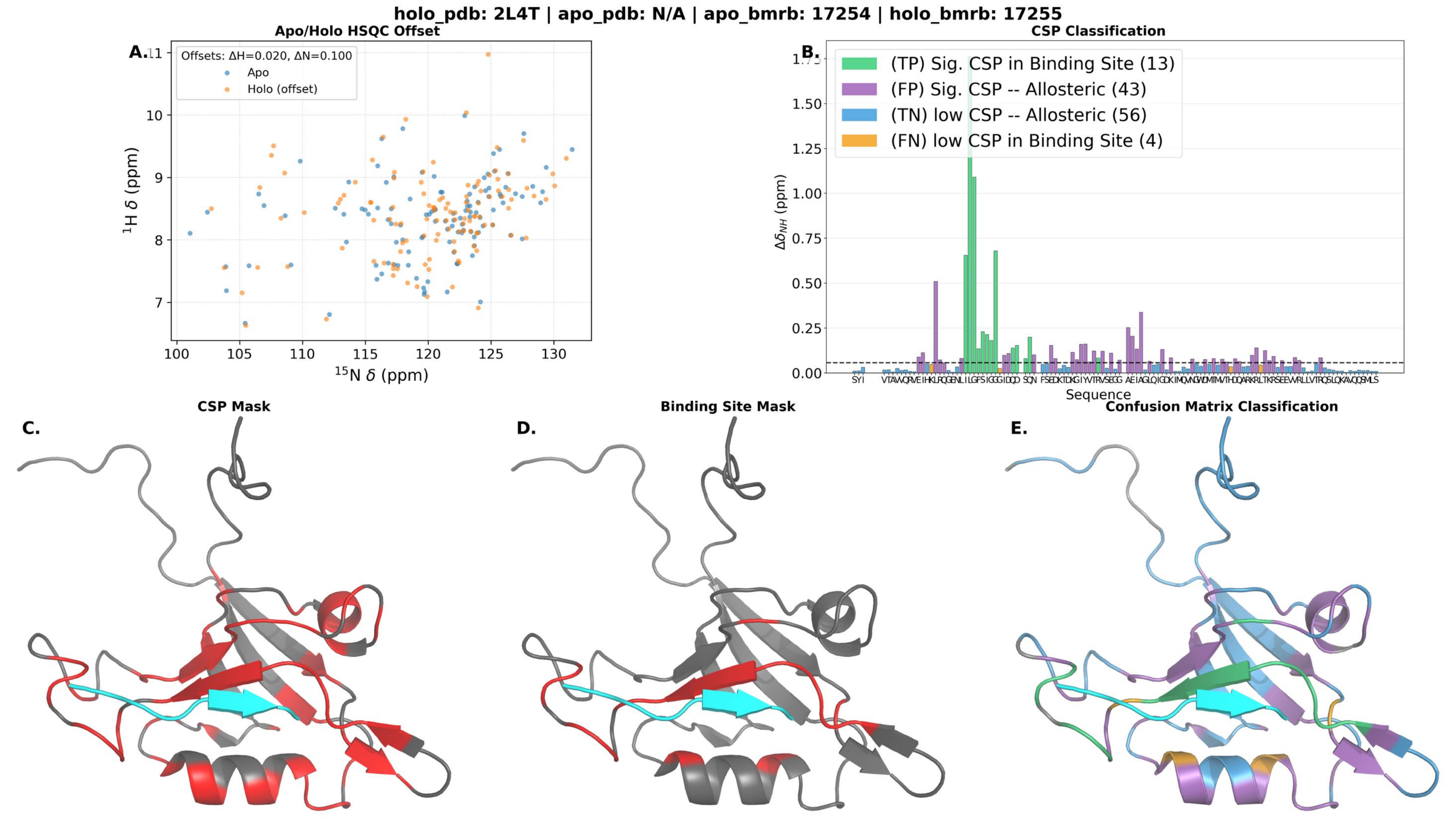

holo_pdb: 2L4T | apo_pdb: N/A | apo_bmrb: 17254 | holo_bmrb: 17255
A.
Apo/Holo HSQC Offset
Offsets: ΔH=0.020, ΔN=0.100
Apo
Holo (offset)
1H δ (ppm)
15N δ (ppm)
B.
CSP Classification
(TP) Sig. CSP in Binding Site (13)
(FP) Sig. CSP -- Allosteric (43)
(TN) low CSP -- Allosteric (56)
(FN) low CSP in Binding Site (4)
ΔδNH (ppm)
Sequence
C.
CSP Mask
D.
Binding Site Mask
E.
Confusion Matrix Classification

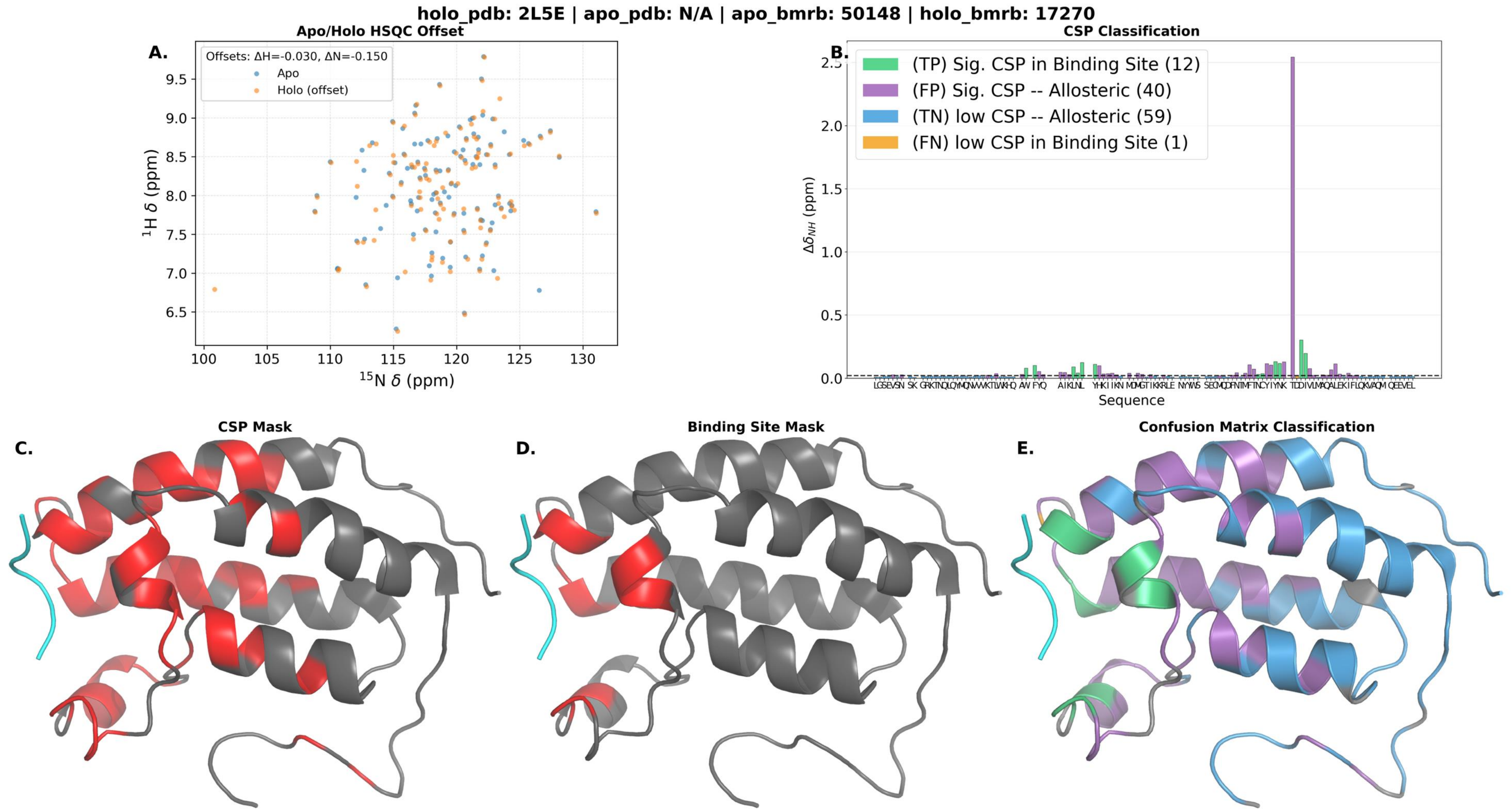

holo_pdb: 2L5E | apo_pdb: N/A | apo_bmrb: 50148 | holo_bmrb: 17270
A.
Apo/Holo HSQC Offset
Offsets: ΔH=-0.030, ΔN=-0.150
Apo
Holo (offset)
¹H δ (ppm)
¹⁵N δ (ppm)
B.
CSP Classification
(TP) Sig. CSP in Binding Site (12)
(FP) Sig. CSP -- Allosteric (40)
(TN) low CSP -- Allosteric (59)
(FN) low CSP in Binding Site (1)
Δδ_NH (ppm)
Sequence
C.
CSP Mask
D.
Binding Site Mask
E.
Confusion Matrix Classification

**holo_pdb: 2L6E | apo_pdb: N/A | apo_bmrb: 16555 | holo_bmrb: 17307**

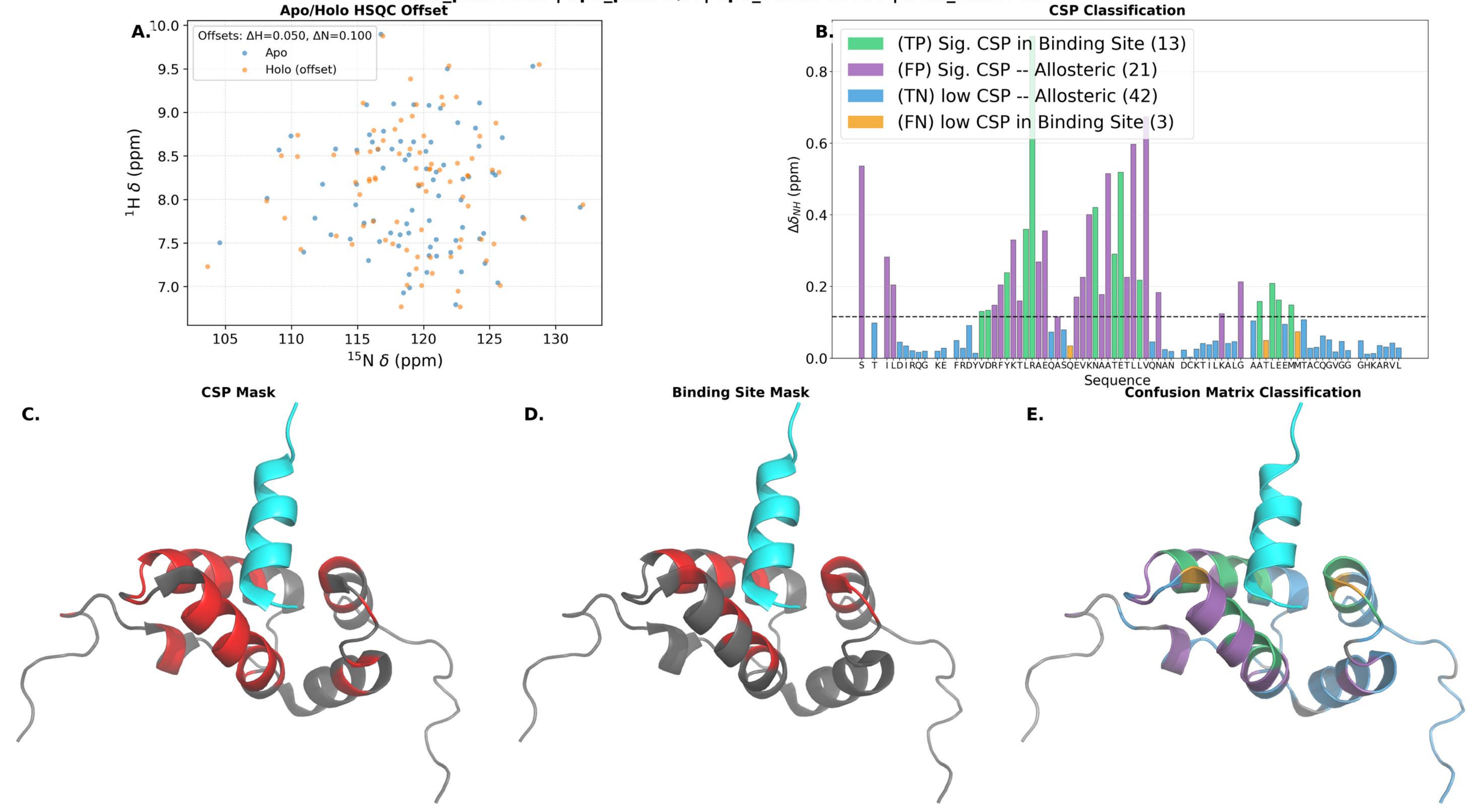

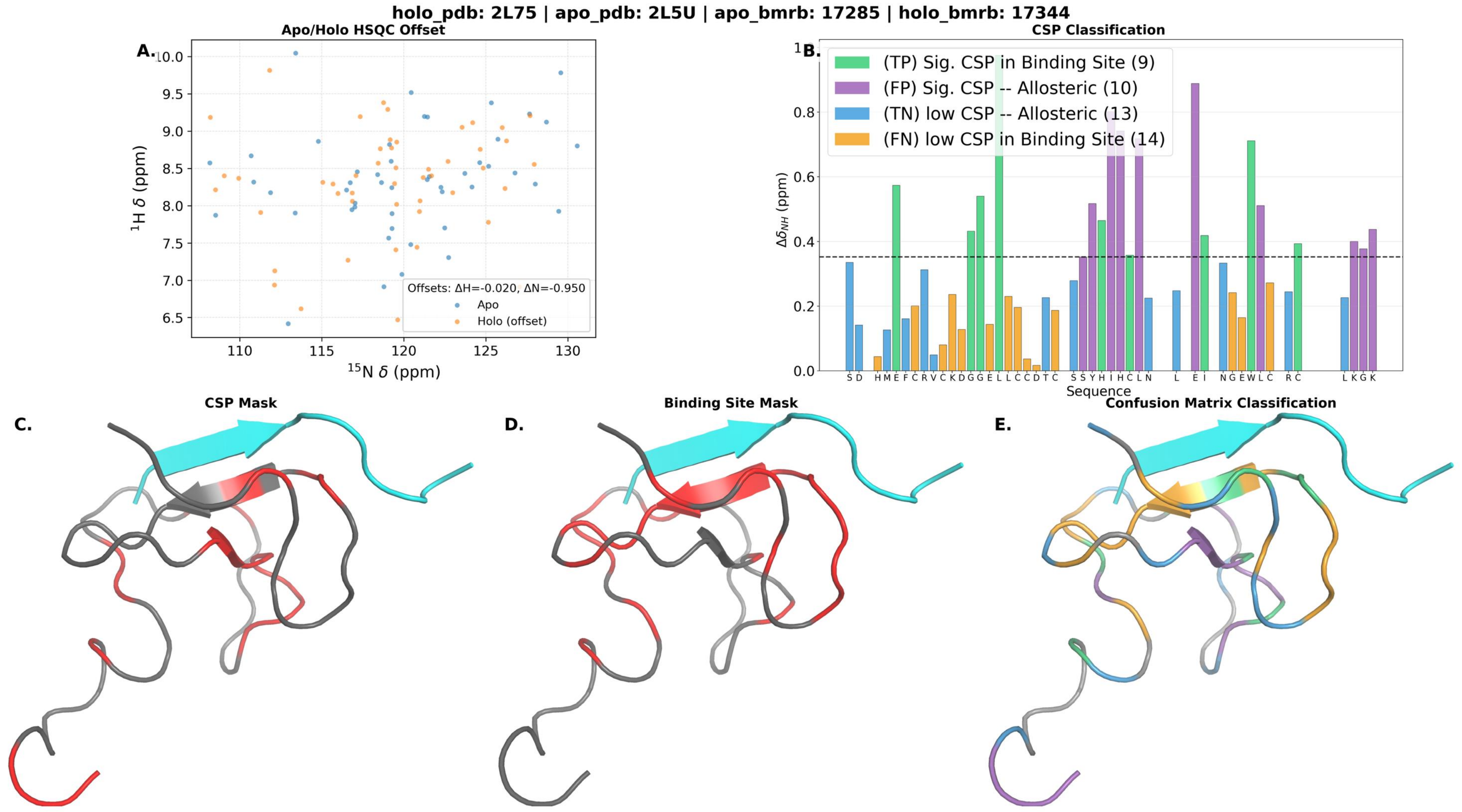

holo_pdb: 2L75 | apo_pdb: 2L5U | apo_bmrb: 17285 | holo_bmrb: 17344
A.
Apo/Holo HSQC Offset
Offsets: ΔH=-0.020, ΔN=-0.950
Apo
Holo (offset)
$^{1}$H δ (ppm)
$^{15}$N δ (ppm)
B.
CSP Classification
(TP) Sig. CSP in Binding Site (9)
(FP) Sig. CSP -- Allosteric (10)
(TN) low CSP -- Allosteric (13)
(FN) low CSP in Binding Site (14)
$\Delta\delta_{NH}$ (ppm)
Sequence
C.
CSP Mask
D.
Binding Site Mask
E.
Confusion Matrix Classification

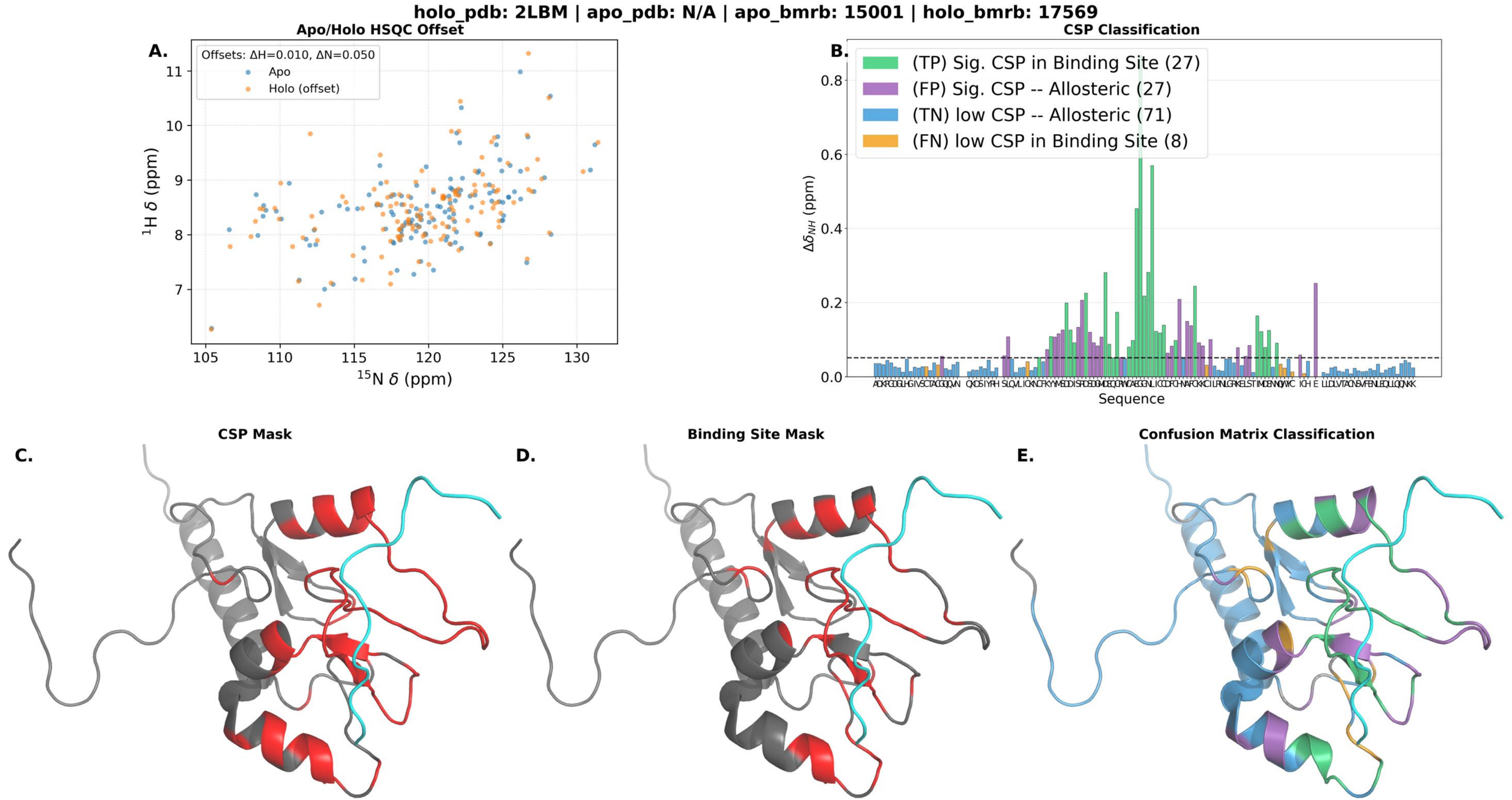
holo_pdb: 2LBM | apo_pdb: N/A | apo_bmrb: 15001 | holo_bmrb: 17569
A.
Apo/Holo HSQC Offset
Offsets: ΔH=0.010, ΔN=0.050
Apo
Holo (offset)
$^{1}$H δ (ppm)
$^{15}$N δ (ppm)
B.
CSP Classification
(TP) Sig. CSP in Binding Site (27)
(FP) Sig. CSP -- Allosteric (27)
(TN) low CSP -- Allosteric (71)
(FN) low CSP in Binding Site (8)
$\Delta\delta_{NH}$ (ppm)
Sequence
C.
CSP Mask
D.
Binding Site Mask
E.
Confusion Matrix Classification

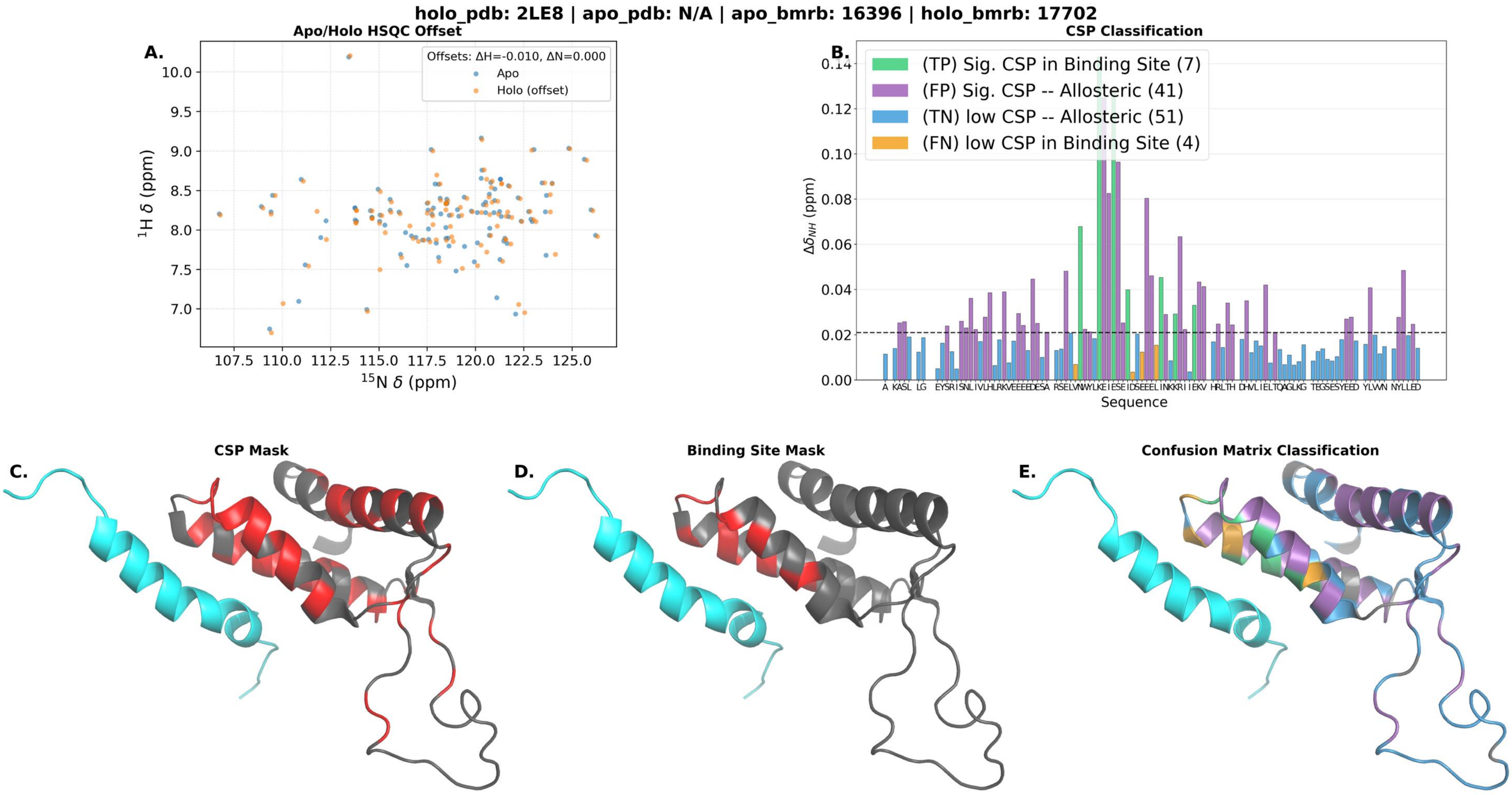
holo_pdb: 2LE8 | apo_pdb: N/A | apo_bmrb: 16396 | holo_bmrb: 17702
Apo/Holo HSQC Offset
A.
Offsets: ΔH=-0.010, ΔN=0.000
Apo
Holo (offset)
1H δ (ppm)
15N δ (ppm)
CSP Classification
B.
(TP) Sig. CSP in Binding Site (7)
(FP) Sig. CSP -- Allosteric (41)
(TN) low CSP -- Allosteric (51)
(FN) low CSP in Binding Site (4)
Δδ_NH (ppm)
Sequence
C.
CSP Mask
D.
Binding Site Mask
E.
Confusion Matrix Classification

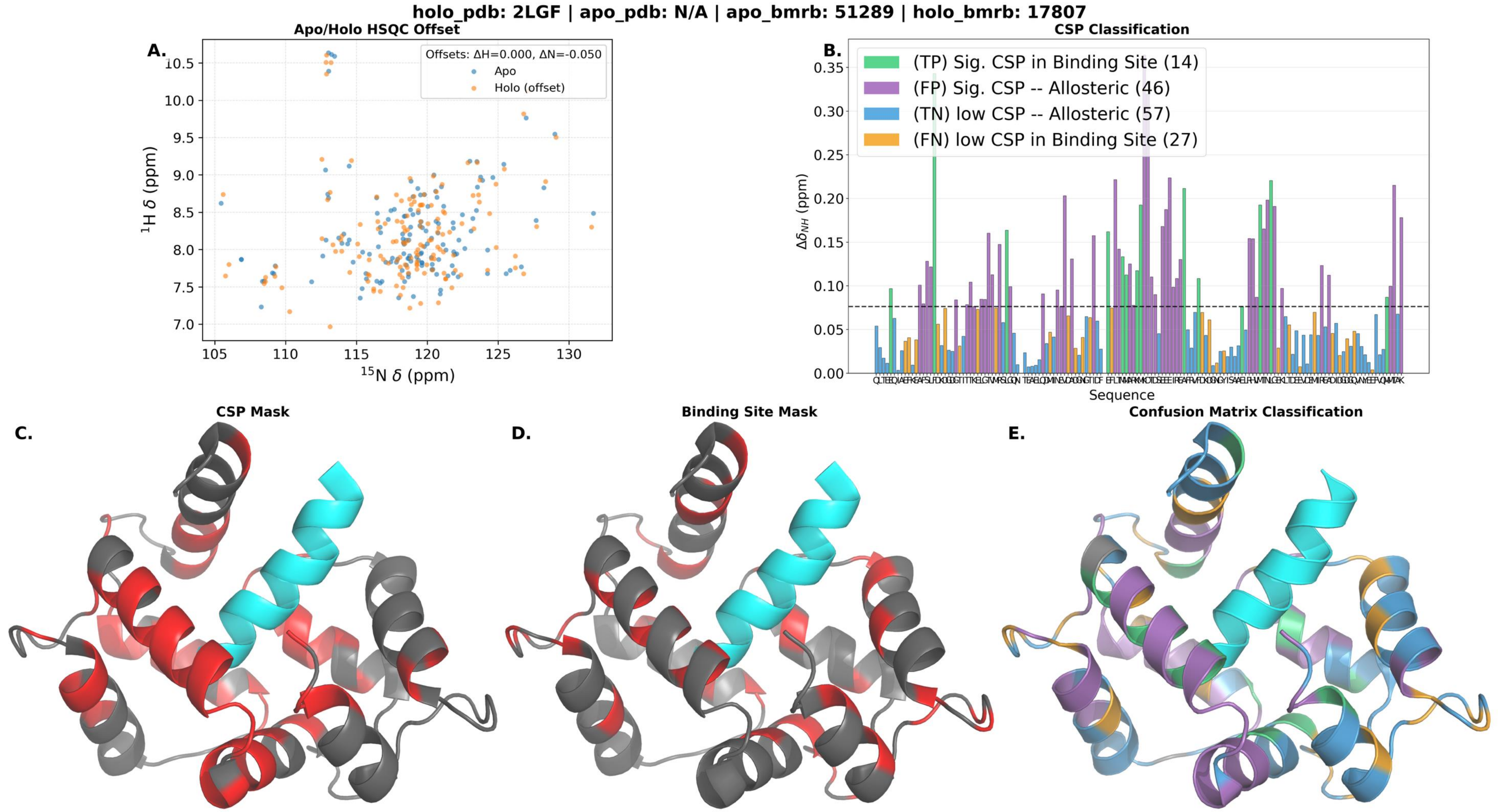

holo_pdb: 2LGF | apo_pdb: N/A | apo_bmrb: 51289 | holo_bmrb: 17807
Apo/Holo HSQC Offset
A.
Offsets: ΔH=0.000, ΔN=-0.050
Apo
Holo (offset)
10.5
10.0
9.5
9.0
8.5
8.0
7.5
7.0
105
110
115
120
125
130
$^{1}$H δ (ppm)
$^{15}$N δ (ppm)
CSP Classification
B.
(TP) Sig. CSP in Binding Site (14)
(FP) Sig. CSP -- Allosteric (46)
(TN) low CSP -- Allosteric (57)
(FN) low CSP in Binding Site (27)
0.35
0.30
0.25
0.20
0.15
0.10
0.05
0.00
$\Delta\delta_{NH}$ (ppm)
Sequence
C.
CSP Mask
D.
Binding Site Mask
E.
Confusion Matrix Classification

**holo_pdb: 2LGG | apo_pdb: N/A | apo_bmrb: 17813 | holo_bmrb: 17808**

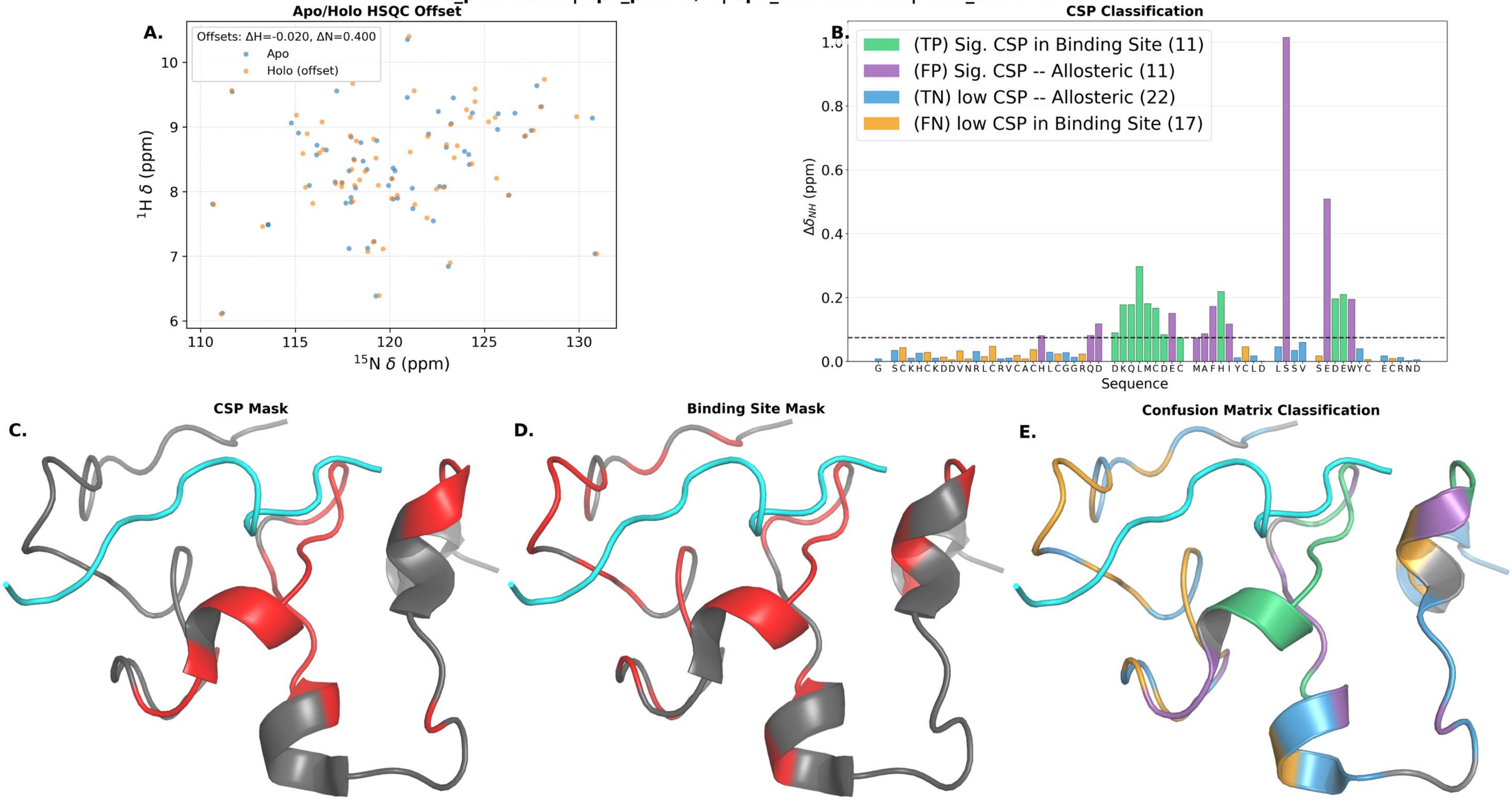

holo_pdb: 2LGK | apo_pdb: N/A | apo_bmrb: 17813 | holo_bmrb: 17812

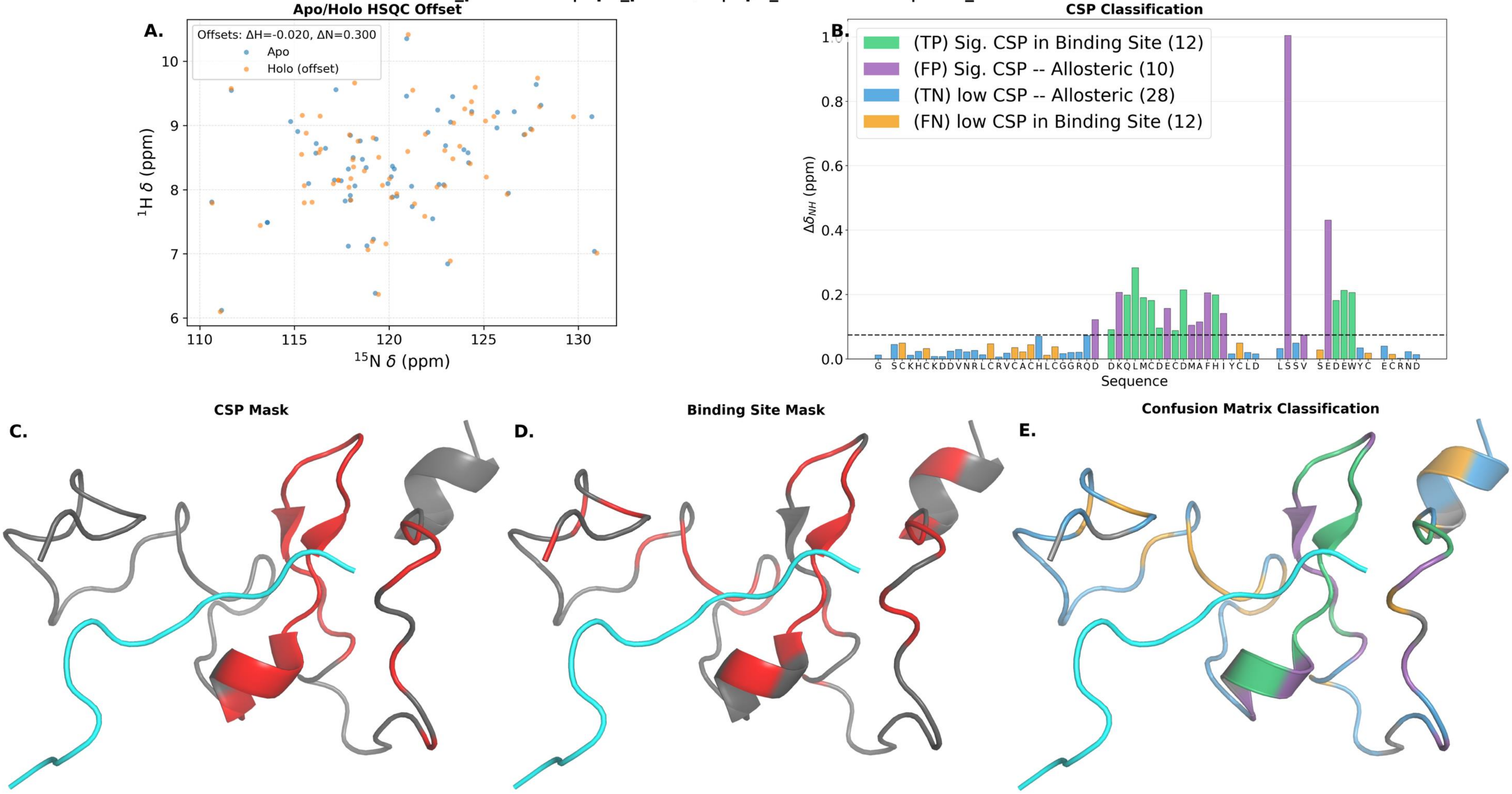

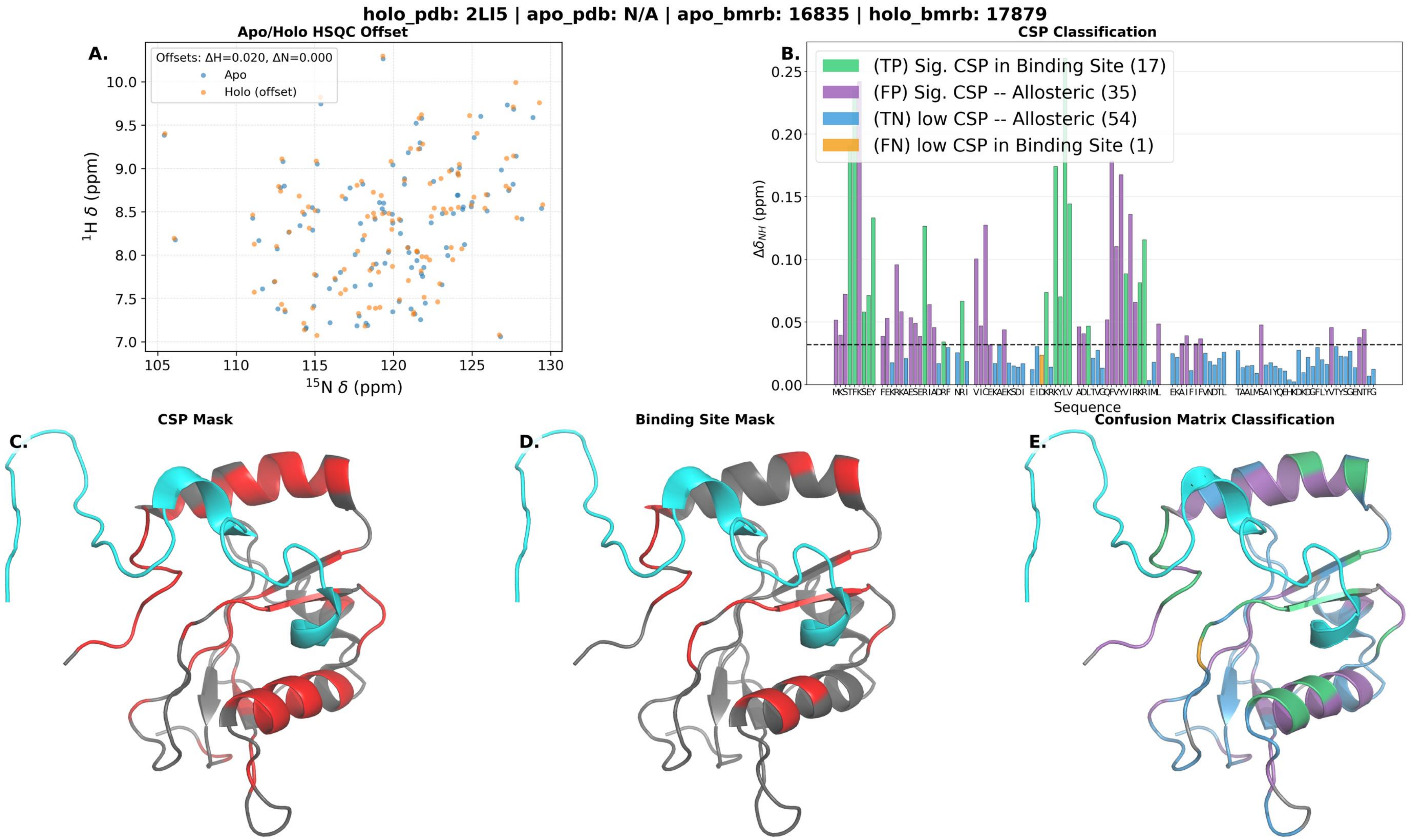

holo_pdb: 2LI5 | apo_pdb: N/A | apo_bmrb: 16835 | holo_bmrb: 17879
A.
Apo/Holo HSQC Offset
Offsets: ΔH=0.020, ΔN=0.000
Apo
Holo (offset)
$^{1}$H δ (ppm)
$^{15}$N δ (ppm)
B.
CSP Classification
(TP) Sig. CSP in Binding Site (17)
(FP) Sig. CSP -- Allosteric (35)
(TN) low CSP -- Allosteric (54)
(FN) low CSP in Binding Site (1)
$\Delta\delta_{NH}$ (ppm)
Sequence
C.
CSP Mask
D.
Binding Site Mask
E.
Confusion Matrix Classification

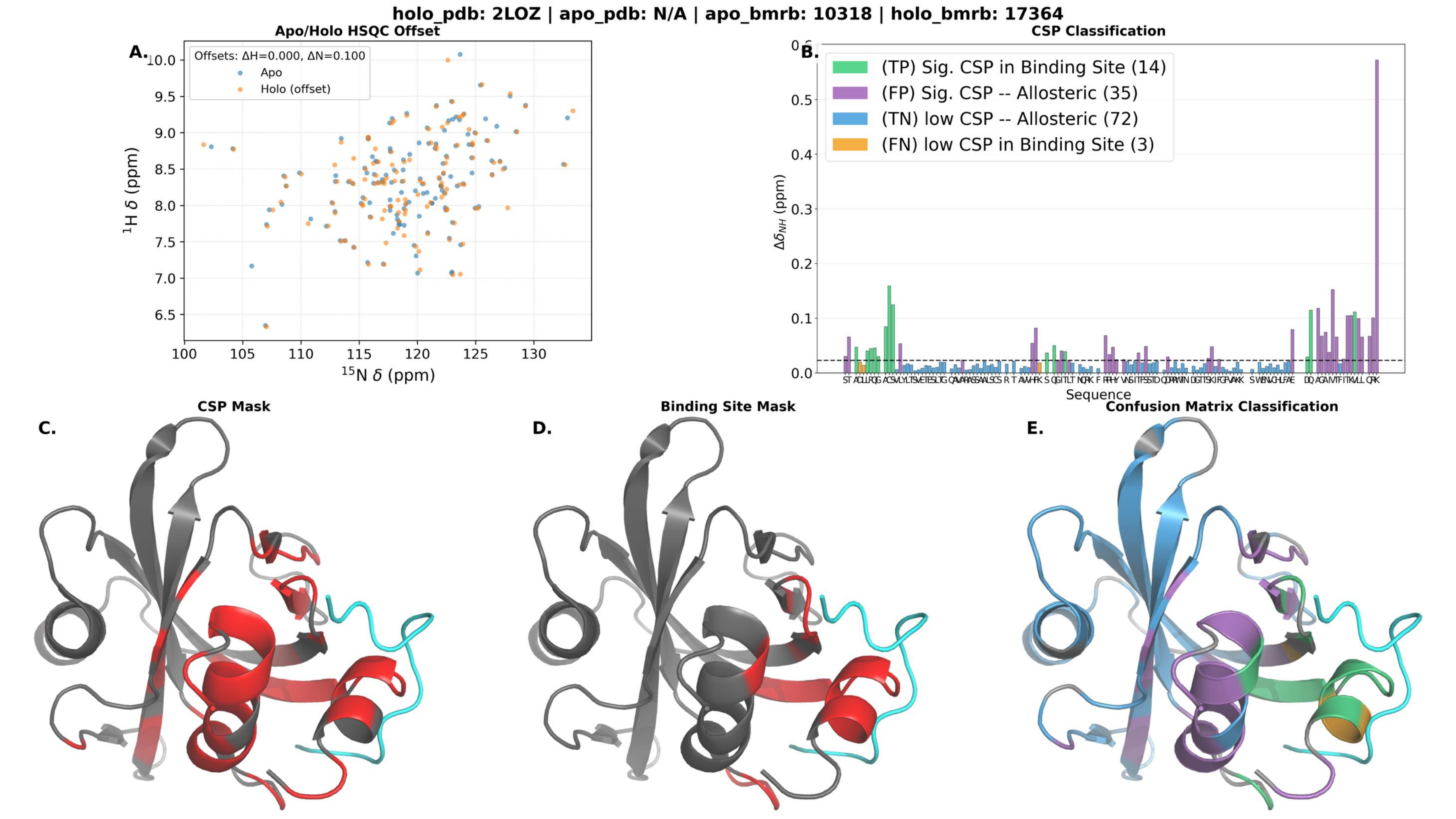
holo_pdb: 2LOZ | apo_pdb: N/A | apo_bmrb: 10318 | holo_bmrb: 17364
Apo/Holo HSQC Offset
A.
Offsets: ΔH=0.000, ΔN=0.100
Apo
Holo (offset)
1H δ (ppm)
15N δ (ppm)
CSP Classification
B.
(TP) Sig. CSP in Binding Site (14)
(FP) Sig. CSP -- Allosteric (35)
(TN) low CSP -- Allosteric (72)
(FN) low CSP in Binding Site (3)
ΔδNH (ppm)
Sequence
CSP Mask
C.
Binding Site Mask
D.
Confusion Matrix Classification
E.

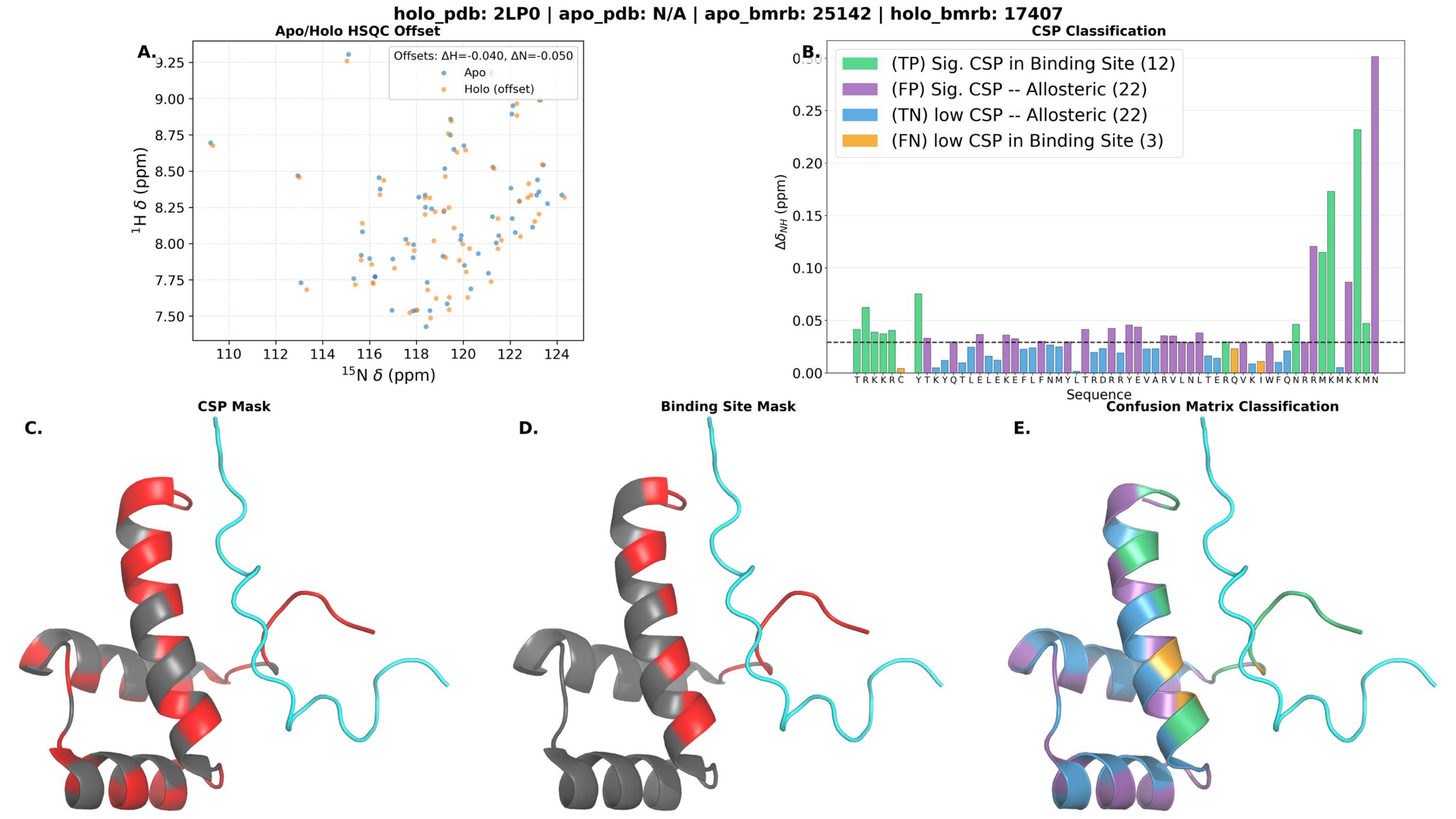
holo_pdb: 2LP0 | apo_pdb: N/A | apo_bmrb: 25142 | holo_bmrb: 17407
A.
Apo/Holo HSQC Offset
Offsets: ΔH=-0.040, ΔN=-0.050
Apo
Holo (offset)
$^{1}$H δ (ppm)
$^{15}$N δ (ppm)
9.25
9.00
8.75
8.50
8.25
8.00
7.75
7.50
110
112
114
116
118
120
122
124
B.
CSP Classification
(TP) Sig. CSP in Binding Site (12)
(FP) Sig. CSP -- Allosteric (22)
(TN) low CSP -- Allosteric (22)
(FN) low CSP in Binding Site (3)
$\Delta\delta_{NH}$ (ppm)
0.30
0.25
0.20
0.15
0.10
0.05
0.00
T R K K R C Y T K Y Q T L E L E K E F L F N M Y L T R D R R Y E V A R V L N L T E R Q V K I W F Q N R R M K M K K M N
Sequence
C.
CSP Mask
D.
Binding Site Mask
E.
Confusion Matrix Classification

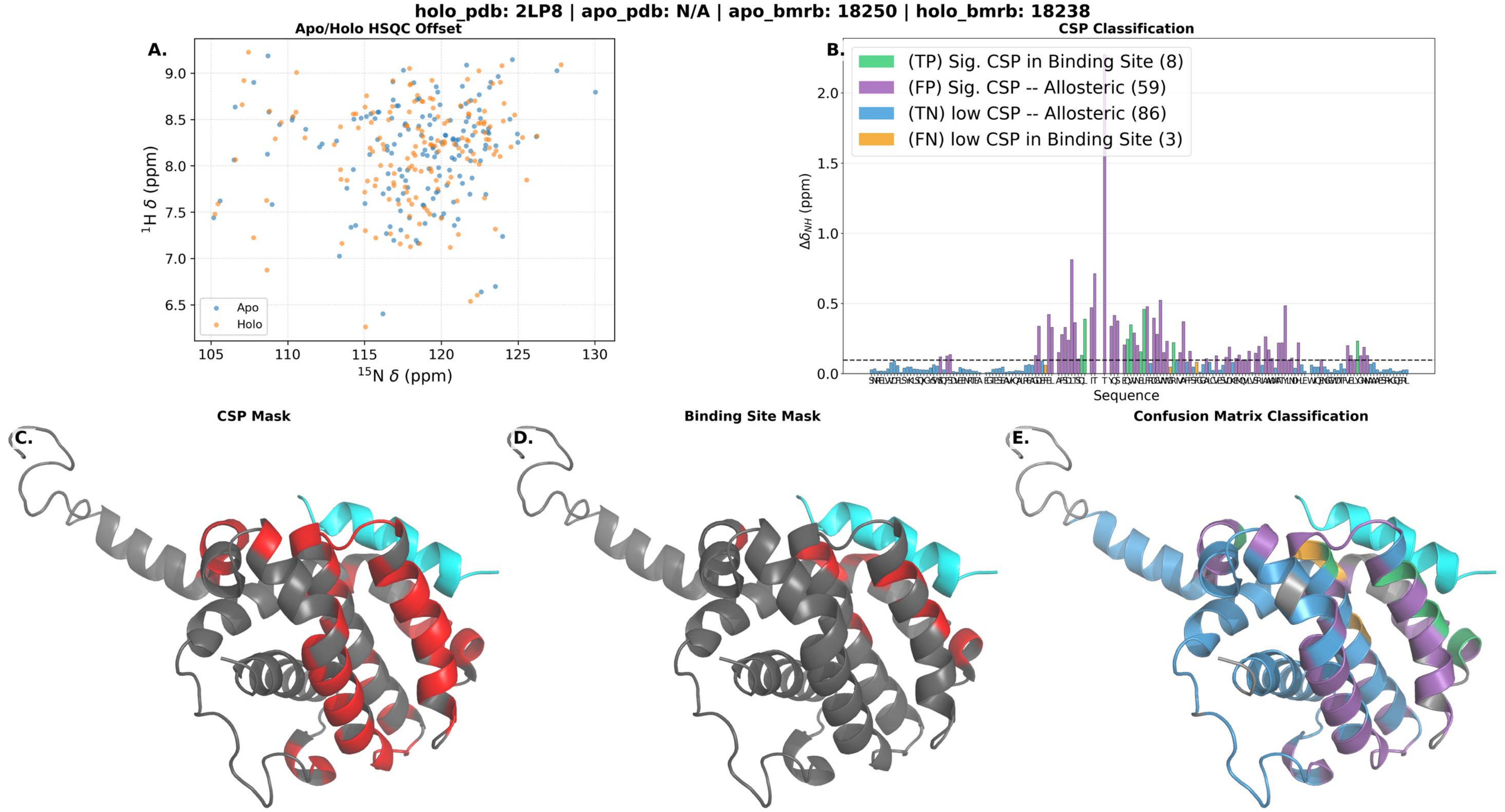
holo_pdb: 2LP8 | apo_pdb: N/A | apo_bmrb: 18250 | holo_bmrb: 18238
Apo/Holo HSQC Offset
A.
$^{1}$H δ (ppm)
$^{15}$N δ (ppm)
Apo
Holo
CSP Classification
B.
(TP) Sig. CSP in Binding Site (8)
(FP) Sig. CSP -- Allosteric (59)
(TN) low CSP -- Allosteric (86)
(FN) low CSP in Binding Site (3)
$\Delta\delta_{NH}$ (ppm)
Sequence
CSP Mask
C.
Binding Site Mask
D.
Confusion Matrix Classification
E.

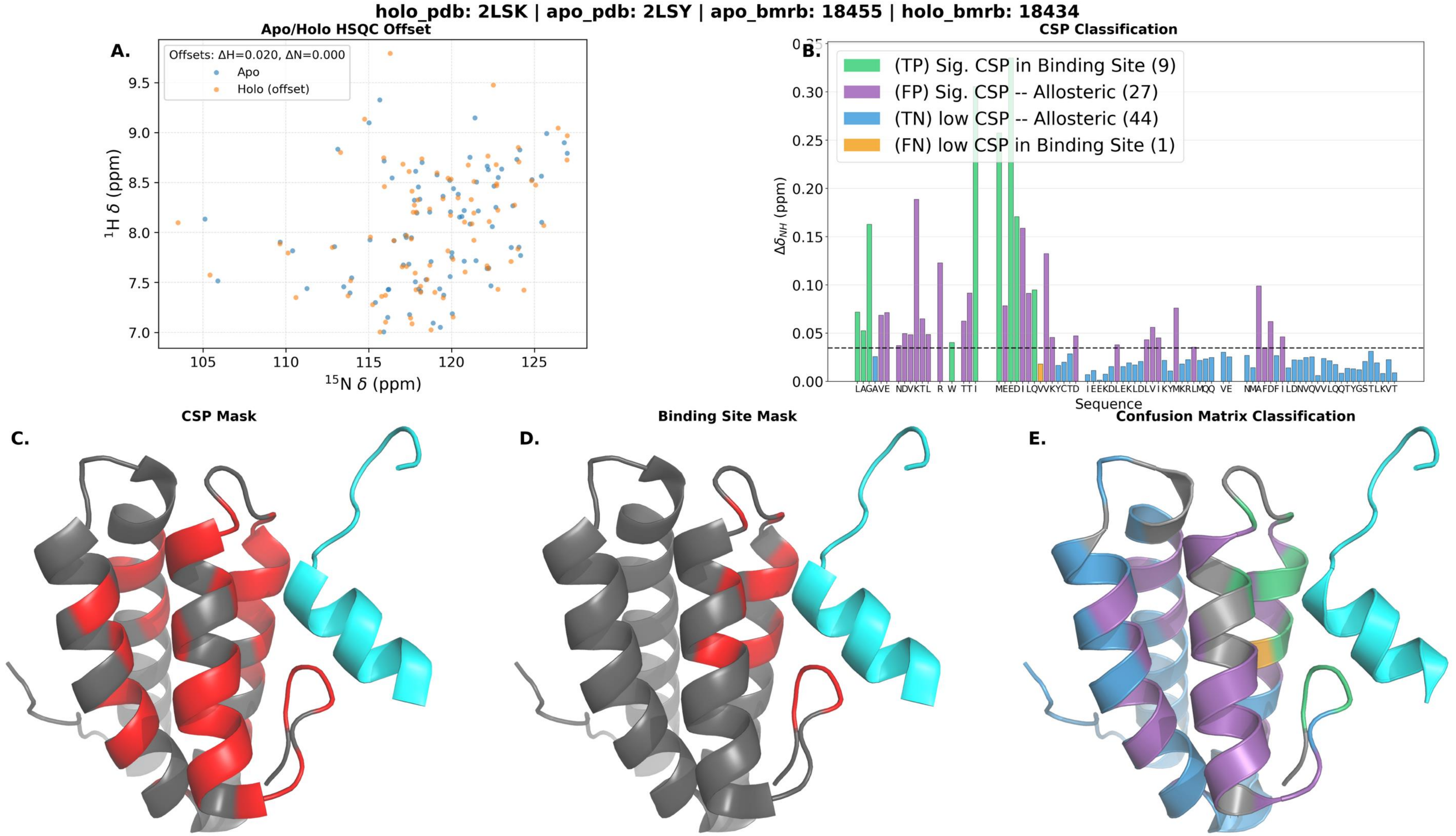

holo_pdb: 2LSK | apo_pdb: 2LSY | apo_bmrb: 18455 | holo_bmrb: 18434
Apo/Holo HSQC Offset
A.
Offsets: ΔH=0.020, ΔN=0.000
Apo
Holo (offset)
¹H δ (ppm)
¹⁵N δ (ppm)
CSP Classification
B.
(TP) Sig. CSP in Binding Site (9)
(FP) Sig. CSP -- Allosteric (27)
(TN) low CSP -- Allosteric (44)
(FN) low CSP in Binding Site (1)
Δδ_NH (ppm)
Sequence
CSP Mask
C.
Binding Site Mask
D.
Confusion Matrix Classification
E.

holo_pdb: 2LSP | apo_pdb: N/A | apo_bmrb: 15057 | holo_bmrb: 18439

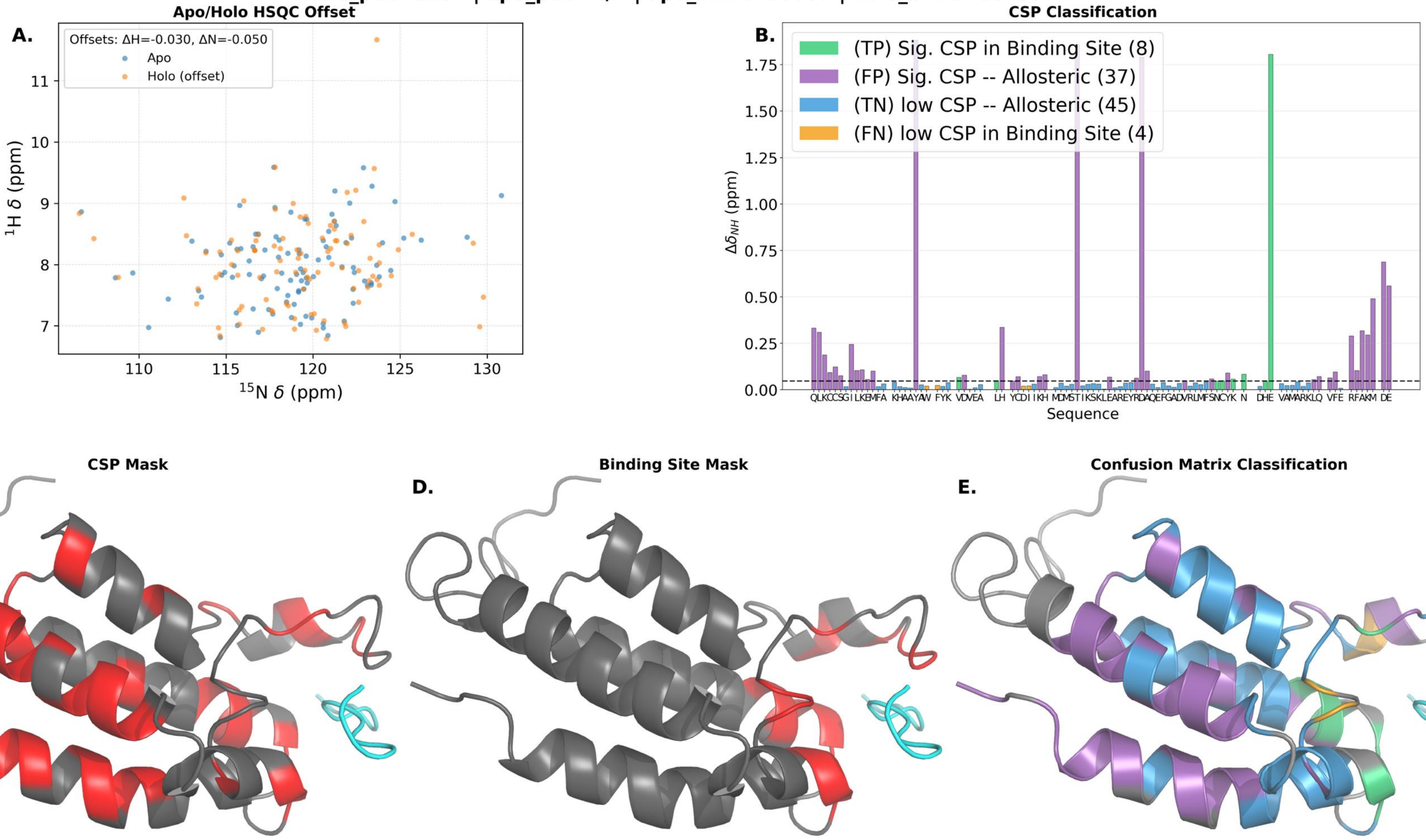

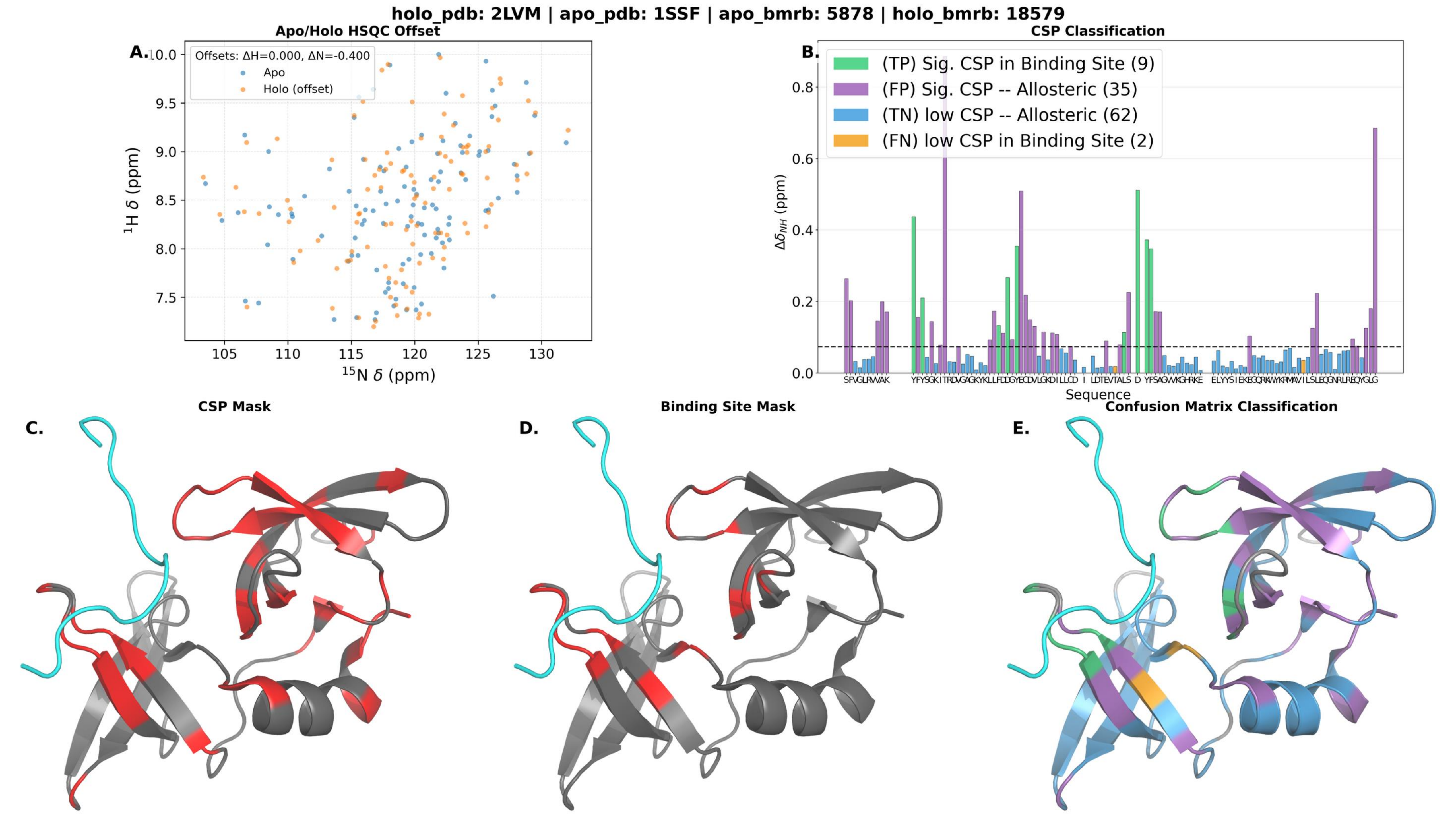
holo_pdb: 2LVM | apo_pdb: 1SSF | apo_bmrb: 5878 | holo_bmrb: 18579
A.
Apo/Holo HSQC Offset
Offsets: ΔH=0.000, ΔN=-0.400
Apo
Holo (offset)
$^{1}$H δ (ppm)
$^{15}$N δ (ppm)
B.
CSP Classification
(TP) Sig. CSP in Binding Site (9)
(FP) Sig. CSP -- Allosteric (35)
(TN) low CSP -- Allosteric (62)
(FN) low CSP in Binding Site (2)
$\Delta\delta_{NH}$ (ppm)
Sequence
C.
CSP Mask
D.
Binding Site Mask
E.
Confusion Matrix Classification

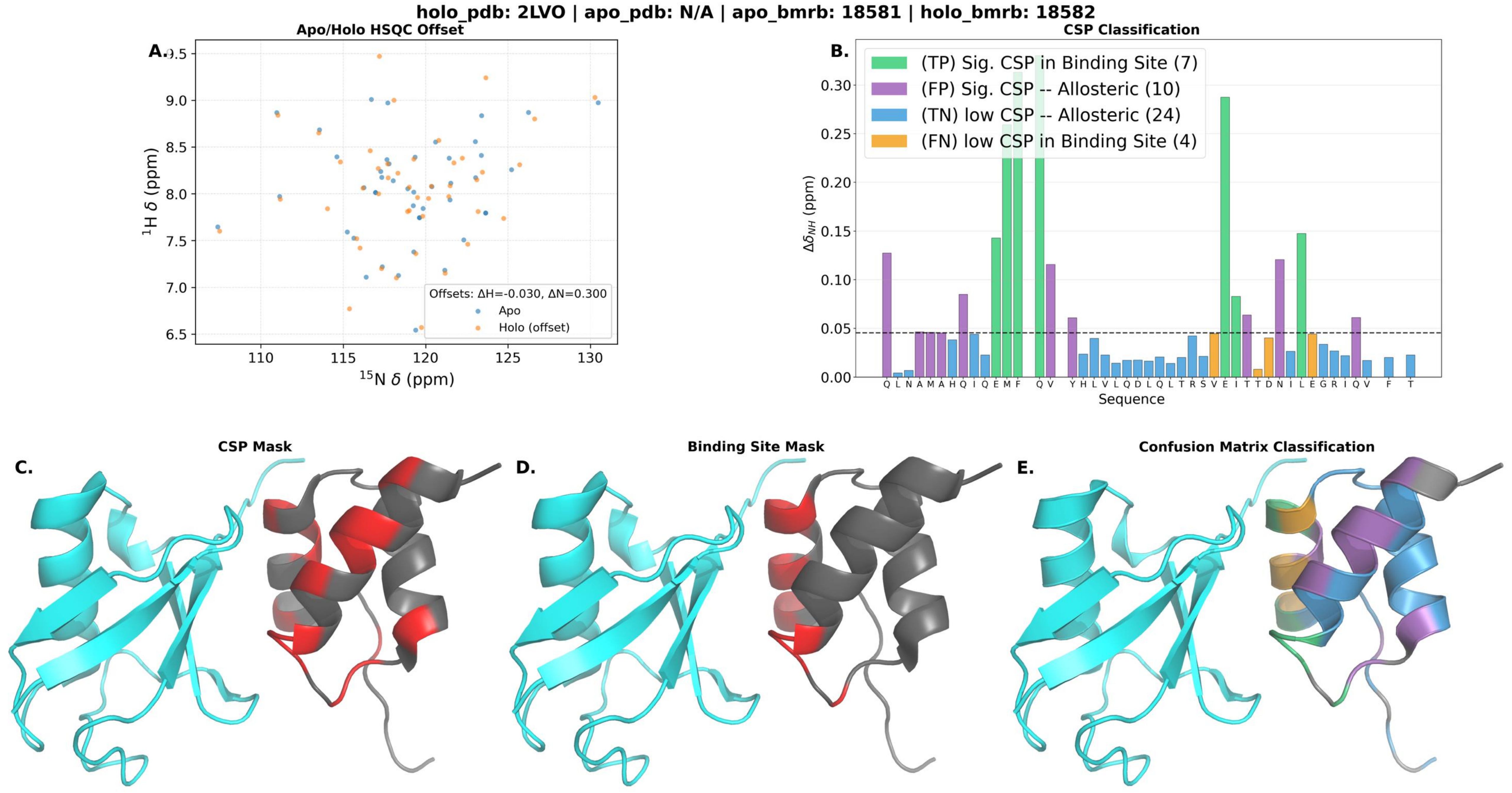

holo_pdb: 2LVO | apo_pdb: N/A | apo_bmrb: 18581 | holo_bmrb: 18582
A.
Apo/Holo HSQC Offset
$^{1}$H δ (ppm)
$^{15}$N δ (ppm)
Offsets: ΔH=-0.030, ΔN=0.300
Apo
Holo (offset)
B.
CSP Classification
(TP) Sig. CSP in Binding Site (7)
(FP) Sig. CSP -- Allosteric (10)
(TN) low CSP -- Allosteric (24)
(FN) low CSP in Binding Site (4)
$\Delta\delta_{NH}$ (ppm)
Sequence
Q L N A M A H Q I Q E M F Q V Y H L V L Q D L Q L T R S V E I T T D N I L E G R I Q V F T
C.
CSP Mask
D.
Binding Site Mask
E.
Confusion Matrix Classification

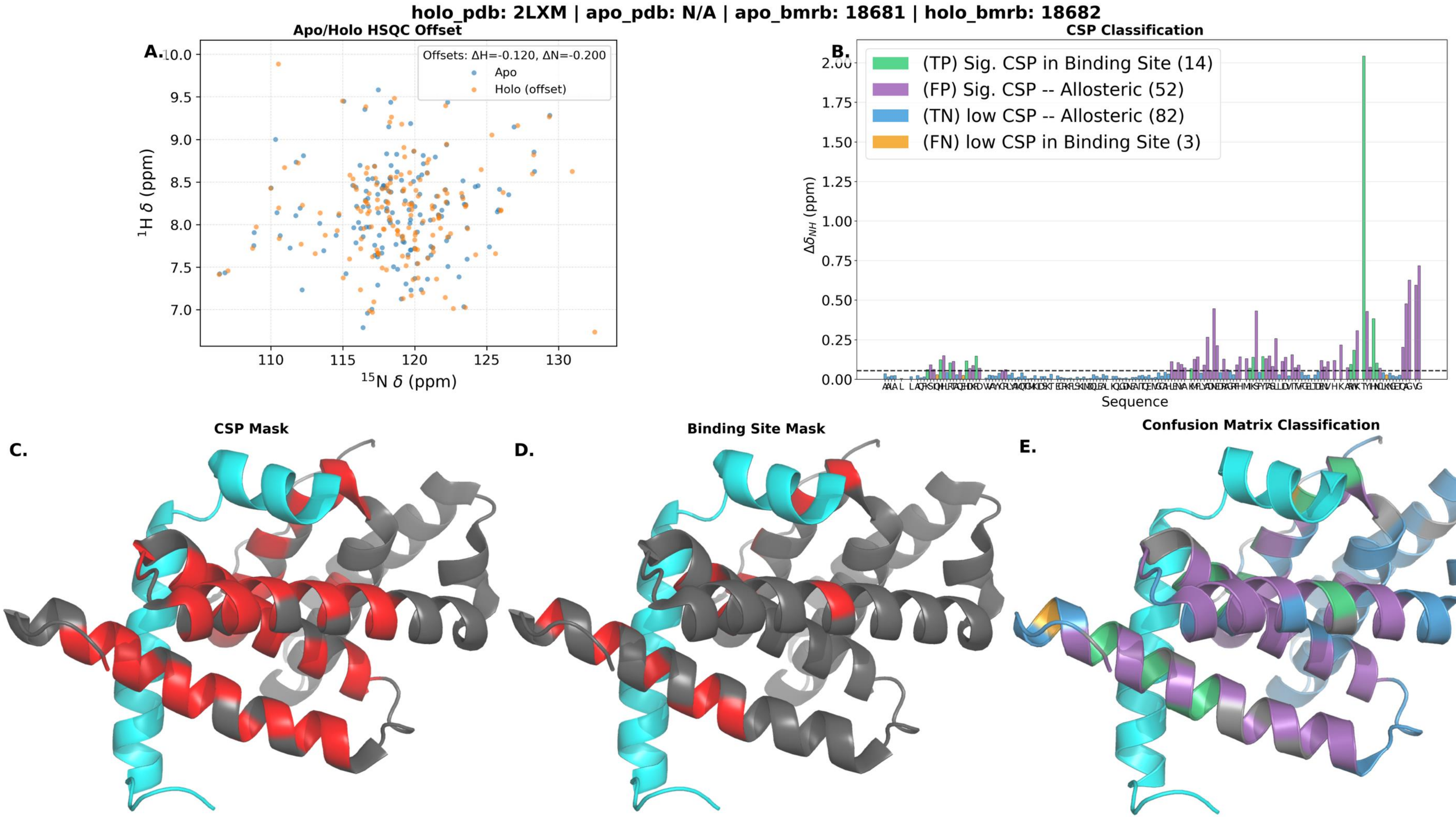

holo_pdb: 2LXM | apo_pdb: N/A | apo_bmrb: 18681 | holo_bmrb: 18682
Apo/Holo HSQC Offset
A.
Offsets: ΔH=-0.120, ΔN=-0.200
Apo
Holo (offset)
$^{1}$H δ (ppm)
$^{15}$N δ (ppm)
10.0
9.5
9.0
8.5
8.0
7.5
7.0
110
115
120
125
130
CSP Classification
B.
(TP) Sig. CSP in Binding Site (14)
(FP) Sig. CSP -- Allosteric (52)
(TN) low CSP -- Allosteric (82)
(FN) low CSP in Binding Site (3)
$\Delta\delta_{NH}$ (ppm)
2.00
1.75
1.50
1.25
1.00
0.75
0.50
0.25
0.00
Sequence
CSP Mask
C.
Binding Site Mask
D.
Confusion Matrix Classification
E.

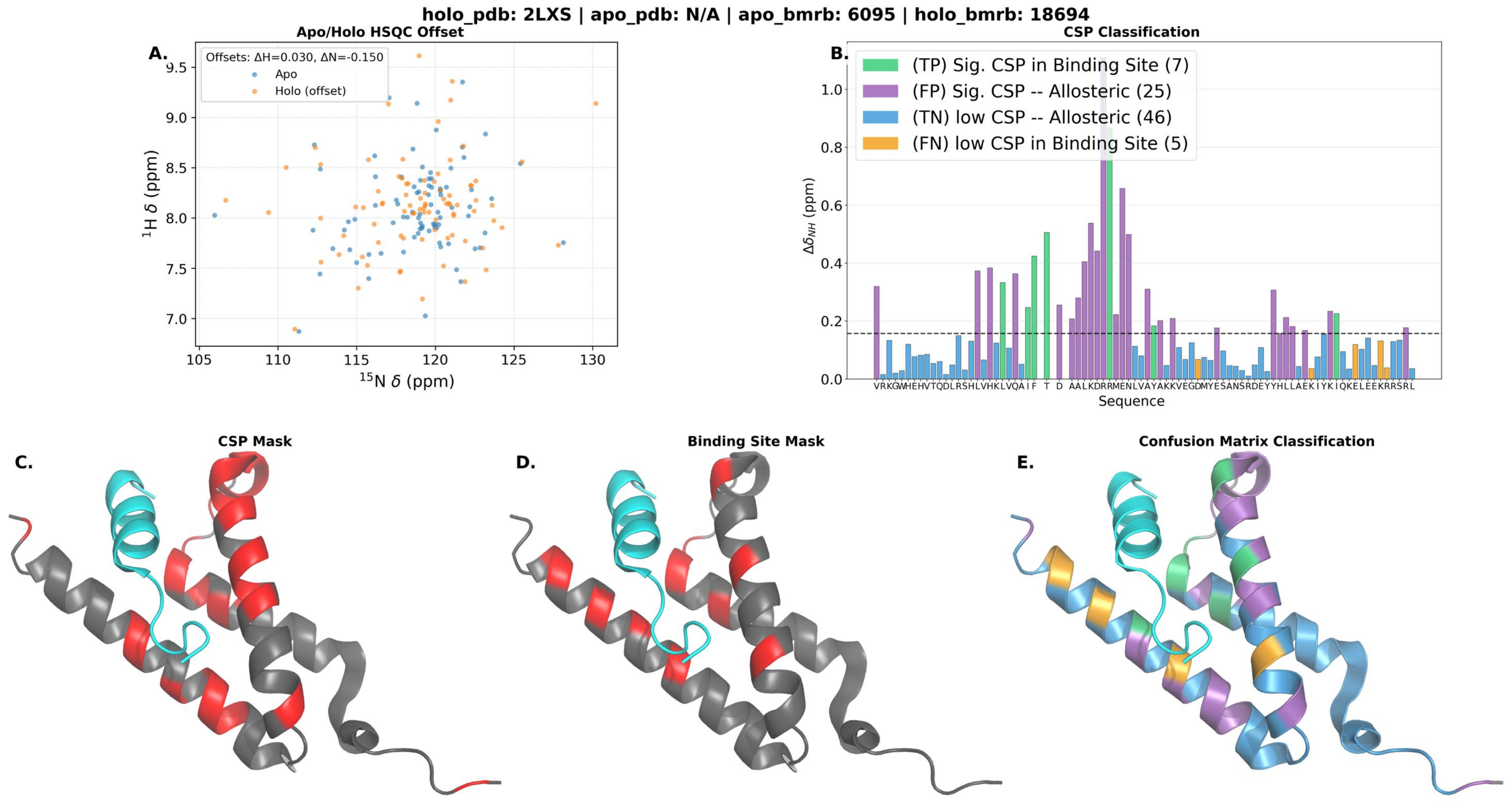

holo_pdb: 2LXS | apo_pdb: N/A | apo_bmrb: 6095 | holo_bmrb: 18694
A.
Apo/Holo HSQC Offset
Offsets: ΔH=0.030, ΔN=-0.150
Apo
Holo (offset)
1H δ (ppm)
15N δ (ppm)
B.
CSP Classification
(TP) Sig. CSP in Binding Site (7)
(FP) Sig. CSP -- Allosteric (25)
(TN) low CSP -- Allosteric (46)
(FN) low CSP in Binding Site (5)
ΔδNH (ppm)
Sequence
C.
CSP Mask
D.
Binding Site Mask
E.
Confusion Matrix Classification

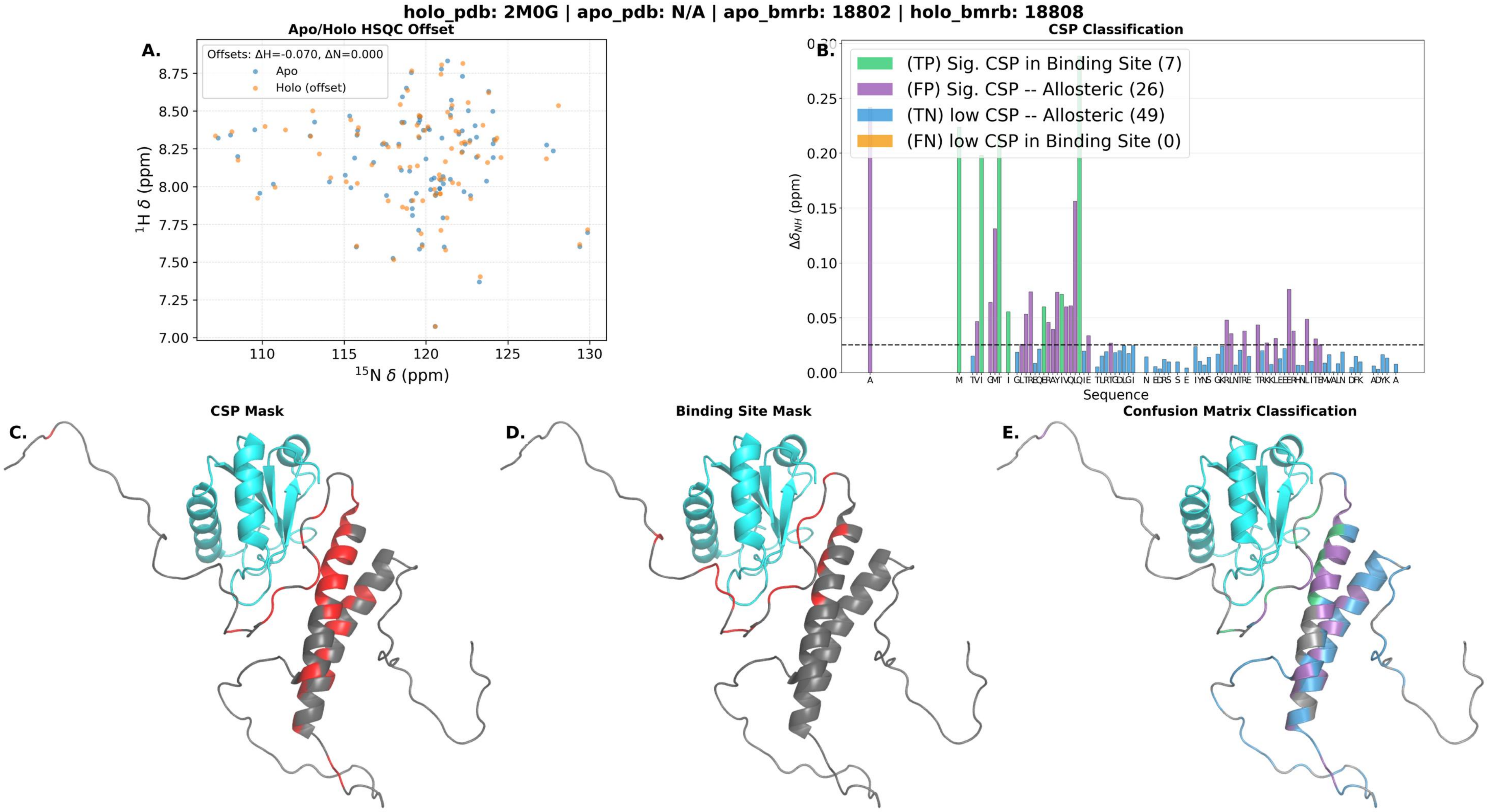

holo_pdb: 2M0G | apo_pdb: N/A | apo_bmrb: 18802 | holo_bmrb: 18808
Apo/Holo HSQC Offset
A.
Offsets: ΔH=-0.070, ΔN=0.000
Apo
Holo (offset)
$^{1}$H δ (ppm)
$^{15}$N δ (ppm)
CSP Classification
B.
(TP) Sig. CSP in Binding Site (7)
(FP) Sig. CSP -- Allosteric (26)
(TN) low CSP -- Allosteric (49)
(FN) low CSP in Binding Site (0)
$\Delta\delta_{NH}$ (ppm)
Sequence
C.
CSP Mask
D.
Binding Site Mask
E.
Confusion Matrix Classification

holo_pdb: 2M0G | apo_pdb: N/A | apo_bmrb: 19034 | holo_bmrb: 18808

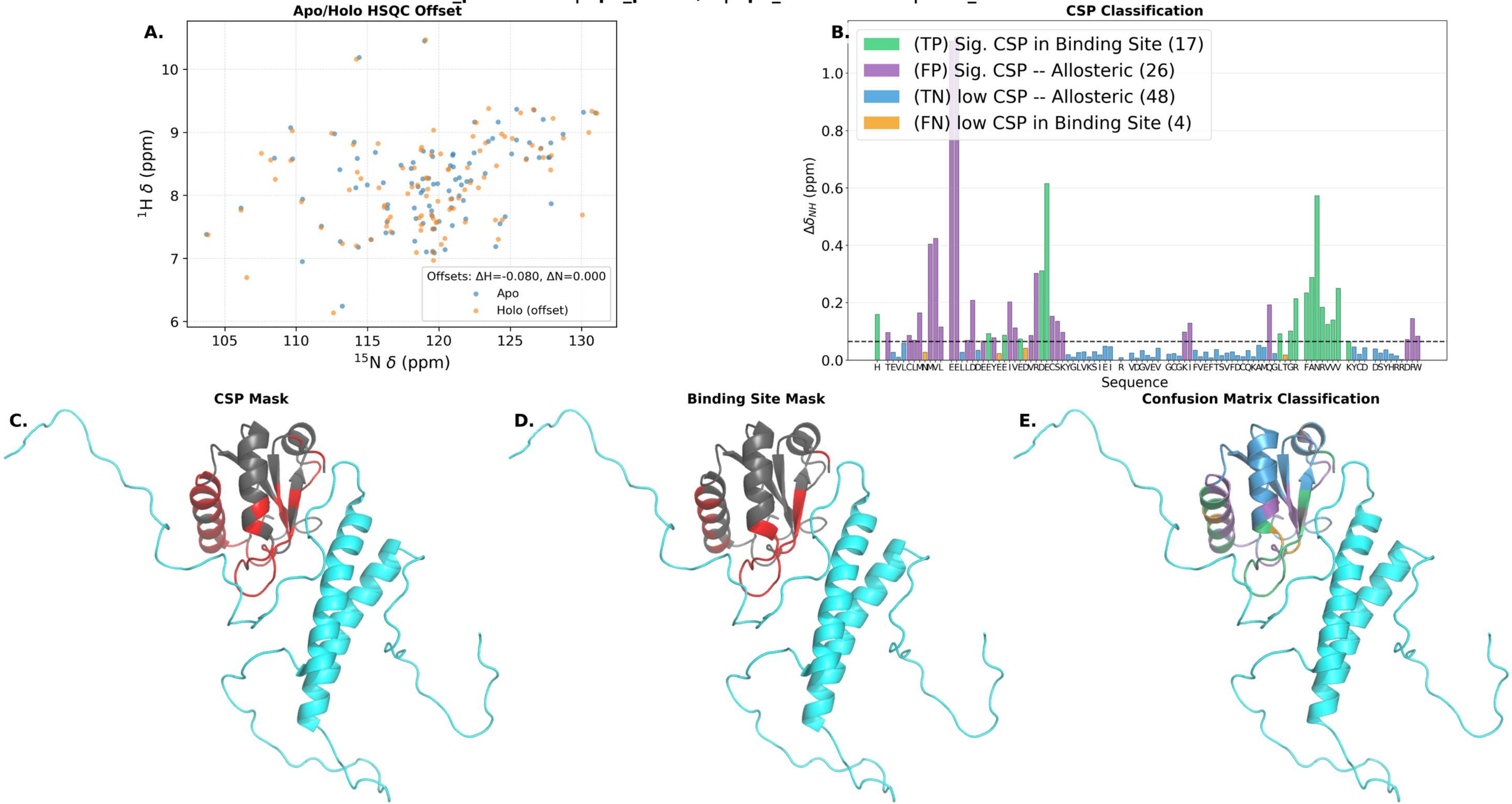

**holo_pdb: 2M0J | apo_pdb: N/A | apo_bmrb: 51289 | holo_bmrb: 17771**

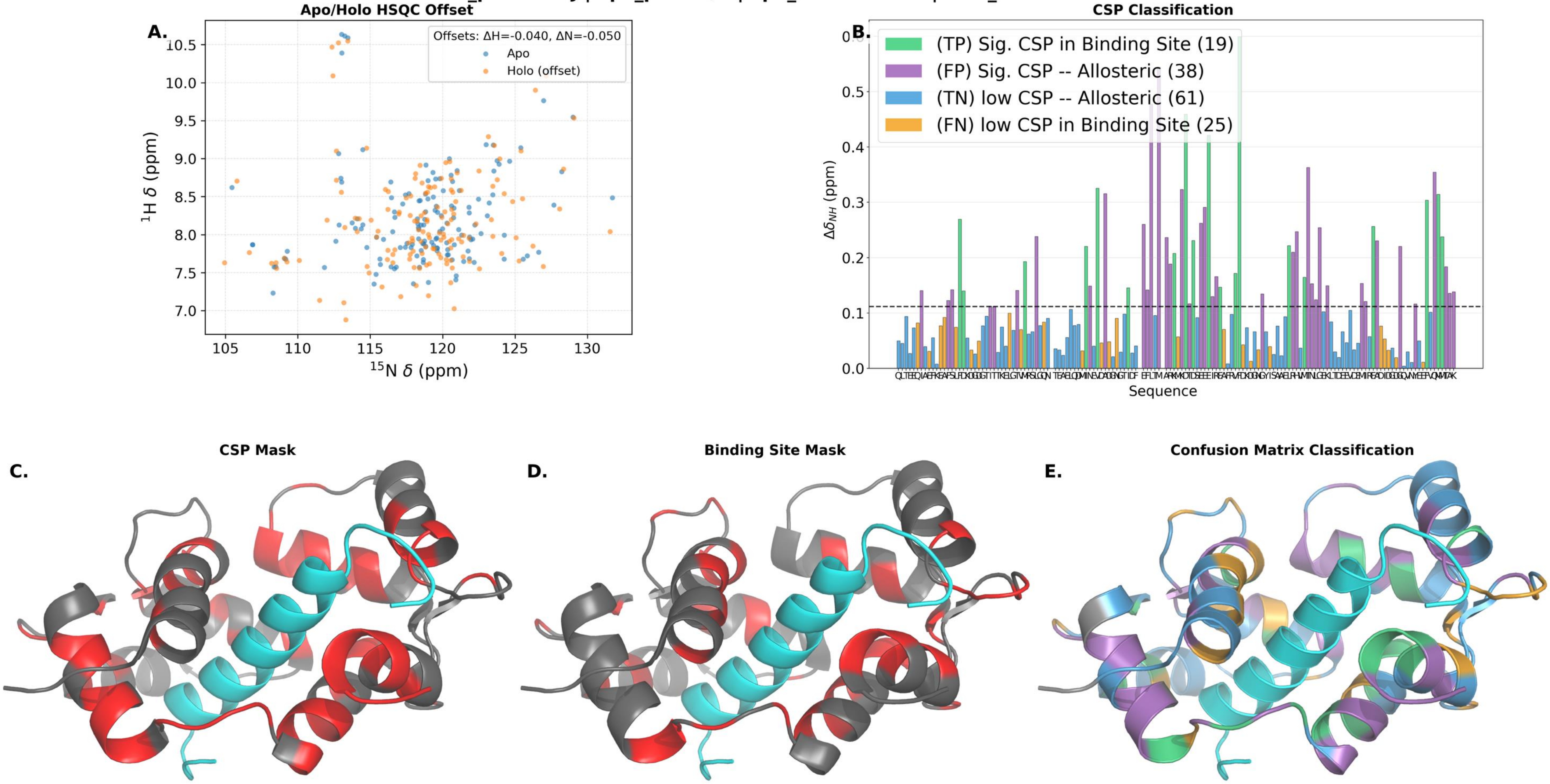

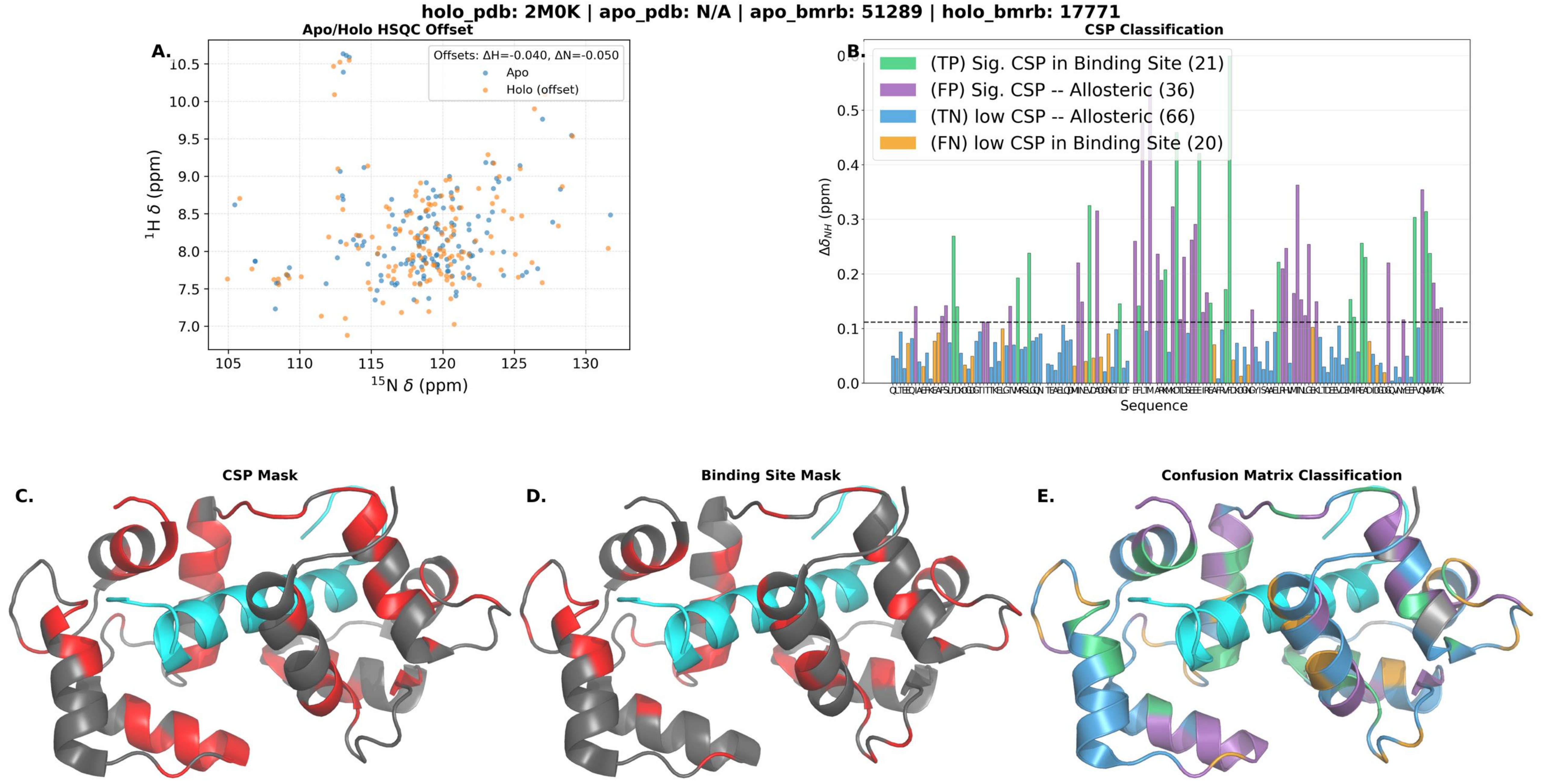

holo_pdb: 2M0K | apo_pdb: N/A | apo_bmrb: 51289 | holo_bmrb: 17771
A.
Apo/Holo HSQC Offset
Offsets: ΔH=-0.040, ΔN=-0.050
Apo
Holo (offset)
1H δ (ppm)
15N δ (ppm)
B.
CSP Classification
(TP) Sig. CSP in Binding Site (21)
(FP) Sig. CSP -- Allosteric (36)
(TN) low CSP -- Allosteric (66)
(FN) low CSP in Binding Site (20)
ΔδNH (ppm)
Sequence
C.
CSP Mask
D.
Binding Site Mask
E.
Confusion Matrix Classification

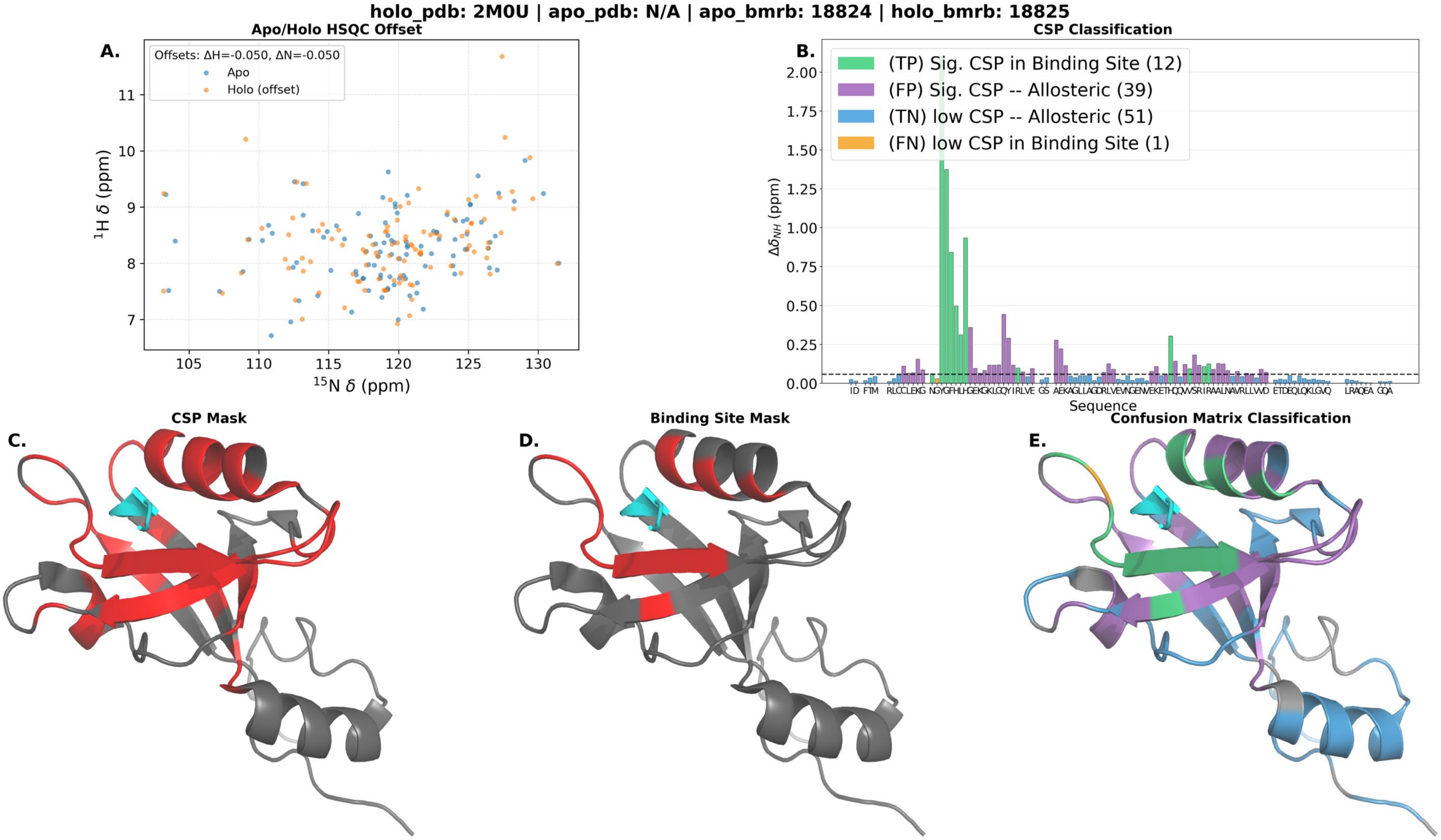
holo_pdb: 2M0U | apo_pdb: N/A | apo_bmrb: 18824 | holo_bmrb: 18825
A.
Apo/Holo HSQC Offset
Offsets: ΔH=-0.050, ΔN=-0.050
Apo
Holo (offset)
$^{1}$H δ (ppm)
$^{15}$N δ (ppm)
B.
CSP Classification
(TP) Sig. CSP in Binding Site (12)
(FP) Sig. CSP -- Allosteric (39)
(TN) low CSP -- Allosteric (51)
(FN) low CSP in Binding Site (1)
$\Delta\delta_{NH}$ (ppm)
Sequence
C.
CSP Mask
D.
Binding Site Mask
E.
Confusion Matrix Classification

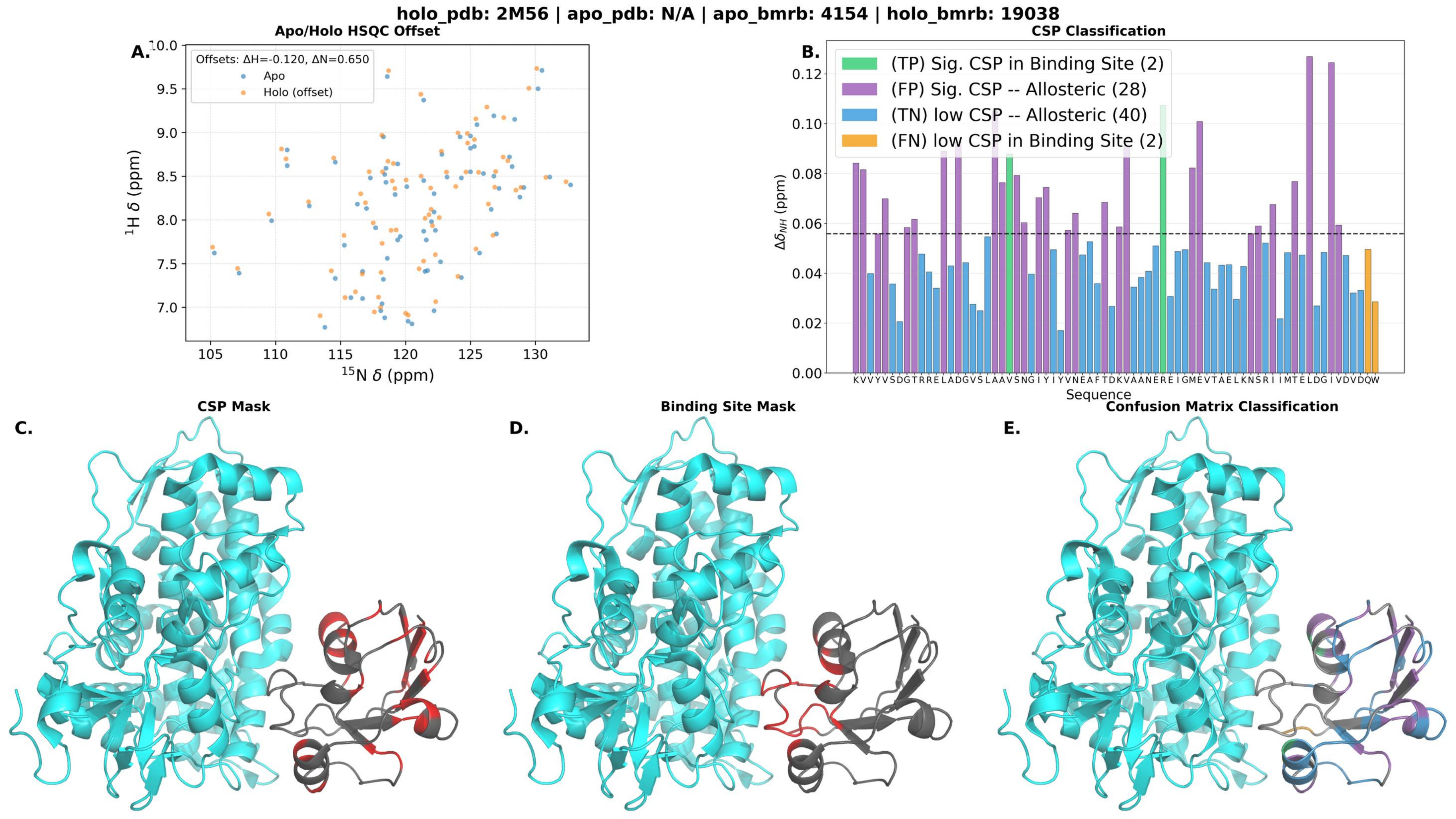
holo_pdb: 2M56 | apo_pdb: N/A | apo_bmrb: 4154 | holo_bmrb: 19038
A.
Apo/Holo HSQC Offset
Offsets: ΔH=-0.120, ΔN=0.650
Apo
Holo (offset)
1H δ (ppm)
15N δ (ppm)
B.
CSP Classification
(TP) Sig. CSP in Binding Site (2)
(FP) Sig. CSP -- Allosteric (28)
(TN) low CSP -- Allosteric (40)
(FN) low CSP in Binding Site (2)
ΔδNH (ppm)
Sequence
KVVYVSDGTRRELADGVSLAAVSNGIYIYVNEAFTDKVAANERE IGMEVTAELKNSRI IMTELDGIVDVDQW
C.
CSP Mask
D.
Binding Site Mask
E.
Confusion Matrix Classification

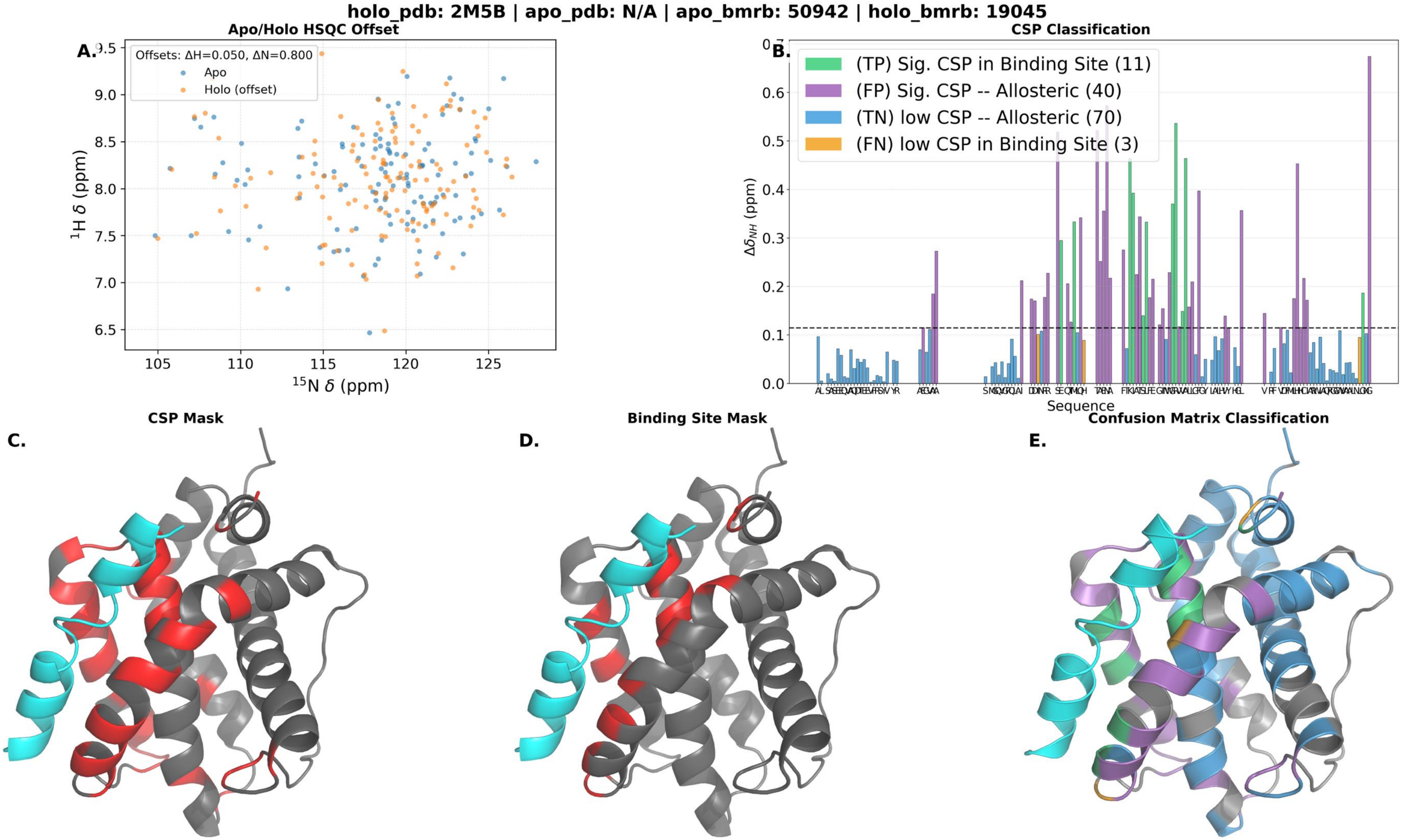

holo_pdb: 2M5B | apo_pdb: N/A | apo_bmrb: 50942 | holo_bmrb: 19045
A.
Apo/Holo HSQC Offset
Offsets: ΔH=0.050, ΔN=0.800
Apo
Holo (offset)
¹H δ (ppm)
¹⁵N δ (ppm)
B.
CSP Classification
(TP) Sig. CSP in Binding Site (11)
(FP) Sig. CSP -- Allosteric (40)
(TN) low CSP -- Allosteric (70)
(FN) low CSP in Binding Site (3)
Δδ_NH (ppm)
Sequence
C.
CSP Mask
D.
Binding Site Mask
E.
Confusion Matrix Classification

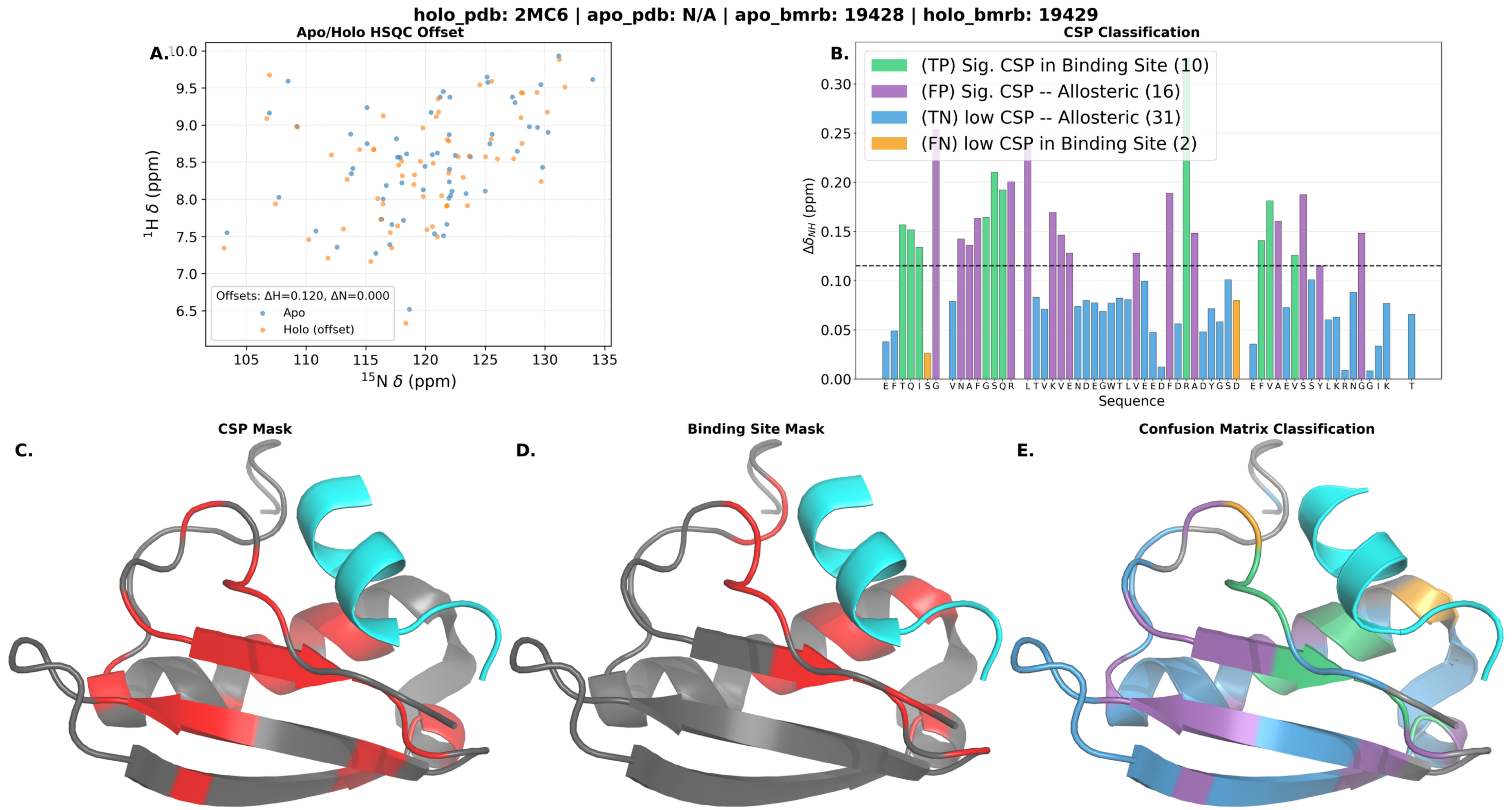
holo_pdb: 2MC6 | apo_pdb: N/A | apo_bmrb: 19428 | holo_bmrb: 19429
A.
Apo/Holo HSQC Offset
$^{1}$H δ (ppm)
$^{15}$N δ (ppm)
Offsets: ΔH=0.120, ΔN=0.000
Apo
Holo (offset)
B.
CSP Classification
(TP) Sig. CSP in Binding Site (10)
(FP) Sig. CSP -- Allosteric (16)
(TN) low CSP -- Allosteric (31)
(FN) low CSP in Binding Site (2)
$\Delta\delta_{NH}$ (ppm)
Sequence
E F T Q I S G V N A F G S Q R L T V K V E N D E G W T L V E E D F D R A D Y G S D E F V A E V S S Y L K R N G G I K T
C.
CSP Mask
D.
Binding Site Mask
E.
Confusion Matrix Classification

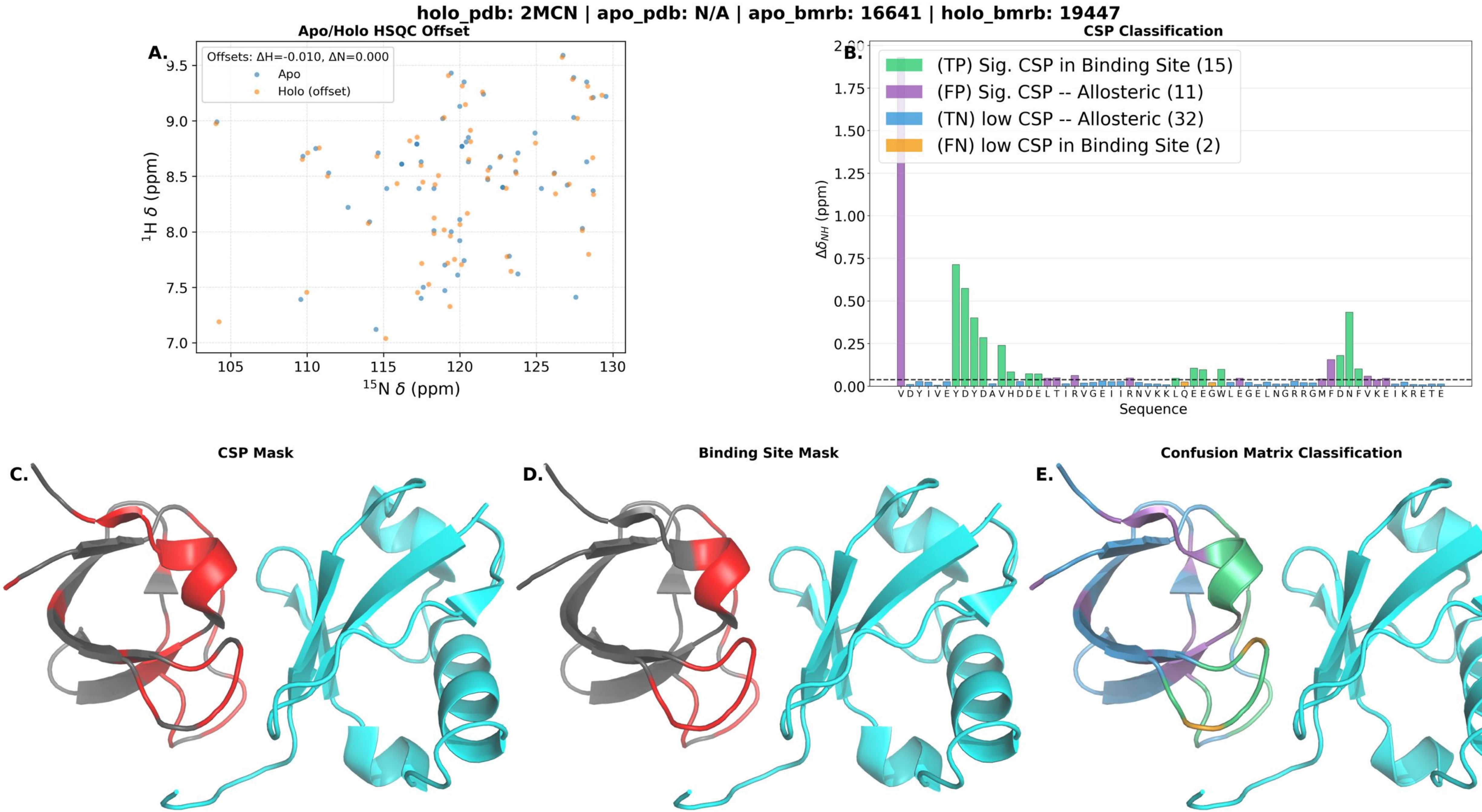
holo_pdb: 2MCN | apo_pdb: N/A | apo_bmrb: 16641 | holo_bmrb: 19447
A.
Apo/Holo HSQC Offset
Offsets: ΔH=-0.010, ΔN=0.000
Apo
Holo (offset)
1H δ (ppm)
15N δ (ppm)
B.
CSP Classification
(TP) Sig. CSP in Binding Site (15)
(FP) Sig. CSP -- Allosteric (11)
(TN) low CSP -- Allosteric (32)
(FN) low CSP in Binding Site (2)
Δδ_NH (ppm)
Sequence
V D Y I V E Y D Y D A V H D D E L T I R V G E I I R N V K K L Q E E G W L E G E L N G R R G M F D N F V K E I K R E T E
C.
CSP Mask
D.
Binding Site Mask
E.
Confusion Matrix Classification

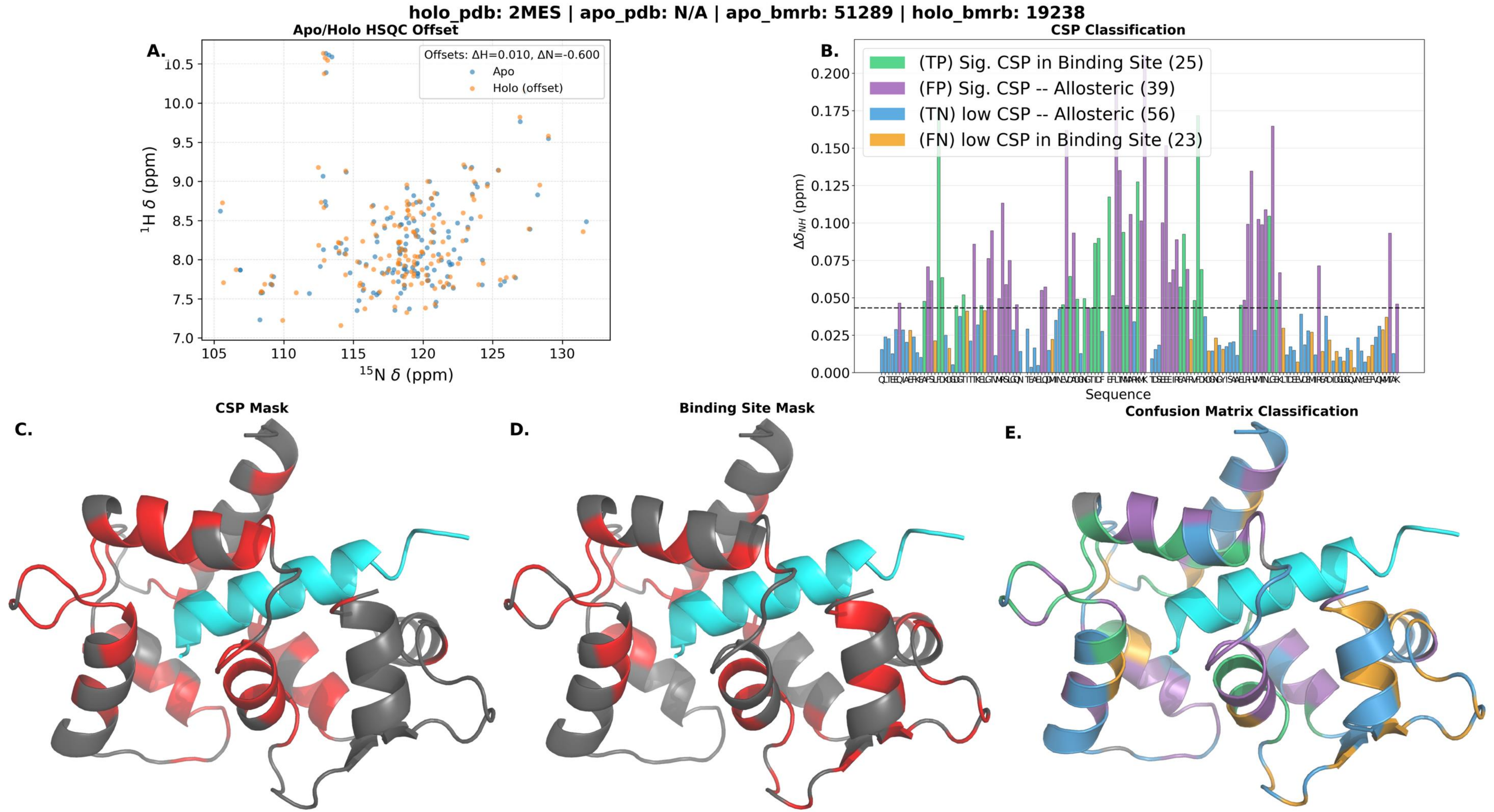
holo_pdb: 2MES | apo_pdb: N/A | apo_bmrb: 51289 | holo_bmrb: 19238
Apo/Holo HSQC Offset
A.
Offsets: ΔH=0.010, ΔN=-0.600
Apo
Holo (offset)
1H δ (ppm)
15N δ (ppm)
CSP Classification
B.
(TP) Sig. CSP in Binding Site (25)
(FP) Sig. CSP -- Allosteric (39)
(TN) low CSP -- Allosteric (56)
(FN) low CSP in Binding Site (23)
Δδ_NH (ppm)
Sequence
CSP Mask
C.
Binding Site Mask
D.
Confusion Matrix Classification
E.

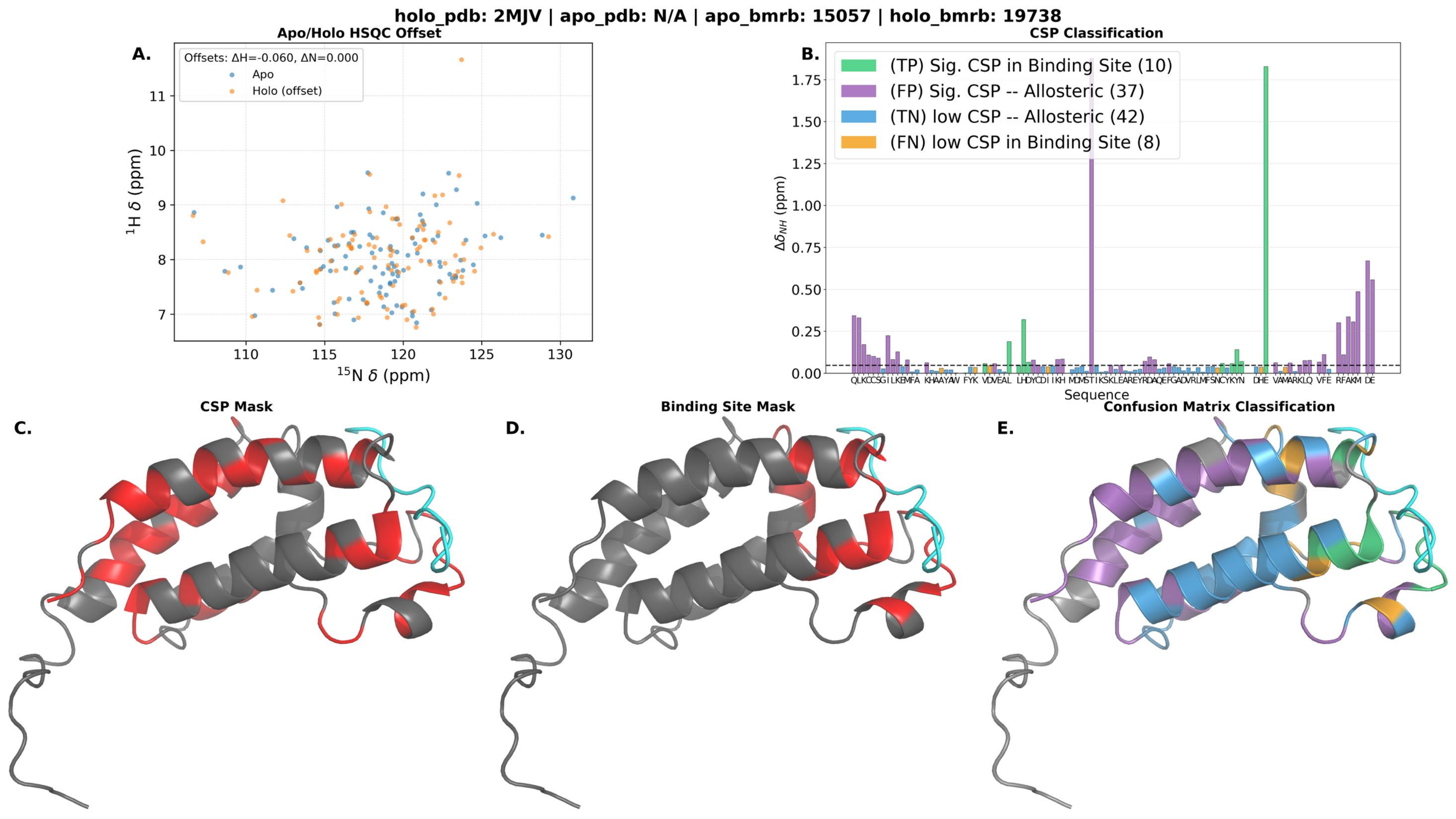

holo_pdb: 2MJV | apo_pdb: N/A | apo_bmrb: 15057 | holo_bmrb: 19738
A.
Apo/Holo HSQC Offset
Offsets: ΔH=-0.060, ΔN=0.000
Apo
Holo (offset)
$^{1}$H δ (ppm)
$^{15}$N δ (ppm)
B.
CSP Classification
(TP) Sig. CSP in Binding Site (10)
(FP) Sig. CSP -- Allosteric (37)
(TN) low CSP -- Allosteric (42)
(FN) low CSP in Binding Site (8)
$\Delta\delta_{NH}$ (ppm)
Sequence
C.
CSP Mask
D.
Binding Site Mask
E.
Confusion Matrix Classification

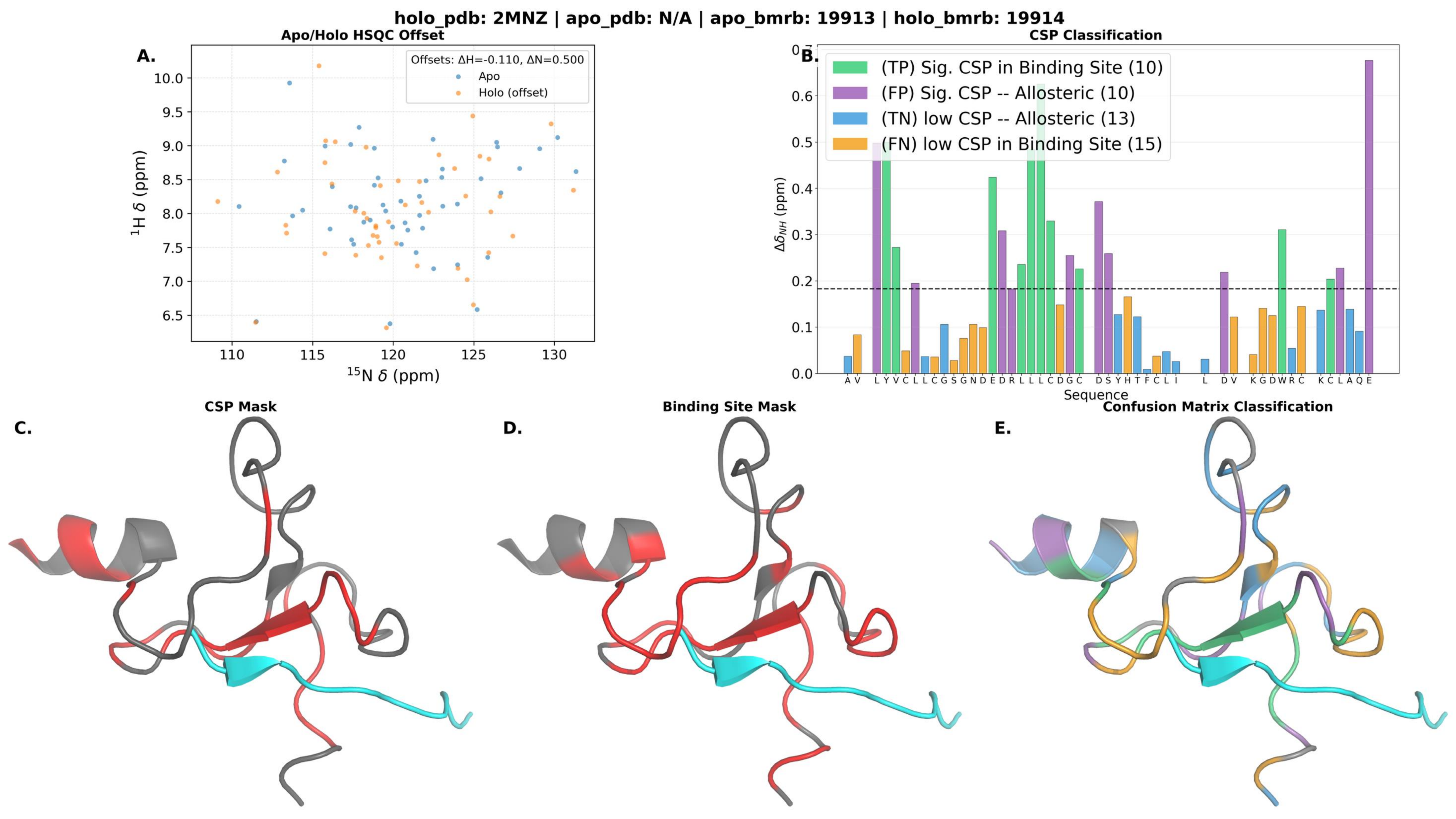

holo_pdb: 2MNZ | apo_pdb: N/A | apo_bmrb: 19913 | holo_bmrb: 19914
Apo/Holo HSQC Offset
A.
Offsets: ΔH=-0.110, ΔN=0.500
Apo
Holo (offset)
$^{1}$H $\delta$ (ppm)
$^{15}$N $\delta$ (ppm)
CSP Classification
B.
(TP) Sig. CSP in Binding Site (10)
(FP) Sig. CSP -- Allosteric (10)
(TN) low CSP -- Allosteric (13)
(FN) low CSP in Binding Site (15)
$\Delta\delta_{NH}$ (ppm)
Sequence
CSP Mask
C.
Binding Site Mask
D.
Confusion Matrix Classification
E.

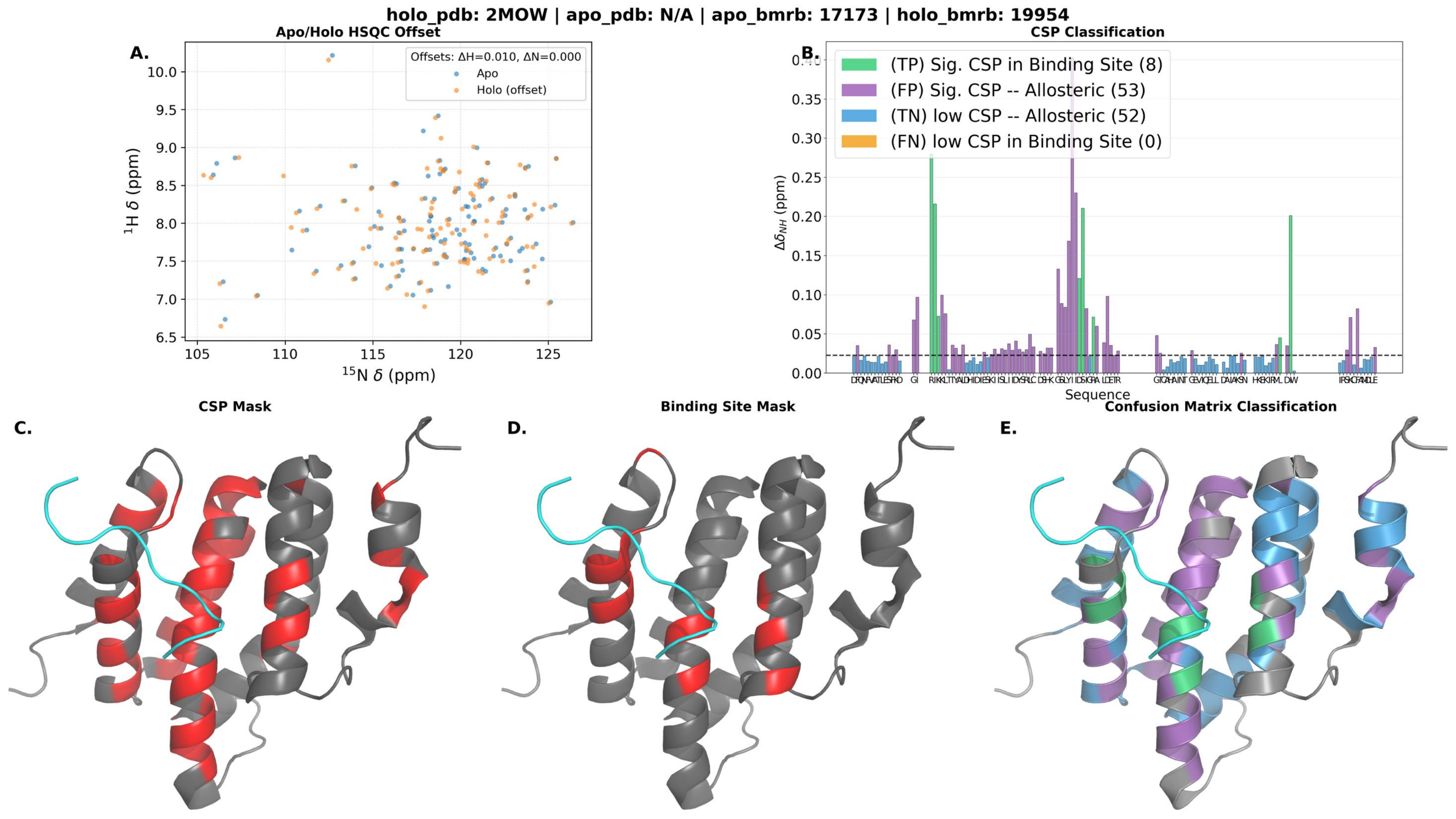
holo_pdb: 2MOW | apo_pdb: N/A | apo_bmrb: 17173 | holo_bmrb: 19954
A.
Apo/Holo HSQC Offset
Offsets: ΔH=0.010, ΔN=0.000
Apo
Holo (offset)
1H δ (ppm)
15N δ (ppm)
B.
CSP Classification
(TP) Sig. CSP in Binding Site (8)
(FP) Sig. CSP -- Allosteric (53)
(TN) low CSP -- Allosteric (52)
(FN) low CSP in Binding Site (0)
Δδ_NH (ppm)
Sequence
C.
CSP Mask
D.
Binding Site Mask
E.
Confusion Matrix Classification

holo_pdb: 2MRE | apo_pdb: N/A | apo_bmrb: 25071 | holo_bmrb: 25070

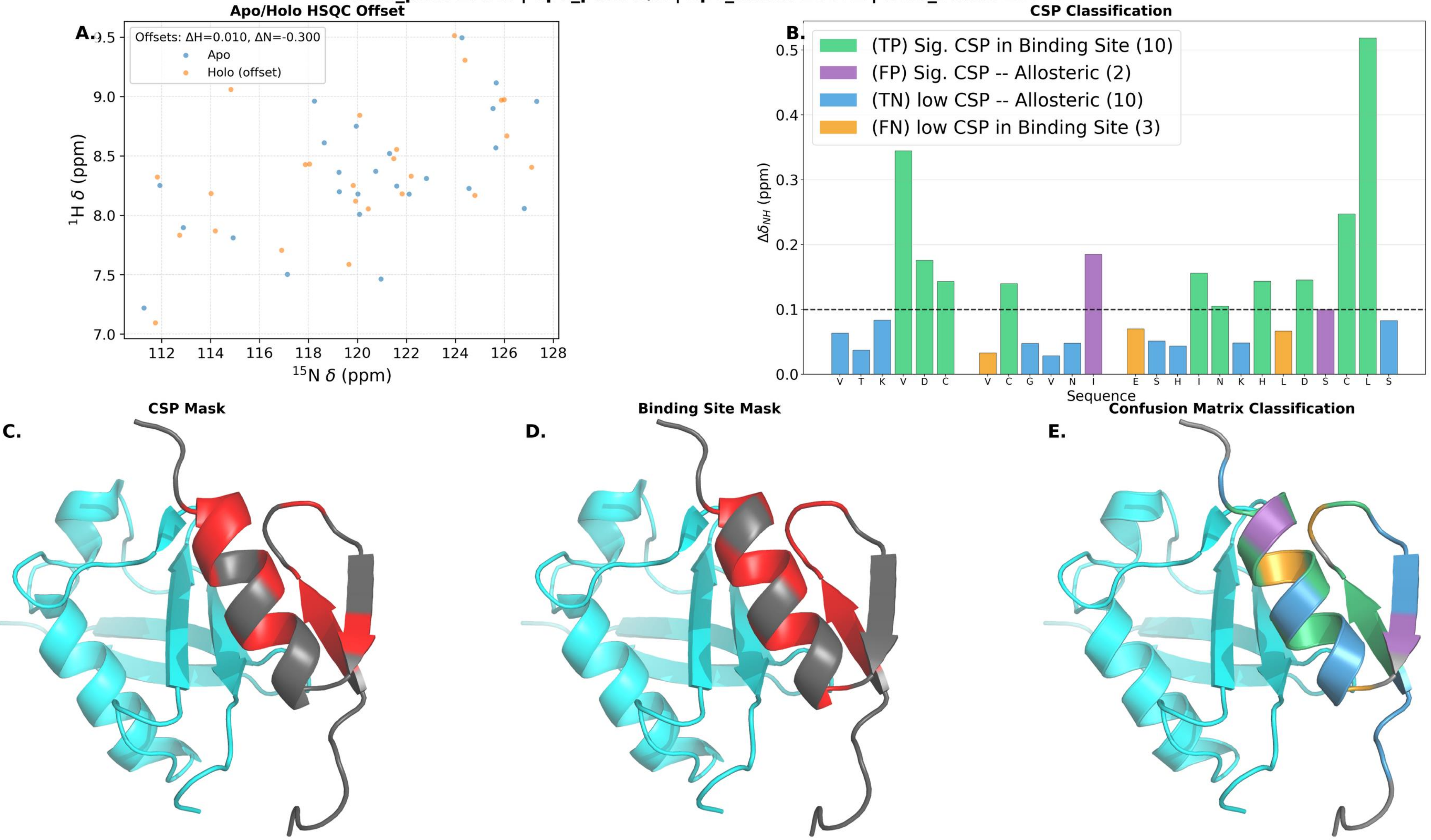

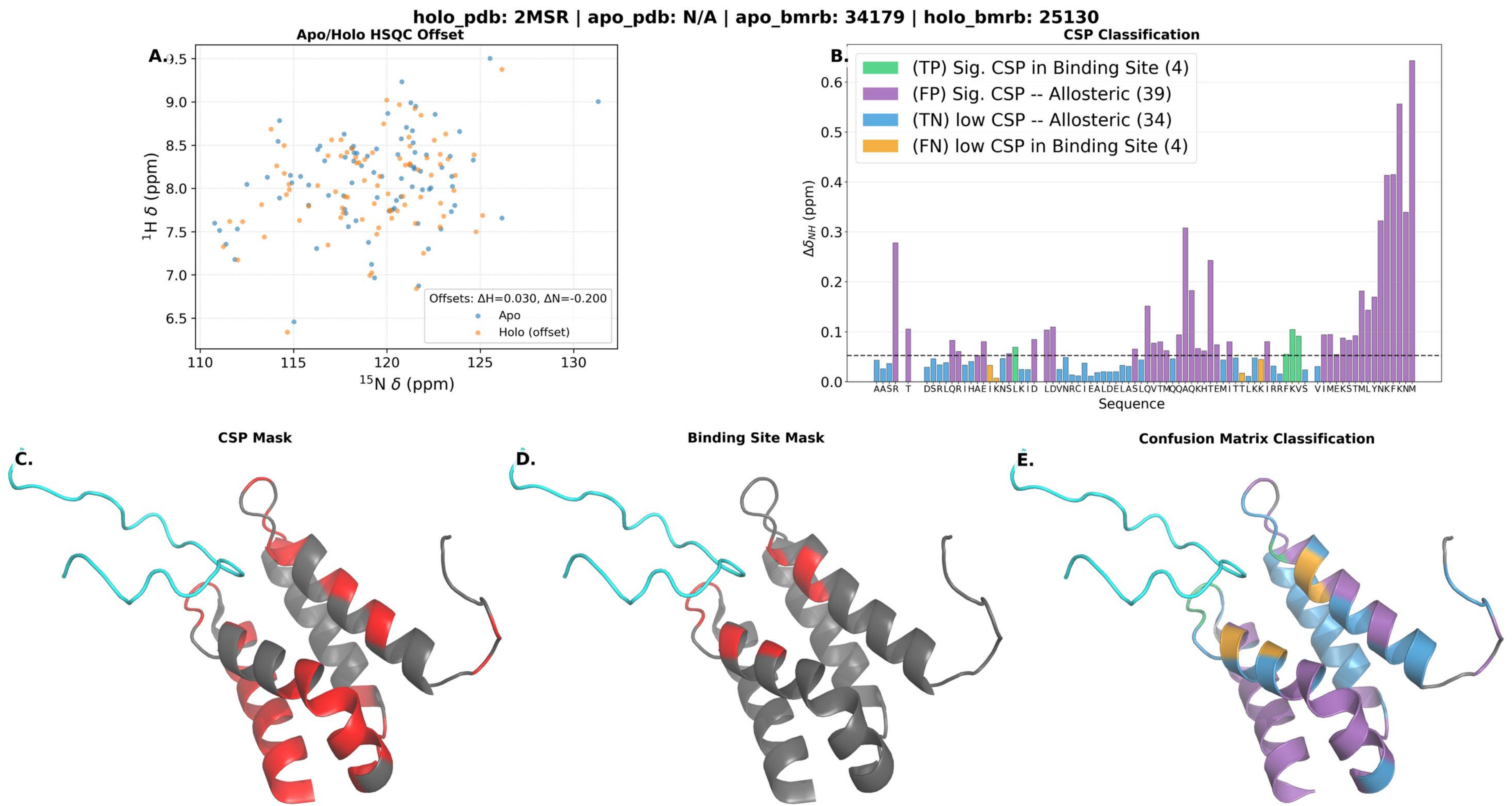
holo_pdb: 2MSR | apo_pdb: N/A | apo_bmrb: 34179 | holo_bmrb: 25130
A.
Apo/Holo HSQC Offset
Offsets: ΔH=0.030, ΔN=-0.200
Apo
Holo (offset)
$^{1}$H δ (ppm)
$^{15}$N δ (ppm)
B.
CSP Classification
(TP) Sig. CSP in Binding Site (4)
(FP) Sig. CSP -- Allosteric (39)
(TN) low CSP -- Allosteric (34)
(FN) low CSP in Binding Site (4)
$\Delta\delta_{NH}$ (ppm)
Sequence
C.
CSP Mask
D.
Binding Site Mask
E.
Confusion Matrix Classification

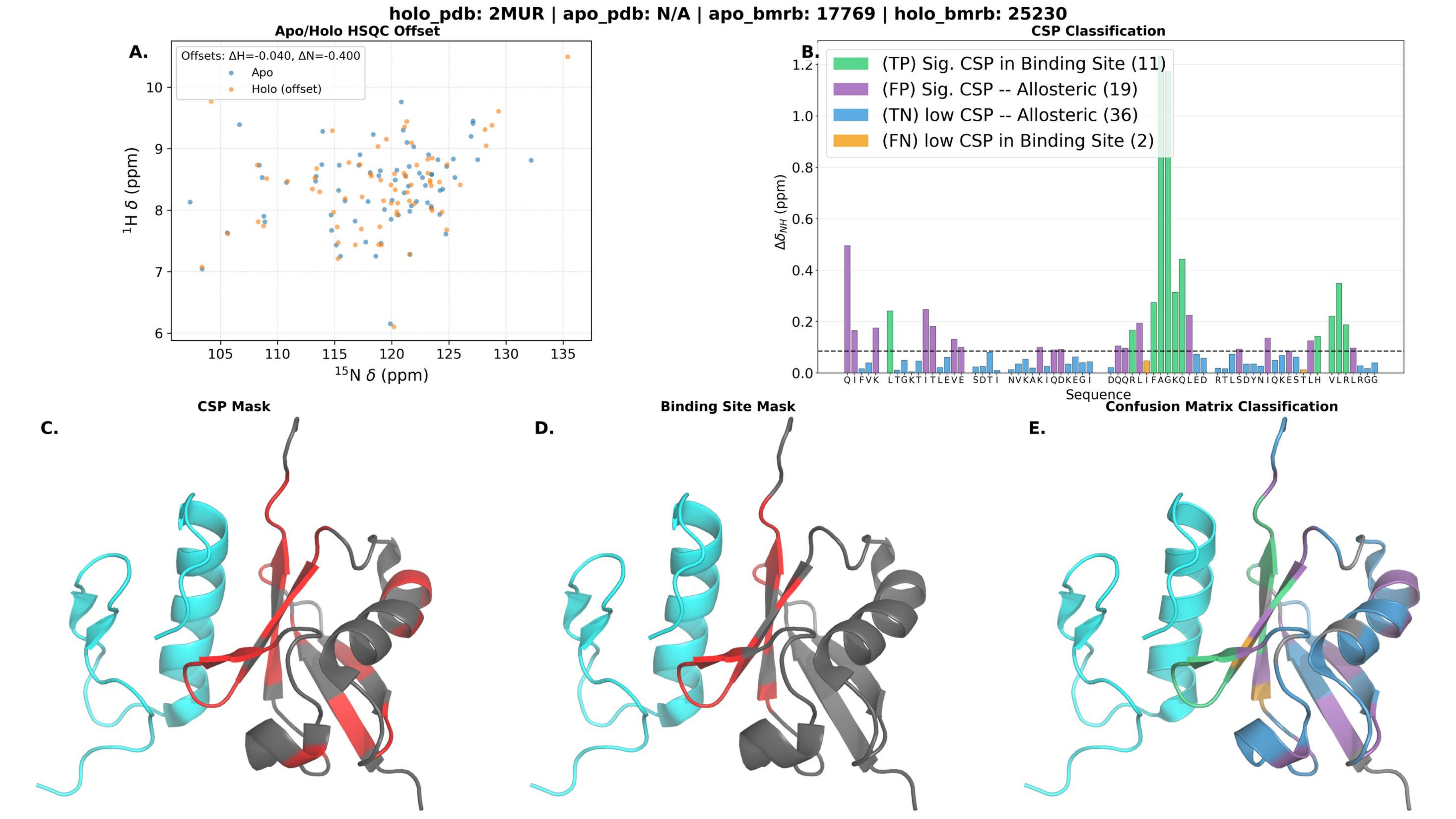
holo_pdb: 2MUR | apo_pdb: N/A | apo_bmrb: 17769 | holo_bmrb: 25230
A.
Apo/Holo HSQC Offset
Offsets: ΔH=-0.040, ΔN=-0.400
Apo
Holo (offset)
¹H δ (ppm)
¹⁵N δ (ppm)
B.
CSP Classification
(TP) Sig. CSP in Binding Site (11)
(FP) Sig. CSP -- Allosteric (19)
(TN) low CSP -- Allosteric (36)
(FN) low CSP in Binding Site (2)
Δδ_NH (ppm)
Sequence
C.
CSP Mask
D.
Binding Site Mask
E.
Confusion Matrix Classification

holo_pdb: 2MUR | apo_pdb: N/A | apo_bmrb: 25229 | holo_bmrb: 25230

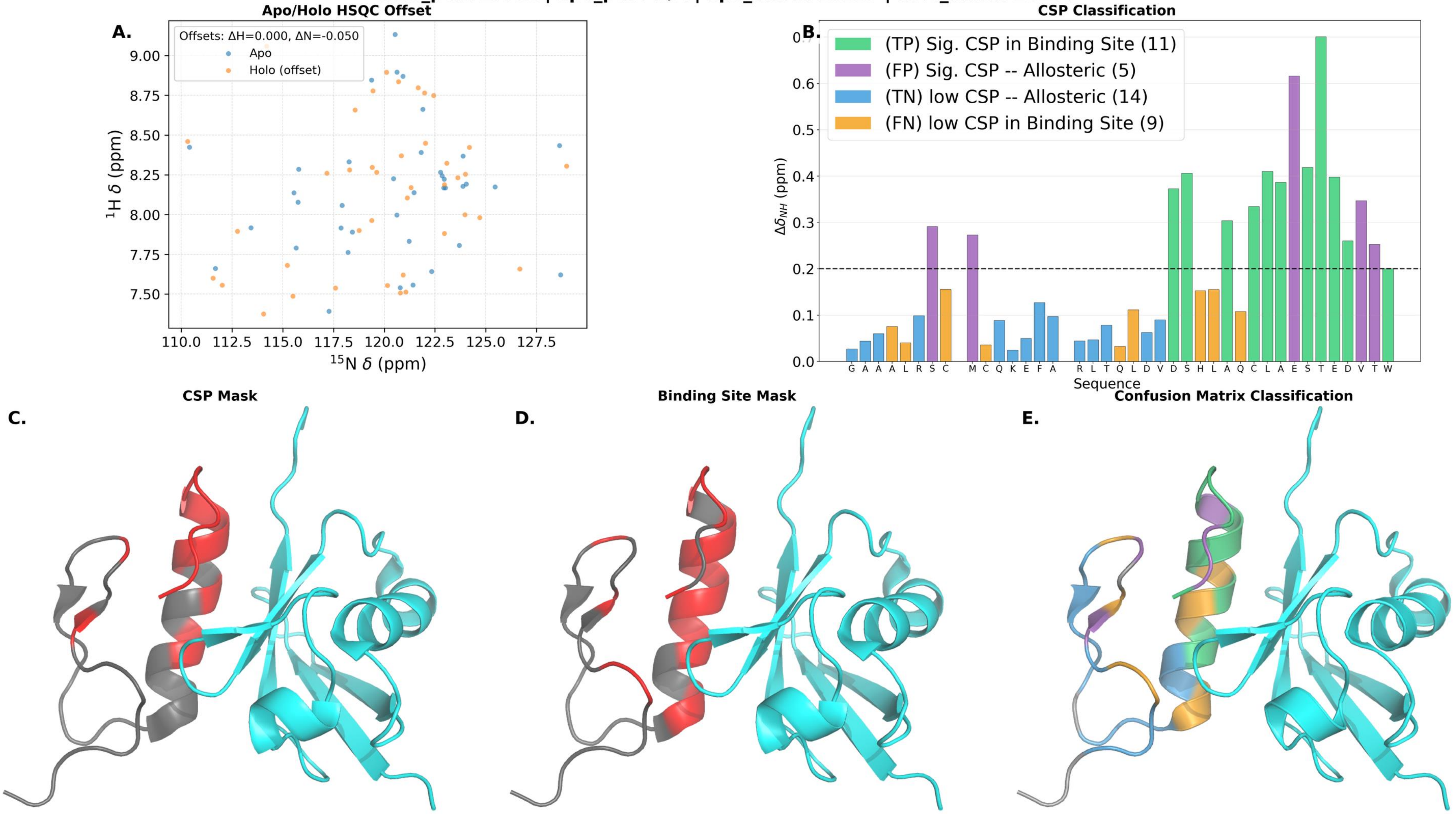

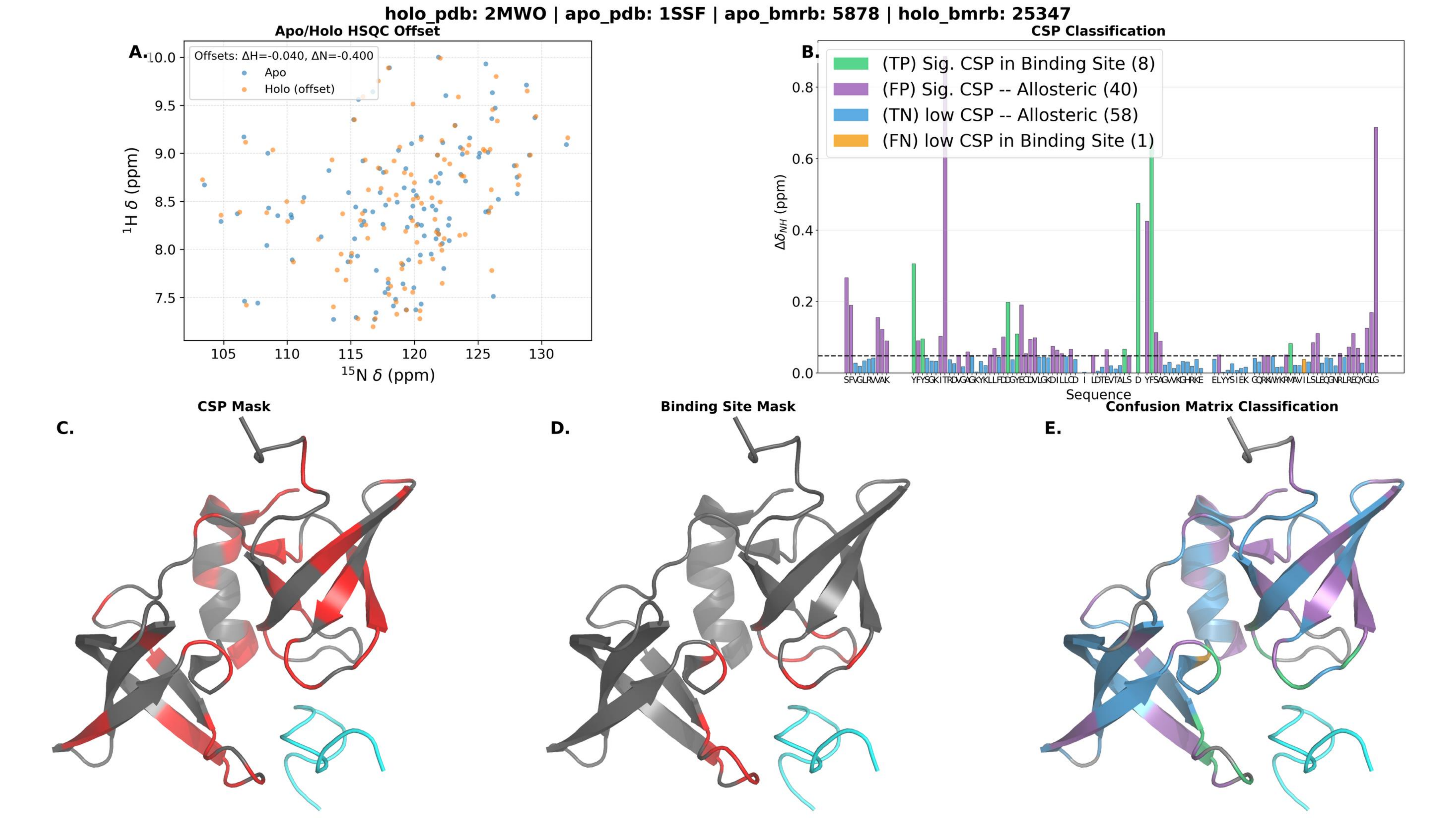
holo_pdb: 2MWO | apo_pdb: 1SSF | apo_bmrb: 5878 | holo_bmrb: 25347
A.
Apo/Holo HSQC Offset
Offsets: ΔH=-0.040, ΔN=-0.400
Apo
Holo (offset)
$^1$H $\delta$ (ppm)
$^{15}$N $\delta$ (ppm)
B.
CSP Classification
(TP) Sig. CSP in Binding Site (8)
(FP) Sig. CSP -- Allosteric (40)
(TN) low CSP -- Allosteric (58)
(FN) low CSP in Binding Site (1)
$\Delta\delta_{NH}$ (ppm)
Sequence
C.
CSP Mask
D.
Binding Site Mask
E.
Confusion Matrix Classification

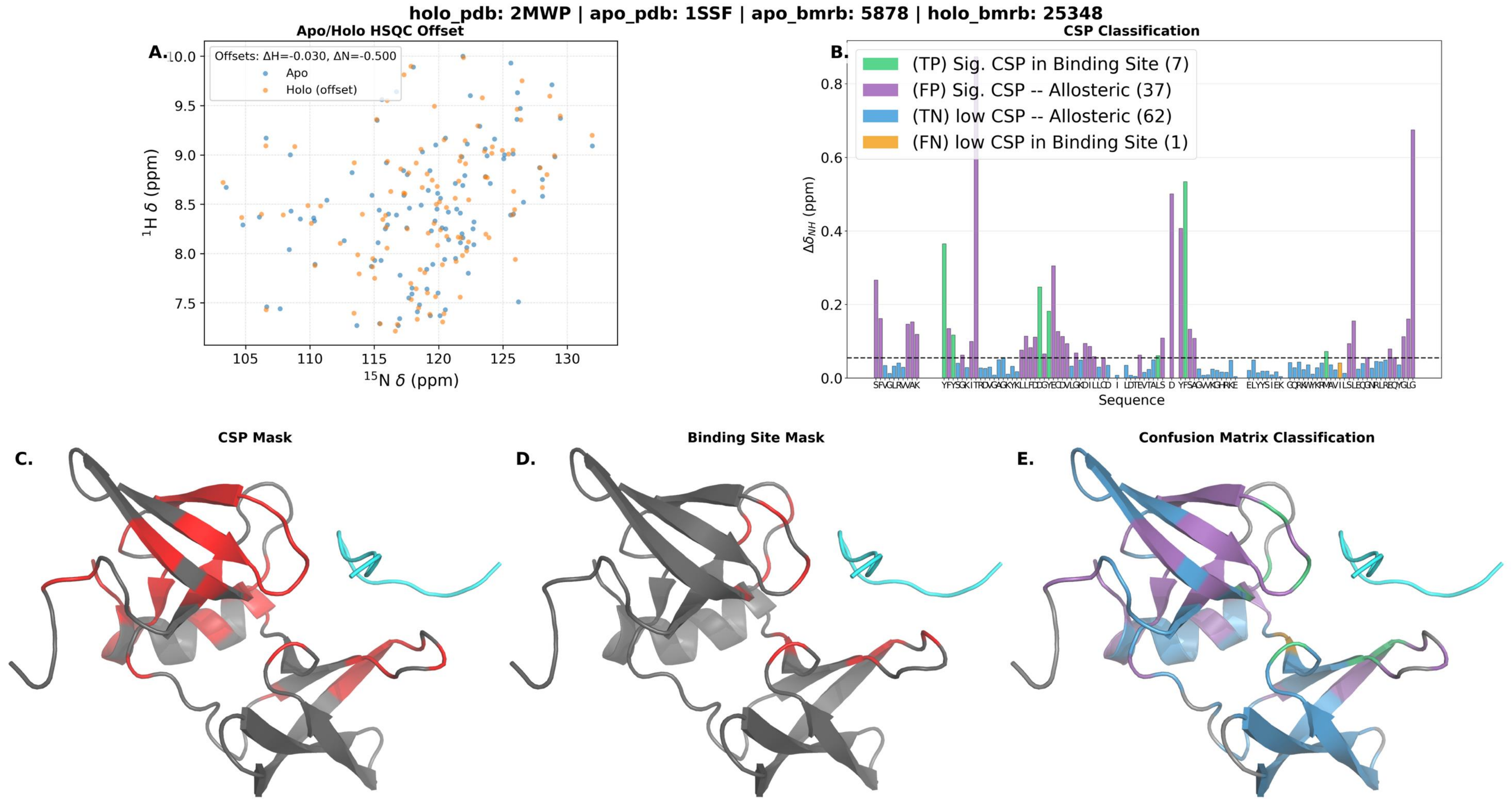

holo_pdb: 2MWP | apo_pdb: 1SSF | apo_bmrb: 5878 | holo_bmrb: 25348
A.
Apo/Holo HSQC Offset
Offsets: ΔH=-0.030, ΔN=-0.500
Apo
Holo (offset)
$^{1}$H δ (ppm)
$^{15}$N δ (ppm)
B.
CSP Classification
(TP) Sig. CSP in Binding Site (7)
(FP) Sig. CSP -- Allosteric (37)
(TN) low CSP -- Allosteric (62)
(FN) low CSP in Binding Site (1)
$\Delta\delta_{NH}$ (ppm)
Sequence
C.
CSP Mask
D.
Binding Site Mask
E.
Confusion Matrix Classification

**holo_pdb: 2MZW | apo_pdb: N/A | apo_bmrb: 25368 | holo_bmrb: 25504**

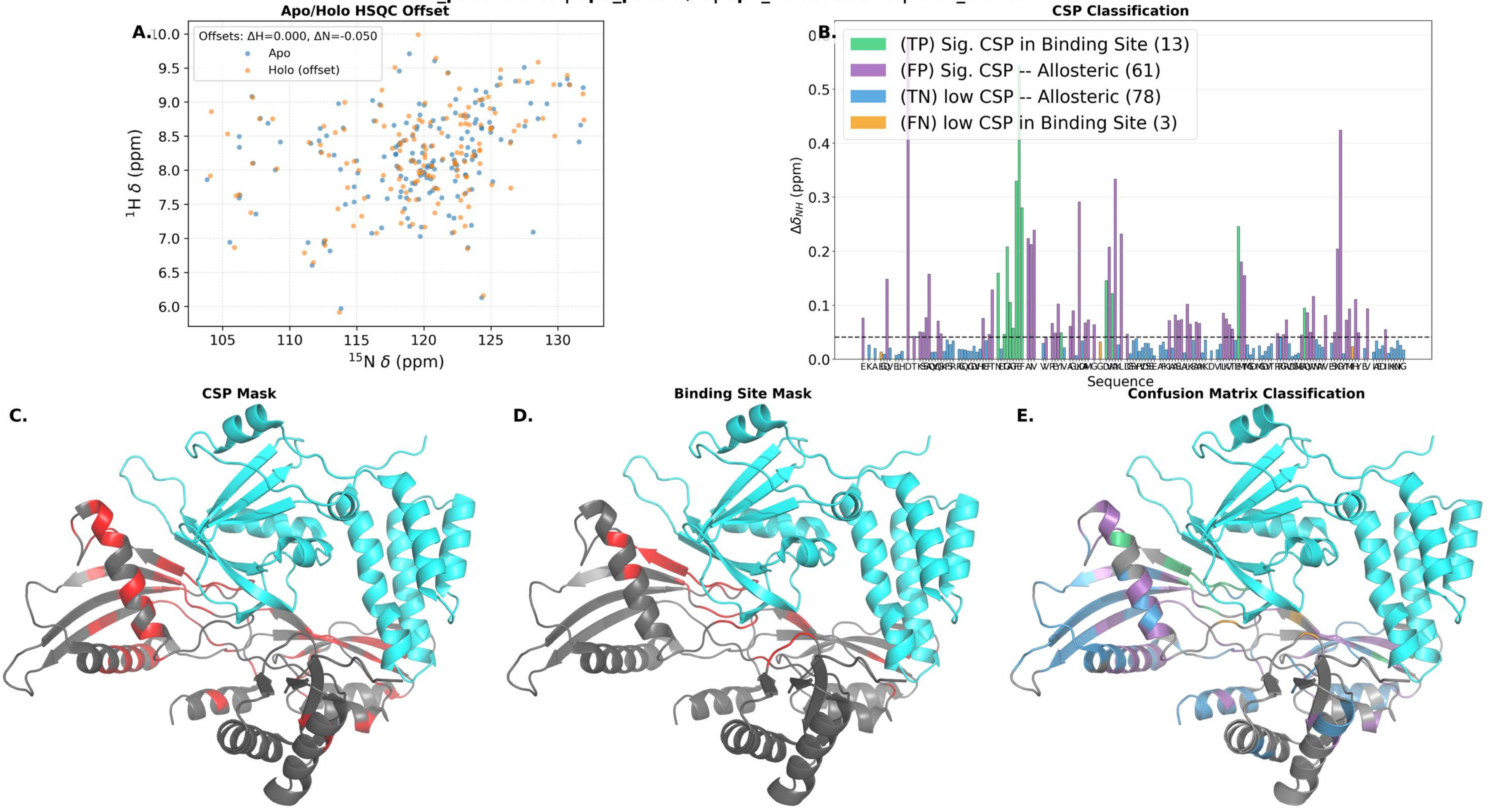

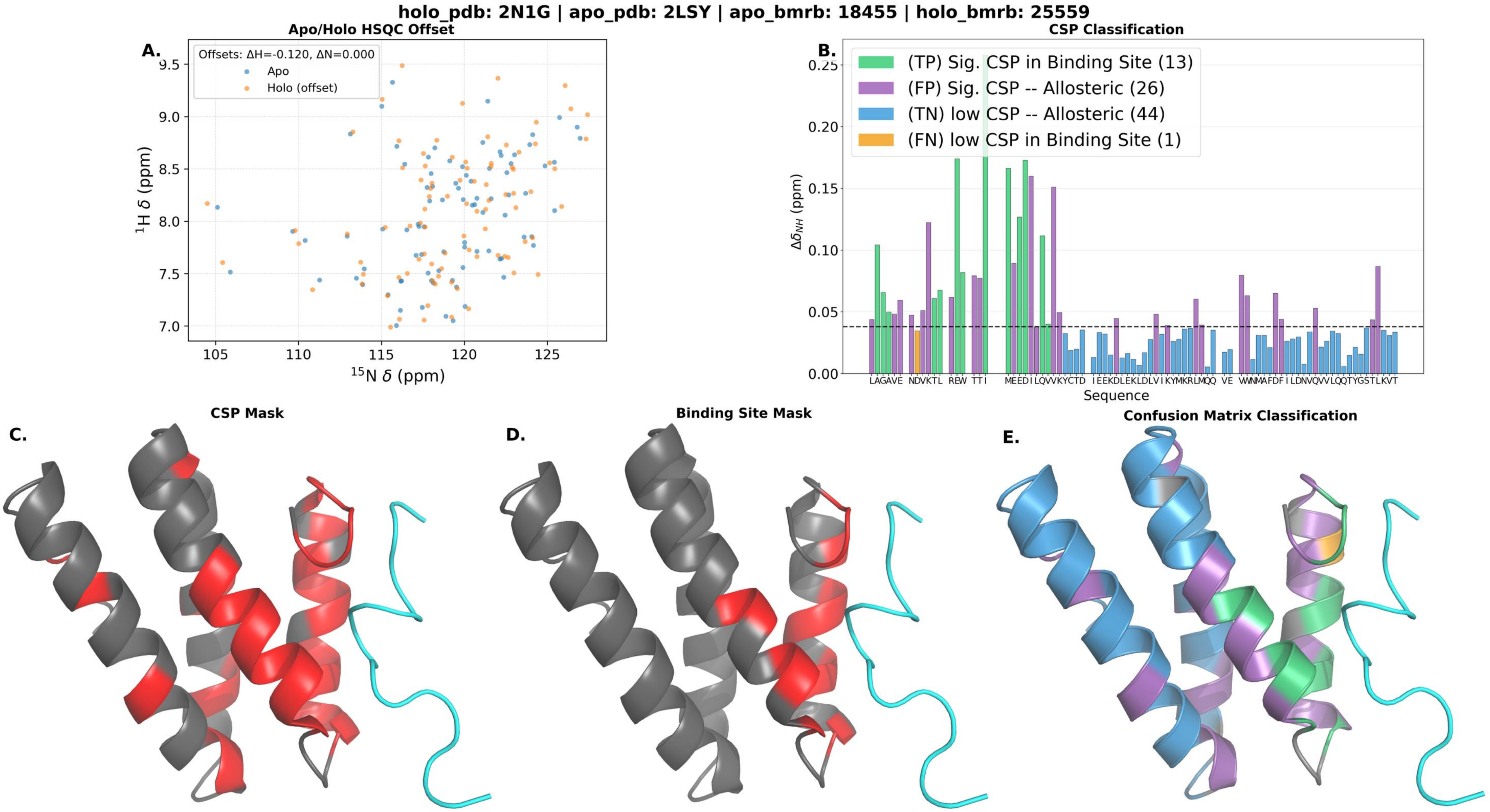
holo_pdb: 2N1G | apo_pdb: 2LSY | apo_bmrb: 18455 | holo_bmrb: 25559
Apo/Holo HSQC Offset
A.
Offsets: ΔH=-0.120, ΔN=0.000
Apo
Holo (offset)
$^{1}$H δ (ppm)
$^{15}$N δ (ppm)
CSP Classification
B.
(TP) Sig. CSP in Binding Site (13)
(FP) Sig. CSP -- Allosteric (26)
(TN) low CSP -- Allosteric (44)
(FN) low CSP in Binding Site (1)
$\Delta\delta_{NH}$ (ppm)
Sequence
C.
CSP Mask
D.
Binding Site Mask
E.
Confusion Matrix Classification

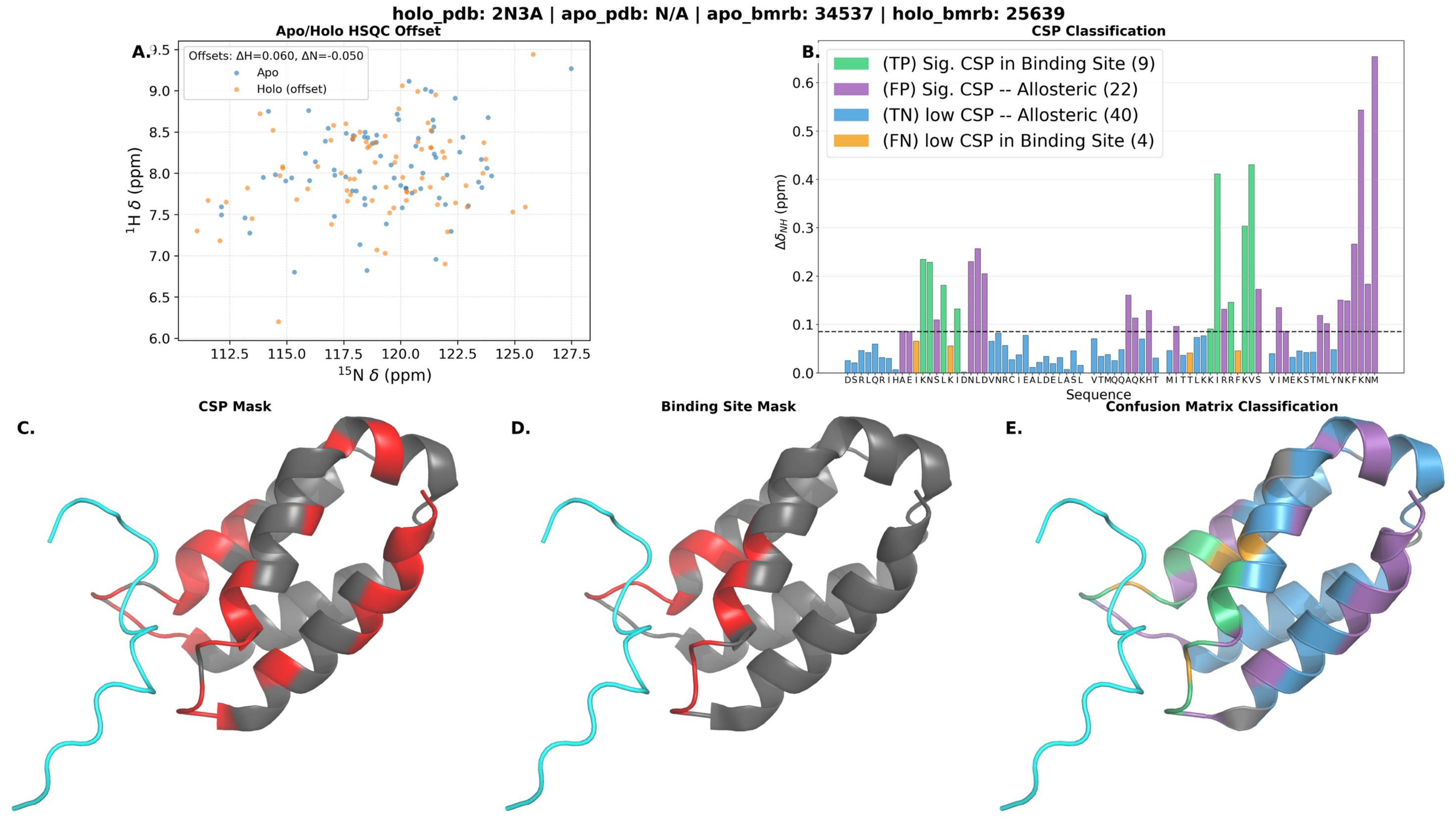
holo_pdb: 2N3A | apo_pdb: N/A | apo_bmrb: 34537 | holo_bmrb: 25639
A.
Apo/Holo HSQC Offset
Offsets: ΔH=0.060, ΔN=-0.050
Apo
Holo (offset)
1H δ (ppm)
15N δ (ppm)
B.
CSP Classification
(TP) Sig. CSP in Binding Site (9)
(FP) Sig. CSP -- Allosteric (22)
(TN) low CSP -- Allosteric (40)
(FN) low CSP in Binding Site (4)
ΔδNH (ppm)
Sequence
C.
CSP Mask
D.
Binding Site Mask
E.
Confusion Matrix Classification

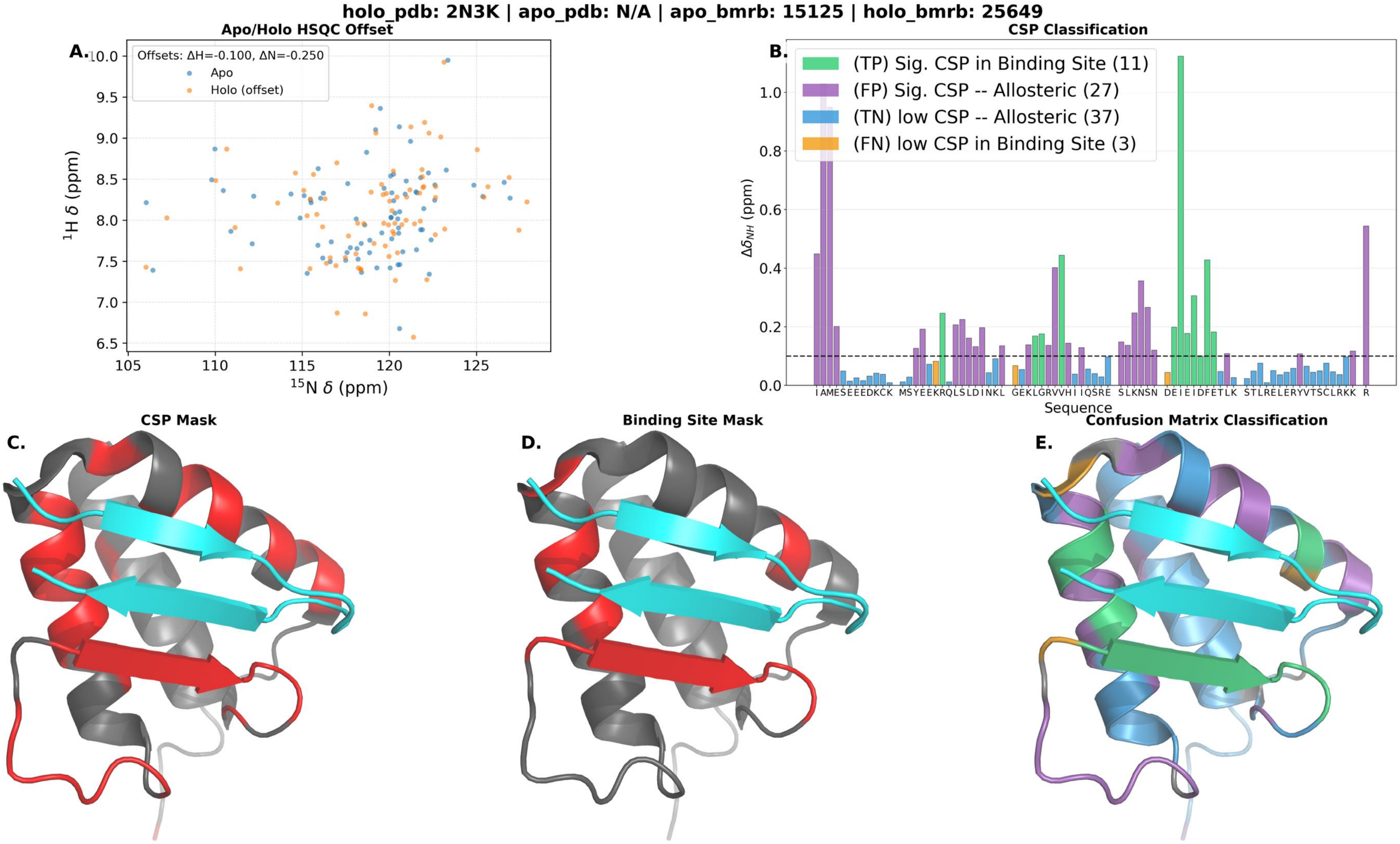

holo_pdb: 2N3K | apo_pdb: N/A | apo_bmrb: 15125 | holo_bmrb: 25649
Apo/Holo HSQC Offset
A.
Offsets: ΔH=-0.100, ΔN=-0.250
Apo
Holo (offset)
$^1$H δ (ppm)
$^{15}$N δ (ppm)
CSP Classification
B.
(TP) Sig. CSP in Binding Site (11)
(FP) Sig. CSP -- Allosteric (27)
(TN) low CSP -- Allosteric (37)
(FN) low CSP in Binding Site (3)
$\Delta\delta_{NH}$ (ppm)
Sequence
CSP Mask
C.
Binding Site Mask
D.
Confusion Matrix Classification
E.

**holo_pdb: 2N55 | apo_pdb: N/A | apo_bmrb: 52209 | holo_bmrb: 25694**

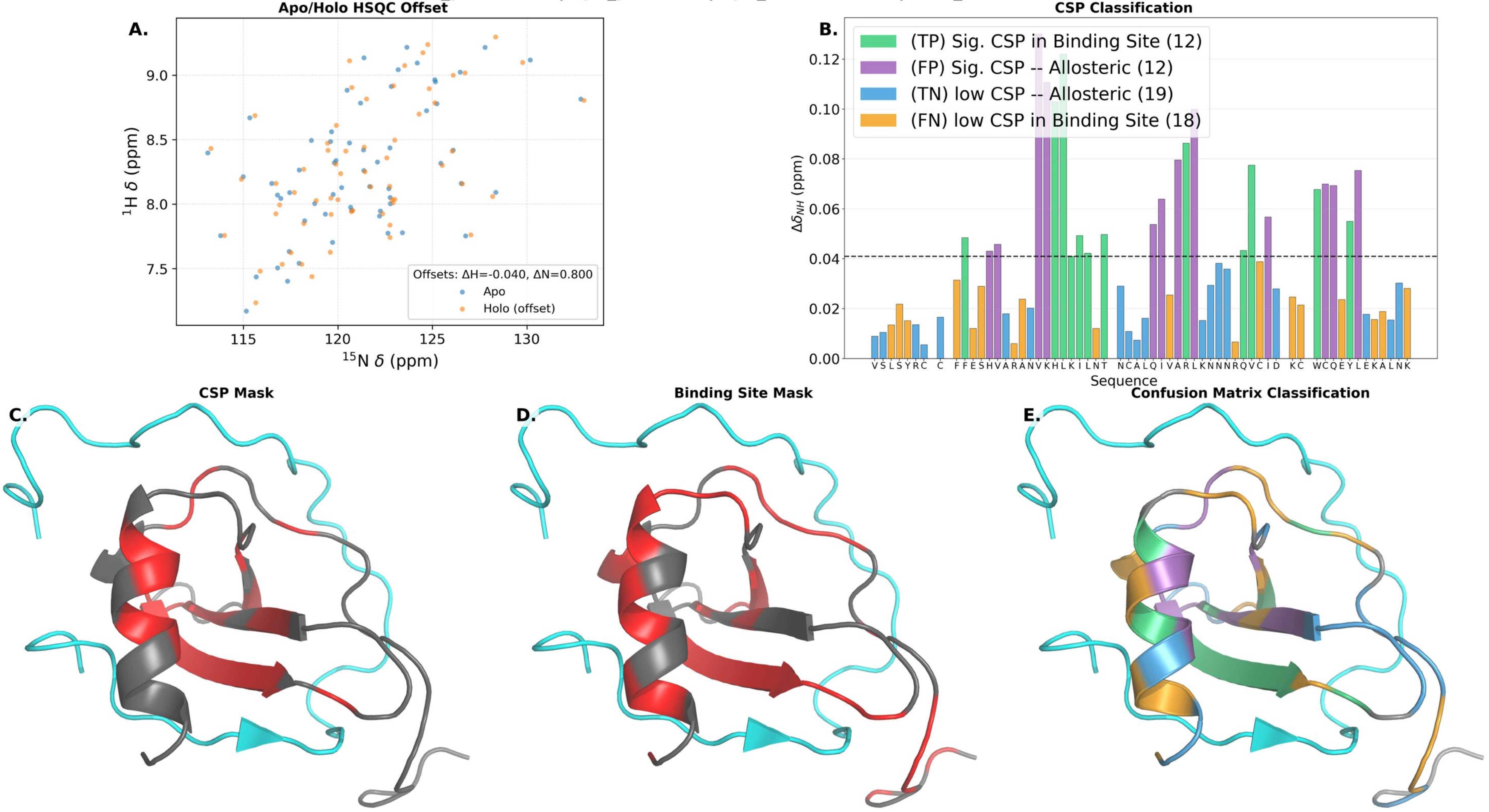

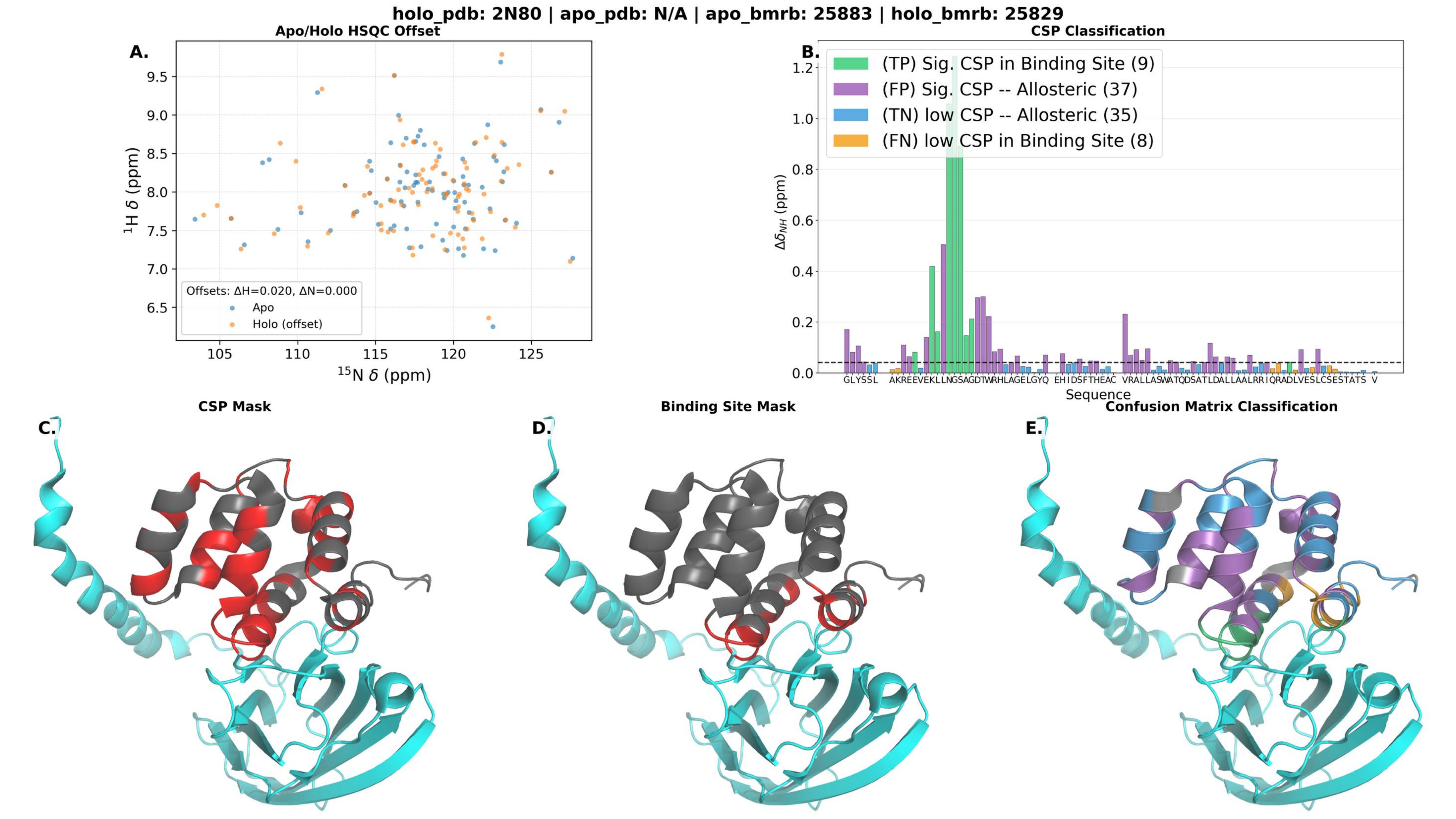

holo_pdb: 2N80 | apo_pdb: N/A | apo_bmrb: 25883 | holo_bmrb: 25829
A.
Apo/Holo HSQC Offset
$^{1}$H δ (ppm)
$^{15}$N δ (ppm)
Offsets: ΔH=0.020, ΔN=0.000
Apo
Holo (offset)
B.
CSP Classification
(TP) Sig. CSP in Binding Site (9)
(FP) Sig. CSP -- Allosteric (37)
(TN) low CSP -- Allosteric (35)
(FN) low CSP in Binding Site (8)
$\Delta\delta_{NH}$ (ppm)
Sequence
C.
CSP Mask
D.
Binding Site Mask
E.
Confusion Matrix Classification

holo_pdb: 2N8T | apo_pdb: N/A | apo_bmrb: 25866 | holo_bmrb: 25865

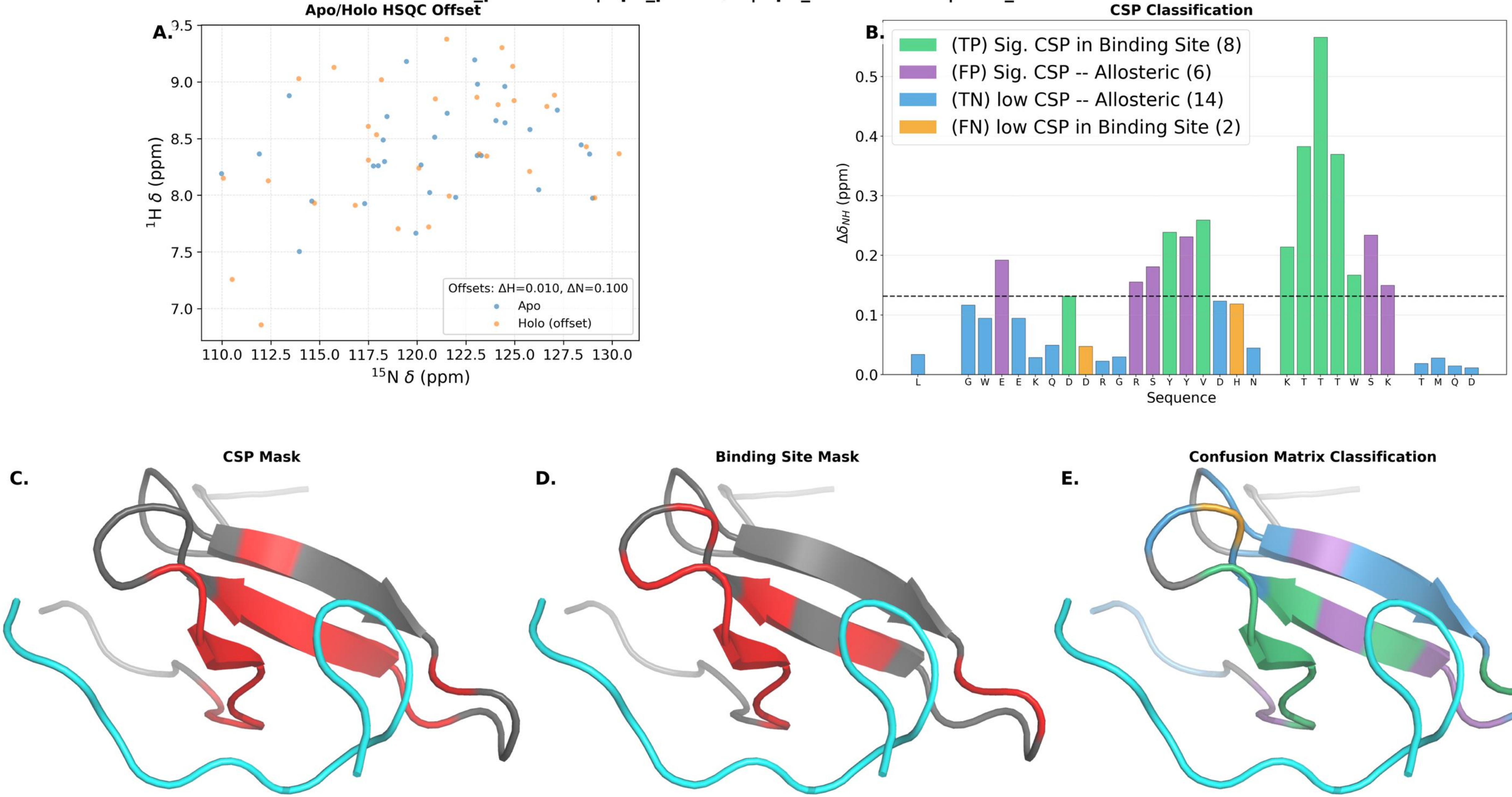

**holo_pdb: 2N9P | apo_pdb: N/A | apo_bmrb: 25913 | holo_bmrb: 25914**

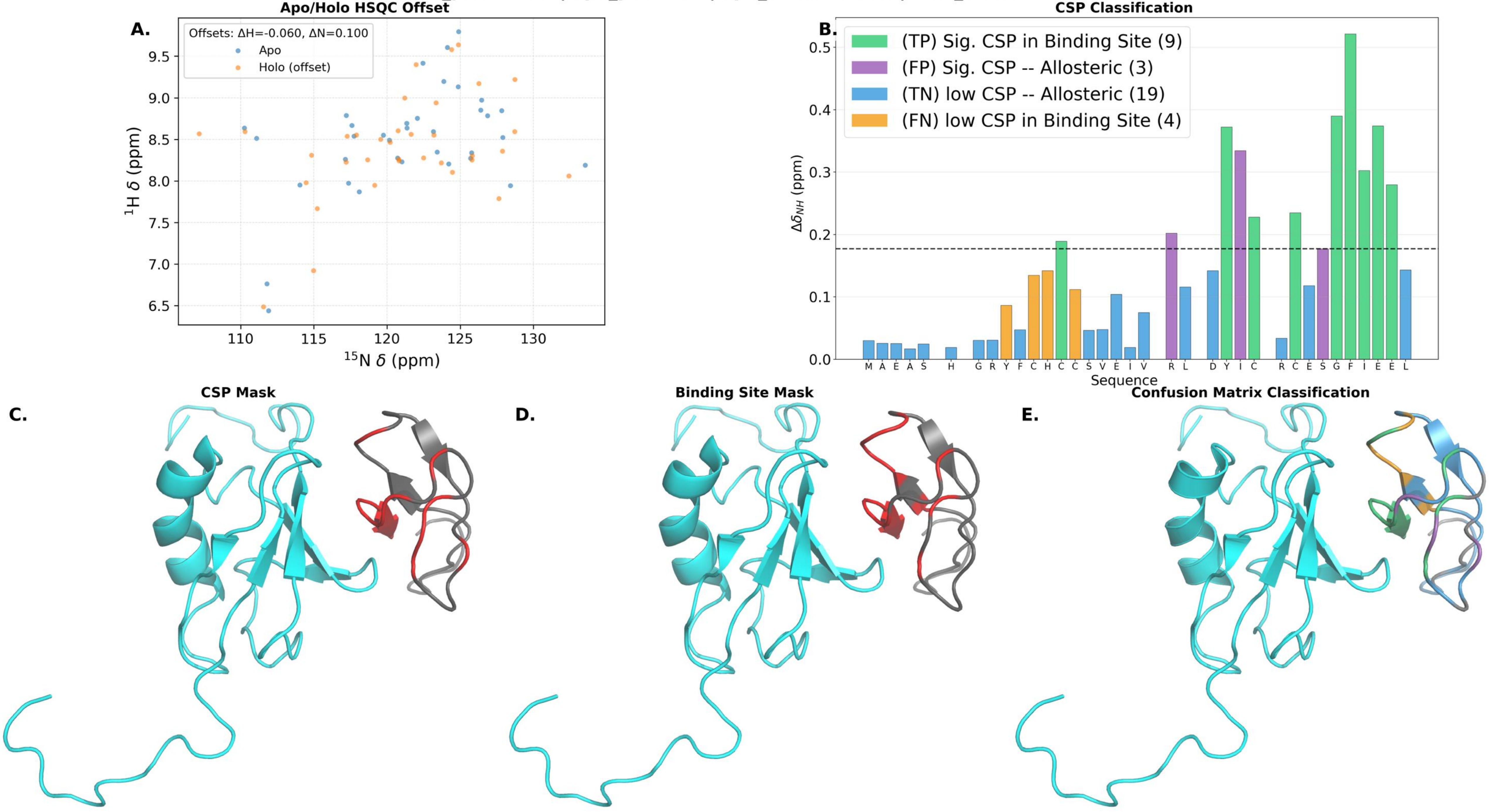

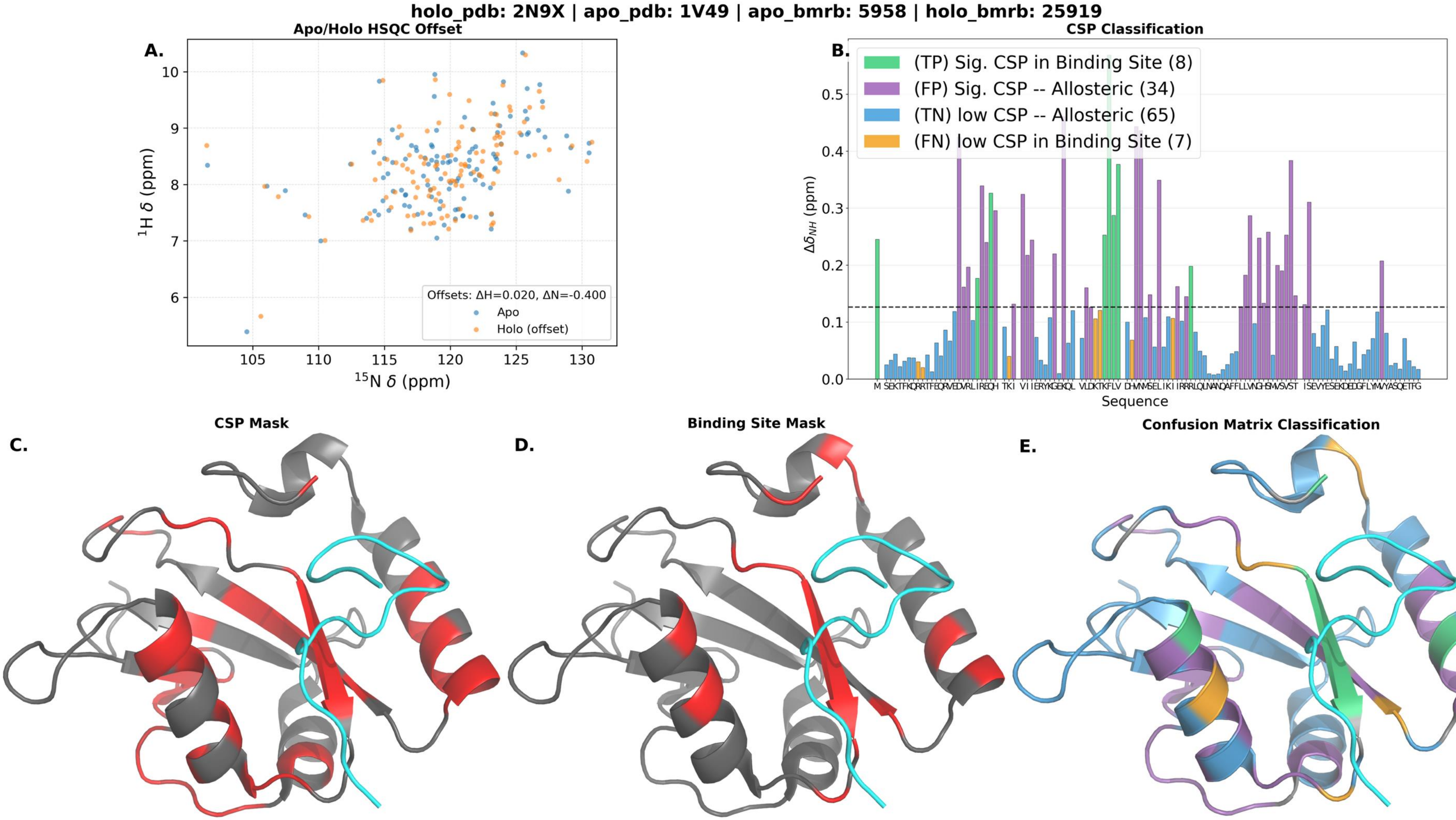
holo_pdb: 2N9X | apo_pdb: 1V49 | apo_bmrb: 5958 | holo_bmrb: 25919
A.
Apo/Holo HSQC Offset
Offsets: ΔH=0.020, ΔN=-0.400
Apo
Holo (offset)
¹H δ (ppm)
¹⁵N δ (ppm)
B.
CSP Classification
(TP) Sig. CSP in Binding Site (8)
(FP) Sig. CSP -- Allosteric (34)
(TN) low CSP -- Allosteric (65)
(FN) low CSP in Binding Site (7)
Δδ_NH (ppm)
Sequence
C.
CSP Mask
D.
Binding Site Mask
E.
Confusion Matrix Classification

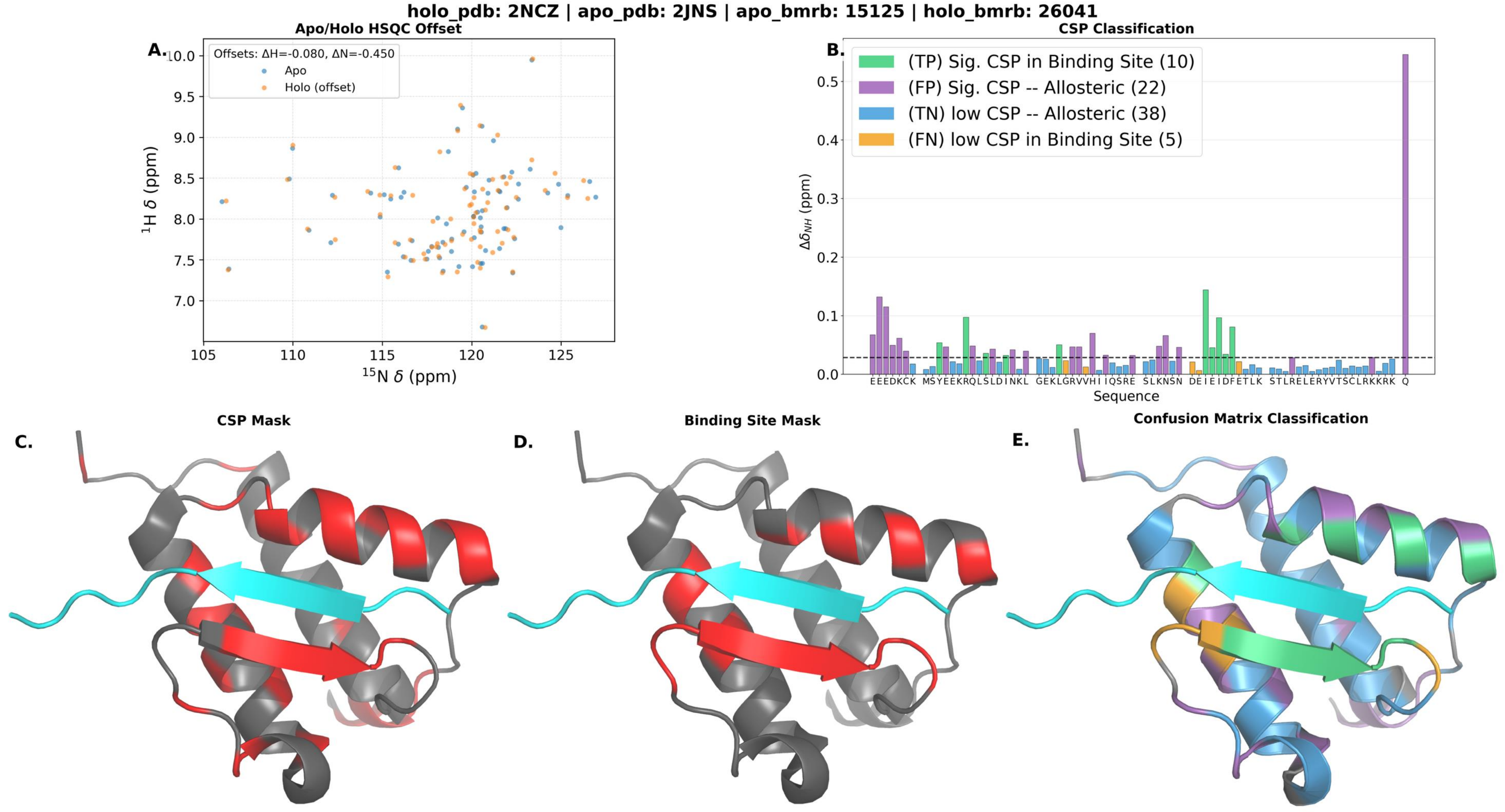

holo_pdb: 2NCZ | apo_pdb: 2JNS | apo_bmrb: 15125 | holo_bmrb: 26041
A.
Apo/Holo HSQC Offset
Offsets: ΔH=-0.080, ΔN=-0.450
Apo
Holo (offset)
1H δ (ppm)
15N δ (ppm)
B.
CSP Classification
(TP) Sig. CSP in Binding Site (10)
(FP) Sig. CSP -- Allosteric (22)
(TN) low CSP -- Allosteric (38)
(FN) low CSP in Binding Site (5)
ΔδNH (ppm)
Sequence
C.
CSP Mask
D.
Binding Site Mask
E.
Confusion Matrix Classification

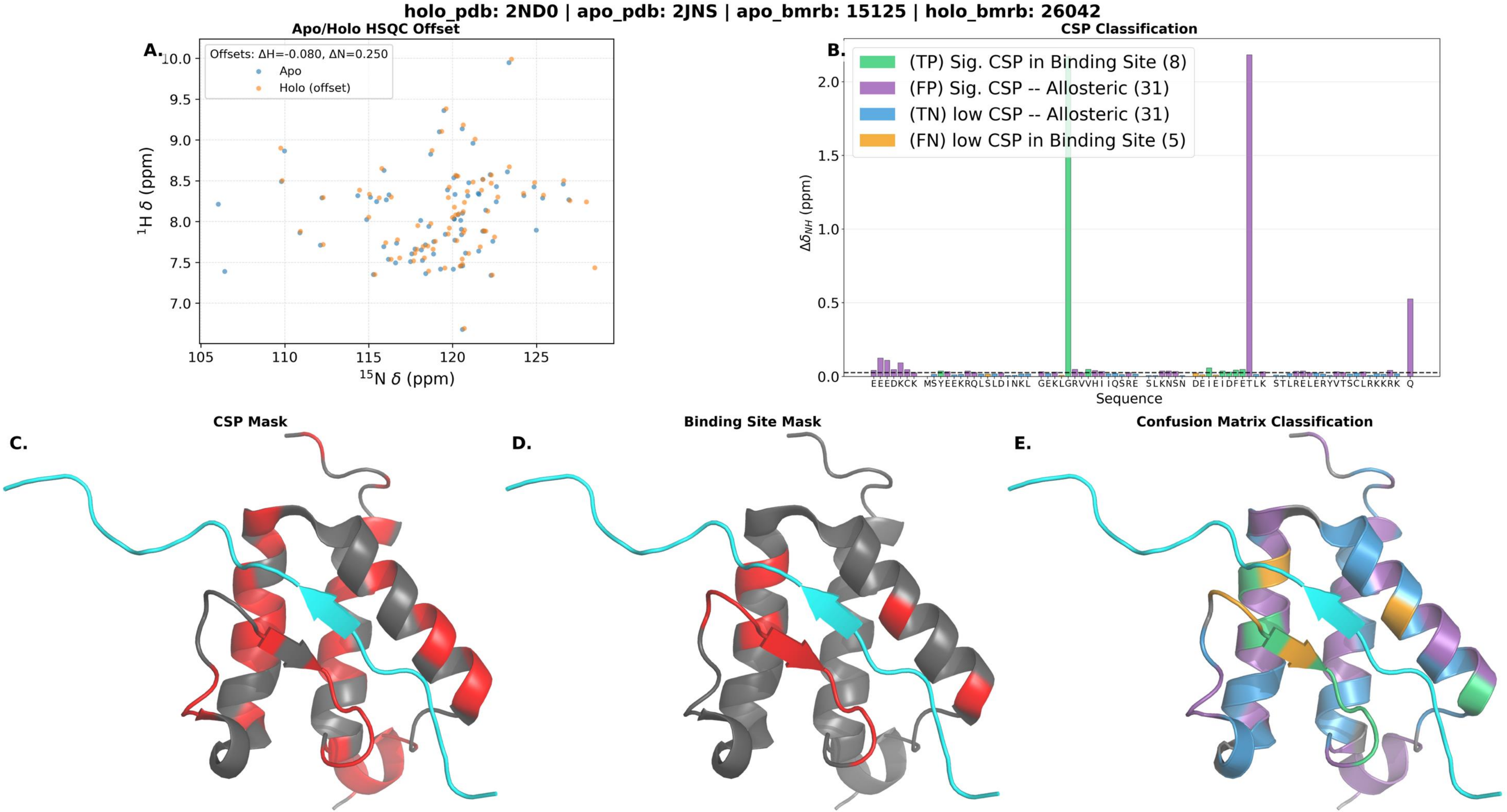

holo_pdb: 2ND0 | apo_pdb: 2JNS | apo_bmrb: 15125 | holo_bmrb: 26042
Apo/Holo HSQC Offset
A.
Offsets: ΔH=-0.080, ΔN=0.250
Apo
Holo (offset)
1H δ (ppm)
15N δ (ppm)
CSP Classification
B.
(TP) Sig. CSP in Binding Site (8)
(FP) Sig. CSP -- Allosteric (31)
(TN) low CSP -- Allosteric (31)
(FN) low CSP in Binding Site (5)
ΔδNH (ppm)
Sequence
CSP Mask
C.
Binding Site Mask
D.
Confusion Matrix Classification
E.

**holo_pdb: 2ND1 | apo_pdb: 2JNS | apo_bmrb: 15125 | holo_bmrb: 26043**

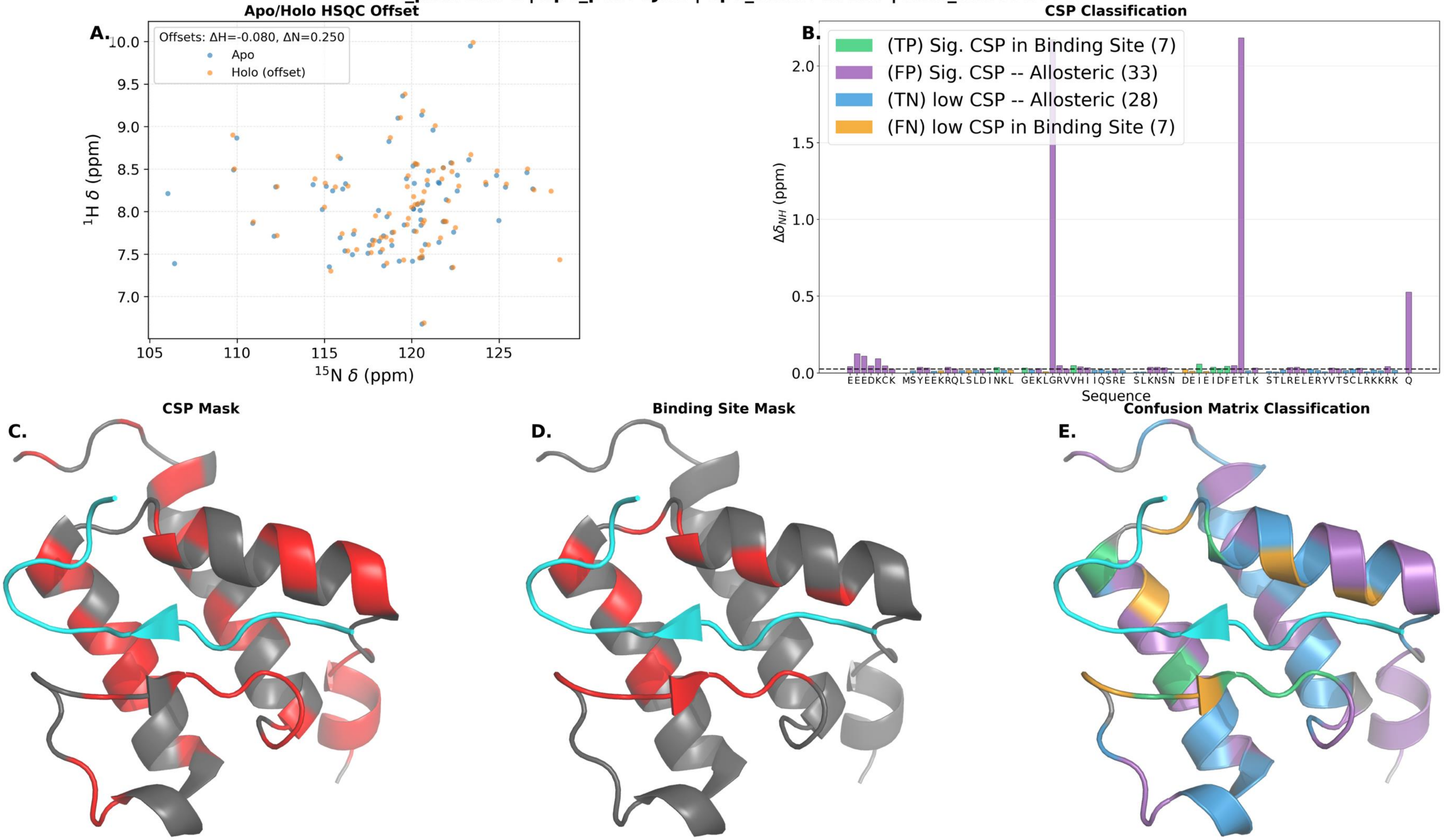

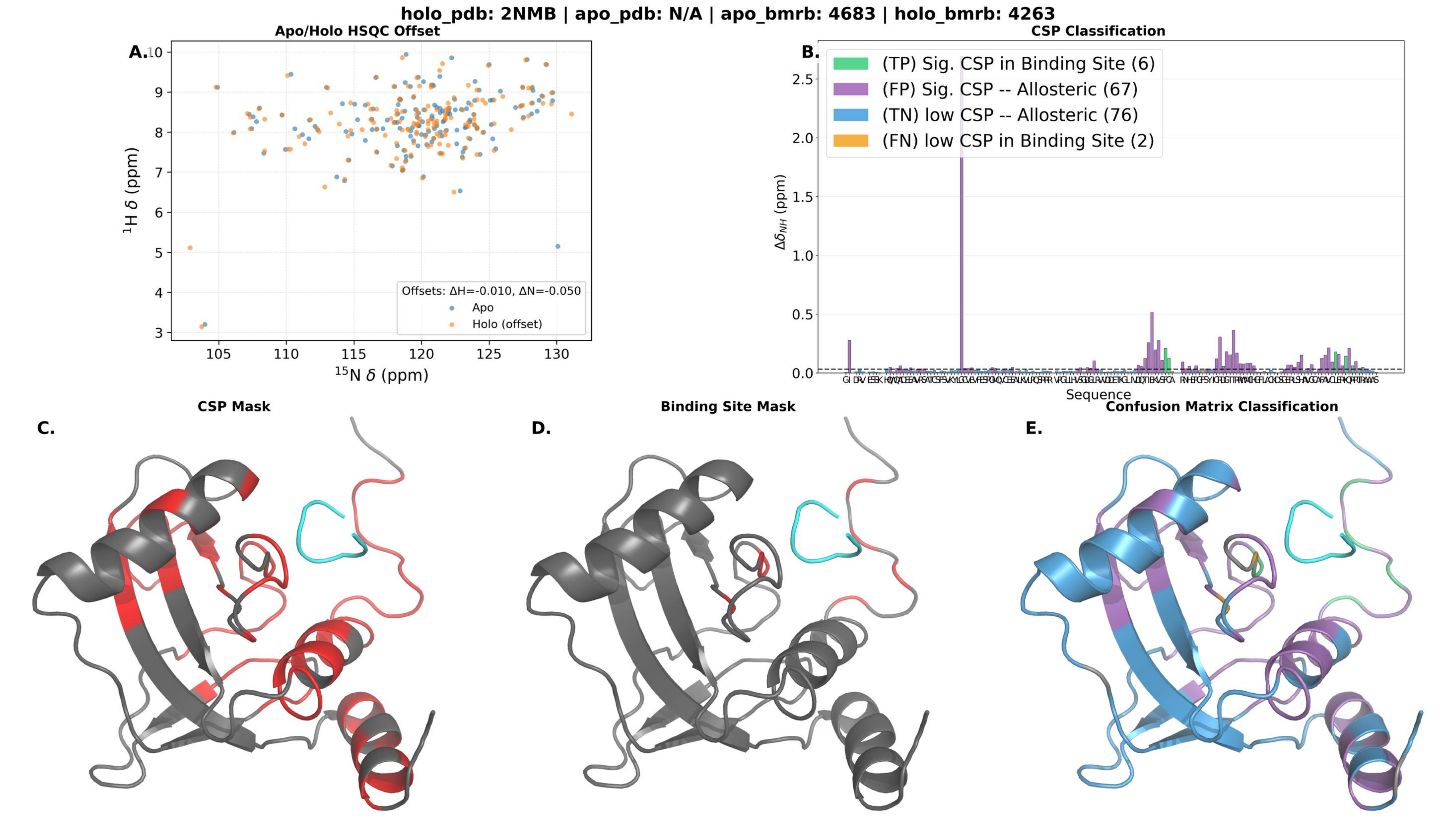
holo_pdb: 2NMB | apo_pdb: N/A | apo_bmrb: 4683 | holo_bmrb: 4263
A.
Apo/Holo HSQC Offset
$^{1}$H δ (ppm)
$^{15}$N δ (ppm)
Offsets: ΔH=-0.010, ΔN=-0.050
Apo
Holo (offset)
B.
CSP Classification
(TP) Sig. CSP in Binding Site (6)
(FP) Sig. CSP -- Allosteric (67)
(TN) low CSP -- Allosteric (76)
(FN) low CSP in Binding Site (2)
$\Delta\delta_{NH}$ (ppm)
Sequence
C.
CSP Mask
D.
Binding Site Mask
E.
Confusion Matrix Classification

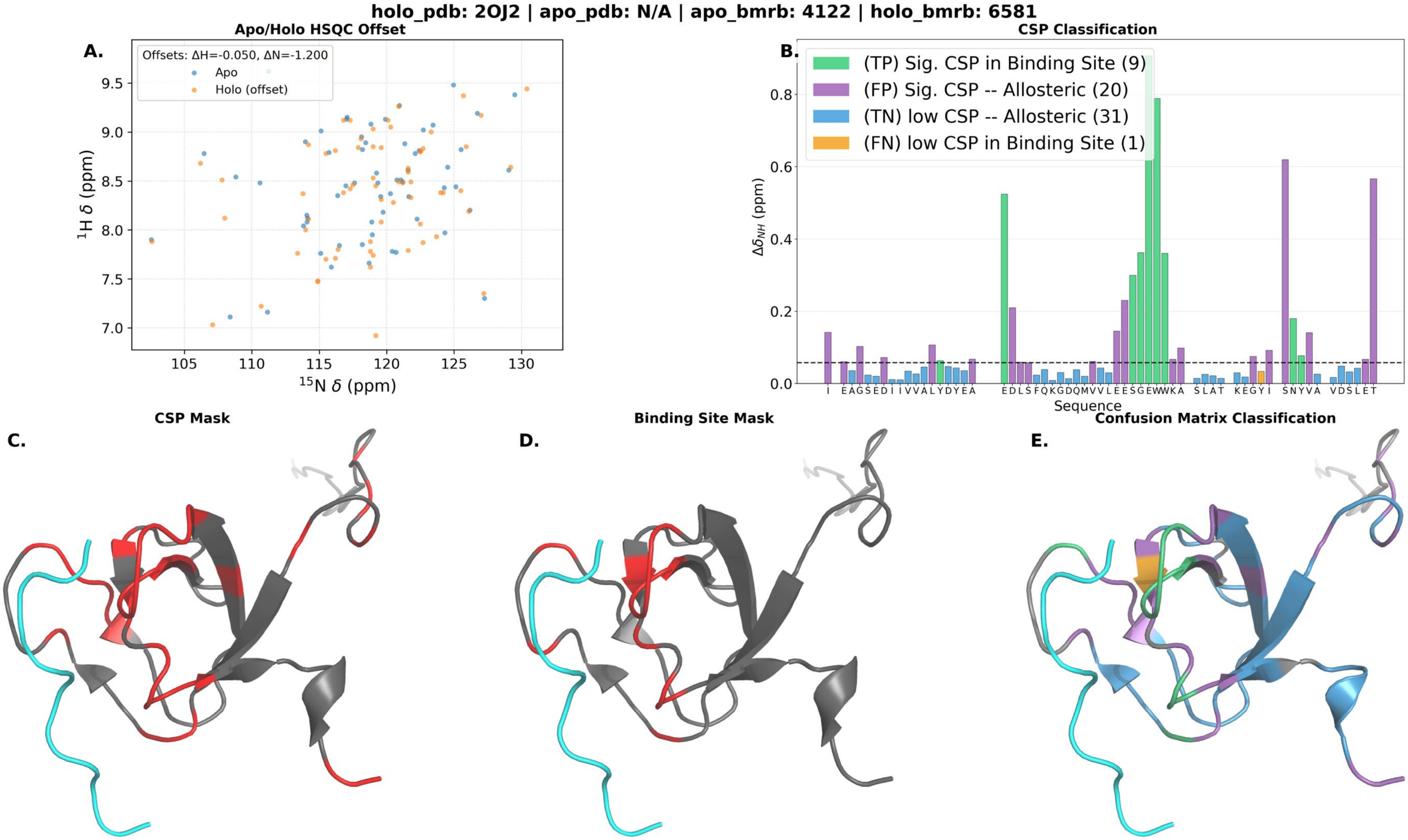
holo_pdb: 2OJ2 | apo_pdb: N/A | apo_bmrb: 4122 | holo_bmrb: 6581
A.
Apo/Holo HSQC Offset
Offsets: ΔH=-0.050, ΔN=-1.200
Apo
Holo (offset)
¹H δ (ppm)
¹⁵N δ (ppm)
9.5
9.0
8.5
8.0
7.5
7.0
105
110
115
120
125
130
B.
CSP Classification
(TP) Sig. CSP in Binding Site (9)
(FP) Sig. CSP -- Allosteric (20)
(TN) low CSP -- Allosteric (31)
(FN) low CSP in Binding Site (1)
Δδ_NH (ppm)
0.0
0.2
0.4
0.6
0.8
I EAGSEDI IVVALYDYEA EDLSFQKGDQMVVLEESGEWWKA SLAT KEGYI SNYVA VDSLET
Sequence
C.
CSP Mask
D.
Binding Site Mask
E.
Confusion Matrix Classification

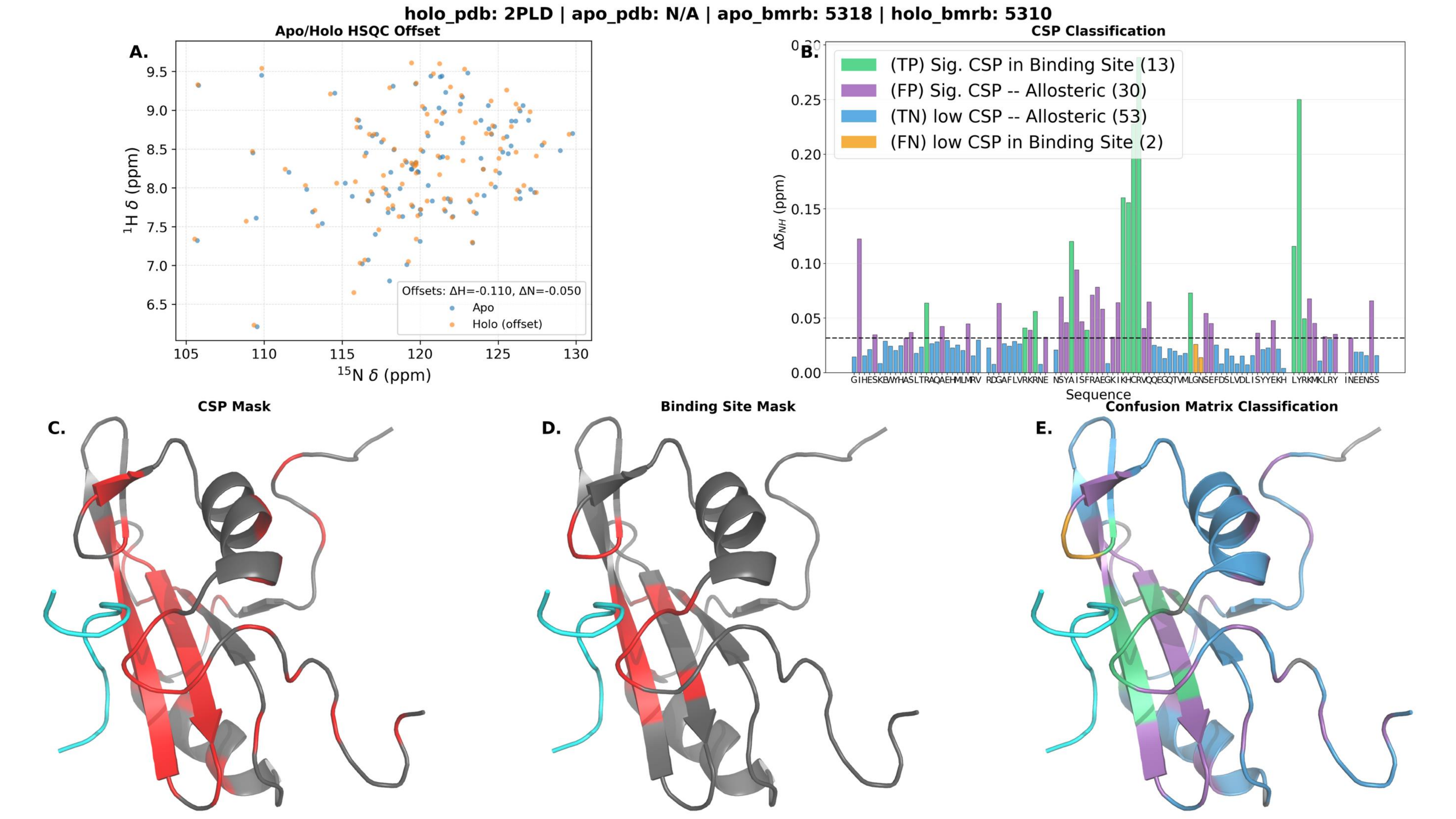

holo_pdb: 2PLD | apo_pdb: N/A | apo_bmrb: 5318 | holo_bmrb: 5310
Apo/Holo HSQC Offset
A.
$^1$H δ (ppm)
$^{15}$N δ (ppm)
Offsets: ΔH=-0.110, ΔN=-0.050
Apo
Holo (offset)
CSP Classification
B.
(TP) Sig. CSP in Binding Site (13)
(FP) Sig. CSP -- Allosteric (30)
(TN) low CSP -- Allosteric (53)
(FN) low CSP in Binding Site (2)
$\Delta\delta_{NH}$ (ppm)
GIHESKEWYHASLTRAQAEHMLMRV RDGAFLVRKRNE NSYAISFRAEGKIKHCRVQQEGQTVMLGNSEFDSLVDLISYYEKH LYRKMKLRY INEENSS
Sequence
CSP Mask
C.
Binding Site Mask
D.
Confusion Matrix Classification
E.

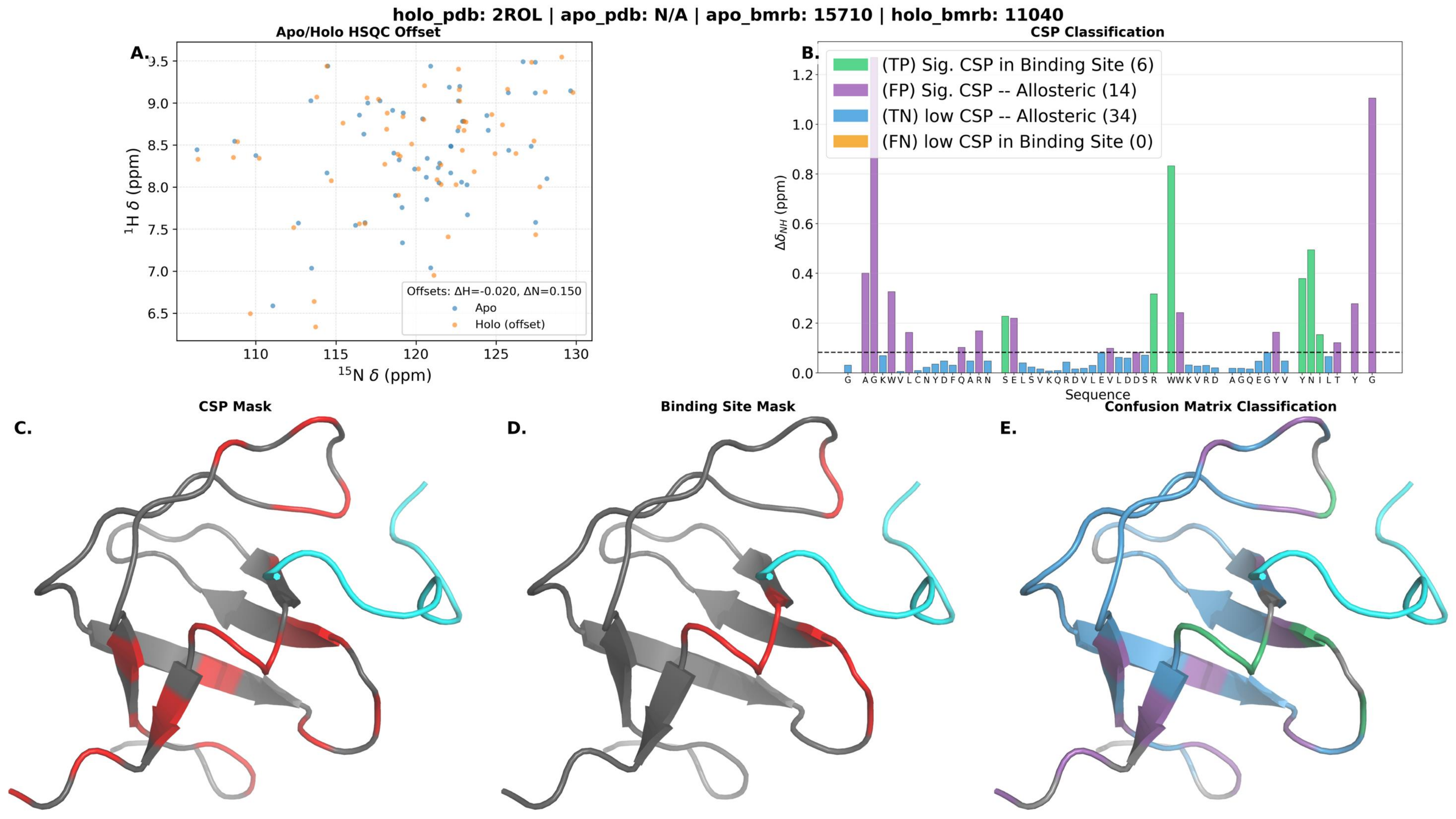

holo_pdb: 2ROL | apo_pdb: N/A | apo_bmrb: 15710 | holo_bmrb: 11040
A.
Apo/Holo HSQC Offset
$^1$H δ (ppm)
$^{15}$N δ (ppm)
Offsets: ΔH=-0.020, ΔN=0.150
Apo
Holo (offset)
B.
CSP Classification
(TP) Sig. CSP in Binding Site (6)
(FP) Sig. CSP -- Allosteric (14)
(TN) low CSP -- Allosteric (34)
(FN) low CSP in Binding Site (0)
$\Delta\delta_{NH}$ (ppm)
Sequence
G A G K W V L C N Y D F Q A R N S E L S V K Q R D V L E V L D D S R W W K V R D A G Q E G Y V Y N I L T Y G
C.
CSP Mask
D.
Binding Site Mask
E.
Confusion Matrix Classification

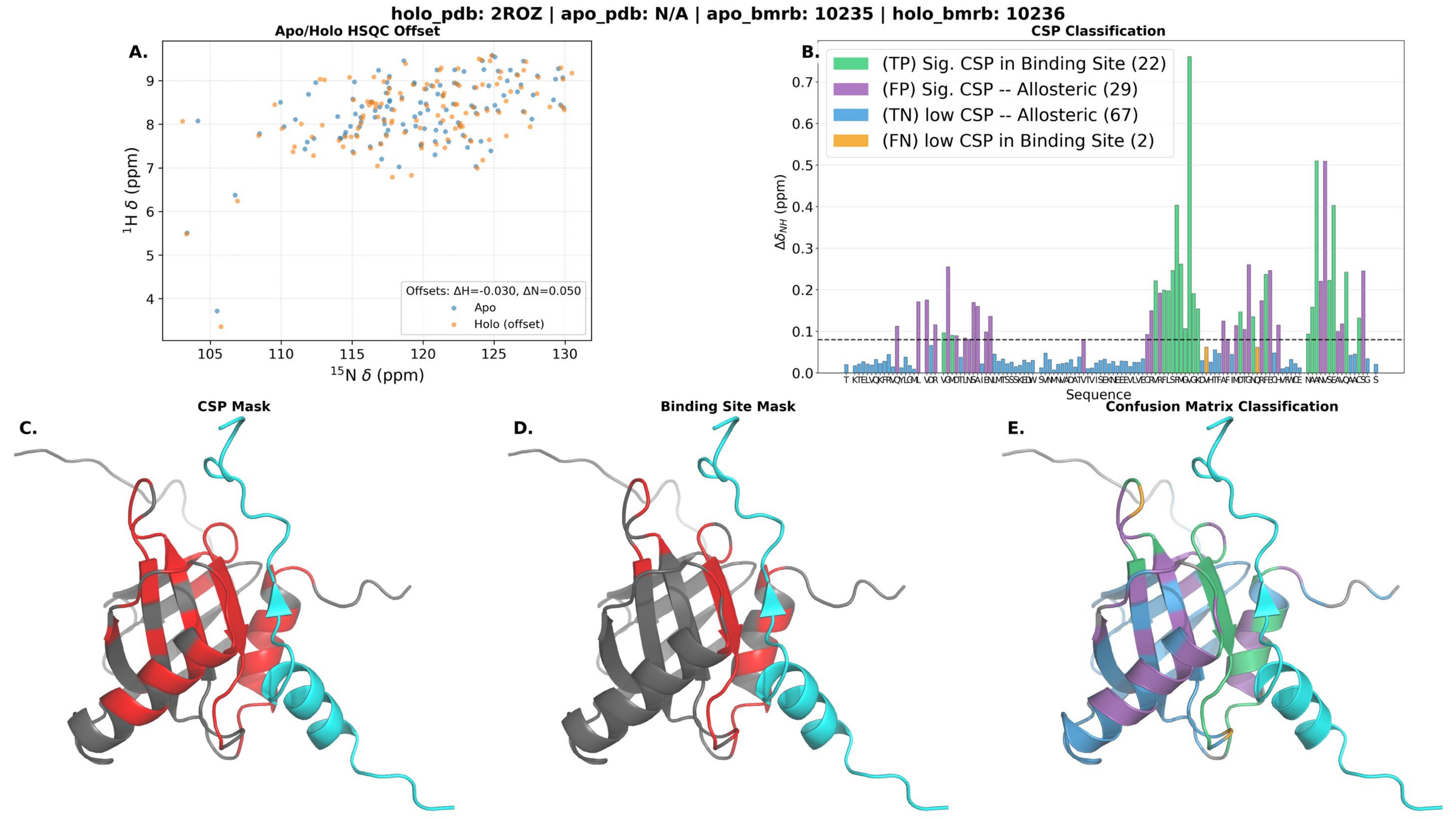
holo_pdb: 2ROZ | apo_pdb: N/A | apo_bmrb: 10235 | holo_bmrb: 10236
A.
Apo/Holo HSQC Offset
Offsets: ΔH=-0.030, ΔN=0.050
Apo
Holo (offset)
1H δ (ppm)
15N δ (ppm)
B.
CSP Classification
(TP) Sig. CSP in Binding Site (22)
(FP) Sig. CSP -- Allosteric (29)
(TN) low CSP -- Allosteric (67)
(FN) low CSP in Binding Site (2)
ΔδNH (ppm)
Sequence
C.
CSP Mask
D.
Binding Site Mask
E.
Confusion Matrix Classification

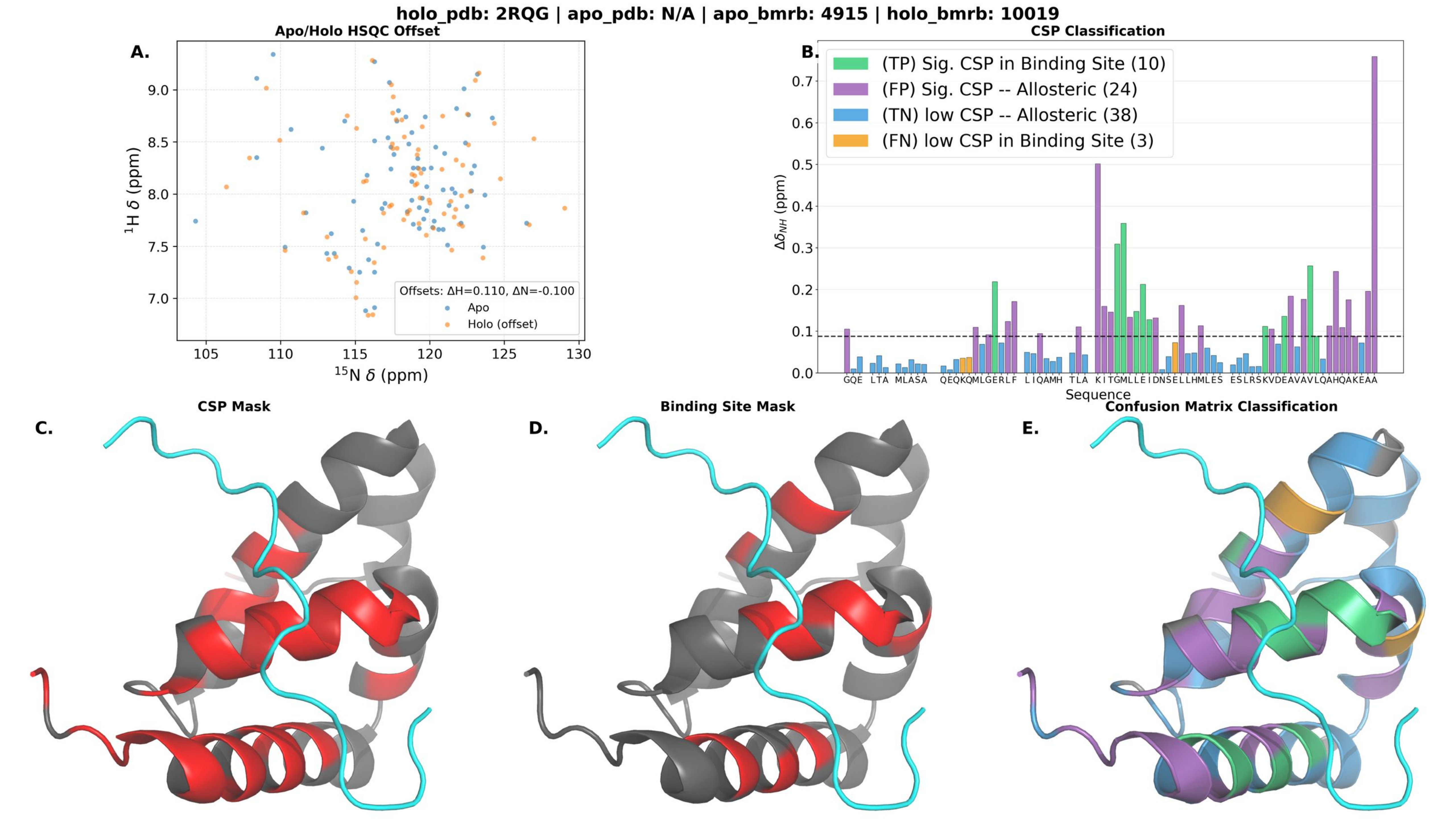

holo_pdb: 2RQG | apo_pdb: N/A | apo_bmrb: 4915 | holo_bmrb: 10019
A.
Apo/Holo HSQC Offset
Offsets: ΔH=0.110, ΔN=-0.100
Apo
Holo (offset)
$^{1}$H δ (ppm)
$^{15}$N δ (ppm)
B.
CSP Classification
(TP) Sig. CSP in Binding Site (10)
(FP) Sig. CSP -- Allosteric (24)
(TN) low CSP -- Allosteric (38)
(FN) low CSP in Binding Site (3)
$\Delta\delta_{NH}$ (ppm)
Sequence
C.
CSP Mask
D.
Binding Site Mask
E.
Confusion Matrix Classification

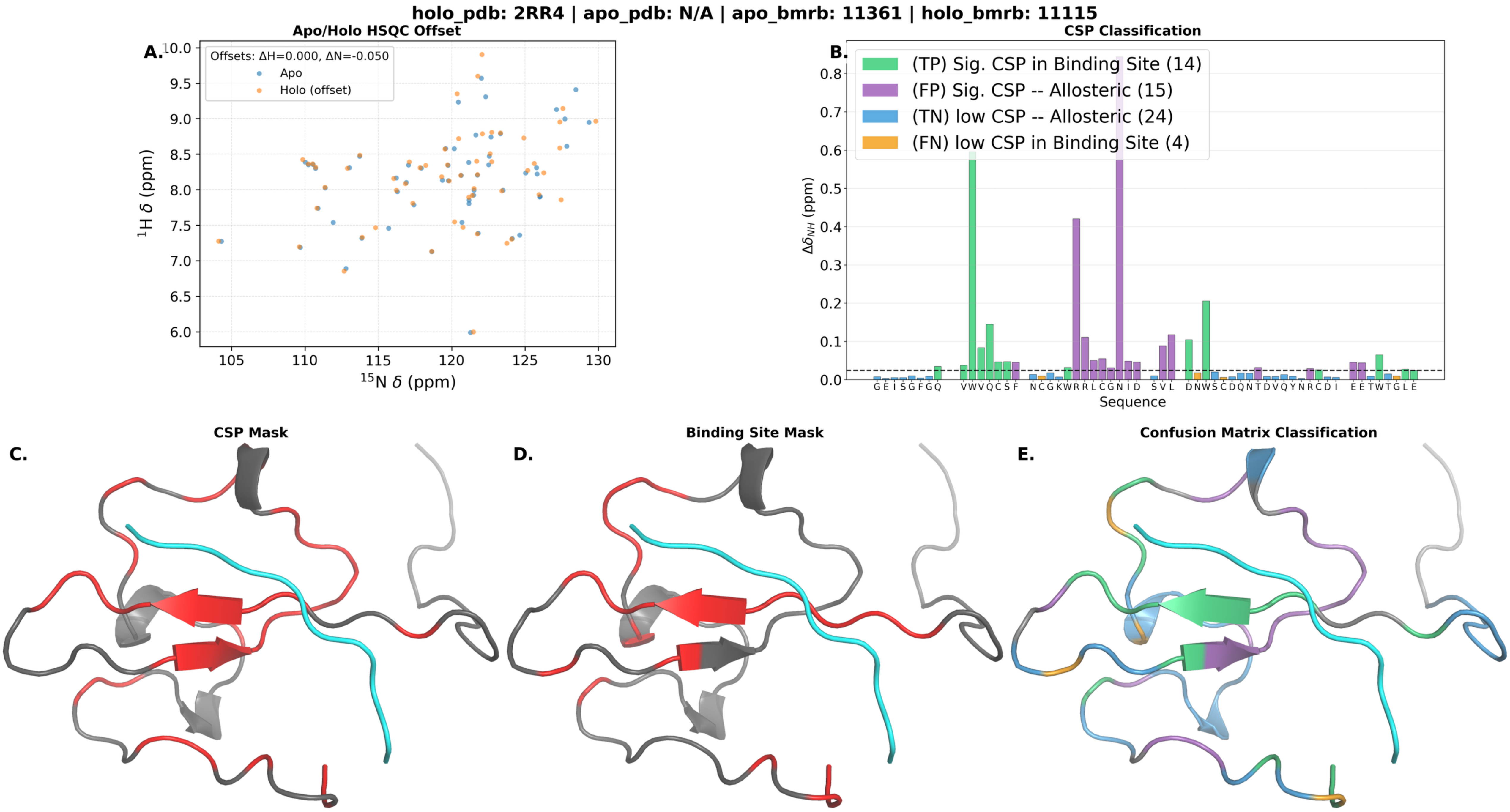
holo_pdb: 2RR4 | apo_pdb: N/A | apo_bmrb: 11361 | holo_bmrb: 11115
A.
Apo/Holo HSQC Offset
Offsets: ΔH=0.000, ΔN=-0.050
Apo
Holo (offset)
$^{1}$H δ (ppm)
$^{15}$N δ (ppm)
B.
CSP Classification
(TP) Sig. CSP in Binding Site (14)
(FP) Sig. CSP -- Allosteric (15)
(TN) low CSP -- Allosteric (24)
(FN) low CSP in Binding Site (4)
$\Delta\delta_{NH}$ (ppm)
G E I S G F G Q V W V Q C S F N C G K W R R L C G N I D S V L D N W S C D Q N T D V Q Y N R C D I E E T W T G L E
Sequence
C.
CSP Mask
D.
Binding Site Mask
E.
Confusion Matrix Classification

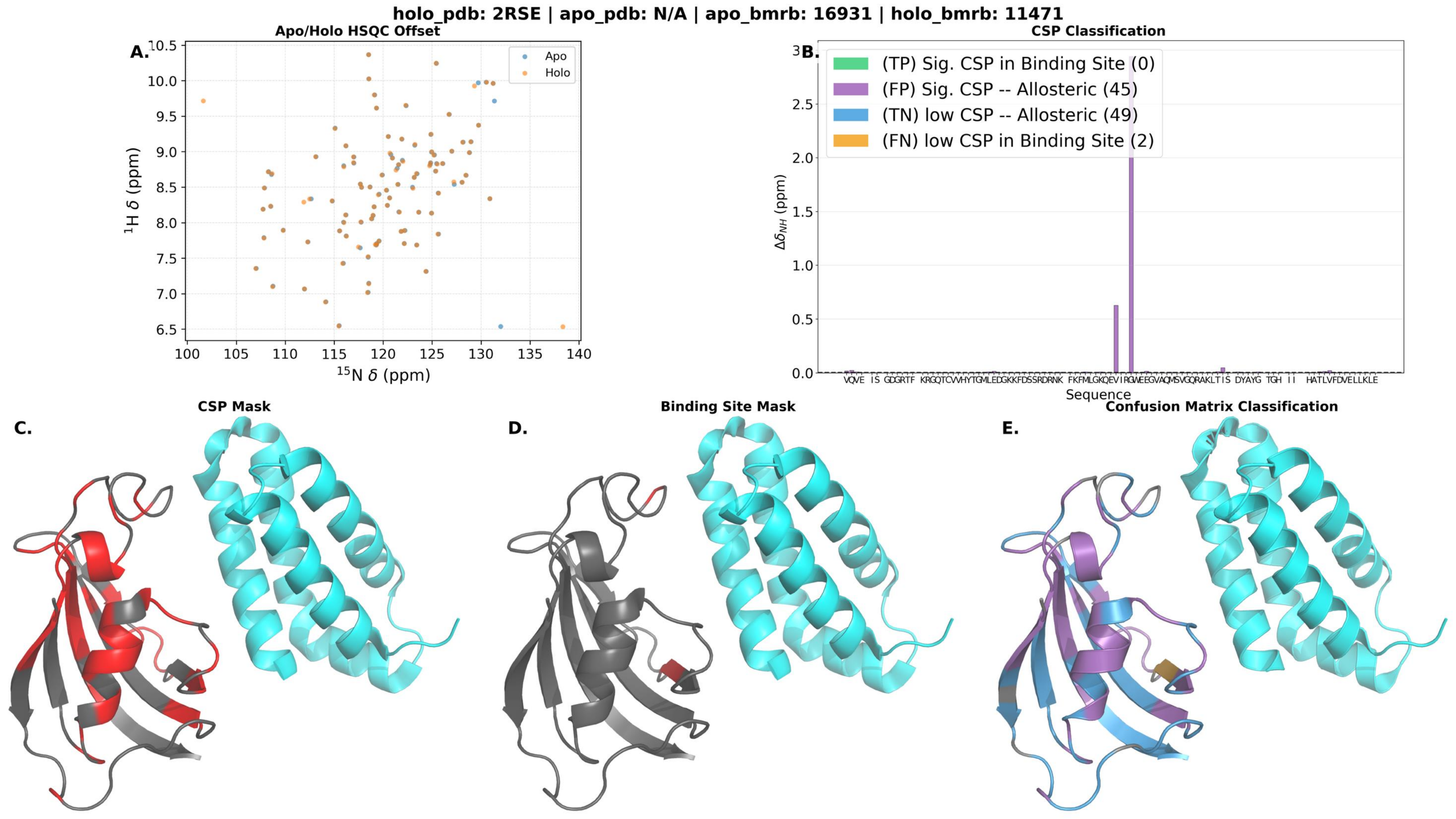

holo_pdb: 2RSE | apo_pdb: N/A | apo_bmrb: 16931 | holo_bmrb: 11471
A.
Apo/Holo HSQC Offset
Apo
Holo
$^{1}$H δ (ppm)
$^{15}$N δ (ppm)
B.
CSP Classification
(TP) Sig. CSP in Binding Site (0)
(FP) Sig. CSP -- Allosteric (45)
(TN) low CSP -- Allosteric (49)
(FN) low CSP in Binding Site (2)
$\Delta\delta_{NH}$ (ppm)
Sequence
C.
CSP Mask
D.
Binding Site Mask
E.
Confusion Matrix Classification

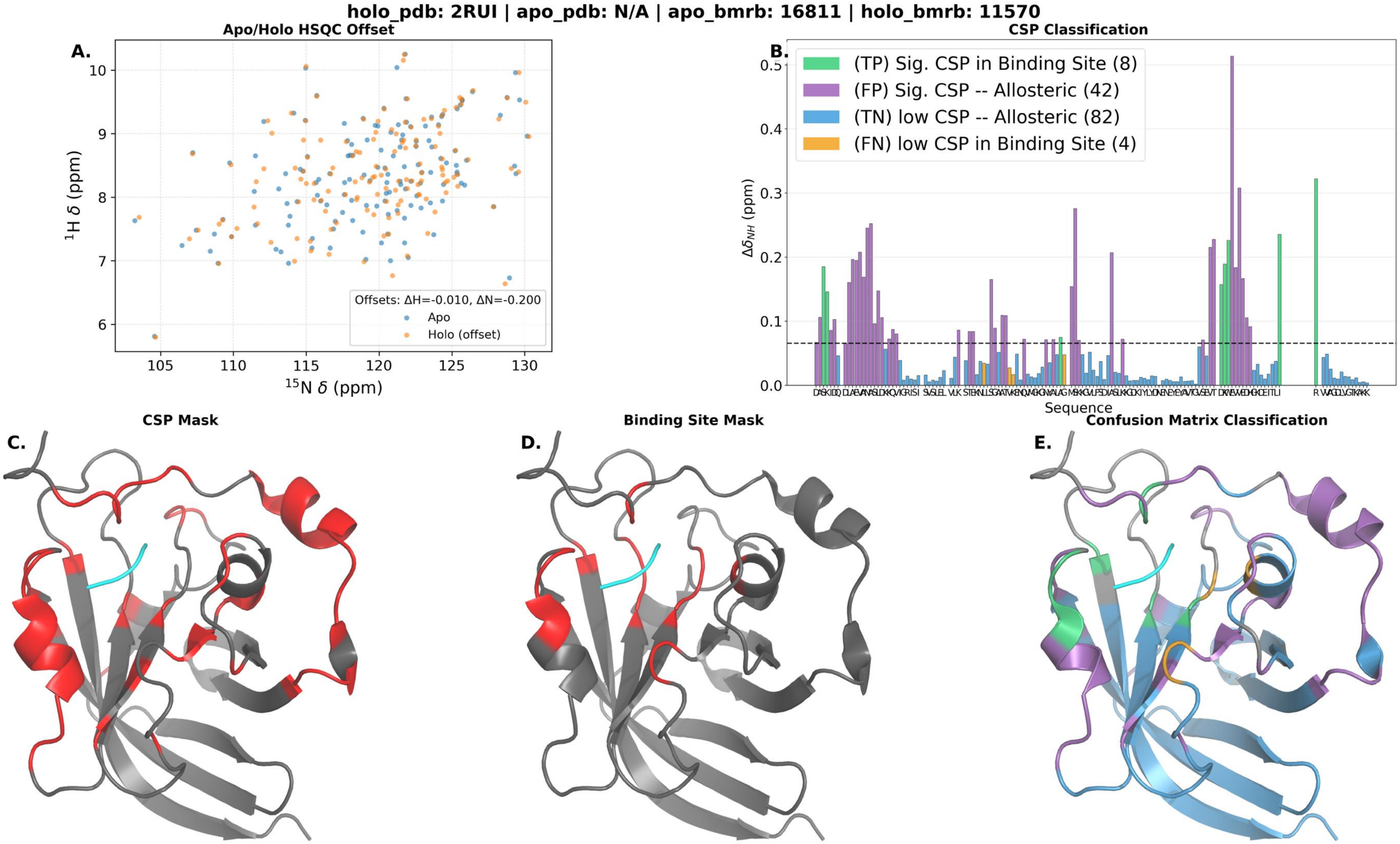

holo_pdb: 2RUI | apo_pdb: N/A | apo_bmrb: 16811 | holo_bmrb: 11570
A.
Apo/Holo HSQC Offset
Offsets: ΔH=-0.010, ΔN=-0.200
Apo
Holo (offset)
$^{1}$H δ (ppm)
$^{15}$N δ (ppm)
B.
CSP Classification
(TP) Sig. CSP in Binding Site (8)
(FP) Sig. CSP -- Allosteric (42)
(TN) low CSP -- Allosteric (82)
(FN) low CSP in Binding Site (4)
$\Delta\delta_{NH}$ (ppm)
Sequence
C.
CSP Mask
D.
Binding Site Mask
E.
Confusion Matrix Classification

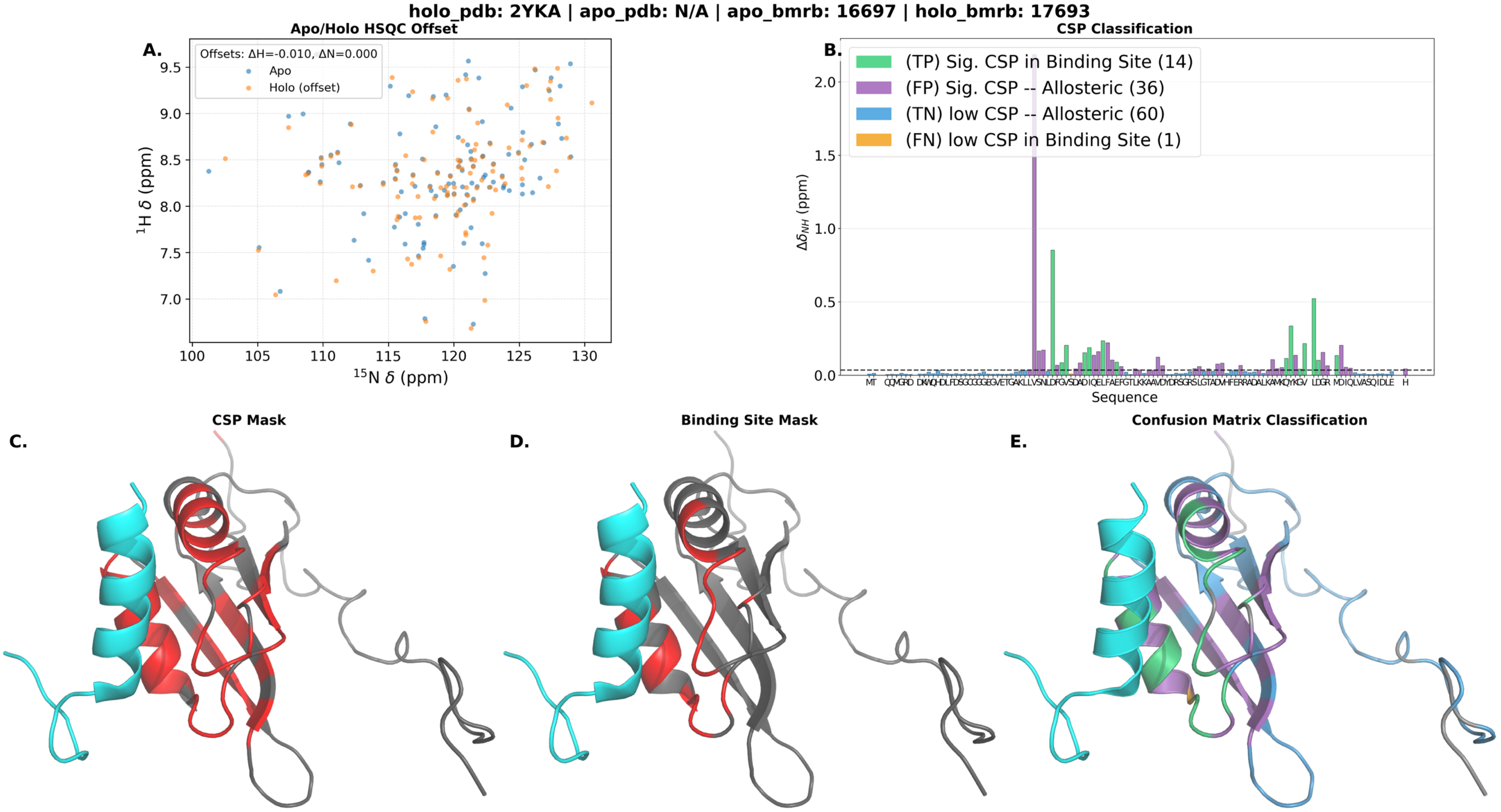
holo_pdb: 2YKA | apo_pdb: N/A | apo_bmrb: 16697 | holo_bmrb: 17693
A.
Apo/Holo HSQC Offset
Offsets: ΔH=-0.010, ΔN=0.000
Apo
Holo (offset)
$^{1}$H δ (ppm)
$^{15}$N δ (ppm)
B.
CSP Classification
(TP) Sig. CSP in Binding Site (14)
(FP) Sig. CSP -- Allosteric (36)
(TN) low CSP -- Allosteric (60)
(FN) low CSP in Binding Site (1)
$\Delta\delta_{NH}$ (ppm)
Sequence
C.
CSP Mask
D.
Binding Site Mask
E.
Confusion Matrix Classification

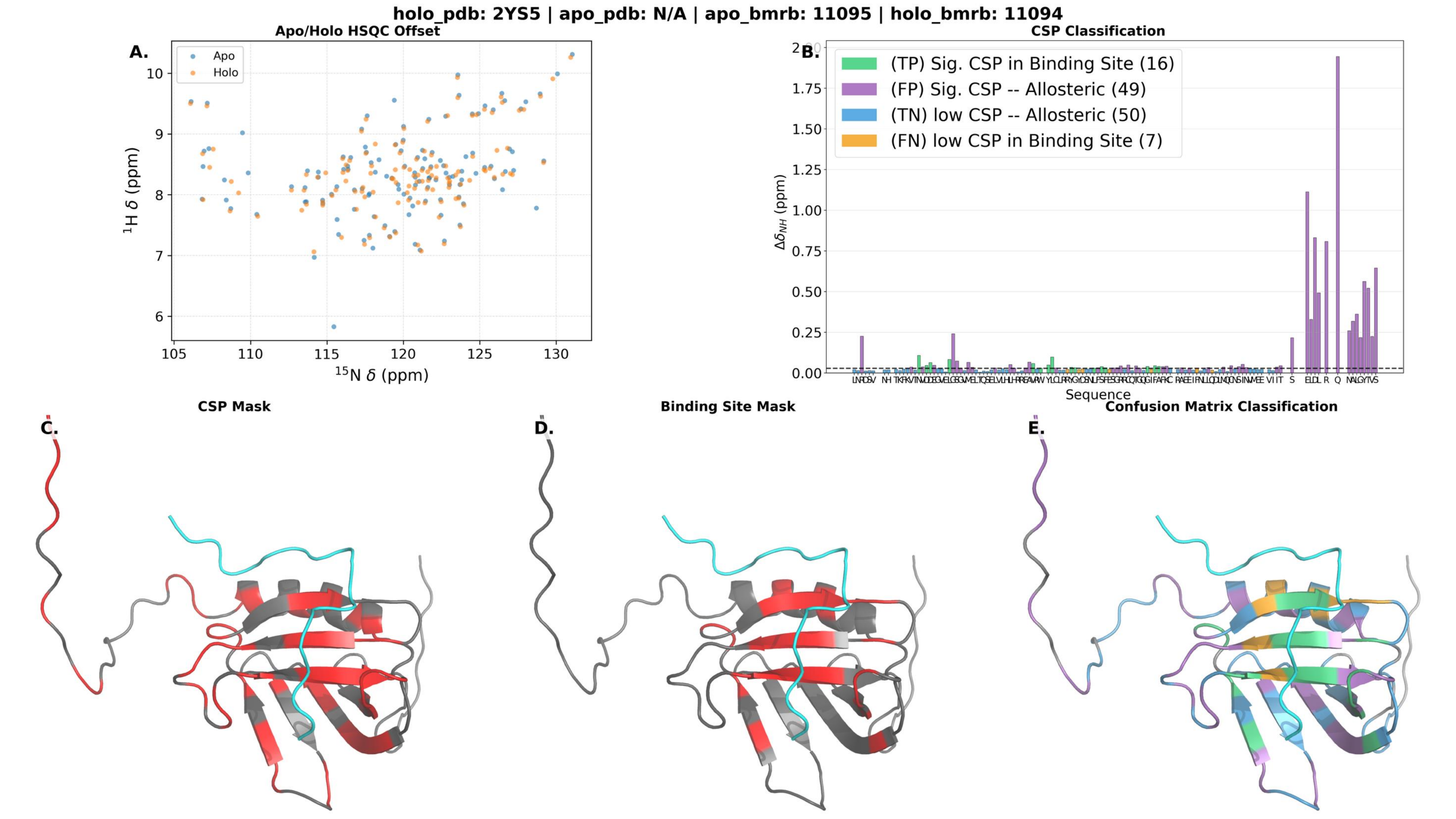

holo_pdb: 2YS5 | apo_pdb: N/A | apo_bmrb: 11095 | holo_bmrb: 11094
A.
Apo/Holo HSQC Offset
Apo
Holo
$^1$H δ (ppm)
$^{15}$N δ (ppm)
B.
CSP Classification
(TP) Sig. CSP in Binding Site (16)
(FP) Sig. CSP -- Allosteric (49)
(TN) low CSP -- Allosteric (50)
(FN) low CSP in Binding Site (7)
$\Delta\delta_{NH}$ (ppm)
Sequence
C.
CSP Mask
D.
Binding Site Mask
E.
Confusion Matrix Classification

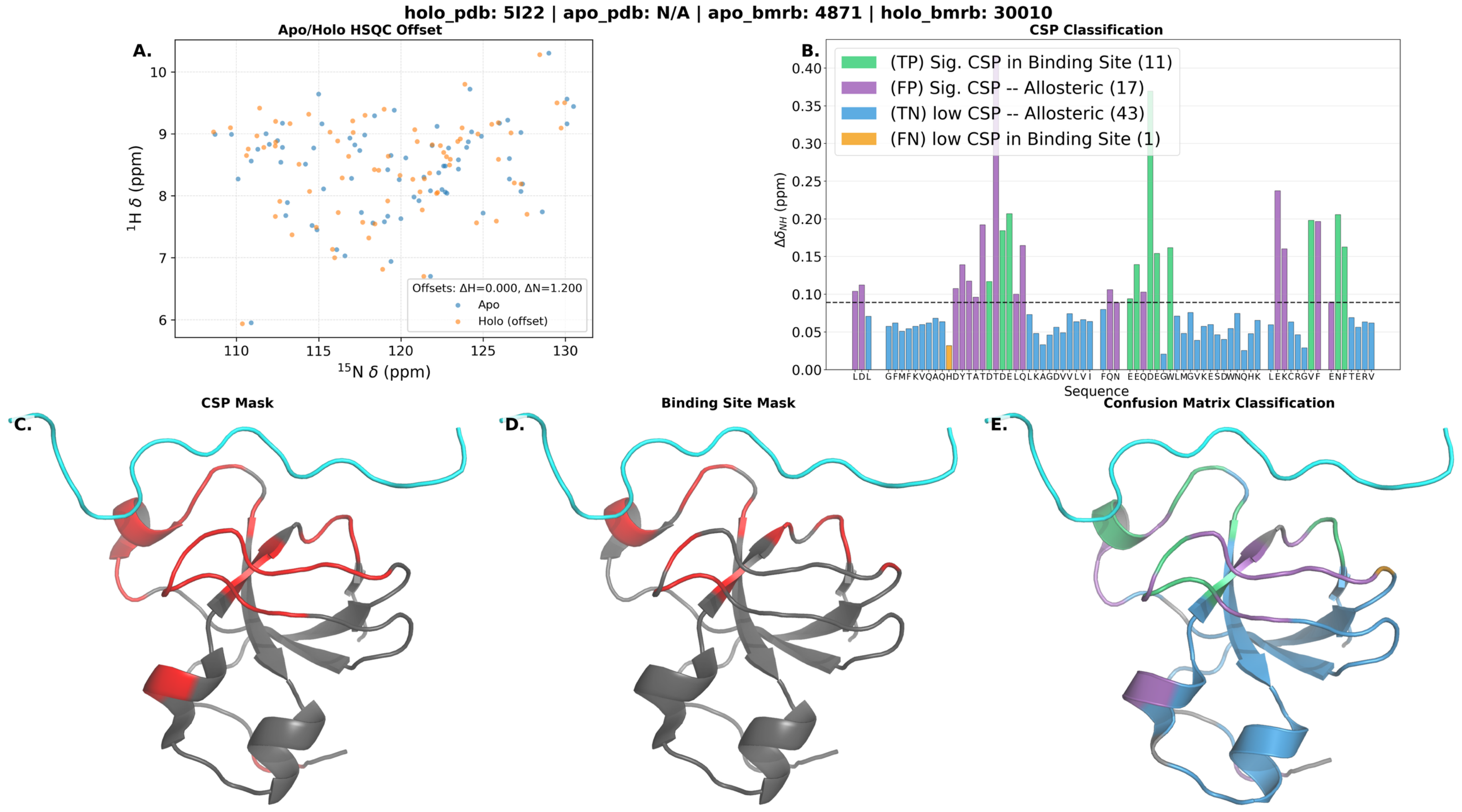
holo_pdb: 5I22 | apo_pdb: N/A | apo_bmrb: 4871 | holo_bmrb: 30010
A.
Apo/Holo HSQC Offset
Offsets: ΔH=0.000, ΔN=1.200
Apo
Holo (offset)
$^{1}$H δ (ppm)
$^{15}$N δ (ppm)
B.
CSP Classification
(TP) Sig. CSP in Binding Site (11)
(FP) Sig. CSP -- Allosteric (17)
(TN) low CSP -- Allosteric (43)
(FN) low CSP in Binding Site (1)
$\Delta\delta_{NH}$ (ppm)
Sequence
C.
CSP Mask
D.
Binding Site Mask
E.
Confusion Matrix Classification

**holo_pdb: 5J8H | apo_pdb: N/A | apo_bmrb: 51289 | holo_bmrb: 30063**

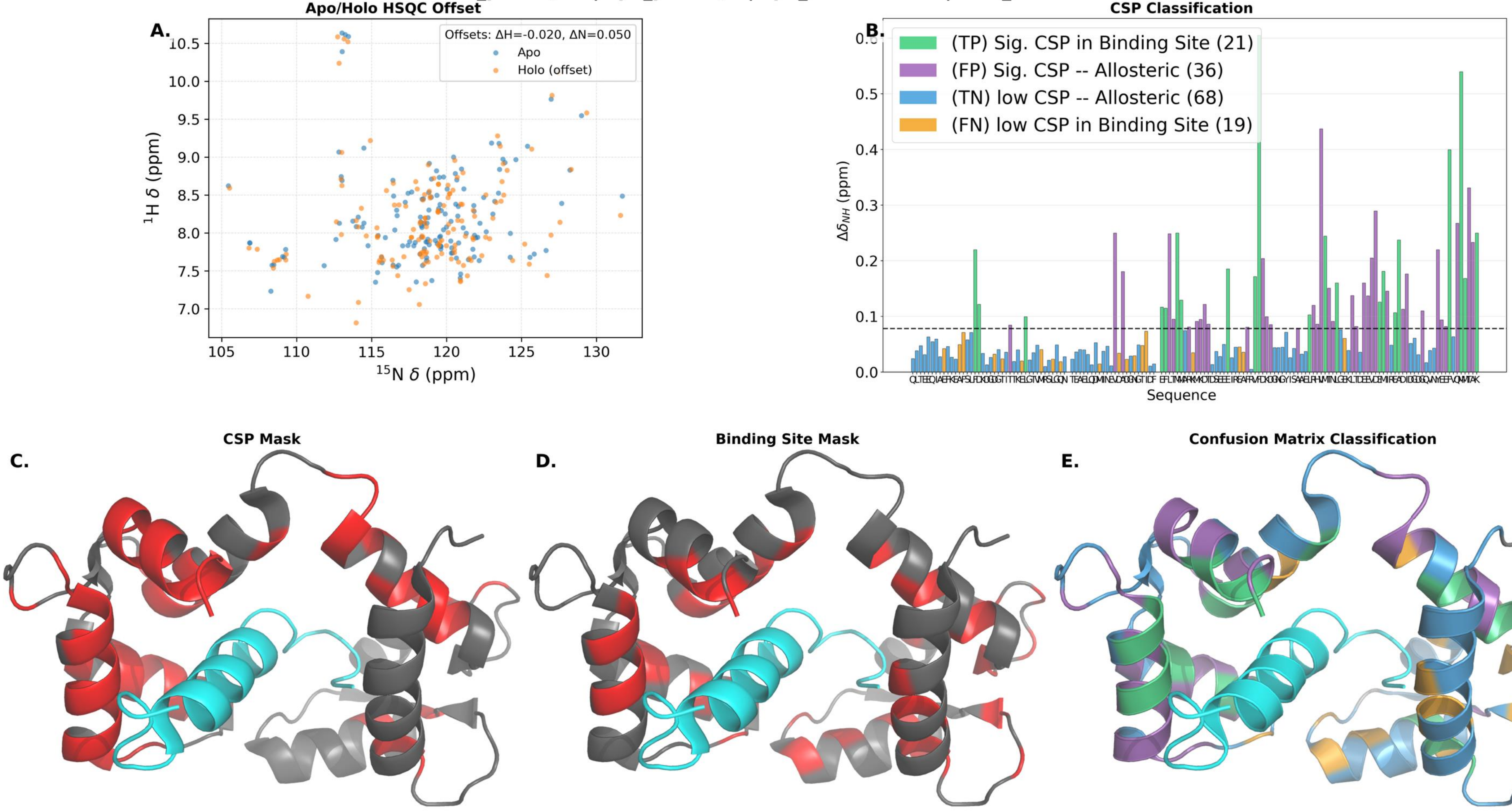

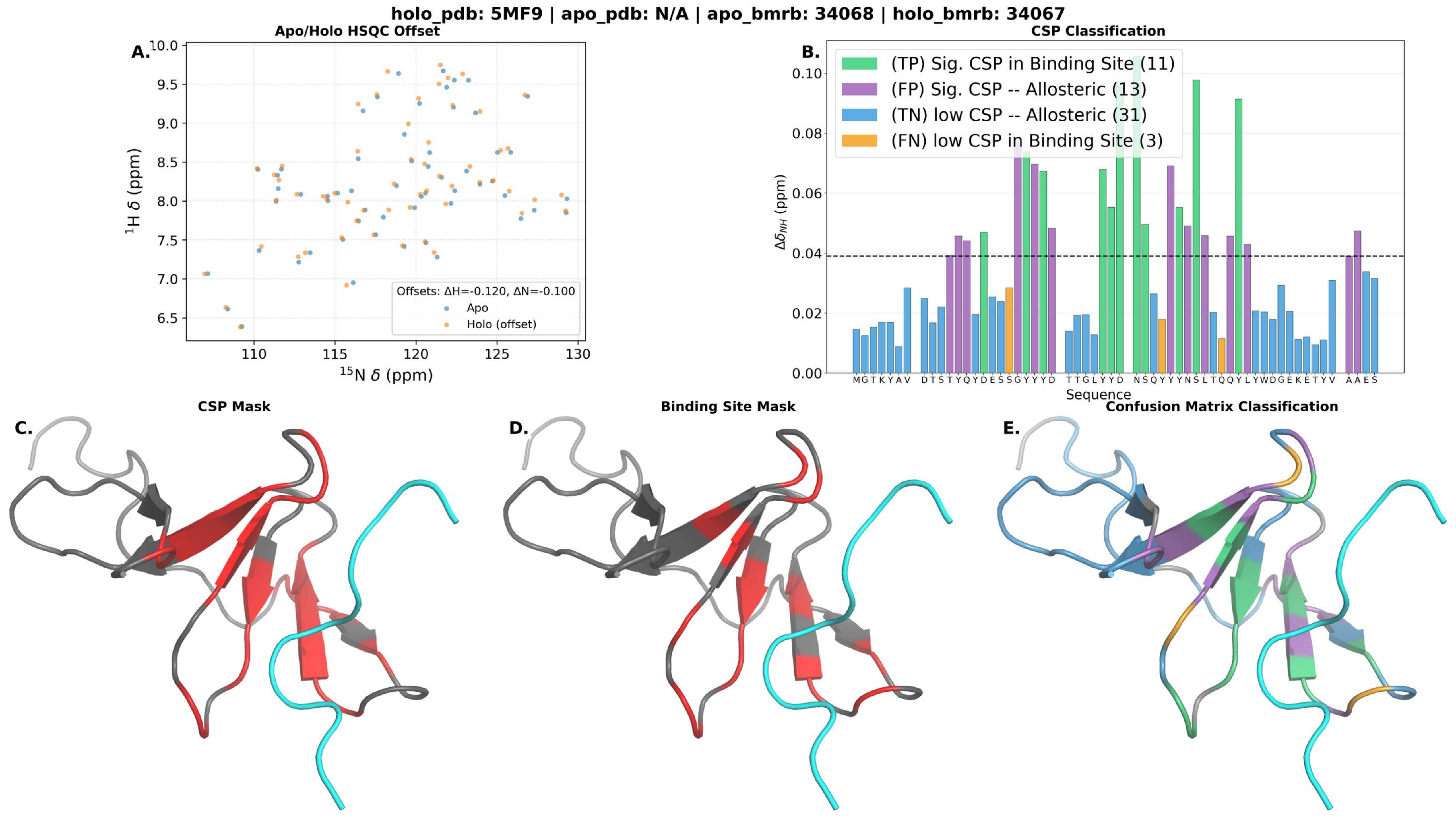

holo_pdb: 5MF9 | apo_pdb: N/A | apo_bmrb: 34068 | holo_bmrb: 34067
A.
Apo/Holo HSQC Offset
10.0
9.5
9.0
8.5
8.0
7.5
7.0
6.5
$^{1}$H $\delta$ (ppm)
110
115
120
125
130
$^{15}$N $\delta$ (ppm)
Offsets: ΔH=-0.120, ΔN=-0.100
Apo
Holo (offset)
B.
CSP Classification
(TP) Sig. CSP in Binding Site (11)
(FP) Sig. CSP -- Allosteric (13)
(TN) low CSP -- Allosteric (31)
(FN) low CSP in Binding Site (3)
0.10
0.08
0.06
0.04
0.02
0.00
$\Delta\delta_{NH}$ (ppm)
M G T K Y A V D T S T Y Q Y D E S S G Y Y Y D T T G L Y Y D N S Q Y Y Y N S L T Q Q Y L Y W D G E K E T Y V A A E S
Sequence
C.
CSP Mask
D.
Binding Site Mask
E.
Confusion Matrix Classification

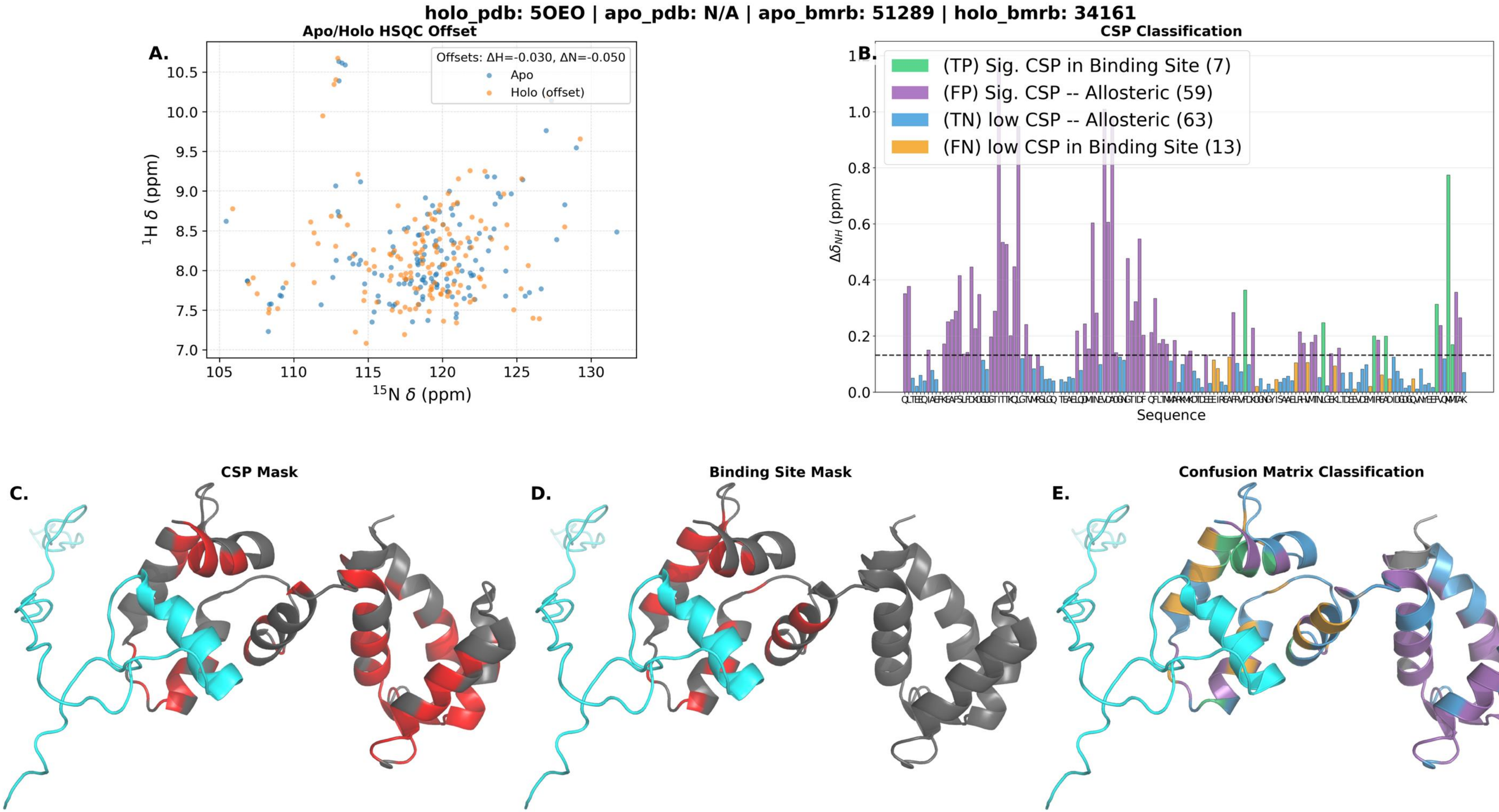

holo_pdb: 5OEO | apo_pdb: N/A | apo_bmrb: 51289 | holo_bmrb: 34161
A.
Apo/Holo HSQC Offset
Offsets: ΔH=-0.030, ΔN=-0.050
Apo
Holo (offset)
$^{1}$H δ (ppm)
$^{15}$N δ (ppm)
B.
CSP Classification
(TP) Sig. CSP in Binding Site (7)
(FP) Sig. CSP -- Allosteric (59)
(TN) low CSP -- Allosteric (63)
(FN) low CSP in Binding Site (13)
$\Delta\delta_{NH}$ (ppm)
Sequence
C.
CSP Mask
D.
Binding Site Mask
E.
Confusion Matrix Classification

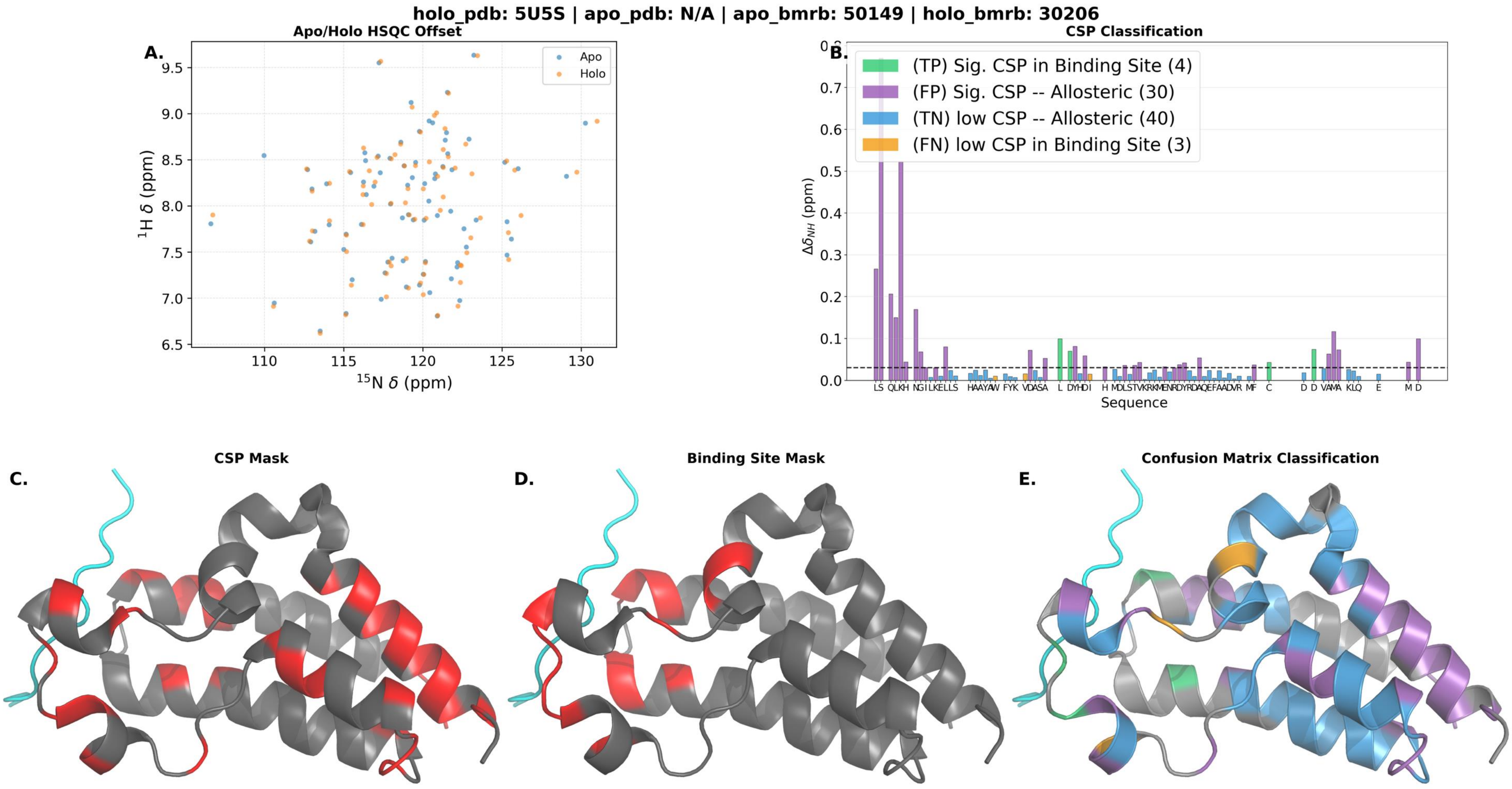
holo_pdb: 5U5S | apo_pdb: N/A | apo_bmrb: 50149 | holo_bmrb: 30206
A.
Apo/Holo HSQC Offset
Apo
Holo
$^{1}$H δ (ppm)
$^{15}$N δ (ppm)
B.
CSP Classification
(TP) Sig. CSP in Binding Site (4)
(FP) Sig. CSP -- Allosteric (30)
(TN) low CSP -- Allosteric (40)
(FN) low CSP in Binding Site (3)
$\Delta\delta_{NH}$ (ppm)
Sequence
C.
CSP Mask
D.
Binding Site Mask
E.
Confusion Matrix Classification

holo_pdb: 5VF0 | apo_pdb: N/A | apo_bmrb: 17769 | holo_bmrb: 30276

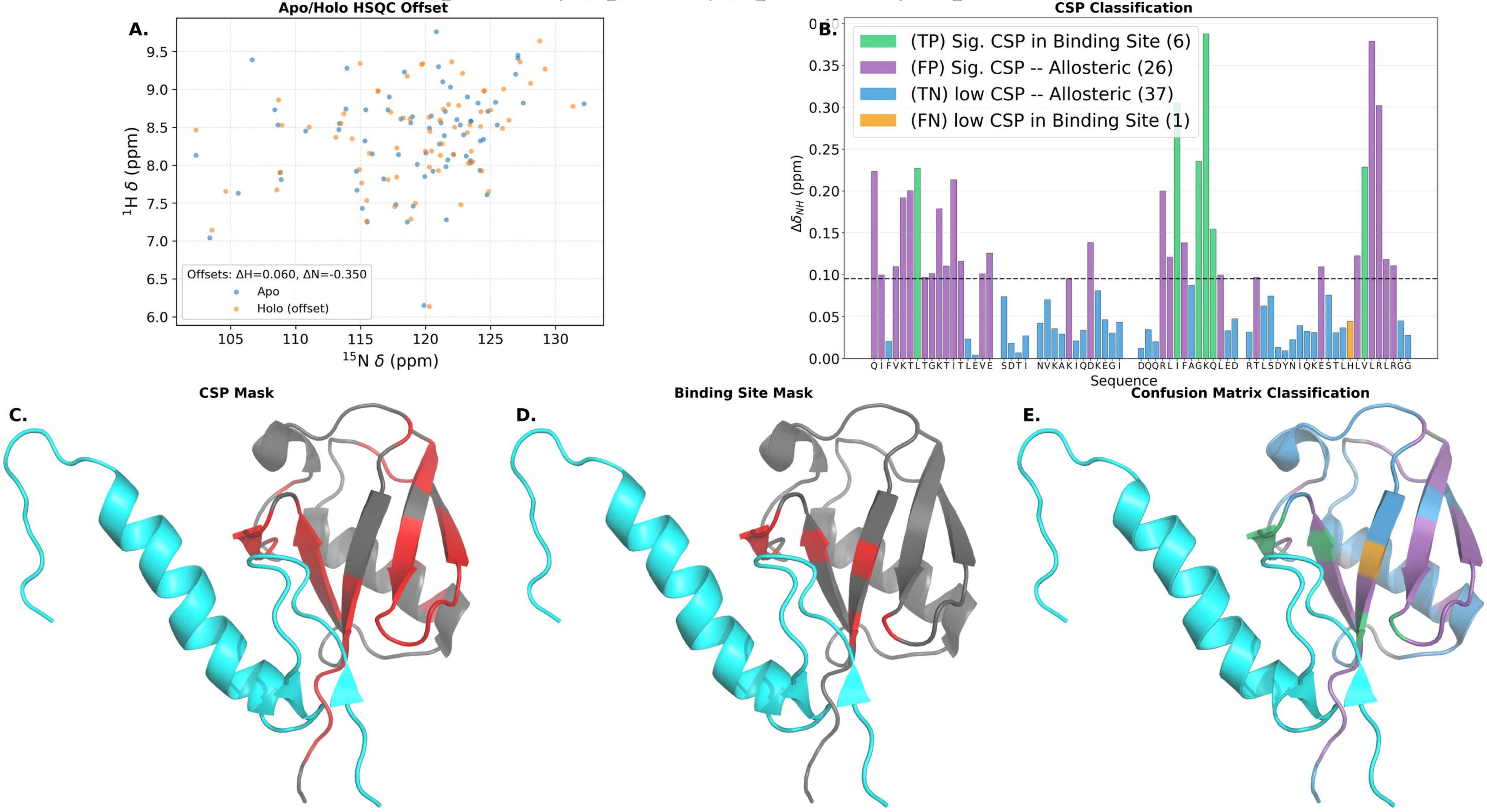

holo_pdb: 6BGG | apo_pdb: 2JNS | apo_bmrb: 15125 | holo_bmrb: 30367

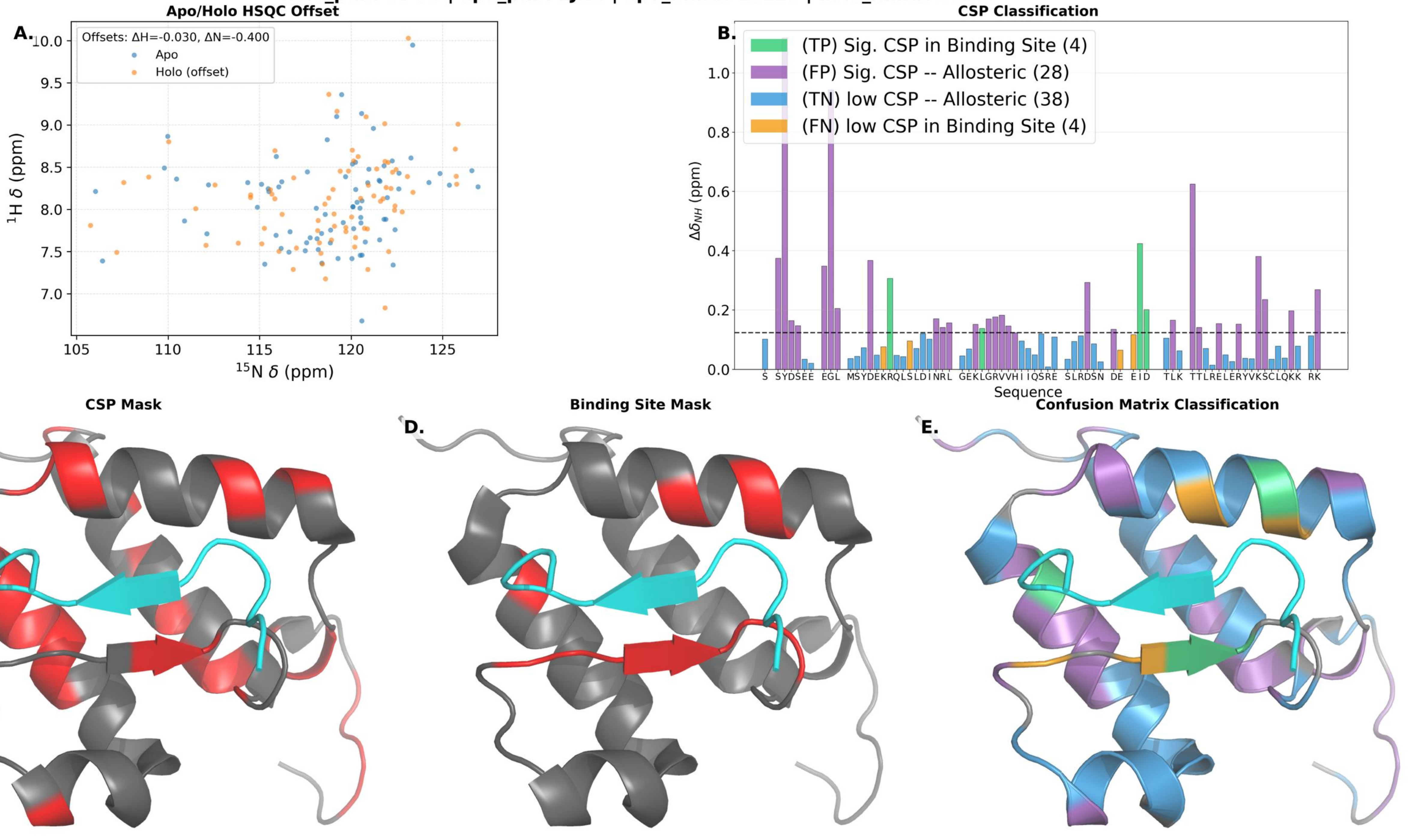

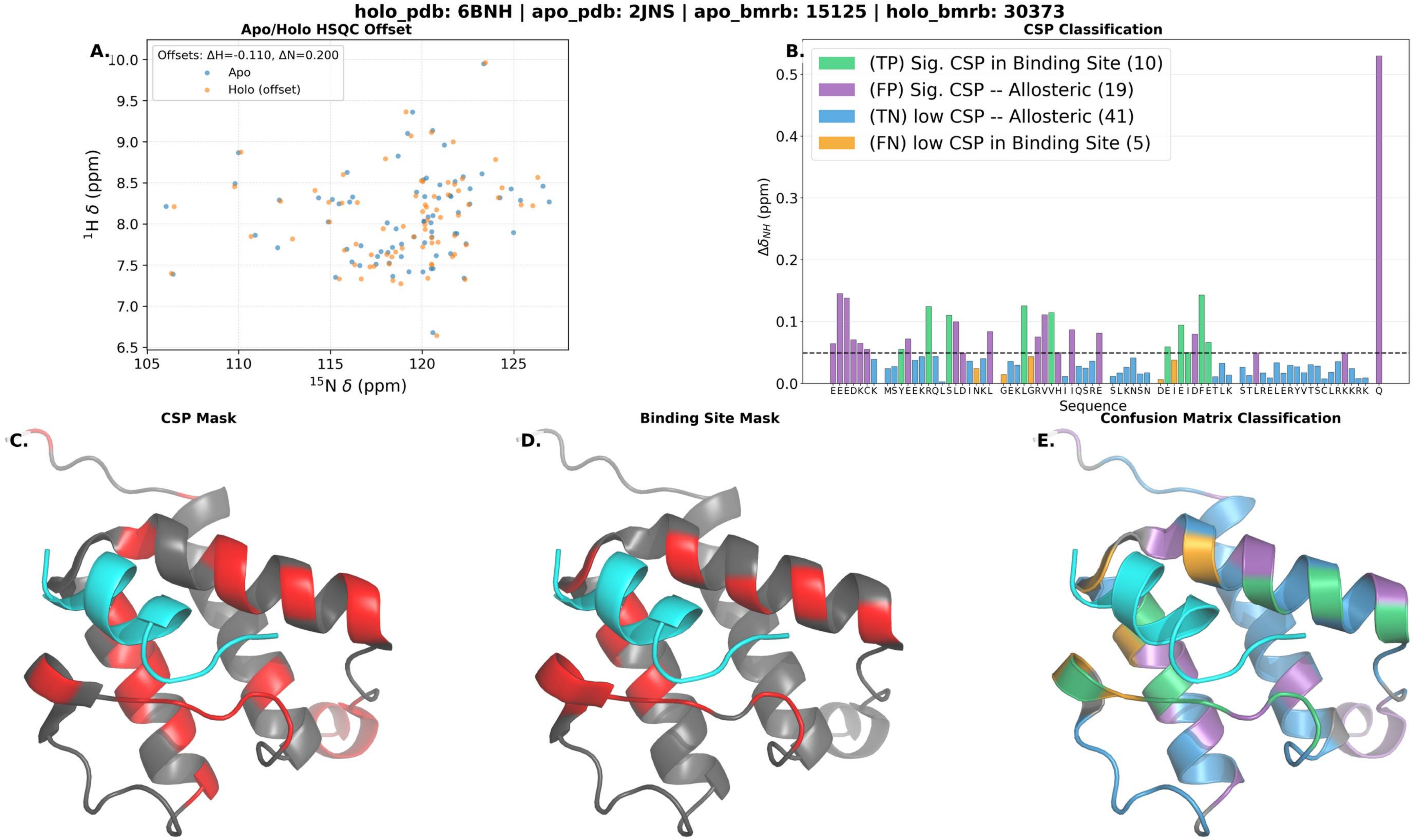
holo_pdb: 6BNH | apo_pdb: 2JNS | apo_bmrb: 15125 | holo_bmrb: 30373
Apo/Holo HSQC Offset
A.
Offsets: ΔH=-0.110, ΔN=0.200
Apo
Holo (offset)
¹H δ (ppm)
¹⁵N δ (ppm)
CSP Classification
B.
(TP) Sig. CSP in Binding Site (10)
(FP) Sig. CSP -- Allosteric (19)
(TN) low CSP -- Allosteric (41)
(FN) low CSP in Binding Site (5)
Δδ_NH (ppm)
Sequence
CSP Mask
C.
Binding Site Mask
D.
Confusion Matrix Classification
E.

**holo_pdb: 6E5N | apo_pdb: N/A | apo_bmrb: 25544 | holo_bmrb: 30500**

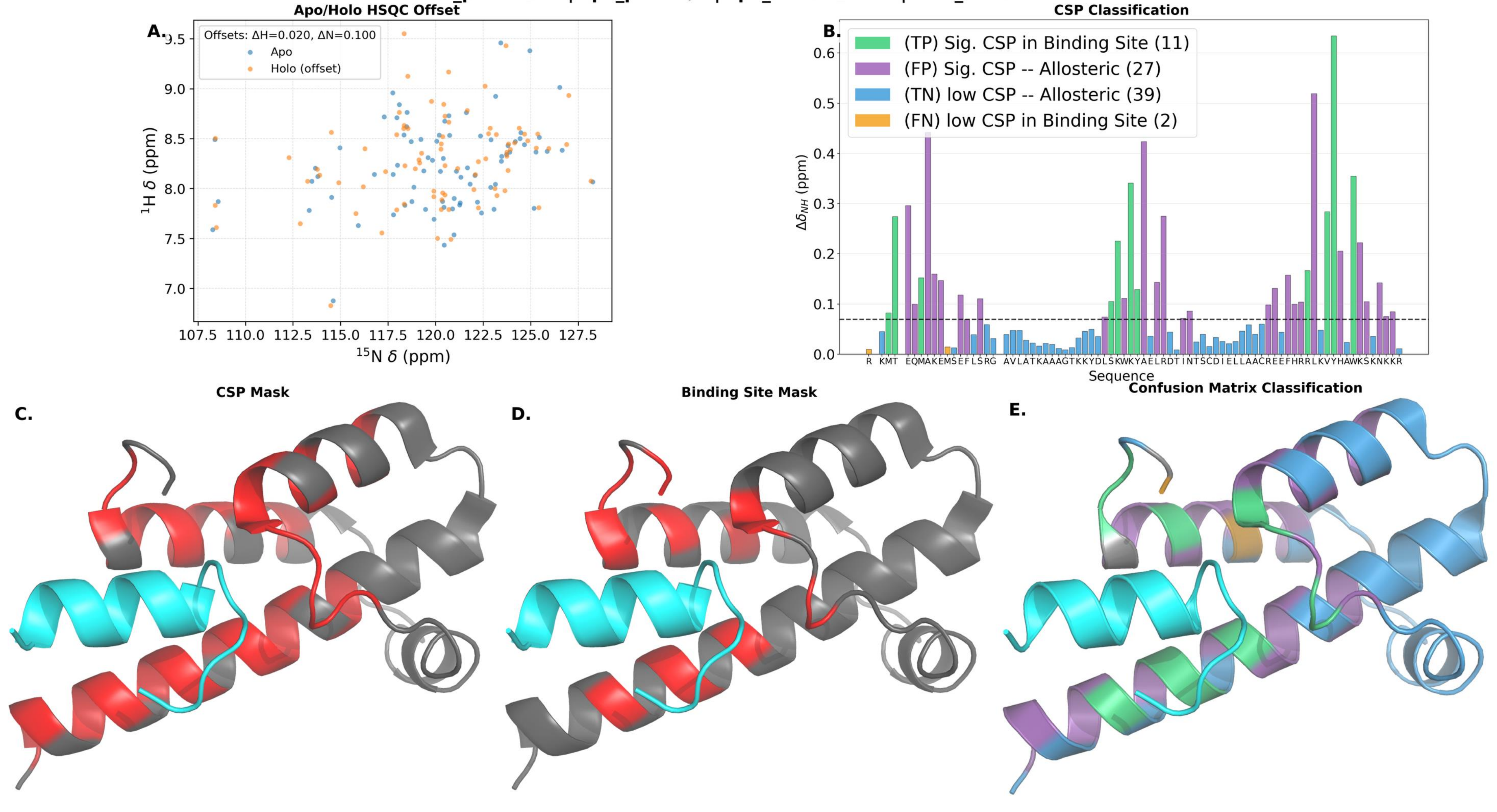

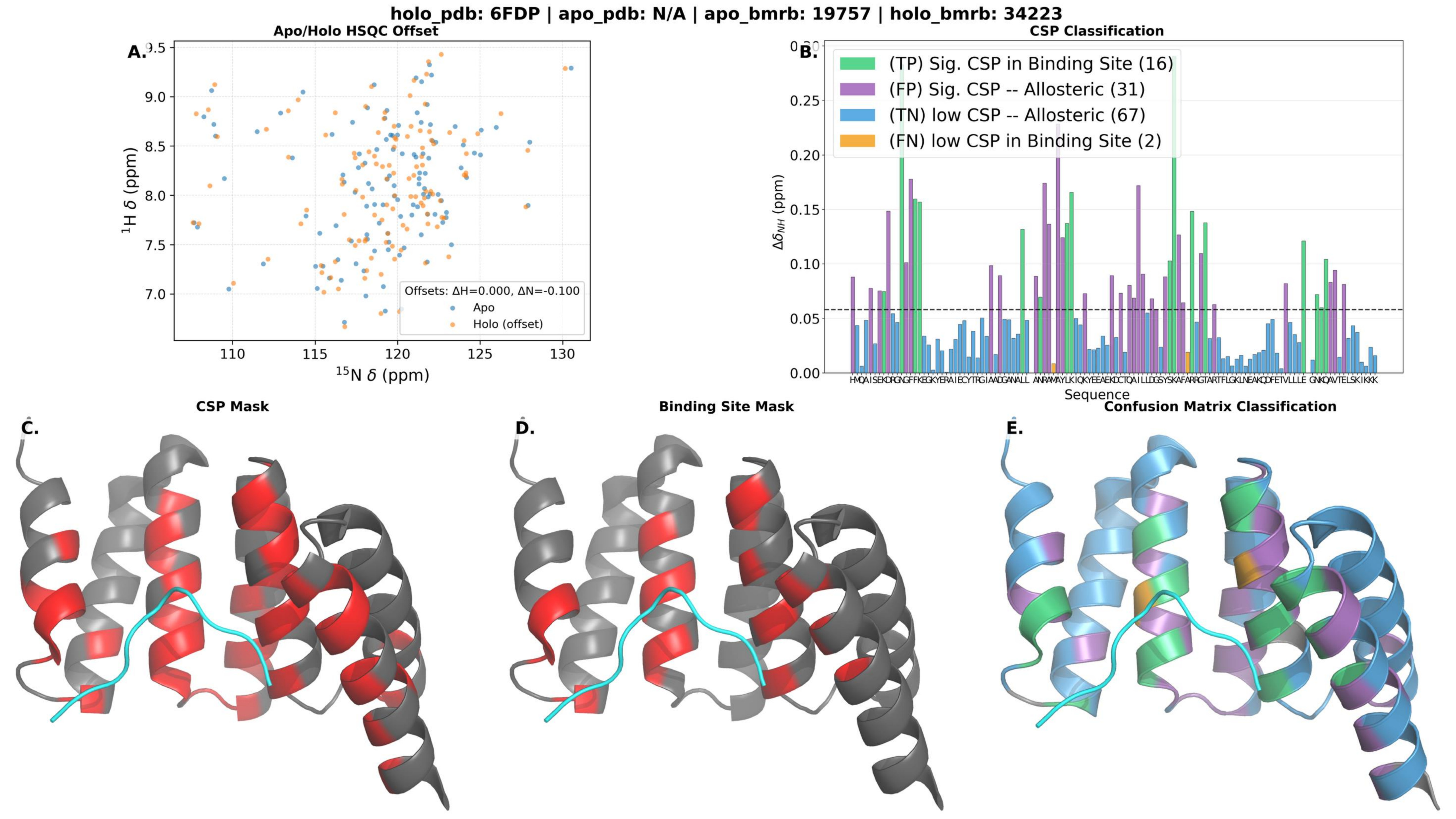

holo_pdb: 6FDP | apo_pdb: N/A | apo_bmrb: 19757 | holo_bmrb: 34223
A.
Apo/Holo HSQC Offset
$^{1}$H δ (ppm)
$^{15}$N δ (ppm)
Offsets: ΔH=0.000, ΔN=-0.100
Apo
Holo (offset)
B.
CSP Classification
(TP) Sig. CSP in Binding Site (16)
(FP) Sig. CSP -- Allosteric (31)
(TN) low CSP -- Allosteric (67)
(FN) low CSP in Binding Site (2)
$\Delta\delta_{NH}$ (ppm)
Sequence
C.
CSP Mask
D.
Binding Site Mask
E.
Confusion Matrix Classification

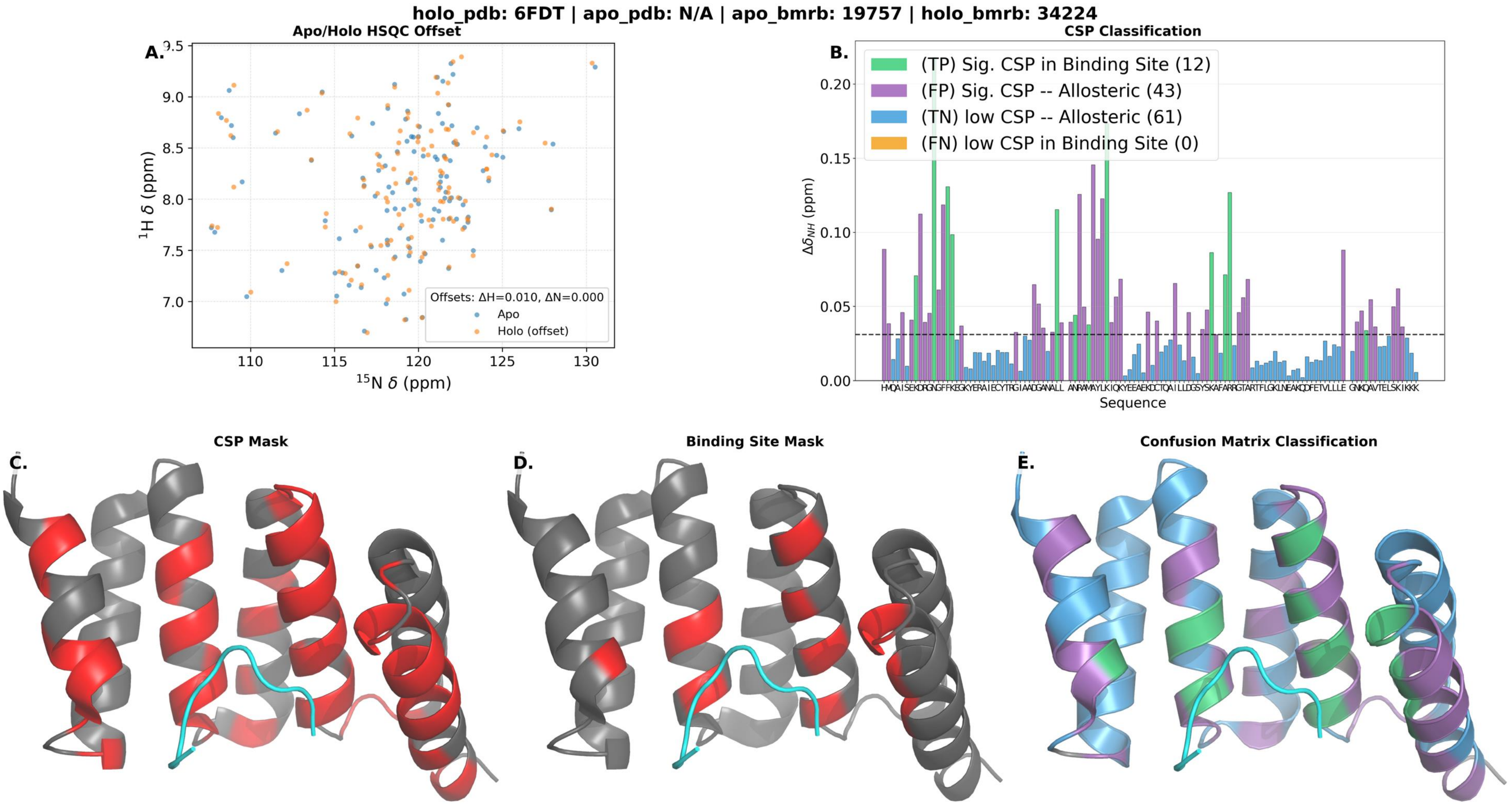

holo_pdb: 6FDT | apo_pdb: N/A | apo_bmrb: 19757 | holo_bmrb: 34224
A.
Apo/Holo HSQC Offset
1H δ (ppm)
15N δ (ppm)
Offsets: ΔH=0.010, ΔN=0.000
Apo
Holo (offset)
B.
CSP Classification
(TP) Sig. CSP in Binding Site (12)
(FP) Sig. CSP -- Allosteric (43)
(TN) low CSP -- Allosteric (61)
(FN) low CSP in Binding Site (0)
ΔδNH (ppm)
Sequence
C.
CSP Mask
D.
Binding Site Mask
E.
Confusion Matrix Classification

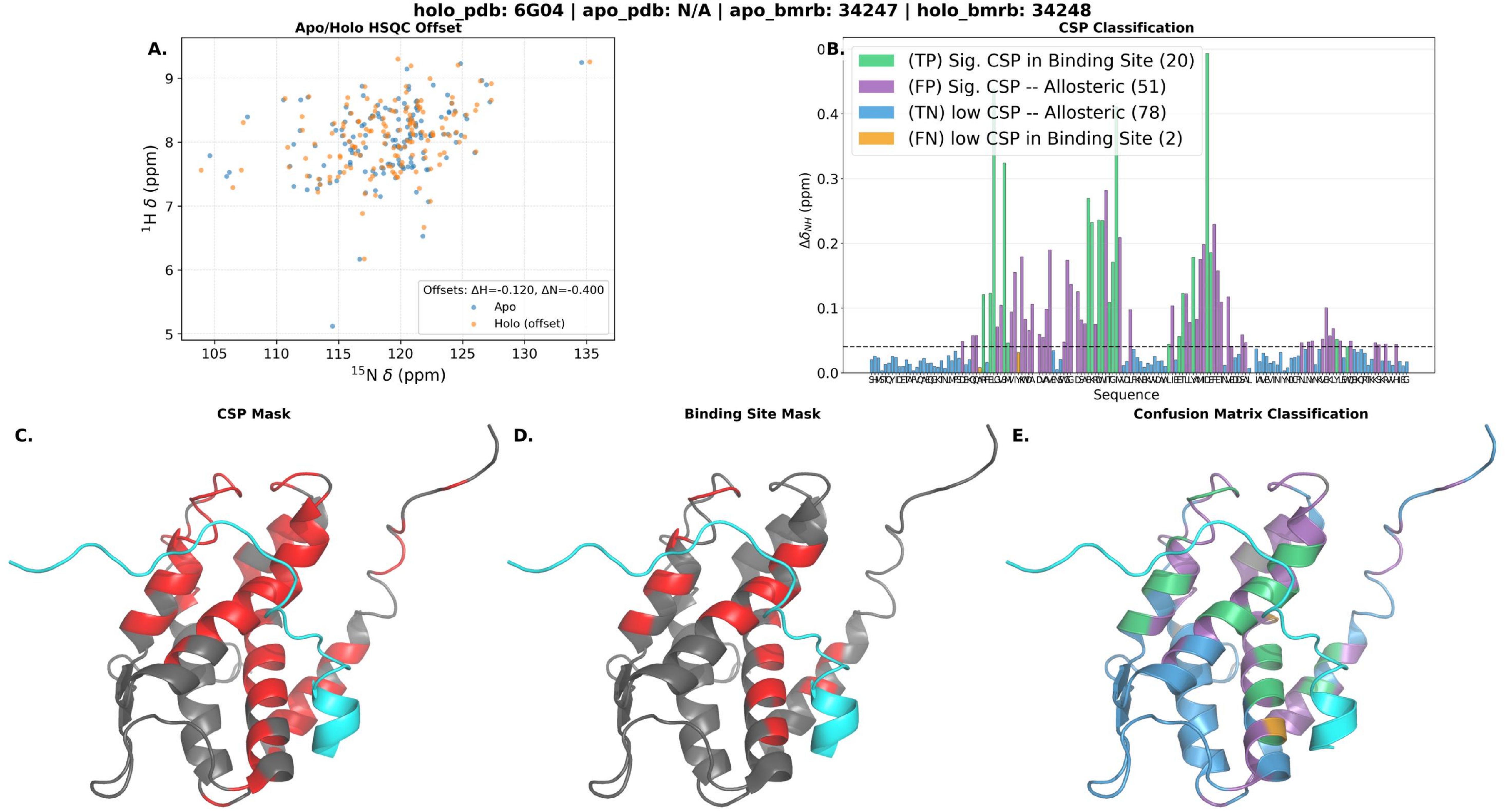

holo_pdb: 6G04 | apo_pdb: N/A | apo_bmrb: 34247 | holo_bmrb: 34248
A.
Apo/Holo HSQC Offset
Offsets: ΔH=-0.120, ΔN=-0.400
Apo
Holo (offset)
$^{1}$H δ (ppm)
$^{15}$N δ (ppm)
B.
CSP Classification
(TP) Sig. CSP in Binding Site (20)
(FP) Sig. CSP -- Allosteric (51)
(TN) low CSP -- Allosteric (78)
(FN) low CSP in Binding Site (2)
$\Delta\delta_{NH}$ (ppm)
Sequence
C.
CSP Mask
D.
Binding Site Mask
E.
Confusion Matrix Classification

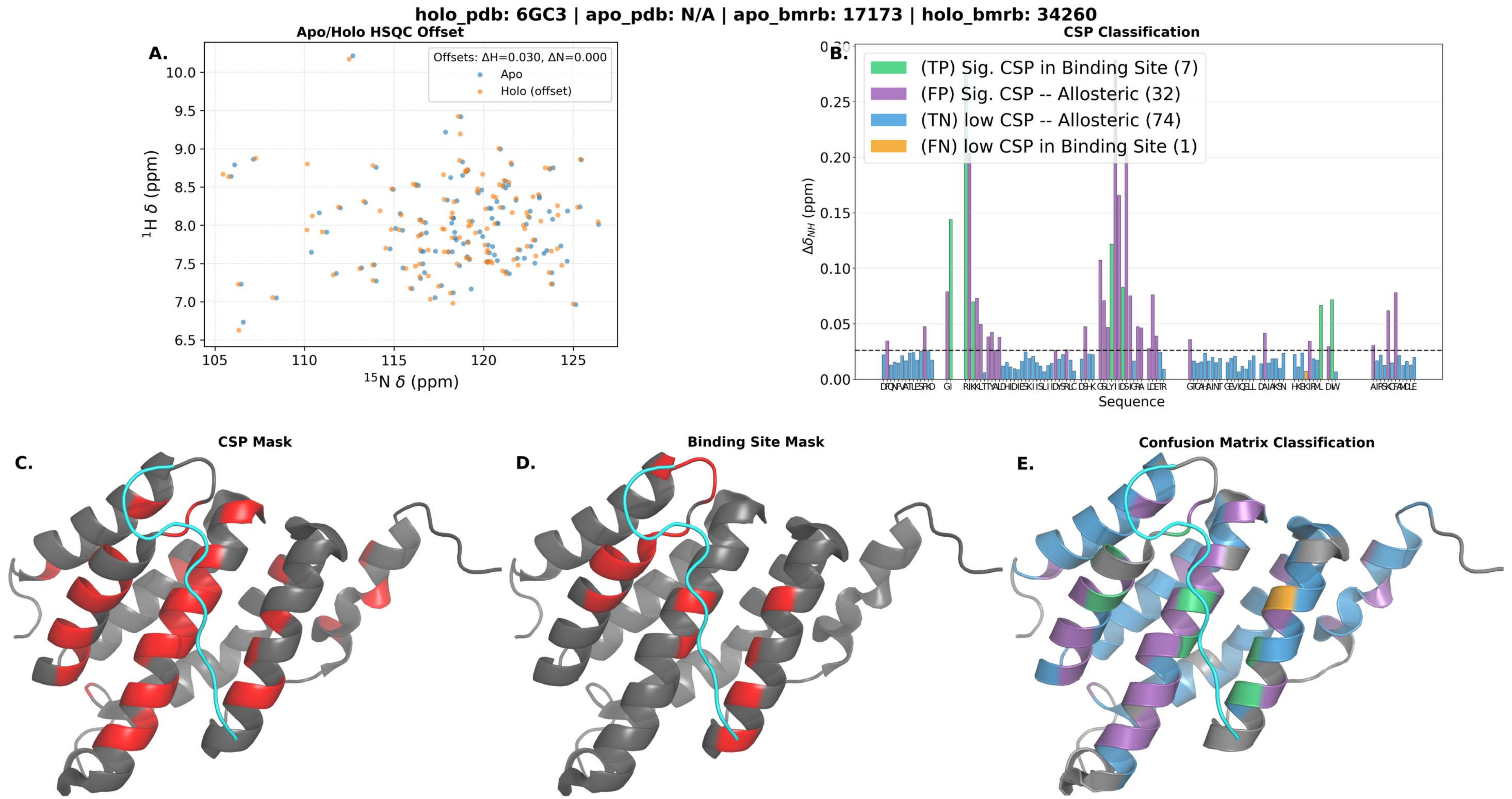

holo_pdb: 6GC3 | apo_pdb: N/A | apo_bmrb: 17173 | holo_bmrb: 34260
A.
Apo/Holo HSQC Offset
Offsets: ΔH=0.030, ΔN=0.000
Apo
Holo (offset)
¹H δ (ppm)
¹⁵N δ (ppm)
B.
CSP Classification
(TP) Sig. CSP in Binding Site (7)
(FP) Sig. CSP -- Allosteric (32)
(TN) low CSP -- Allosteric (74)
(FN) low CSP in Binding Site (1)
Δδ_NH (ppm)
Sequence
C.
CSP Mask
D.
Binding Site Mask
E.
Confusion Matrix Classification

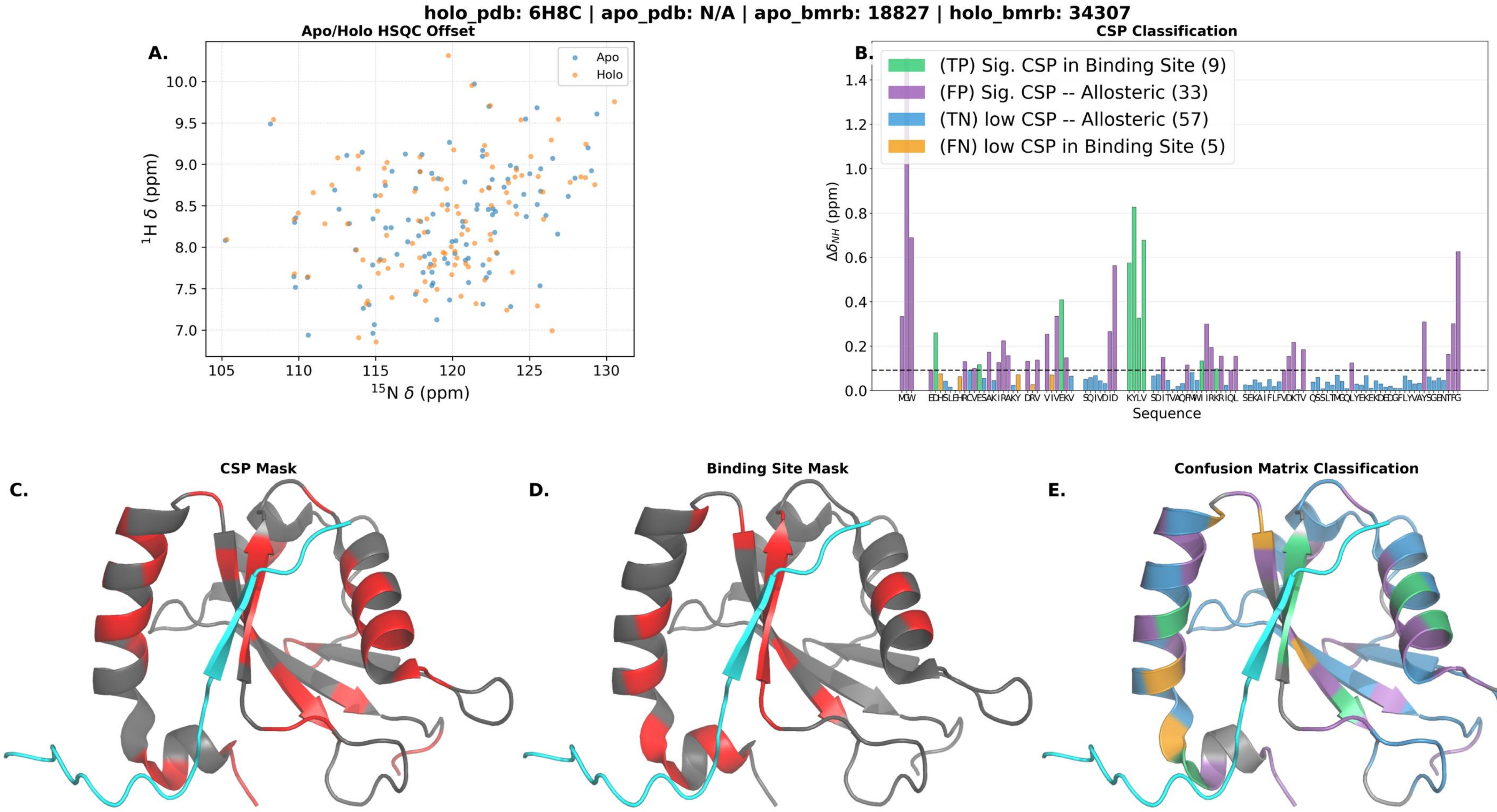
holo_pdb: 6H8C | apo_pdb: N/A | apo_bmrb: 18827 | holo_bmrb: 34307
A.
Apo/Holo HSQC Offset
Apo
Holo
$^{1}$H δ (ppm)
$^{15}$N δ (ppm)
B.
CSP Classification
(TP) Sig. CSP in Binding Site (9)
(FP) Sig. CSP -- Allosteric (33)
(TN) low CSP -- Allosteric (57)
(FN) low CSP in Binding Site (5)
$\Delta\delta_{NH}$ (ppm)
Sequence
C.
CSP Mask
D.
Binding Site Mask
E.
Confusion Matrix Classification

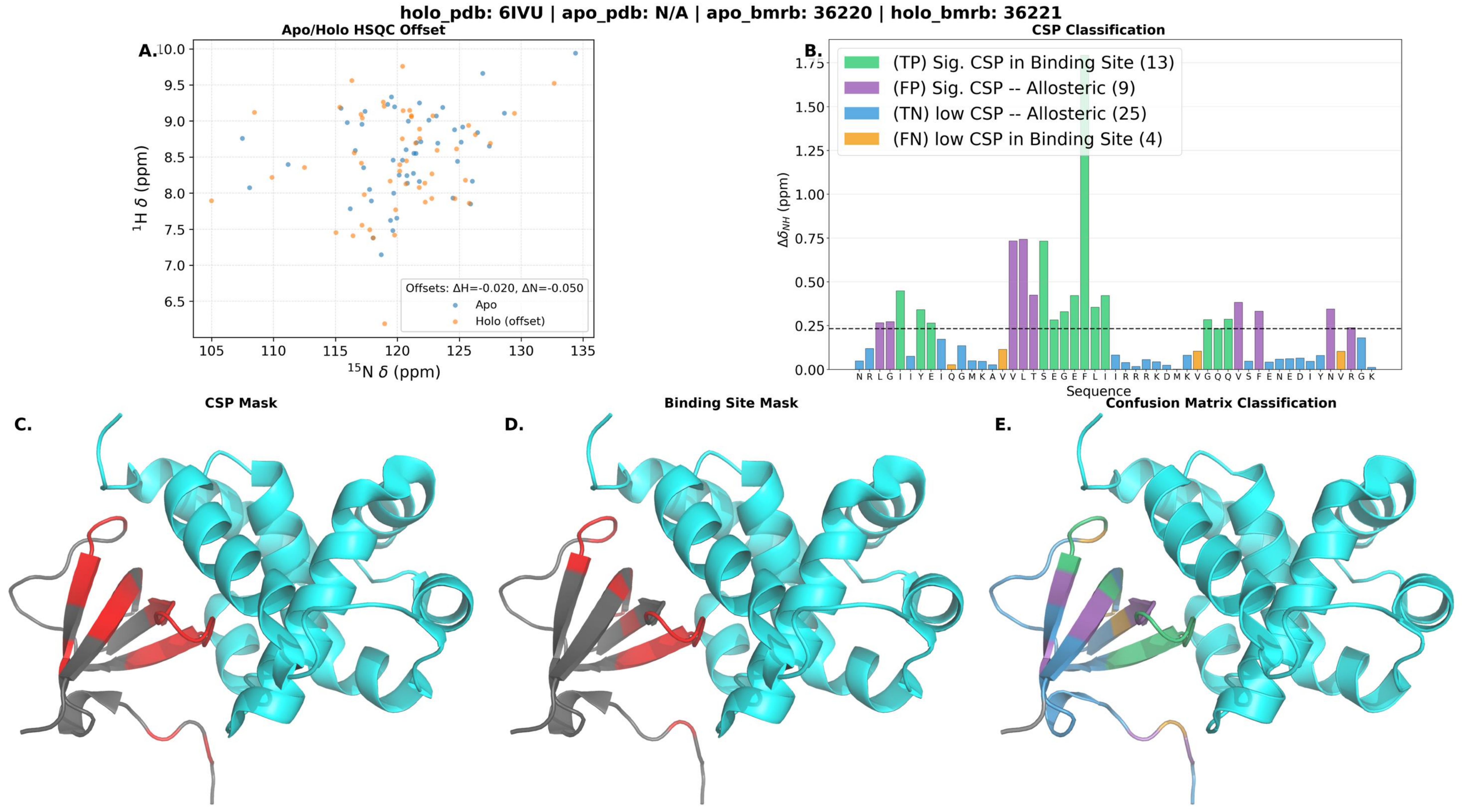

holo_pdb: 6IVU | apo_pdb: N/A | apo_bmrb: 36220 | holo_bmrb: 36221
Apo/Holo HSQC Offset
A.
Offsets: ΔH=-0.020, ΔN=-0.050
Apo
Holo (offset)
$^{1}$H δ (ppm)
$^{15}$N δ (ppm)
CSP Classification
B.
(TP) Sig. CSP in Binding Site (13)
(FP) Sig. CSP -- Allosteric (9)
(TN) low CSP -- Allosteric (25)
(FN) low CSP in Binding Site (4)
$\Delta\delta_{NH}$ (ppm)
N R L G I I Y E I Q G M K A V V L T S E G E F L I I R R R K D M K V G Q Q V S F E N E D I Y N V R G K
Sequence
CSP Mask
C.
Binding Site Mask
D.
Confusion Matrix Classification
E.

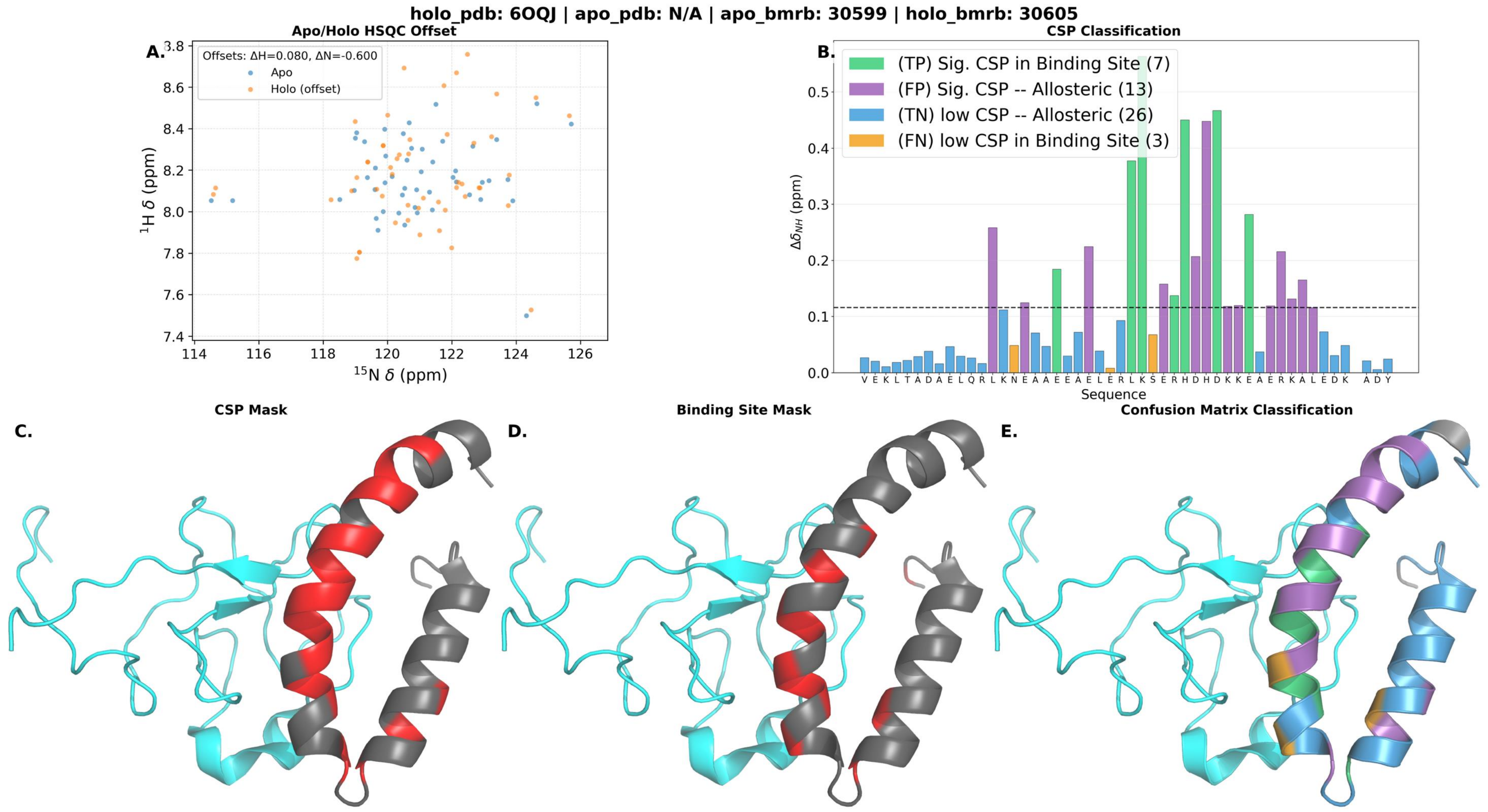

holo_pdb: 6OQJ | apo_pdb: N/A | apo_bmrb: 30599 | holo_bmrb: 30605
Apo/Holo HSQC Offset
A.
Offsets: ΔH=0.080, ΔN=-0.600
Apo
Holo (offset)
$^{1}$H δ (ppm)
$^{15}$N δ (ppm)
CSP Classification
B.
(TP) Sig. CSP in Binding Site (7)
(FP) Sig. CSP -- Allosteric (13)
(TN) low CSP -- Allosteric (26)
(FN) low CSP in Binding Site (3)
$\Delta\delta_{NH}$ (ppm)
Sequence
V E K L T A D A E L Q R L K N E A A E E A E L E R L K S E R H D H D K K E A E R K A L E D K A D Y
CSP Mask
C.
Binding Site Mask
D.
Confusion Matrix Classification
E.

### holo_pdb: 6QXZ | apo_pdb: N/A | apo_bmrb: 27250 | holo_bmrb: 27251

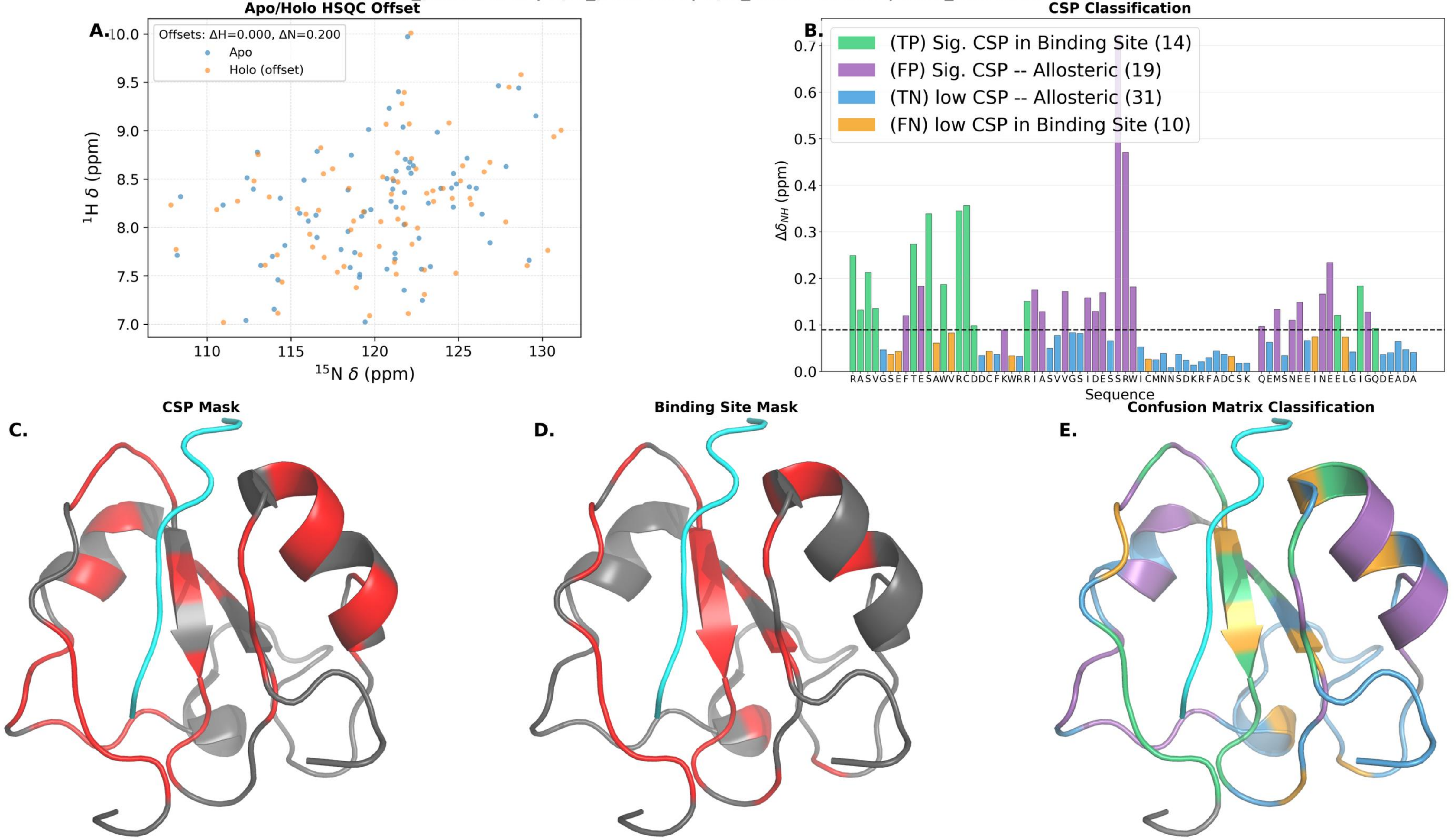

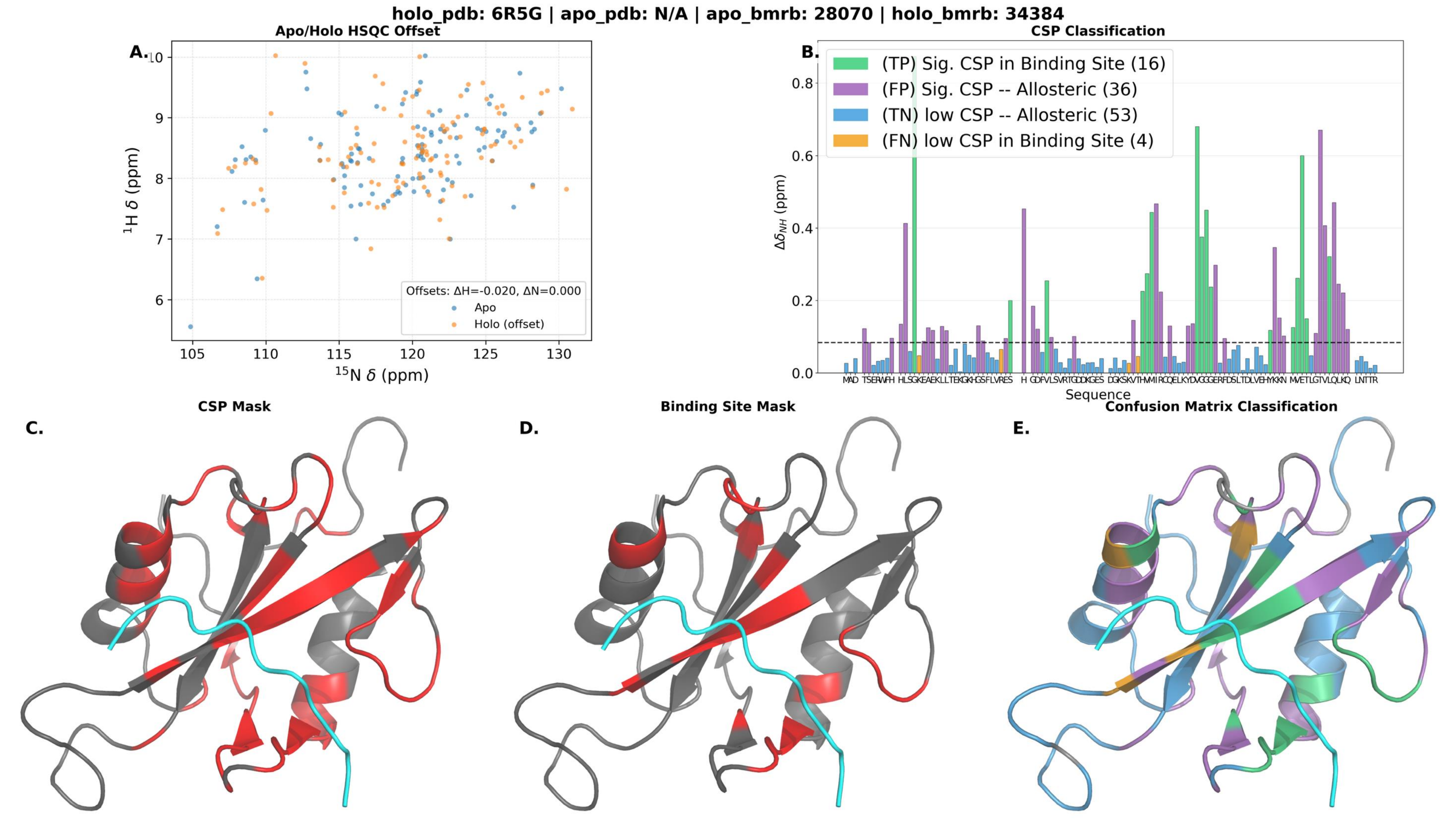
holo_pdb: 6R5G | apo_pdb: N/A | apo_bmrb: 28070 | holo_bmrb: 34384
A.
Apo/Holo HSQC Offset
$^{1}$H δ (ppm)
$^{15}$N δ (ppm)
Offsets: ΔH=-0.020, ΔN=0.000
Apo
Holo (offset)
B.
CSP Classification
(TP) Sig. CSP in Binding Site (16)
(FP) Sig. CSP -- Allosteric (36)
(TN) low CSP -- Allosteric (53)
(FN) low CSP in Binding Site (4)
$\Delta\delta_{NH}$ (ppm)
Sequence
C.
CSP Mask
D.
Binding Site Mask
E.
Confusion Matrix Classification

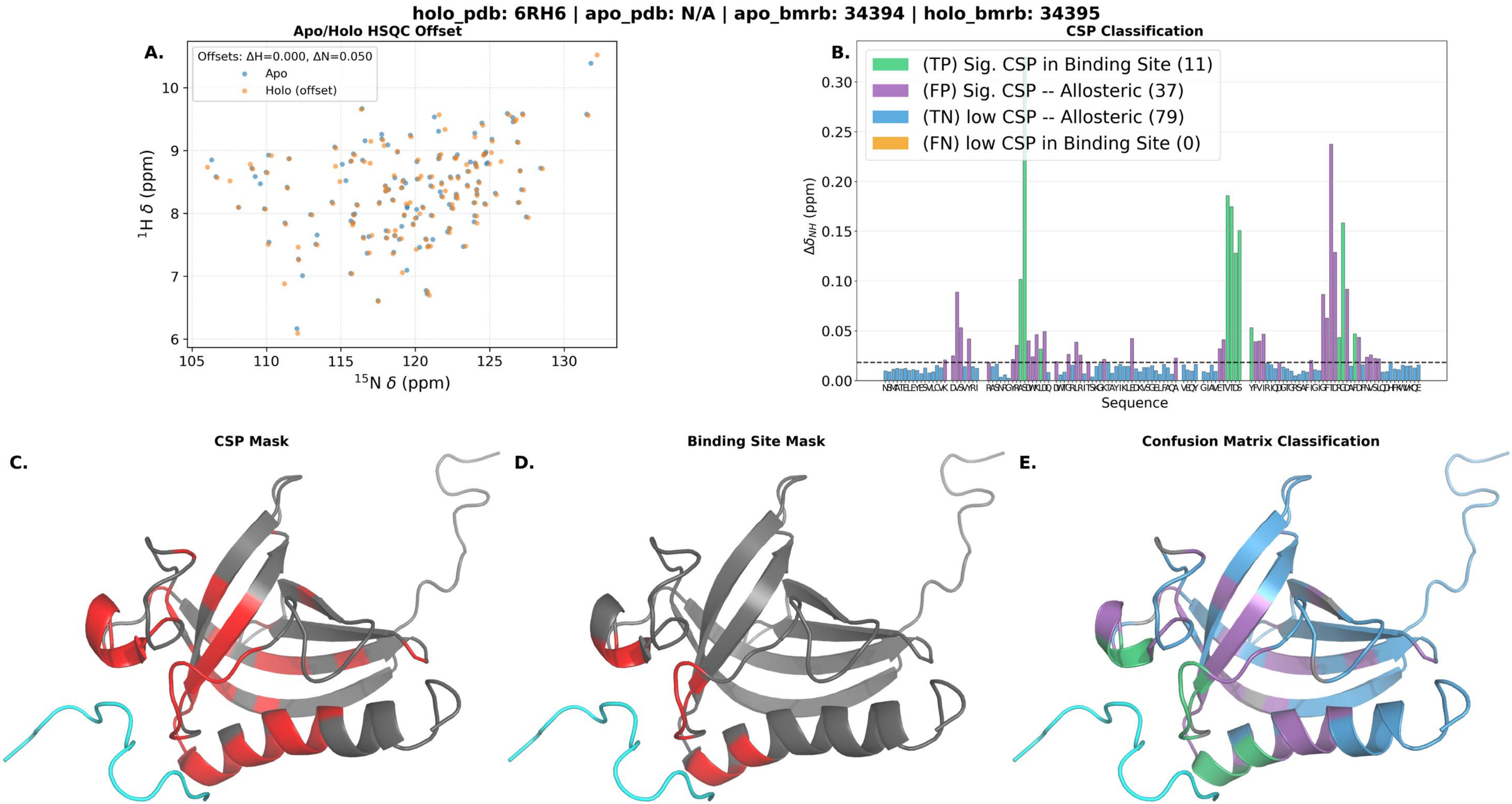

holo_pdb: 6RH6 | apo_pdb: N/A | apo_bmrb: 34394 | holo_bmrb: 34395
A.
Apo/Holo HSQC Offset
Offsets: ΔH=0.000, ΔN=0.050
Apo
Holo (offset)
1H δ (ppm)
15N δ (ppm)
B.
CSP Classification
(TP) Sig. CSP in Binding Site (11)
(FP) Sig. CSP -- Allosteric (37)
(TN) low CSP -- Allosteric (79)
(FN) low CSP in Binding Site (0)
Δδ_NH (ppm)
Sequence
C.
CSP Mask
D.
Binding Site Mask
E.
Confusion Matrix Classification

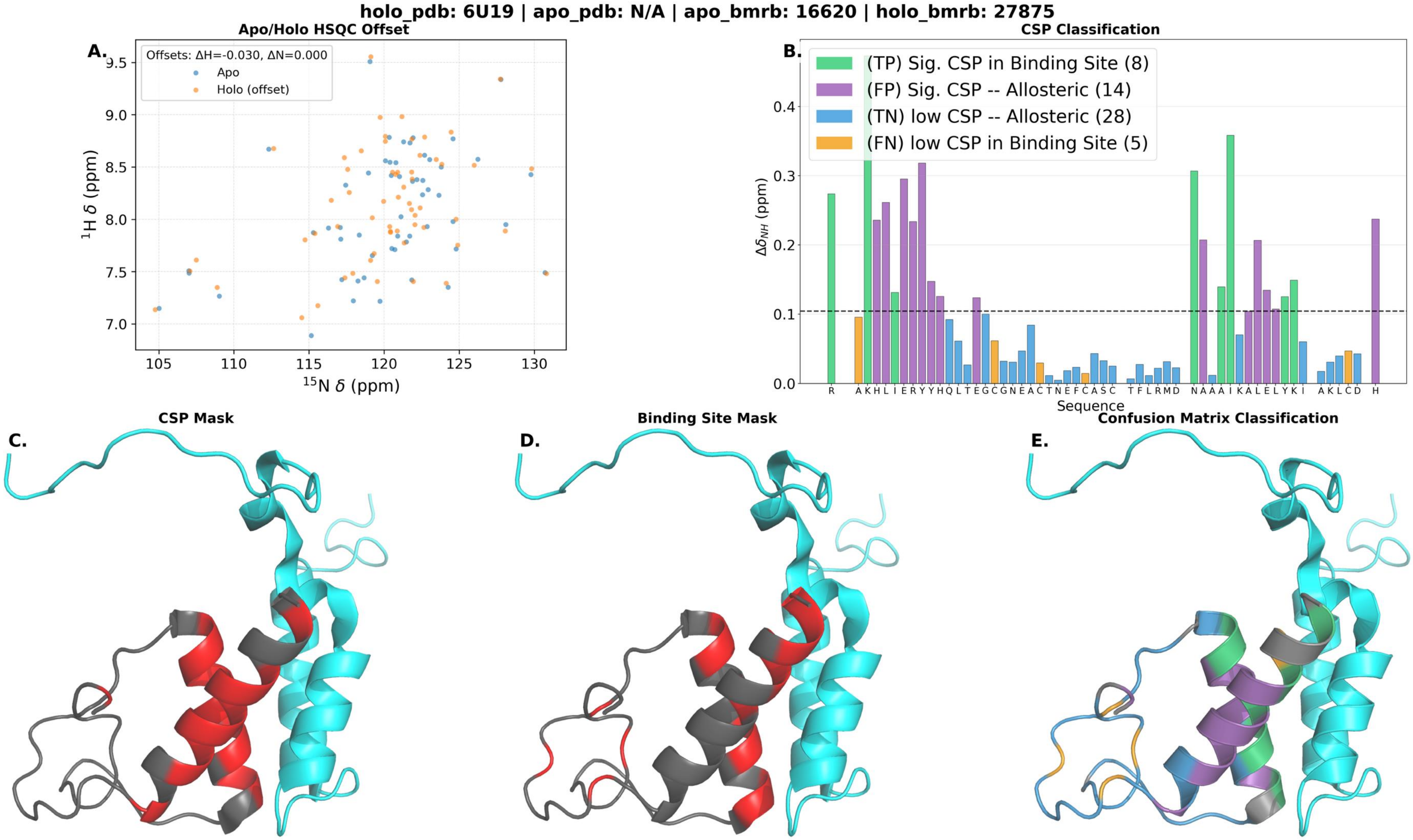

holo_pdb: 6U19 | apo_pdb: N/A | apo_bmrb: 16620 | holo_bmrb: 27875
Apo/Holo HSQC Offset
A.
Offsets: ΔH=-0.030, ΔN=0.000
Apo
Holo (offset)
¹H δ (ppm)
¹⁵N δ (ppm)
CSP Classification
B.
(TP) Sig. CSP in Binding Site (8)
(FP) Sig. CSP -- Allosteric (14)
(TN) low CSP -- Allosteric (28)
(FN) low CSP in Binding Site (5)
Δδ_NH (ppm)
Sequence
CSP Mask
C.
Binding Site Mask
D.
Confusion Matrix Classification
E.

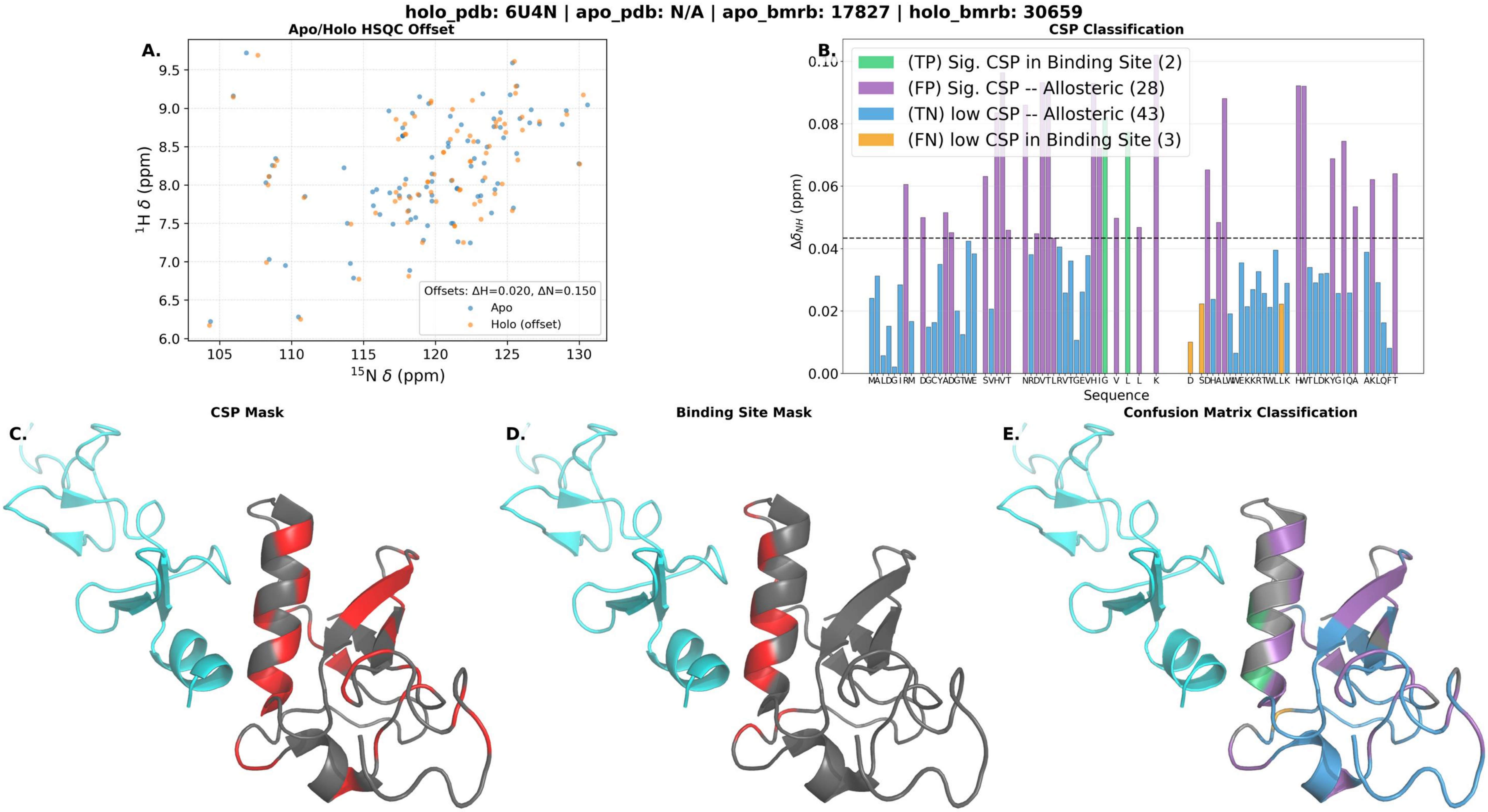

holo_pdb: 6U4N | apo_pdb: N/A | apo_bmrb: 17827 | holo_bmrb: 30659
Apo/Holo HSQC Offset
A.
Offsets: ΔH=0.020, ΔN=0.150
Apo
Holo (offset)
¹H δ (ppm)
¹⁵N δ (ppm)
CSP Classification
B.
(TP) Sig. CSP in Binding Site (2)
(FP) Sig. CSP -- Allosteric (28)
(TN) low CSP -- Allosteric (43)
(FN) low CSP in Binding Site (3)
Δδ_NH (ppm)
Sequence
CSP Mask
C.
Binding Site Mask
D.
Confusion Matrix Classification
E.

**holo_pdb: 7JQ8 | apo_pdb: 7JMY | apo_bmrb: 30782 | holo_bmrb: 30786**

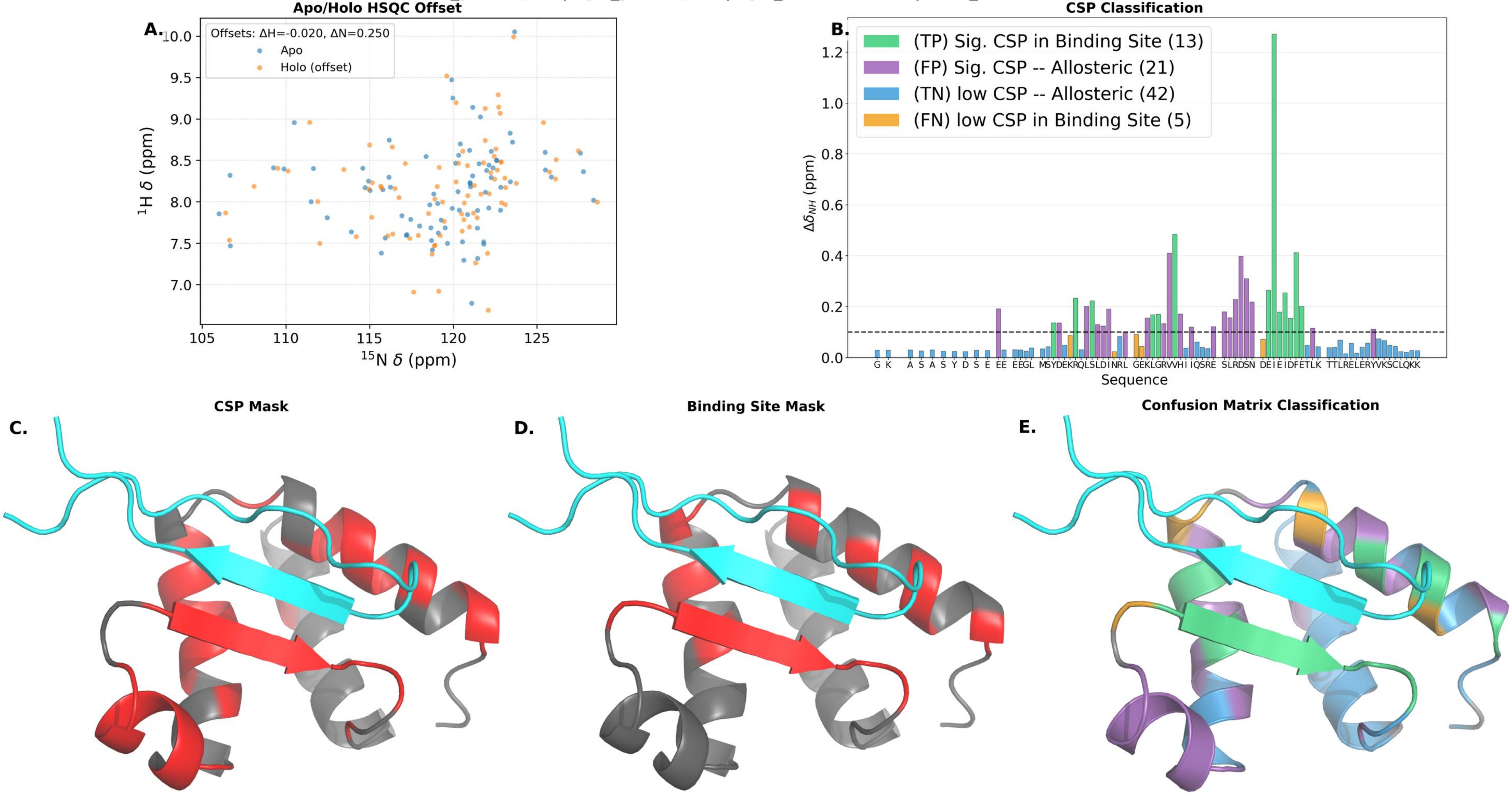

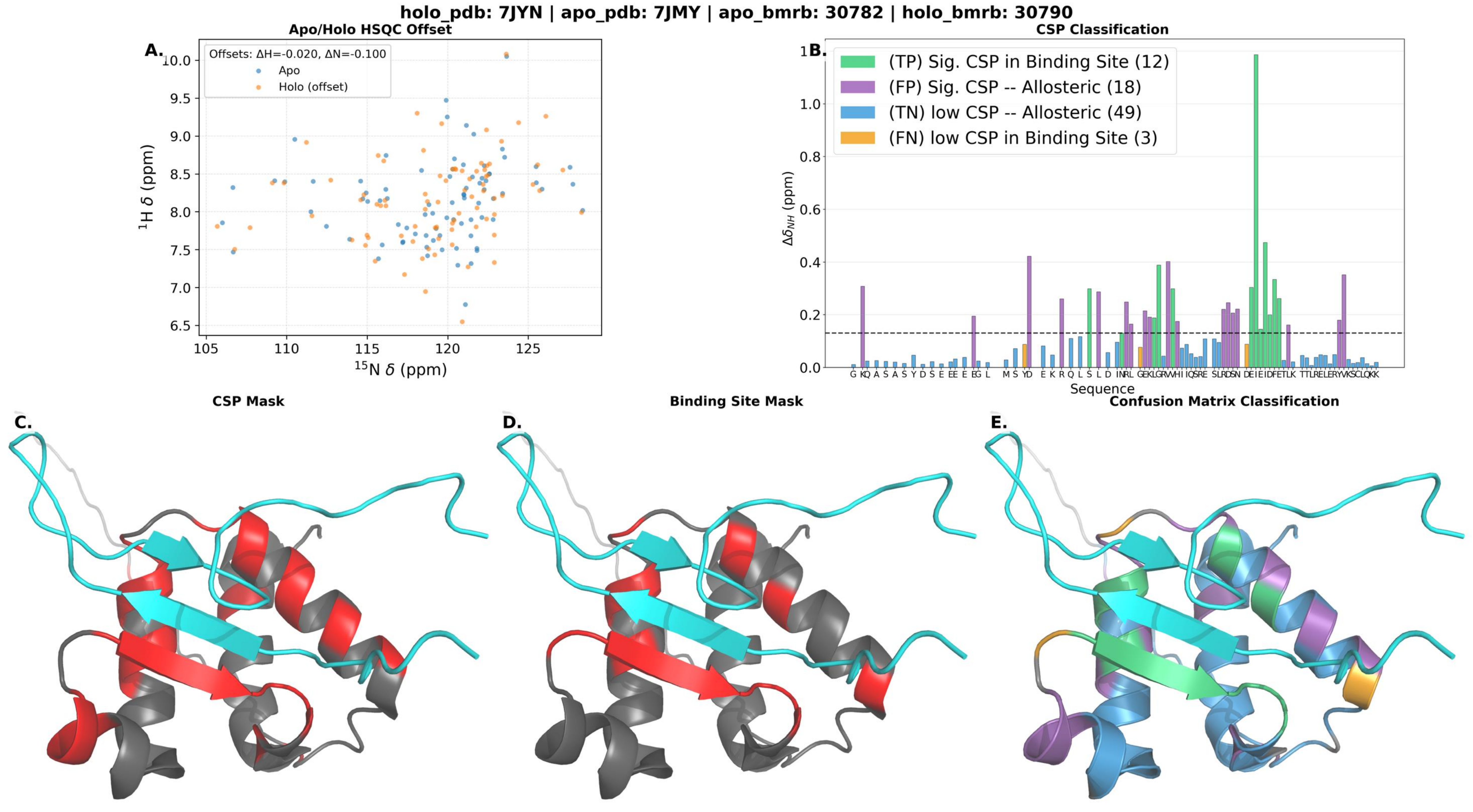

holo_pdb: 7JYN | apo_pdb: 7JMY | apo_bmrb: 30782 | holo_bmrb: 30790
A.
Apo/Holo HSQC Offset
Offsets: ΔH=-0.020, ΔN=-0.100
Apo
Holo (offset)
¹H δ (ppm)
¹⁵N δ (ppm)
B.
CSP Classification
(TP) Sig. CSP in Binding Site (12)
(FP) Sig. CSP -- Allosteric (18)
(TN) low CSP -- Allosteric (49)
(FN) low CSP in Binding Site (3)
Δδ_NH (ppm)
Sequence
C.
CSP Mask
D.
Binding Site Mask
E.
Confusion Matrix Classification

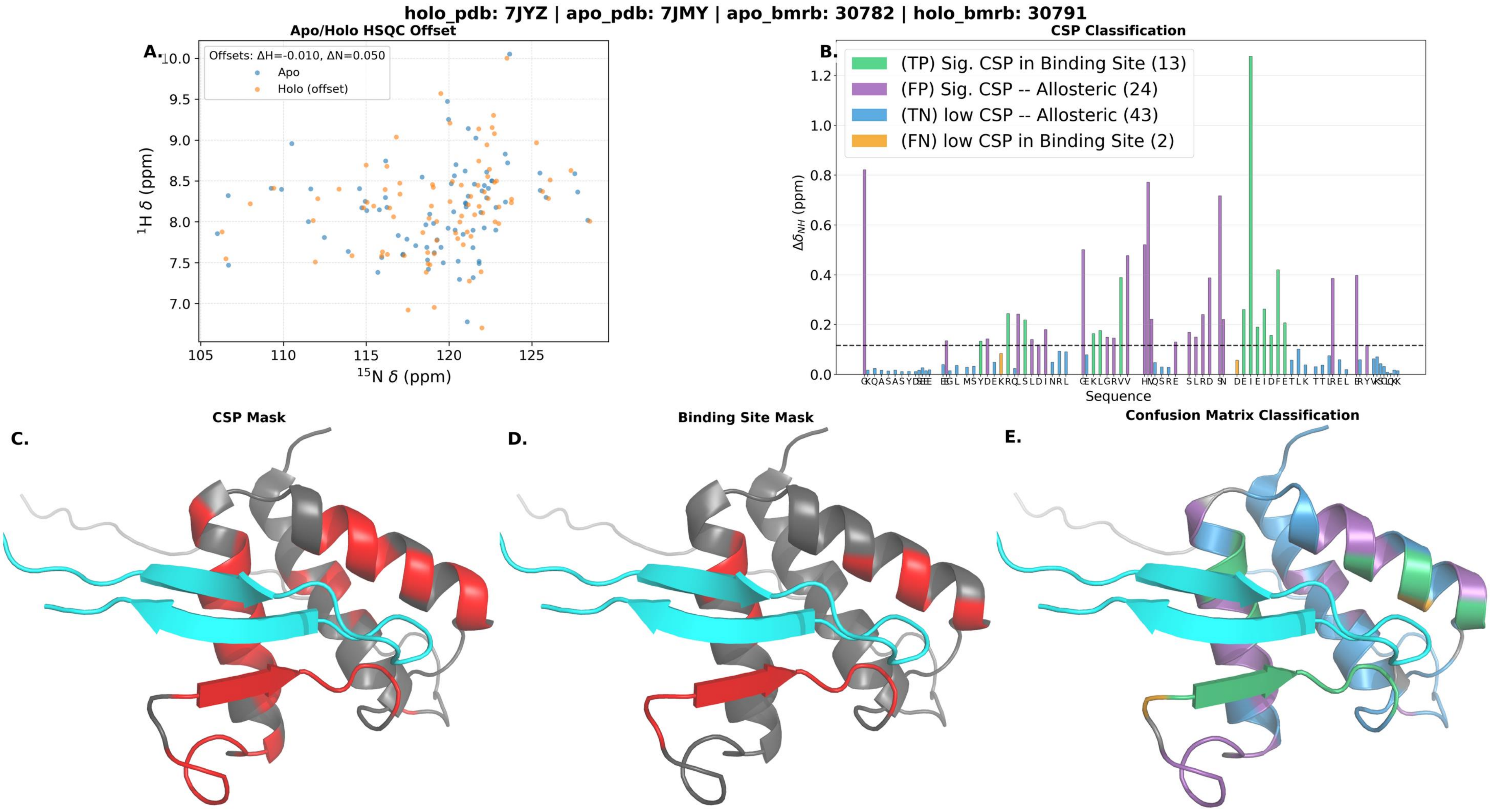
holo_pdb: 7JYZ | apo_pdb: 7JMY | apo_bmrb: 30782 | holo_bmrb: 30791
A.
Apo/Holo HSQC Offset
Offsets: ΔH=-0.010, ΔN=0.050
Apo
Holo (offset)
1H δ (ppm)
15N δ (ppm)
B.
CSP Classification
(TP) Sig. CSP in Binding Site (13)
(FP) Sig. CSP -- Allosteric (24)
(TN) low CSP -- Allosteric (43)
(FN) low CSP in Binding Site (2)
Δδ_NH (ppm)
Sequence
C.
CSP Mask
D.
Binding Site Mask
E.
Confusion Matrix Classification

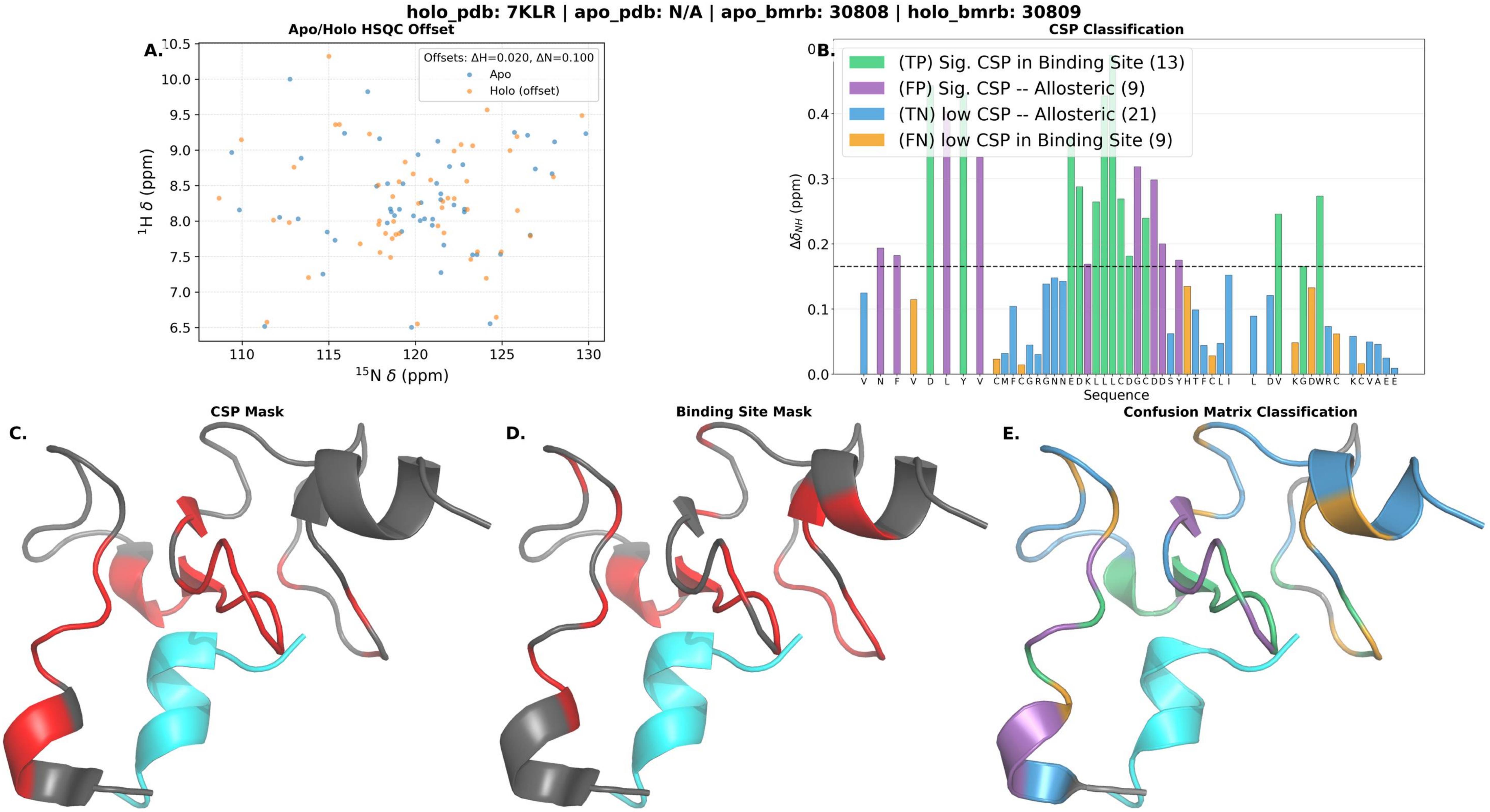
holo_pdb: 7KLR | apo_pdb: N/A | apo_bmrb: 30808 | holo_bmrb: 30809
A.
Apo/Holo HSQC Offset
Offsets: ΔH=0.020, ΔN=0.100
Apo
Holo (offset)
$^{1}$H δ (ppm)
$^{15}$N δ (ppm)
B.
CSP Classification
(TP) Sig. CSP in Binding Site (13)
(FP) Sig. CSP -- Allosteric (9)
(TN) low CSP -- Allosteric (21)
(FN) low CSP in Binding Site (9)
$\Delta\delta_{NH}$ (ppm)
Sequence
V N F V D L Y V CMFCGRGNNEDKLLLLCDGCDDSYHTFCLI L DV KGDWRC KCVAEE
C.
CSP Mask
D.
Binding Site Mask
E.
Confusion Matrix Classification

**holo_pdb: 7PKU | apo_pdb: N/A | apo_bmrb: 50446 | holo_bmrb: 34661**

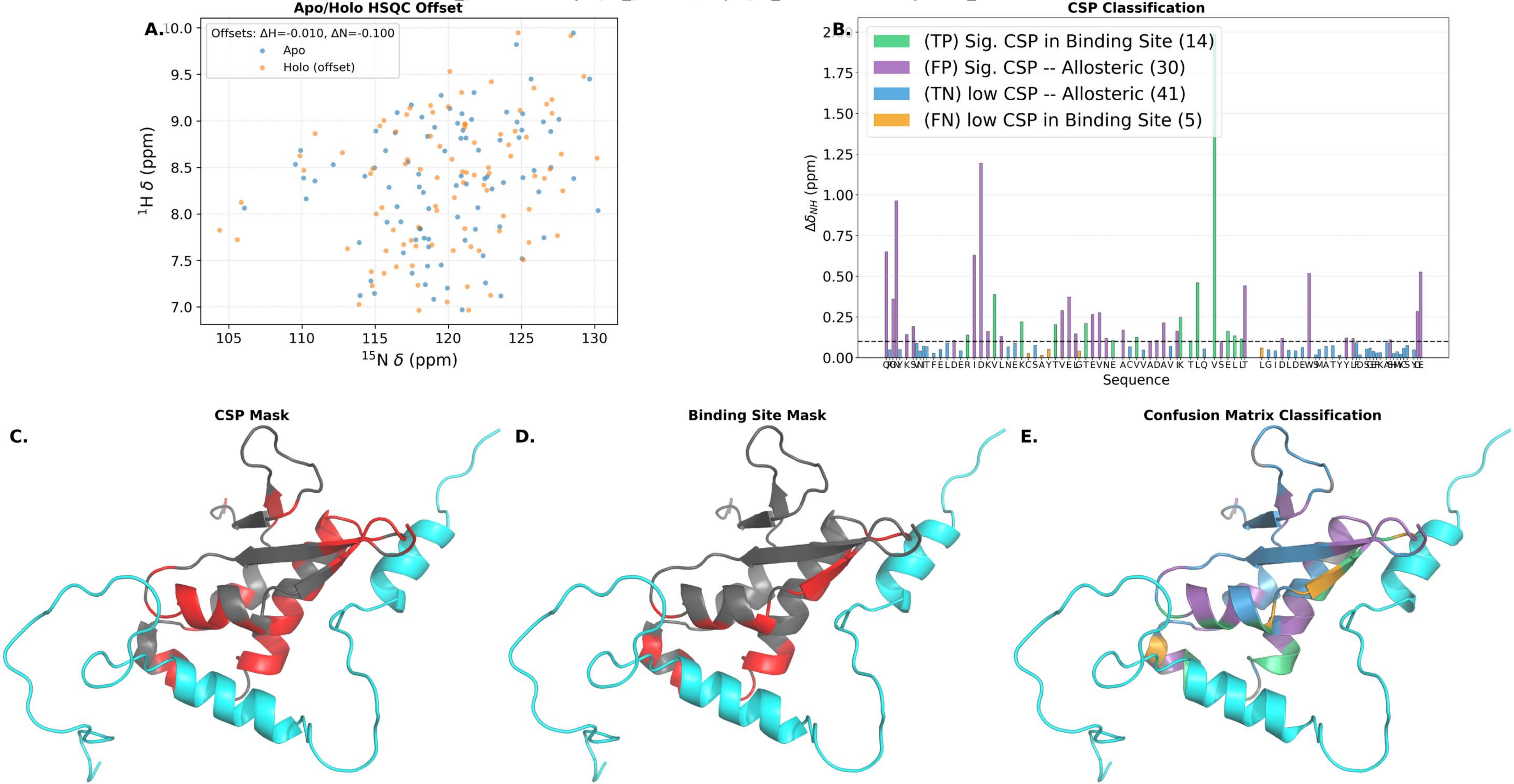

**holo_pdb: 7S5J | apo_pdb: N/A | apo_bmrb: 30926 | holo_bmrb: 30950**

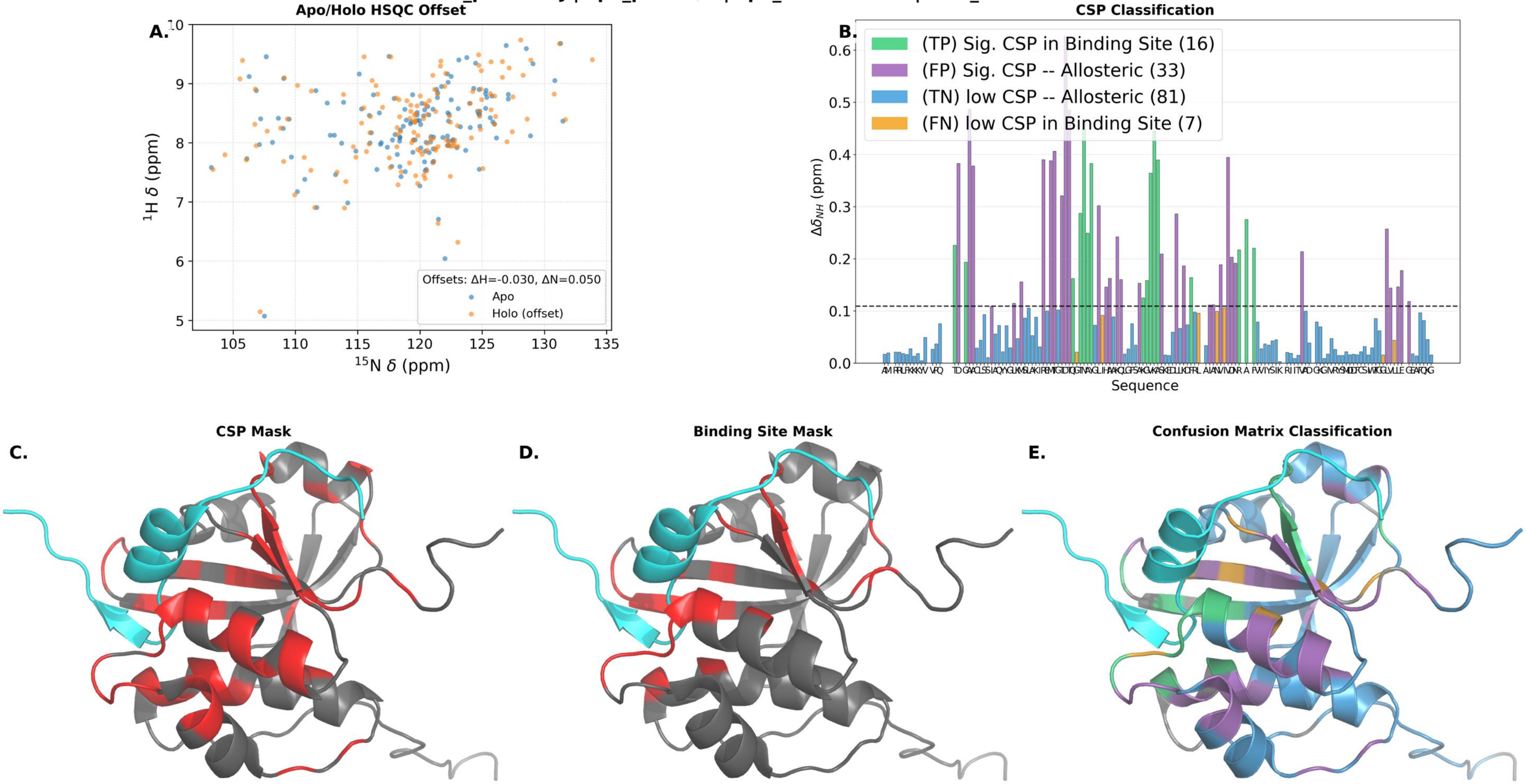

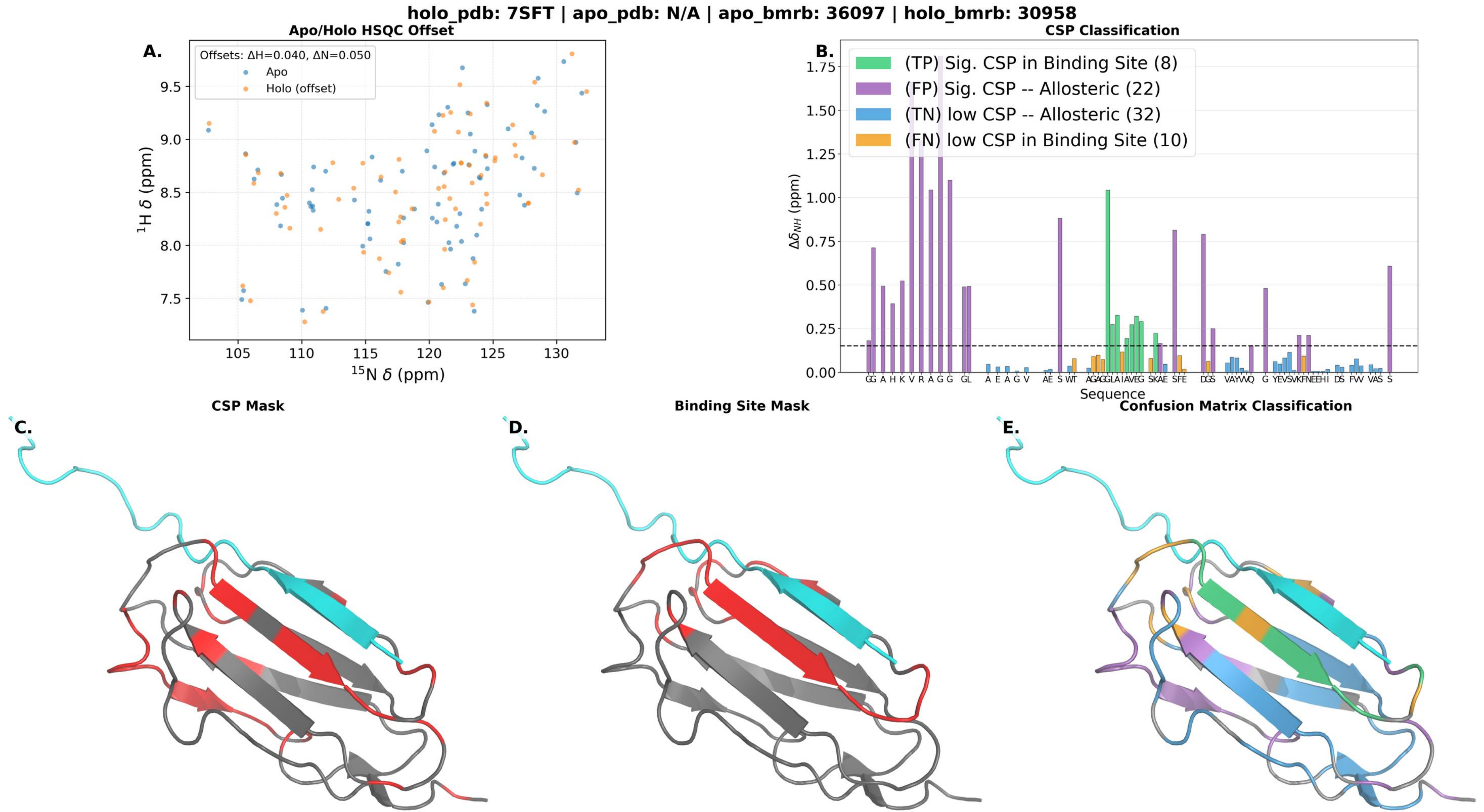

holo_pdb: 7SFT | apo_pdb: N/A | apo_bmrb: 36097 | holo_bmrb: 30958
Apo/Holo HSQC Offset
A.
Offsets: ΔH=0.040, ΔN=0.050
Apo
Holo (offset)
$^{1}$H δ (ppm)
$^{15}$N δ (ppm)
CSP Classification
B.
(TP) Sig. CSP in Binding Site (8)
(FP) Sig. CSP -- Allosteric (22)
(TN) low CSP -- Allosteric (32)
(FN) low CSP in Binding Site (10)
$\Delta\delta_{NH}$ (ppm)
Sequence
CSP Mask
C.
Binding Site Mask
D.
Confusion Matrix Classification
E.

**holo_pdb: 7X5C | apo_pdb: N/A | apo_bmrb: 51441 | holo_bmrb: 36473**

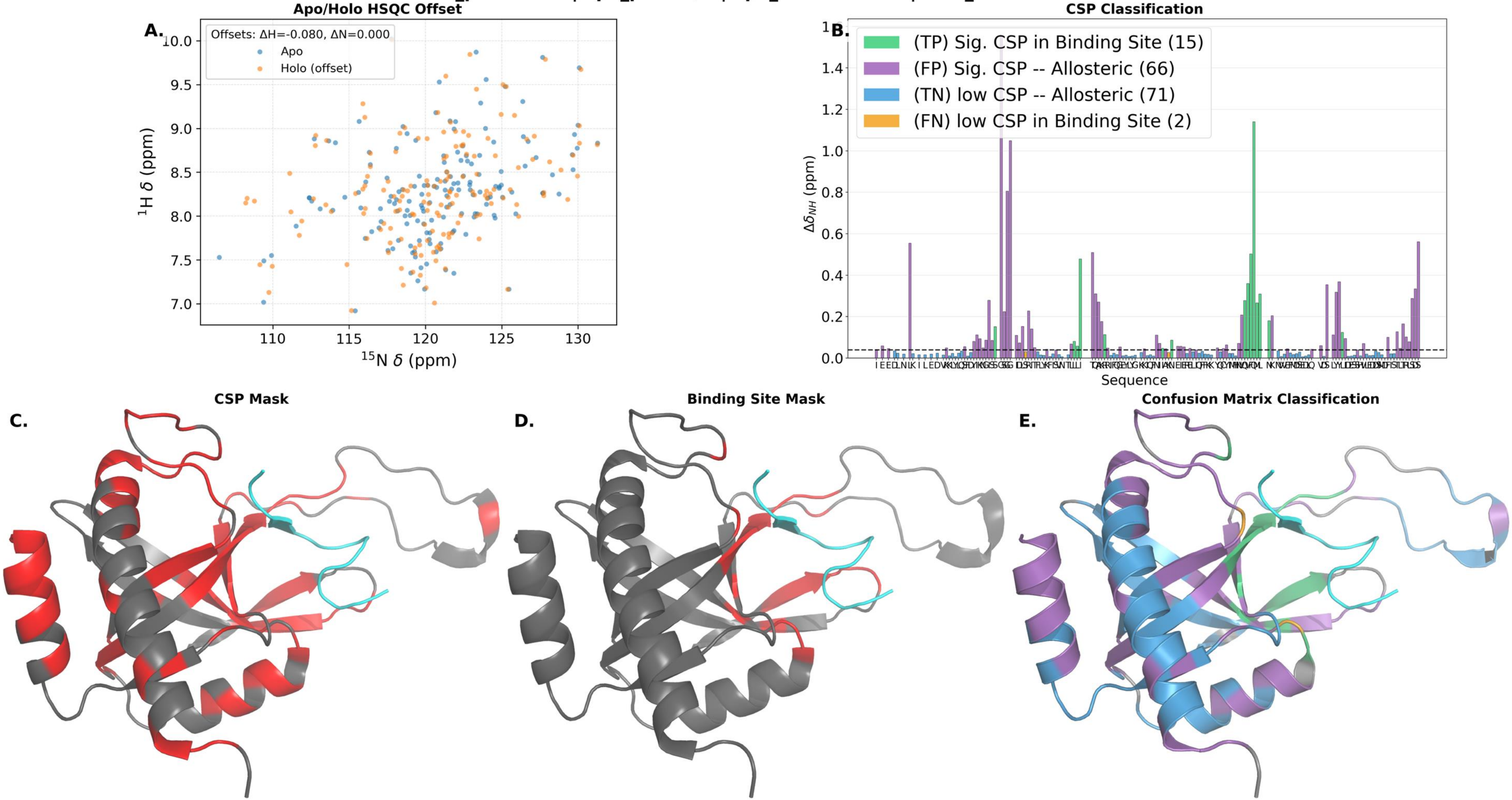

holo_pdb: 8SG2 | apo_pdb: N/A | apo_bmrb: 27579 | holo_bmrb: 31080

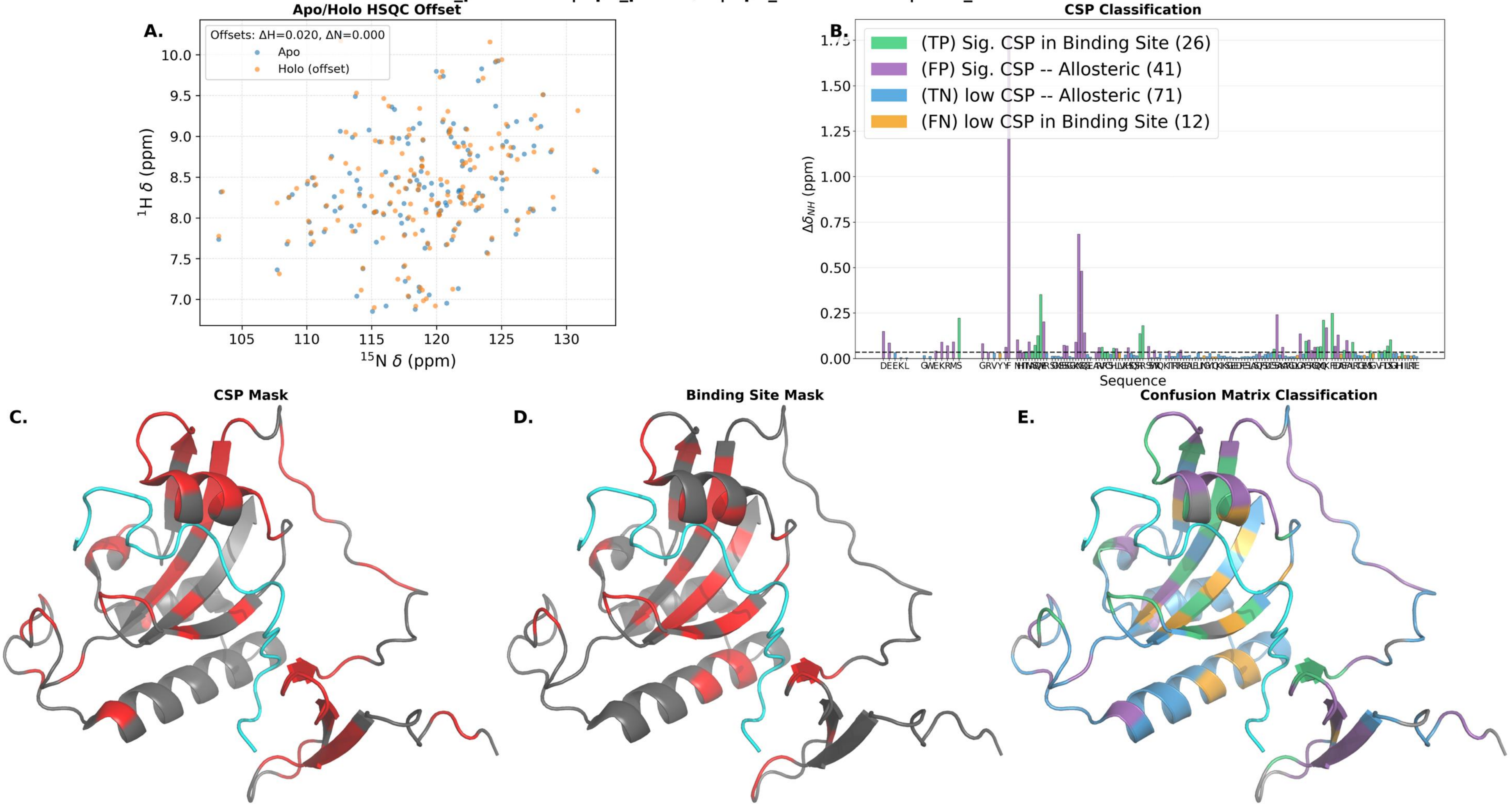

holo_pdb: 9C5E | apo_pdb: N/A | apo_bmrb: 18642 | holo_bmrb: 31177

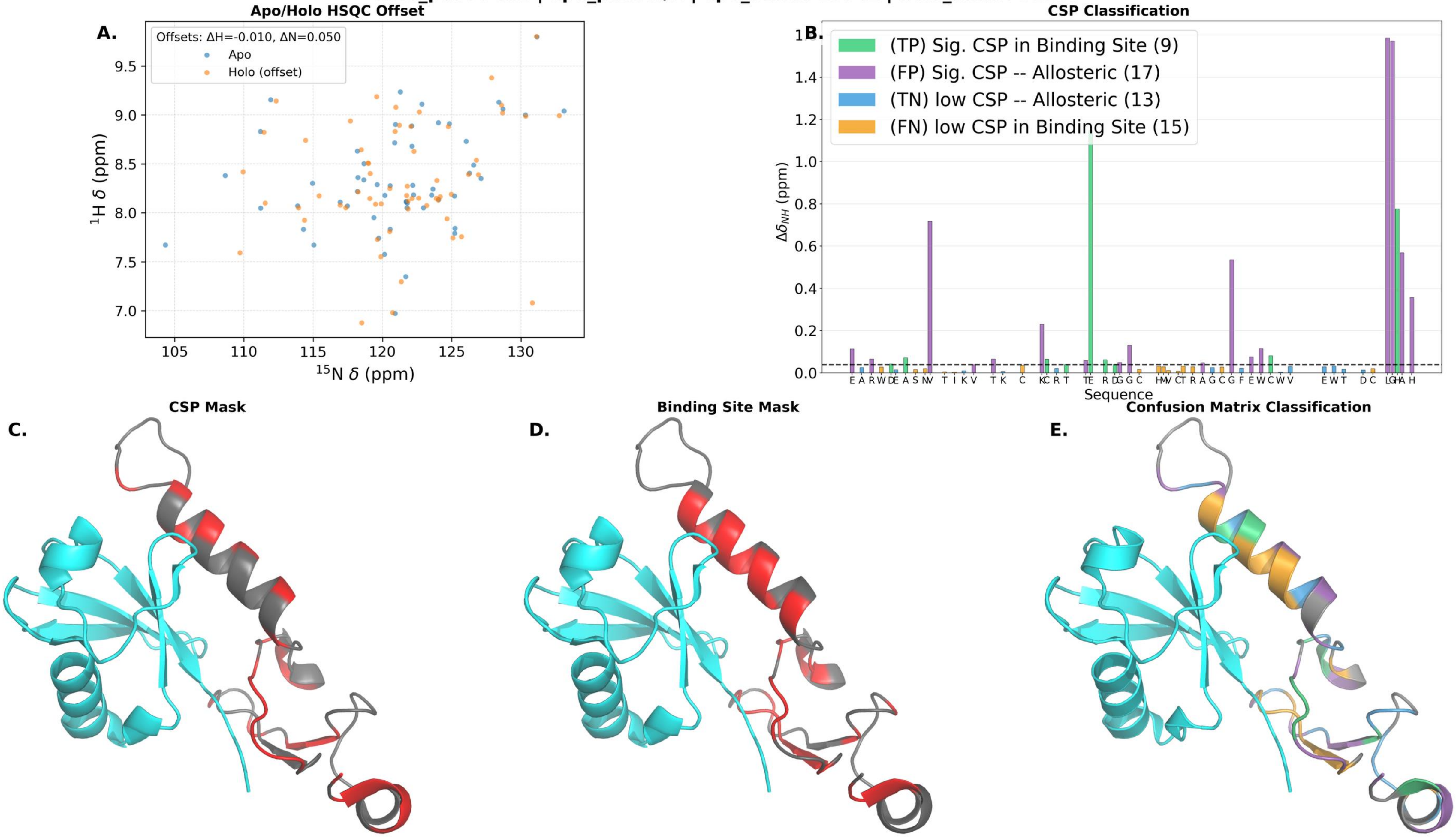

holo_pdb: 9QLM | apo_pdb: N/A | apo_bmrb: 15670 | holo_bmrb: 34986

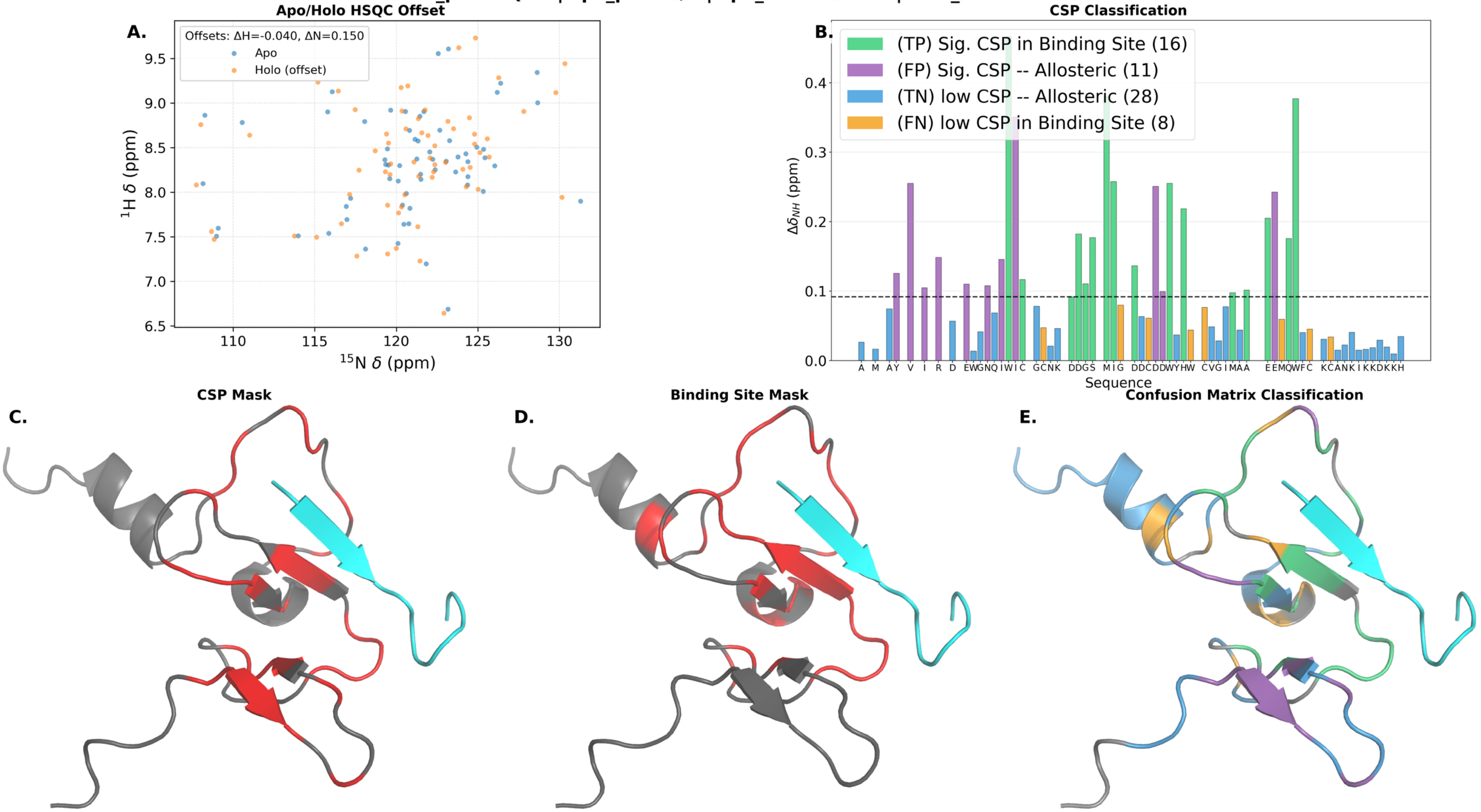

**holo_pdb: 9R3Y | apo_pdb: N/A | apo_bmrb: 26643 | holo_bmrb: 34992**

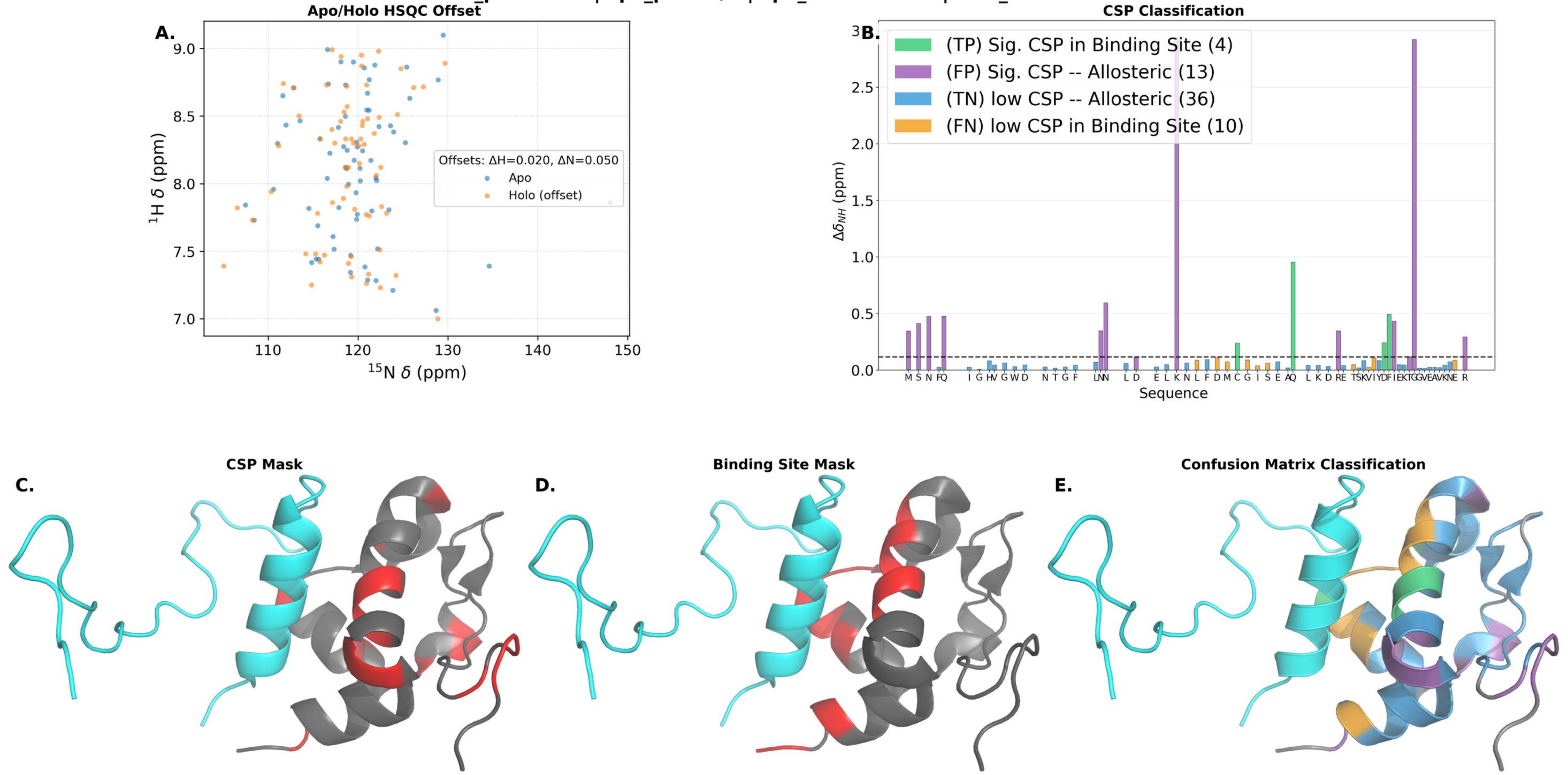
